# Allosteric regulation of 3CL protease of SARS-CoV-2 and SARS-CoV observed in the crystal structure ensemble

**DOI:** 10.1101/2021.08.29.458083

**Authors:** Akinori Kidera, Kei Moritsugu, Toru Ekimoto, Mitsunori Ikeguchi

## Abstract

The 3C-like protease (3CL^pro^) of SARS-CoV-2 is a potential therapeutic target for COVID-19. Importantly, it has an abundance of structural information solved as a complex with various drug candidate compounds. Collecting these crystal structures (83 Protein Data Bank (PDB) entries) together with those of the highly homologous 3CL^pro^ of SARS-CoV (101 PDB entries), we constructed the crystal structure ensemble of 3CL^pro^ to analyze the dynamic regulation of its catalytic function. The structural dynamics of the 3CL^pro^ dimer observed in the ensemble were characterized by the motions of four separate loops (the C-loop, E-loop, H-loop, and Linker) and the C-terminal domain III on the rigid core of the chymotrypsin fold. Among the four moving loops, the C-loop (also known as the oxyanion binding loop) causes the order (active)–disorder (collapsed) transition, which is regulated cooperatively by five hydrogen bonds made with the surrounding residues. The C-loop, E-loop, and Linker constitute the major ligand binding sites, which consist of a limited variety of binding residues including the substrate binding subsites. Ligand binding causes a ligand size dependent conformational change to the E-loop and Linker, which further stabilize the C-loop via the hydrogen bond between the C-loop and E-loop. The T285A mutation from SARS-CoV 3CL^pro^ to SARS-CoV-2 3CL^pro^ significantly closes the interface of the domain III dimer and allosterically stabilizes the active conformation of the C-loop via hydrogen bonds with Ser1 and Gly2; thus, SARS-CoV-2 3CL^pro^ seems to have increased activity relative to that of SARS-CoV 3CL^pro^.

## Introduction

During structure-based drug discovery, the three-dimensional structure of a therapeutic target protein is often determined redundantly in the form of a complex with various drug candidates, which reveals their binding modes at the atomic level [1–4]. Consequently, an abundance of structural information has been accumulated for some target proteins in Protein Data Bank (PDB) and other databases [5–10]. Ligand interactions can alter the structure of a receptor protein in different ways to produce structural variation in the protein [11, 12] When examining protein structures, it becomes clear that ligand binding is not the sole cause of structural variation in crystals: crystal packing and sequence alteration also affect structure [13, 14]. Within a set of PDB entries for a protein, different space groups or different crystal packing are frequently found. Crystallographic experiments often use proteins with altered sequences either for mutagenesis experiments or in sample preparation. These factors generate the structural ensemble of crystal structures, which is termed a “crystal structure ensemble.”

According to the concept of protein dynamics, conformational selection [15], and linear response theory [16], any structural changes of a protein, whether they occur naturally or artificially, are reflections of its intrinsic dynamics. Therefore, the crystal structure ensemble can be understood as a sampled subset of the true structural ensemble, although the samples may be limited and biased to some extent because they are not randomly sampled. In respect of the ensemble’s reliability, the structural variations in the ensemble are statistically significant observations derived from repeated measurements. For these reasons, the crystal structure ensemble is an important source for the study of protein dynamics [17].

In the present study, we assembled a crystal structure ensemble of 3C-like protease (3CL^pro^, also known as main protease) from severe acute respiratory syndrome coronavirus 2 (SARS-CoV-2) as well as the highly homologous 3CL^pro^ of SARS-CoV, the etiological agent for SARS in 2002 (Fig. 1). 3CL^pro^ is a cysteine protease belonging to the C30 family (coronavirus 3C-like proteases) of the PA clan (serine/cysteine proteases with a chymotrypsin fold) [7]; it plays a crucial role in viral replication by cleaving the polyprotein to release functional proteins [18–20]. The COVID-19 pandemic caused by SARS-CoV-2 poses an urgent challenge for the development of antiviral agents [21–23]. The inhibitor of 3CL^pro^ is a potential candidate as an antiviral agent. The crystal structures of 3CL^pro^ complexed with drug candidate molecules have been solved intensively and accumulated in PDB since the first structure was released in February 2020 (PDB:6lu7 [24]; Supporting data S1 and S2). Here, we also utilize the abundant structure data of 3CL^pro^ from SARS-CoV because of its 96% sequence identity and superimposable structures (Fig. 1). The compiled data in this study are of version 10/25/2020, which contains SARS-CoV-2 3CL^pro^ (83 entries; 113 independent chains) and SARS-CoV 3CL^pro^ (101 entries; 145 independent chains) (Supporting data S1 and S2; see Data used in this study in Materials and Methods for details).

**Figure 1.**
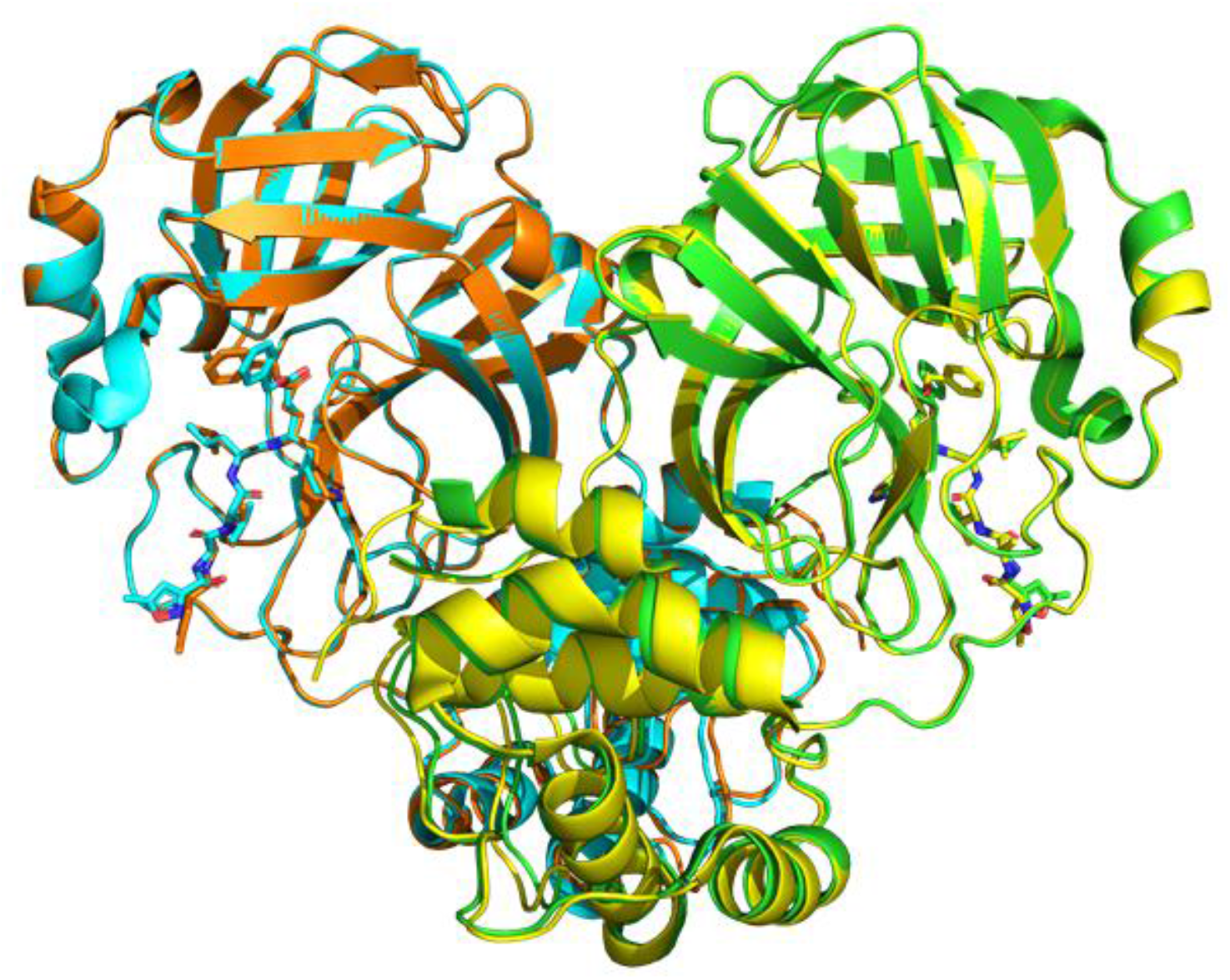
Illustrations of SARS-CoV-2 3CL^pro^ (PDB:7bqy, Chain A (green) and Chain B (cyan)) and SARS-CoV 3CL^pro^ (PDB:2hob, Chain A (yellow) and Chain B (orange)), superimposed at the core region of the chymotrypsin fold. Both have homodimeric structures of C2 symmetry complexed with N3 (stick; a peptide-mimic compound), which are superimposable (Cα RMSD = 0.68 Å), although the terminal moieties of N3 have a deviation. The upper domain is the chymotrypsin fold named domains I and II, whereas the lower helical domain is domain III.

3CL^pro^ is known for its highly dynamic nature. The function of 3CL^pro^ is regulated by the structural transition between the active and collapsed states of the catalytic loop (termed the “C-loop” hereafter) [25]. The activation requires dimerization through the additional C-terminal helical domain (domain III) [26]. The ligand binding sites of 3CL^pro^ are located in flexible loops [7]. These dynamical behaviors were investigated in detail based on the crystal structure ensemble. The results of each entry are summarized in Supporting data S1 and S2 for the ligand-bound and ligand-free chains, respectively. First, the overall dynamics is explored to identify the elements of the dynamic structure. Second, the dynamic regulation of the catalytic function in the C-loop is studied. Third, the ligand binding sites and the influence of ligand binding on the protein structure are investigated, and the ligand-induced activation is explained. Finally, the effects of the mutation between the two 3CL^pro^ species on the dimeric structure and catalytic function are examined.

## Results and Discussion

### Overall motions in the crystal structure ensemble of 3CL^pro^

To understand the dynamic regulation of the catalytic function of 3CL^pro^, the crystal structure ensemble was analyzed according to the following two steps (see Materials and Methods for more details). As the first step, the elements of motion regulating the catalytic function were identified by applying the Motion Tree to the crystal structural ensemble [27, 28], which is a hierarchical cluster analysis of the variance of interresidue distances (therefore, it is superposition free). In the Motion Tree based on all 258 independent chains, clusters are defined as groups of residues moving cooperatively, represented by branches of the Motion Tree; thus, they are termed “moving clusters,” in which the residues move as one group (Fig. 2A). The Motion Tree delineates the dynamics of 3CL^pro^ as a set of moving clusters consisting of four flexible loops and domain III situated on the rigid core of the chymotrypsin fold (Fig. 2B). The moving clusters are as follows: the “C-loop” (residues 138–143; the catalysis-related loop, also known as the L1 or oxyanion binding loop [29]; the target fragment for functional regulation), “E-loop” (residues 166–178; loop starting at E166, L2), “H-loop” (residues 38–64; a mostly helical loop), “Linker” (residues 188–196; a <u>linker</u> to domain III, L3), and “domain III” (residues 200–300). All of these clusters appear to move independently of each other (Table S1). As detailed below, three of the four loops, namely the C-loop, E-loop, and Linker, contain major ligand binding sites. Domain III has been reported to play a crucial role in the dimerization required for the activation [25, 26] and is discussed below in relation to mutation effects and the allosteric regulation of catalytic function. The rigid core consists of the core part of domain I and II (residues 13–17, 33–37, 71–111, 129–137, 154–165, 179–187, and 197–199; chymotrypsin fold) and the dimer interface region (residues 3–12, 18–32, 65–70, 112–128, and 144–153) including the N-finger (residues 1–7) of the essential structural component for dimerization and activity [25, 30–32], whereas residues 1 and 2 are highly flexible and play an important role in the functional regulation.

**Figure 2.**
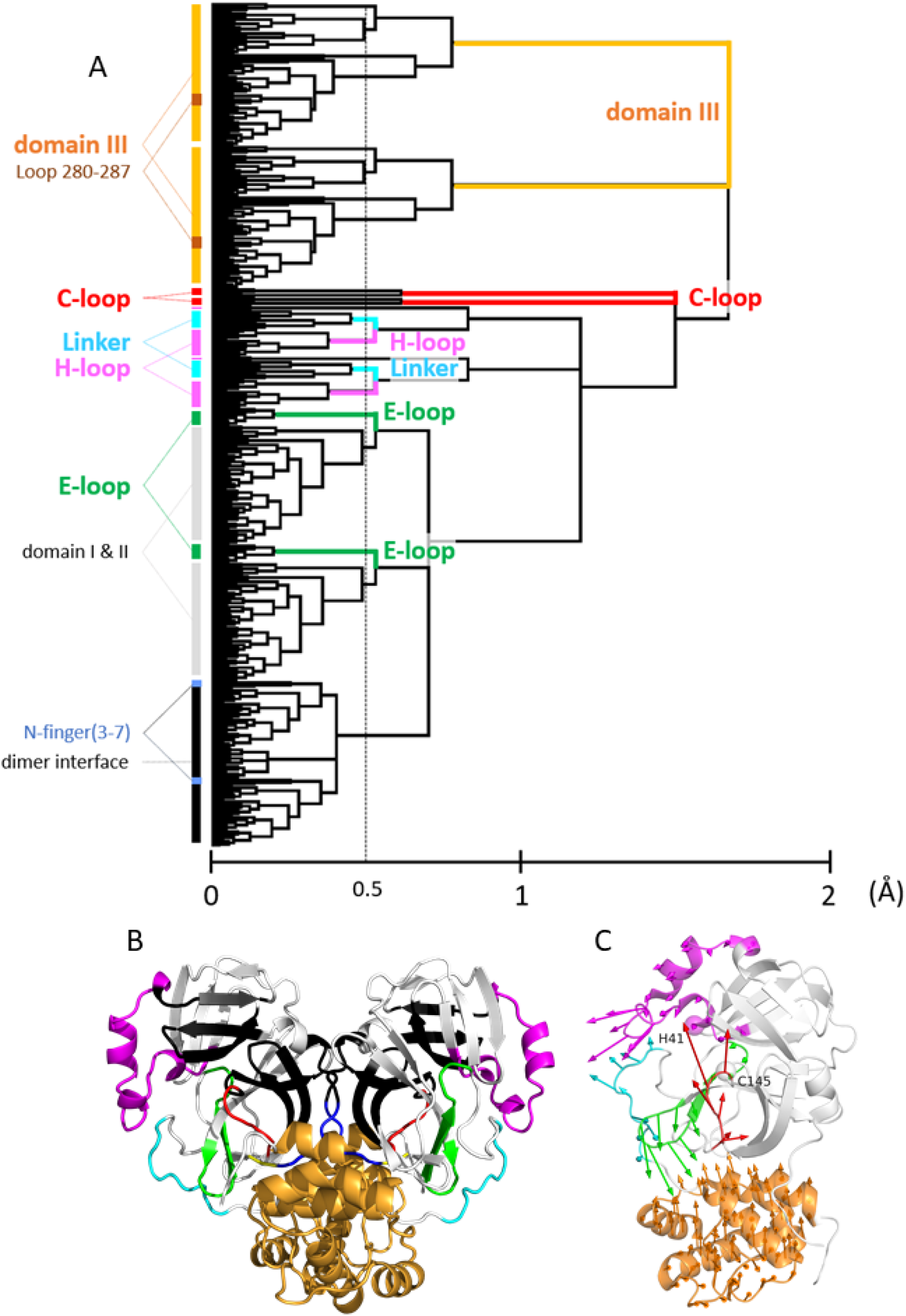
(A) Motion Tree, a hierarchical clustering of the variance of Cα distances [27, 28], calculated for the crystal structure ensemble consisting of all 258 chains of SARS-CoV 3CL^pro^ and SARS-CoV-2 3CL^pro^. The variances *D_mn_* were symmetrized for two chains, or *D_mn_* = *D_MN_* (residues *m* and *n* in a protomer correspond to residues *M* and *N* in the other protomer, respectively), to treat the homodimer of 3CL^pro^; thus, two sets of equivalent clusters corresponding to each protomer exist in the tree (see Methods). In asymmetric dimers, the average of the two distances was used. A taller branch with a high Motion Tree (MT) score, the scale of the abscissa, indicates a cluster moving to a greater extent. The amplitudes of motions follow the order: domain III > C-loop > E-loop ~ H-loop ~ Linker. The clusters over the threshold (a thin broken vertical line; MT score = 0.5) are defined as the moving clusters. Nevertheless, domain III is treated as a single cluster for simplicity. Based on the Motion Tree, we defined five moving clusters (the C-loop, E-loop, H-loop, Linker, and domain III). The N-finger (residues 3–7) is shown to be a part of the core of the dimer interface region. Highly mobile Ser1 and Gly2 and the C-terminal residues 301–306 are not included in the analysis because their coordinates are often missing. Loop 280–287 belongs to domain III (see discussion in “Influences of T285A mutation …”). (b) The moving clusters of SARS-CoV-2 3CL^pro^ (PDB: 6lu7 as a representative). The color scheme for the moving clusters defined by the Motion Tree (Fig. 2A) is as follows: the C-loop, red; E-loop, green; H-loop, magenta; Linker, cyan; and domain III, bright orange. The core regions consist of the core part of domains I and II (white) and the dimer interface region (black). The C-terminal part of the N-finger (residues 3–7; blue) is within the dimer interface region, whereas residues 1 and 2 (yellow) are highly flexible. Notably, each of the N-termini is located near the C-loop of the other protomer. (C) The first principal components (PCs) of the moving clusters within a protomer are illustrated by arrows using the same color scheme used in (B). The variances explained by PC1 are 0.59 (C-loop), 0.58 (E-loop), 0.35 (H-loop), 0.41 (Linker), and 0.53 (domain III). The catalytic dyad, Cys145 and His41, is depicted by spheres at Cα atoms.

As the second step, the motions of each moving cluster were delineated by calculating the principal components (PCs) of the motions relative to the core region for the four loops and domain III (Fig. 2C). Domain III was treated in two ways, one as the motion within a protomer (Fig. 2C) and the other as the motion of the dimer (see Fig. S1 and Supporting Text 1 and 2 for details). The motion of domain III within a protomer is a typical domain motion against the core region, as DynDom, a domain motion analysis program, identifies it as the domain motion [33].

### C-loop: the switch controlling catalytic activity and the target of regulation

#### Hydrogen bond between Gly143 and Asn28 regulates the formation of the oxyanion hole: a marker distinguishing active and collapsed states

The conformational change of the C-loop (residues 138–143) between the active and collapsed states switches on and off the catalytic activity of 3CL^pro^ by forming and collapsing the oxyanion hole stabilizing the catalytic intermediate [25, 34]. This activation mechanism of the C-loop conformational change is largely identical to that of zymogen activation in serine proteases, such as chymotrypsinogen (inactive/collapsed) to chymotrypsin (active) [35], although the activation of 3CL^pro^ also requires dimerization. The representative active and collapsed structures are shown in Fig. 3A. The comparison of these structures suggests that the hydrogen bond (HB) between Gly143 and Asn28 is a sensitive signature of the oxyanion hole (hydrogen bonds are defined by LigPlot [36]). In the active state, HB 143-28 together with HB 145-28 correctly arranges the main-chain NH groups of Gly143 and Cys145 to form the oxyanion hole. When the C-loop is collapsed or Gly143 moves downward as shown in the right panel of Fig. 3A (PC1(C) in Fig. 2C is drawn conversely as the activation process), HB 143-28 is cleaved but HB 145-28 remains intact. Consequently, Gly143 is separated from Cys145 to break down the oxyanion hole. As shown below, the formation of HB 143-28 is highly correlated with the C-loop conformation. Contrastingly, HB 145-28 exists in almost all chains (257 of 258 chains), with the exception of the mutant N28A (PDB:3fzd [37]), as both Cys145 and Asn28 belong to the stable dimer interface region. The importance of Asn28 is confirmed by the mutant N28A, in which the activity is completely lost and the affinity for dimerization is largely impaired [37]. Therefore, in the present study, we define the conformational states of the C-loop by HB 143-28: the active state occurs when HB 143-28 exists, whereas the collapsed state occurs when HB 143-28 is absent. The role of the two hydrogen bonds, HB 143-28 and HB 145-28, may correspond to those of the salt bridge between Asp194 and Ile16 (existing only in chymotrypsin; Ile16 has become the N-terminus due to the autolysis) and the hydrogen bond between Ser195 and Gly43 (existing in both chymotrypsin and chymotrypsinogen) [35]. Actually, any hydrogen bond listed in Table 1 (explained below), particularly HB 140A-1B formed along with dimerization, may correspond to HB 194-16 in chymotrypsin. Figure 3A also shows that the catalytic dyad, Cys145 and His41, maintains the close arrangement in both states. Indeed, the distance between the two residues fluctuates by only a small extent (see the distance histogram in Fig. S3) because His41 is located at the less mobile hinge region of the H-loop.

**Table 1.** Statistics of C-loop conformation.

|  |  | Number of chains |  |  |  |  |  |  |  |  |  |  |
| --- | --- | --- | --- | --- | --- | --- | --- | --- | --- | --- | --- | --- |
|  |  | HB G143-N28 | PC1(C) |  | HB G138-H172 |  | HB S139-Y126 |  | HB F140A-S1B |  | HB L141-Y118 |  |
|  |  |  | > -1<sup>#</sup> | < -1 | Y<sup>+</sup> | N | Y | N | Y | N | Y | N |
| active* | bound | 166 | 165 | 1 | 164 | 2 | 165 | 1 | 130 (9)<sup>§</sup> | 36 (31) | 165 | 1 |
|  | free | 64 | 61 | 1 | 54 | 10 | 56 | 8 | 51 (2) | 13 (9) | 57 | 7 |
| collapsed | bound | 4 | 0 | 4 | 1 | 3 | 0 | 4 | 0 (0) | 4 (3) | 1 | 3 |
|  | free | 24 | 2 | 20 | 6 | 18 | 2 | 22 | 0 (0) | 20 (11) | 2 | 22 |
| rate of agreement<sup>%</sup> |  |  | 0.98 |  | 0.93 |  | 0.96 |  | 0.81, 0.95<sup>&</sup> |  | 0.96 |  |

\*active/collapsed is defined by whether HB G143-N28 exist or not. There are six indices to describe the C-loop conformation, including HB G143-N28.
<sup>#</sup>PC1(C) has the threshold of -1 separating the active/collapsed state.
<sup>+</sup>Y: HB exists; N: HB does not exist.
<sup>\$</sup>The number in the parenthesis is that of chains with the uncharged N-terminus due to some appended amino acids at Ser1.
<sup>%</sup>The rate of agreement with the numbers of chains of HB G1443-N28 are calculated by [(active/Y)+(collapsed/N)]/(total number of chains). The total number of chains is not necessarily 258 because of missing residues preventing from the assignment.
<sup>&</sup>The rate of agreement in HB 140-1 is given in two ways, the left value corresponds to the value counting all chains, and the right value was calculated for the chains with the native charged N-terminus.

**Figure 3.**
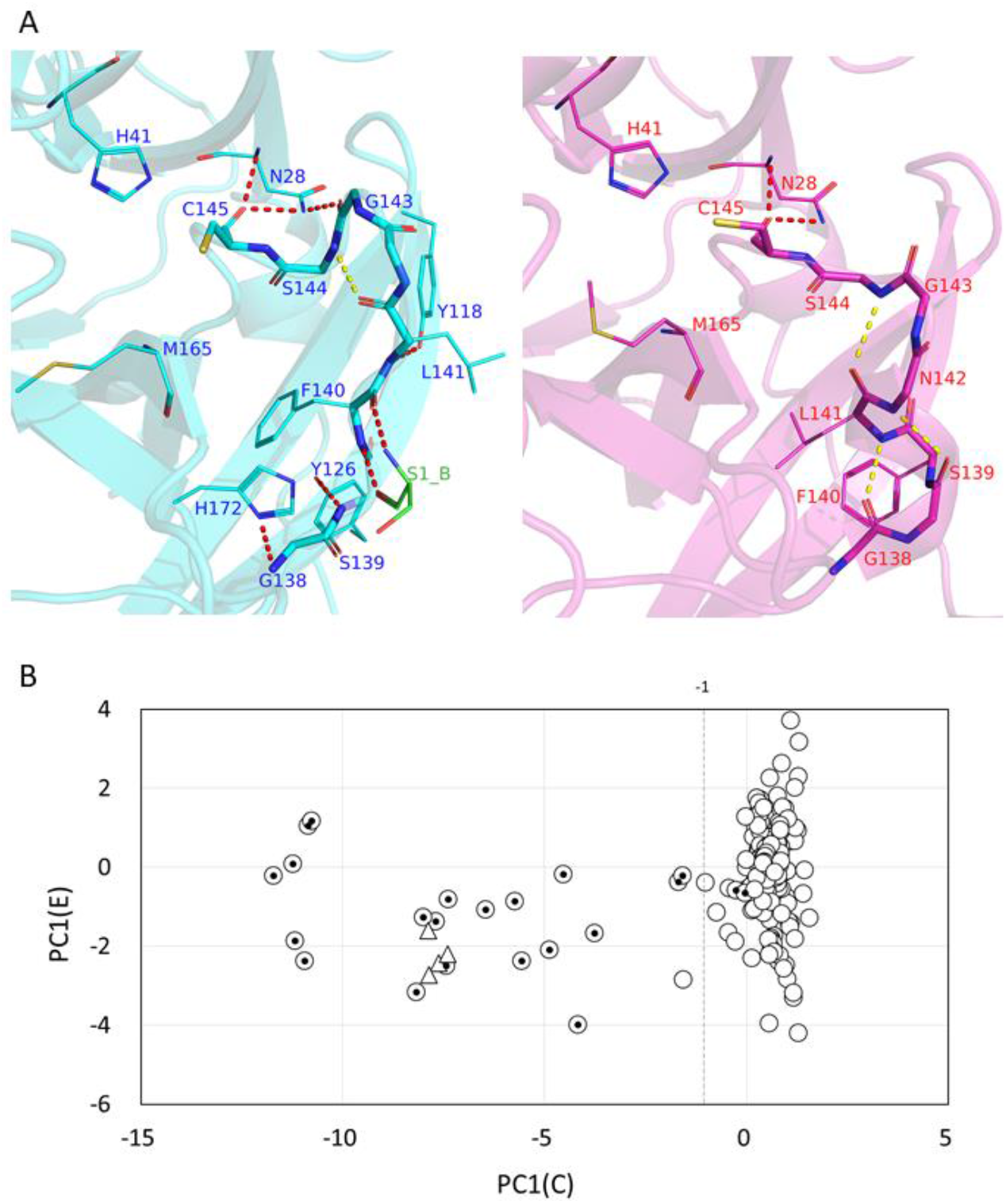
(A) The structures of the C-loop, the active state (left; PDB: 6lu7A, cyan; Ser1 of B chain, green), and the collapsed state (right, 2qcy, magenta). The hydrogen bonds are shown as broken lines; those with the other part of the molecule are in red and those within the main-chain of the C-loop are in yellow. In the active state, HB 143-28 locates Gly143 close to Cys145 to form the oxyanion hole (the main-chain NH groups of Gly143 and Cys145 recognize the oxyanion of the reaction intermediate). However, the collapsed state loses HB 143-28 to move Gly143 down and to separate it from Cys145. The main-chain HB 141-144 is found in both the active and collapsed states (HB 141-144 is found in 214 chains of 230 active chains, and in 11 chains of 28 collapsed chains). The reference atom for the measurement of the side-chain position of Phe140 and Leu141, the main-chain of Met165, and the catalytic dyad, Cys145 and His41, are also shown here. (B) The plot of PC1(C) against PC1(E) for 254 chains: dimer chains (circle) and monomer chains (triangles), representing the typical collapsed state. The inner dots indicate the collapsed chains defined by the absence of HB 143-28. The value PC1(C) = −1 is the threshold separating the active and collapsed states. PC1(E) is used here simply to display PC1(C) in a scatter plot; the distribution of PC1(E) is discussed elsewhere. All the statistics were calculated for the independent chains instead of the entries to deal with the asymmetric dimers.

#### Hydrogen bonds between the C-loop and surrounding residues cooperatively regulate catalytic function

When expanding the focus from the oxyanion hole to the whole C-loop, the conformation is described by various structural characteristics: the continuous principal component PC1 for the C-loop, PC1(C) (Fig. 2C), and five hydrogen bonds between the C-loop and residues surrounding the C-loop, HB 138-172, HB 139-126, HB 140A-1B (A and B indicate the two protomers), and HB 141-118, along with HB 143-28 (Fig. 3A). Among these HBs, HB 138-172 and HB 140A-1B are the hydrogen bonds with the moving clusters, i.e., the E-loop and the flexible fragment of the N-finger coupled with dimerization, respectively. These dynamic couplings with the C-loop are discussed below. The formation of these hydrogen bonds in each conformational state of the C-loop are summarized in Table 1; their formation was highly correlated with those of HB 143-28 (as indicated by the rate of agreement at the bottom of Table 1), except for HB 140-1 (explained below). Therefore, they represent almost the same structural information on the C-loop as that of HB 143-28; in other words, the conformational change occurs cooperatively involving these HBs as a structural transition. PC1(C) can be divided into the two categories with the threshold value of −1, namely the active state for PC1(C) > −1 and the collapsed state for PC1(C) <1, for which the threshold value was roughly optimized to give a high correlation. Figure 3B shows the distribution of PC1(C), in which the active state has a definite value, whereas the collapsed state shows large variation. Thus, the structural change in the C-loop between the two states can be regarded as an order–disorder transition (SupportingText 3 for details).

#### Charged states of the N-terminus affect the conformation of the C-loop

A rather poor agreement, 0.81, was found in the interprotomer HB 140-1. It has been reported that HB 140-1 is involved in an important interprotomer interaction coupled with dimerization to stabilize the active state, as well as another interprotomer hydrogen bond between Glu166A and Ser1B that occurs concurrently with HB 140-1 (160 chains have both of HB 140-1 and HB 166-1; 21 chains have only HB 140-1; and 5 chains have only HB 166-1) [25, 38]. Nevertheless, as shown in Table 1, fewer active chains have HB 140-1 than have the other HBs. This can be explained by the uncharged NH group of Ser1 produced by some amino acids appended to Ser1 [39] (data are summarized in Supporting data S1 and S2). The uncharged N-terminus weakens the interaction to Phe140 O to destabilize the polar contact (83% (= 54/65) of chains with the uncharged N-terminus do not have HB 140-1) and mimics the state prior to the cleavage of the polyprotein at the N-terminus. When only the chains with the innate charged N-terminus are counted in the statistics, the agreement with HB 143-28 increases up to 0.95, i.e., the same level as the other indices.

#### Absence of HB 140-1 has different effects in ligand-free and ligand-bound chains: ligand-induced activation

It is still necessary to clarify the reason why many chains in the active state do not need stabilization via HB 140-1 (Table 1). This situation differs between ligand-free chains and ligand-bound chains. For ligand-free chains, 10 of 13 active/ligand-free chains without HB 140-1 lose two more hydrogen bonds of the three HBs, 138-172, 139-126, and 141-118, on average (Supporting data S2). Thus, the absence of HB 140-1 destabilizes the C-loop to produce a marginally active state. The active/ligand-bound chains without HB 140-1 show a completely different feature, i.e., the other four HBs remain intact as well as PC1(C) > −1 (Supporting data S1). This can be explained by ligand-induced activation [40, 41]; the ligand interactions with the C-loop always contain interactions between a moiety mimicking the main-chain carbonyl group of the P1 site and the main-chain NH groups of Gly143 and Cys145, indicating that Gly143 and Cys145 are in the position of the active state. Table 1 shows that the probability of the chains found in the active state is 0.98 (= 166/(166+4)) in ligand-bound chains, whereas the probability decreases to 0.73 (= 64/(64+24)) in ligand-free chains. Stabilization by ligand molecules is also confirmed by the collapsed/ligand-bound chains; they are in the collapsed state because they do not have any ligand interaction at the C-loop (Supporting data S1; see the discussion on ligand binding below).

### Ligand binding and its influence on the structure leading to activation

#### Ligand binding sites are in the moving clusters

The variety of ligand binding is another subject of the crystal structure ensemble; the dataset contains 167 chains complexed with 92 different ligands (Supporting data S1). Ligand binding was analyzed in terms of the ligand binding sites defined by LigPlot [36]. As shown in Supporting data S3, binding is characterized by whether each residue has polar/nonpolar/covalent interactions with the ligand; 25 residues in 3CL^pro^ are identified as the major binding sites shared by 29–150 chains of the 167 ligand-bound chains. The major binding sites consist of hydrogen bonding residues known as the substrate binding subsites (S1–S6) [25, 34] and their neighboring residues forming the nonpolar contacts. Another 16 minor sites (see the caption of Supporting data S3) were found adjacent to the major binding sites and contained only 51 ligand-residue contacts in total. Notably, the major binding sites mostly overlap with the moving clusters defined by the Motion Tree, i.e., the C-loop, E-loop, Linker, and H-loop, and these binding sites are named as the following five “binding clusters”: the dimer interface region, H-loop, C-loop, E-loop, and Linker with some minor changes to the assignment (Supporting data S3). This observation indicates that the ligand binding sites, composed of the moving four loops, have a highly dynamic character that contrasts with a druggable rigid binding pocket; thus, it may not be easy to achieve high affinity without a covalent interaction at Cys145 (139 of the 167 ligand-bound chains have covalently bound ligands; Table 2A and Supporting data S3).

**Table 2.** Summary of the ligand binding analysis.

**A. Number of ligand contacts in each chain**
| ligand | Number of chains | Average number of binding sites/chain* |  |
| --- | --- | --- | --- |
|  |  | polar | nonpolar |
| peptide and peptide-mimic<sup>#</sup> | 124 (112)<sup>§</sup> | 6.1 | 10.7 |
| nonpeptide | 43 (27) | 2.5 | 7.1 |

\*These values were calculated by LigPlot [36].
<sup>#</sup>Peptide-mimic compounds are defined when a ligand contains more than two peptide moiety, and it is admitted that a carbonyl group is replaced by alcohol, and that Cα is in a part of an aromatic group.
<sup>§</sup>The number in the parenthesis is the number of chains with covalently-bound ligands. The number of nonpeptide ligands contains 2vj1A which has a ligand bound at residues other than Cys145.

**B. Number of chains in each binding mode**
| Binding mode | Number of chains |  |  |
| --- | --- | --- | --- |
|  | w | E | EL |
| peptide and peptide-mimic | 4 (3)<sup>§</sup> | 29 (28) | 92 (82) |
| nonpeptide | 25 (16) | 16 (7) | 0 |

<sup>§</sup>The number in the parenthesis is the number of chains with covalently-bound ligand.

#### Peptide substrates and nonpeptide compounds use essentially the same polar contact sites

To illustrate the structural details of ligand binding, we classified the ligands into the three types: the peptide substrate, peptide-mimic compound, and nonpeptide compound (see the caption of Table 2A for the definitions). As shown in Table 2A, the peptide/peptide-mimic compounds have more binding sites than are found in nonpeptide compounds, which is due to the size difference (average molecular weight: 578 for peptide/peptide-mimic compounds; 322 for nonpeptide compounds). The representative structures of the complexes are shown in Fig. S5. The peptide substrates are recognized by hydrogen bonds with the subsites: S1:C-loop (F140, G143, S144, and C145); E-loop (H163 and H164); H-loop (H41); S3:E-loop (E166); S4:Linker (Q189); and S6:Linker (Q192) (Fig. S5A). The peptide-mimic compounds are also bound at the peptide moieties by the subsites, although the hydrophobic side-chains do not necessarily have a definite orientation (Fig. S5B). In contrast, the nonpeptide compounds show a large variety of binding poses (Fig. S5C). However, when the polar interactions are focused, the subsites correctly make hydrogen bonds with the polar atoms of the nonpeptide compounds (Fig. S5D). Consequently, the variation in binding sites is strictly limited to a small set of the subsites and their adjacent nonpolar residues.

#### E-loop and Linker cooperatively change their conformations depending on the ligand size

We also examined the influence of ligand binding on the structure of 3CL^pro^. The effects of the C-loop have been discussed above; here, the conformations of the E-loop and Linker are examined. The conformation of the H-loop is not influenced by ligand binding but does exhibit intrinsic flexibility (Fig. S6). Figure 4A plots the PC1 values of the E-loop against those of the Linker, the distributions of which are not correlated (correlation coefficient: 0.181; Table S1). However, when the conformations of the E-loop and Linker are classified in terms of the binding cluster (Supporting data S3), a weak but definite binding pose dependence of the conformation is observed. Here, the binding poses are classified as “EL” (binding both the <u>E</u>-loop and <u>L</u>inker), “E” (binding only the <u>E</u>-loop), “w” (<u>w</u>eak; binding neither the E-loop nor Linker), and ligand-free (see the caption of Supporting data S3 for the definition). A one-dimensional histogram drawn along the collective variable, PC1(E)–PC1(L), more clearly shows the binding pose dependence (Fig. 4B). In the ascending order of PC1(E)–PC1(L), the ligand-free, “w,” “E,” and “EL” poses appear in the histogram. The histograms of the ligand-free and “w” poses almost overlap because the ligand binding of the “w” pose does not have an interaction with either the E-loop or Linker. However, the four monomeric crystal structures are situated in the histogram at the smallest extreme because the largely skewed position of domain III makes PC1(L) have largely negative values (Supporting data S2). These data indicate that the value of PC1(E)–PC1(L) increases when the binding sites expand more to the E-loop and Linker. The representative structures of these groups show that when the collective variable increases, the E-loop shifts downward to accommodate larger ligands (Fig. 4D). At the same time, the N-terminal (upper) part of the Linker shifts inward to make more interactions with the ligand, whereas the C-terminal (lower) part that does not participate in binding moves outward. The collective variable PC1(E)–PC1(L) correctly represents these motions in one-dimension. As shown in Fig. 4C, the motion along the collective variable of PC1(E)+PC1(L), perpendicular to PC1(E)–PC1(L), is an opening/closing motion of the two loops that exhibit almost no binding pose dependence. This suggests that PC1(E)+PC1(L) represents the intrinsic fluctuations. The representative structures are shown in Fig. S7. The majority of the nonpeptide compounds have the binding pose “w” because of their small size (Table 2B).

**Figure 4.**
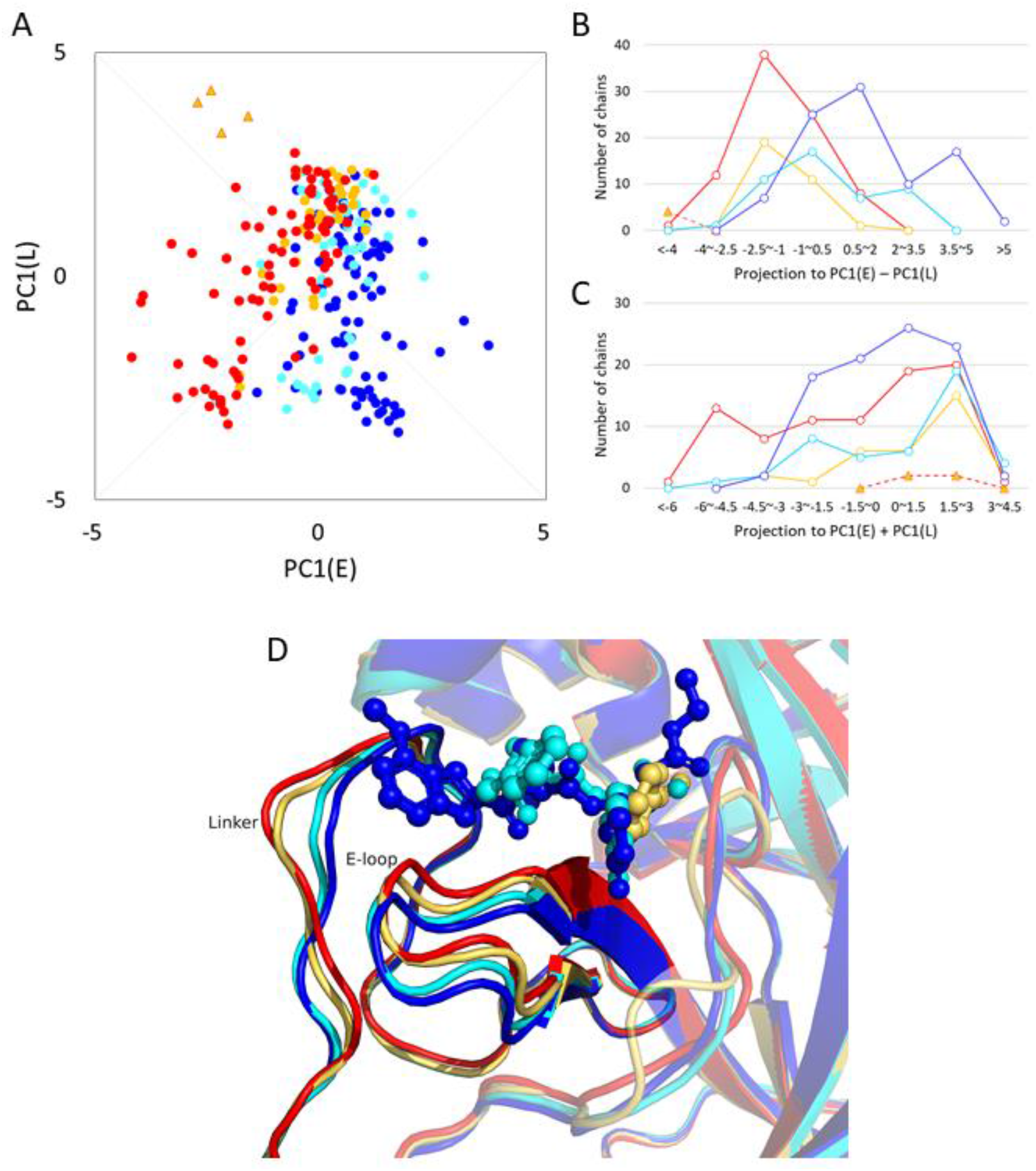
(A) A scatter plot of PC1(E) vs. PC1(L) for the ligand-bound chains. The color scheme for the binding poses is as follows: the ligand-free, red; “w,” light orange; “E,” cyan; “EL,” blue, and the monomeric structure, orange triangle. The collective variables, PC1(E)–PC1(L) and PC1(E)+PC1(L), are shown by gray lines. (B) Histogram of the projection of the data shown in (A) onto the collective variable, PC1(E)–PC1(L). The color scheme is the same as that used in (A). (C) Histogram of the projection of the data shown in (A) onto the collective variable, PC1(E)+PC1(L). (D) The representative structures of the binding poses: ligand-free (PDB:3m3vB; red; PC1(E)–PC1(L) = −3.875), “w” (2vjlB; light orange; −1.566), “E” (7d1mB; cyan; 2.715), and “EL” (6xhoA; blue; 5.181). Superposition was for the core region. With an increasing value of PC1(E)– PC1(L), the E-loop shifts downward, and the N-terminal (upper) part of the Linker moves inward toward the ligand position, whereas the C-terminal (lower) part, not involved in ligand binding, moves outward.

#### Downward motion of the E-loop caused by ligand binding stabilizes the active state of the C-loop

We also investigated the influence of the E-loop on the C-loop, or on catalytic activity. As shown in Table 1, one of the hydrogen bonds stabilizing the active state of the C-loop, HB 138-172, represents the direct interaction between the C-loop (Gly138) and E-loop (His172). As explained above, the E-loop makes downward motions depending on the size of the bound ligand (the larger the ligand becomes, the more the E-loop shifts downward). Figure 5A shows the representative structures with the E-loop situated at the lower position due to ligand binding and with the E-loop at the upper position in the ligand-free chain (PDB: 7brpA and 7bro, respectively); the former structure forms a hydrogen bond between His172 and Gly138, whereas the latter structure does not form this bond. Statistics for the PC1(E) dependence of the hydrogen bond formation are shown in Fig. 5B; a monotonous increase was observed for the probability of formation of HB 138-172 with increasing PC1(E) (the downward motion). Simultaneously, the probability of the C-loop in the active state increases. To avoid the influence of ligand-induced activation, we also calculated the same quantities for ligand-free chains. The difference between the values for all chains and those for the ligand-free chains can be ascribed to the effect of ligand-induced activation, but the behavior of the increase with PC1(E) is found in both chains. Therefore, the downward motion of the E-loop stabilizes HB 138-172, which then stabilizes the active state of the C-loop. The interrelation among the three features, the bound ligand size, the motion of the E-loop, and the formation of HB 138-172, suggests that the activity of 3CL^pro^ is maximized in large native substrates.

**Figure 5.**
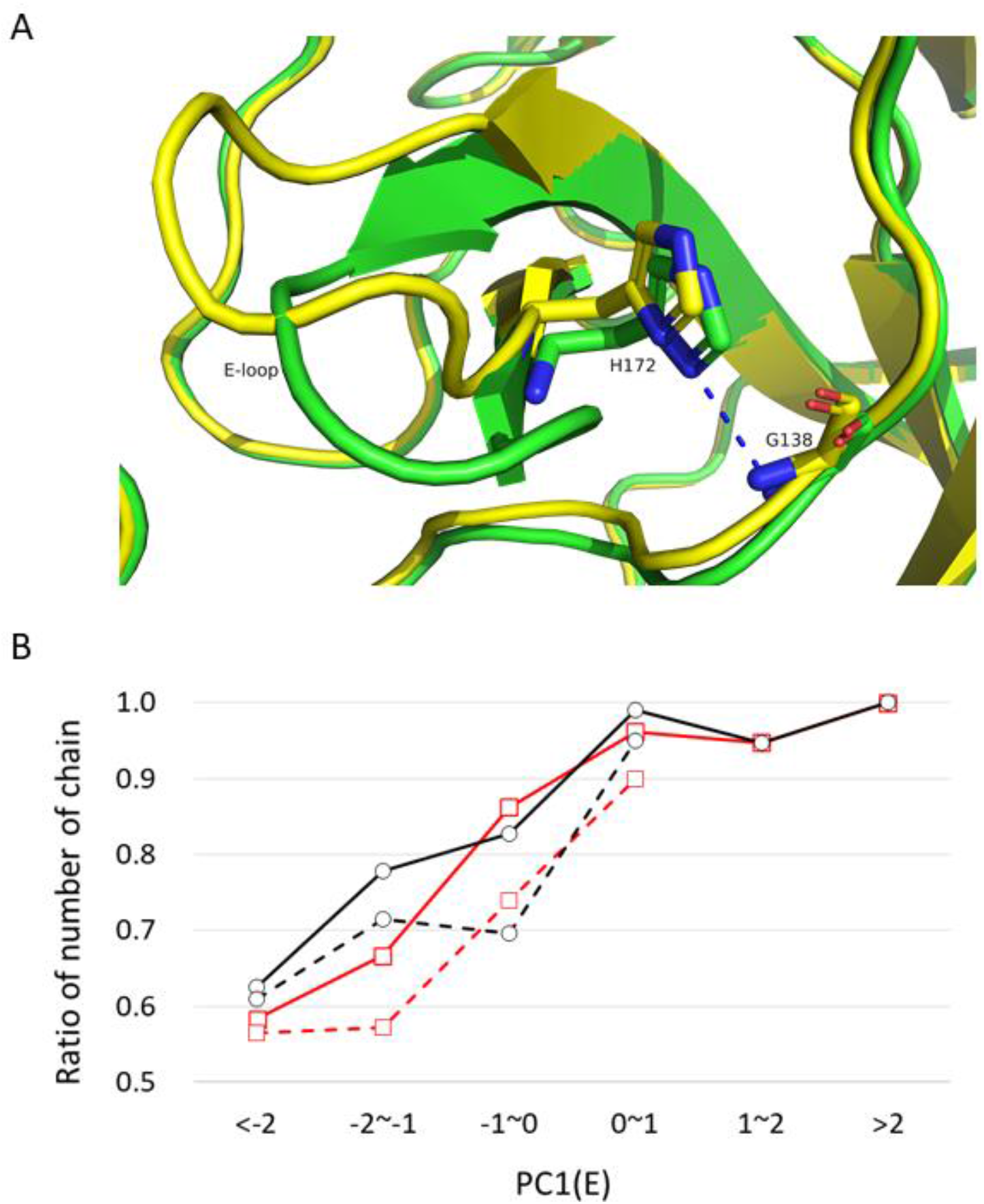
(A) Representative structures exhibiting the position of the E-loop in the lower position (PDB:7brpA (ligand-bound); green; PC1(E) = 3.712) and upper position (PDB:7bro (ligand-free); yellow; −4.201), drawn after superposition at the core region. The ligand of 7brpA is drawn as lines. HB 138-172 is formed in 7brpA and depicted as a blue dotted line, whereas 7bro does not form HB 138-172. (B) PC1(E) dependence of the probability of formation of hydrogen bonds. Solid curves are those for all chains and broken curves are for ligand-free chains. Black curves with circles are for HB143-28 defining the active state of the C-loop; red curves with squares are for HB 138-172 representing the interaction between the C-loop and E-loop. The total numbers of chains at each interval of PC1(E) are, in ascending order, 24, 27, 58, 105, 38, and 6 for all chains, and 23, 21, 23, and 20 for ligand-free chains (for ligand-free chains, data on intervals > 1 are not presented because the number of chains is not sufficient to calculate statistics).

### Influences of T285A mutation and allosteric functional regulation through domain III

#### T285A mutation closes the interface of the domain III dimer

In the above sections, we did not distinguish between SARS-CoV 3CL^pro^ and SARS-CoV-2 3CL^pro^ in our analysis of the crystal structure ensemble. However, since there are 12 amino acid alterations between the two 3CL^pro^, the influence of the mutations is discussed in this section. Ten sites among the twelve mutations are mostly located on the surface of the core part of domains I and II (solvent accessibility: ~0.8); therefore, they have no substantial influence on the structure. However, two mutations, T285A and I286L, are located on the interface of the domain III dimer and have substantial effects on structure. Figure 6A shows the distributions of the interprotomer Cα distance between Thr(Ala)285A and Thr(Ala)285B; the distance of SARS-CoV-2 3CL^pro^ is much shorter than that of SARS-CoV 3CL^pro^, as has already been observed in the structure of a triple mutant S284-T285-I286/A of SARS-CoV 3CL^pro^ [42]. In Fig. 6C, the representative configurations of the interface of the domain III dimer are compared; the interprotomer hydrophobic contacts are formed among Ala285A(B), Ala285B(A), and Leu286B(A) in SARS-CoV-2 3CL^pro^, whereas Thr285 of SARS-CoV 3CL^pro^ is distant from its counterpart. The smaller size of the alanine side-chain relative to that of threonine enables a shorter interprotomer distance. Furthermore, the hydrophobic packing of a pair of the alanine/leucine residues has greater affinity than that of a weak hydrogen bond between the two hydroxyl groups of threonine (a hydroxyl group may preferably make a hydrogen bond with water).

**Figure 6.**
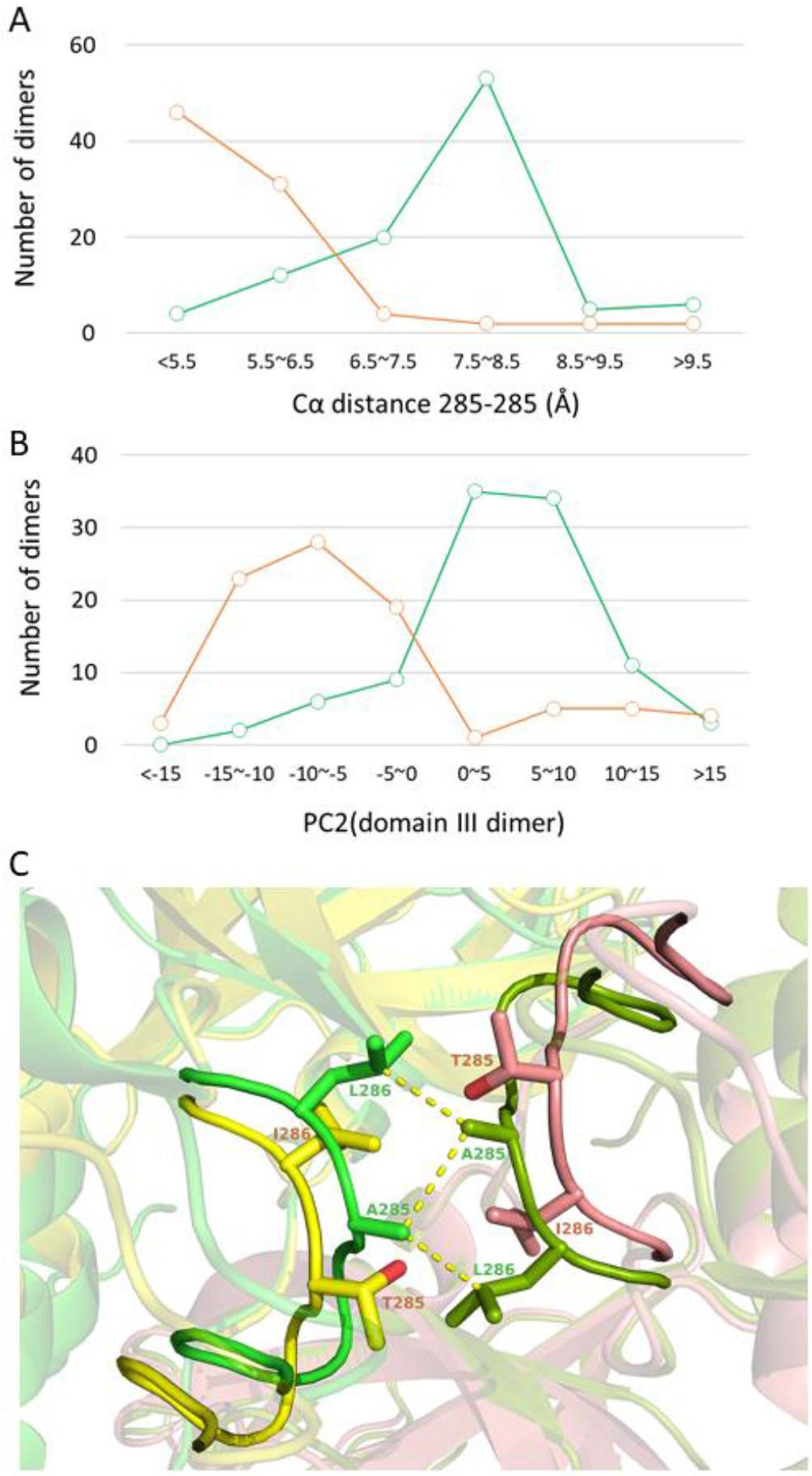
(A) The distributions of Cα distance 285-285: SARS-CoV 3CL^pro^ (green; mean = 7.7 ± 1.0 Å) and SARS-CoV-2 3CL^pro^ (orange; mean = 5.8 ± 1.0 Å). Note that the data of SARS-CoV 3CL^pro^ contain ten chains of the triple mutant S284-T285-I286/A, for which the mean value was 6.2 ± 1.0 Å. The minimum value of distance 285-285 is 5.0 Å. (B) The distributions of PC2(domain III dimer), a parameter representing the width of the interface of the domain III dimer (larger values of PC2 correspond to more open forms) for SARS-CoV 3CL^pro^ (green) and SARS-CoV-2 3CL^pro^ (orange). (C) Representative structures of the interface of the domain III dimer: 2gt8 (SARS-CoV 3CL^pro^; yellow (chain A) and salmon (chain B); distance 285-285 = 9.131 Å and PC2(domain III dimer) = 11.365) and 6m03 (SARS-CoV-2 3CL^pro^; green and pale green; 5.227 Å and −8.573), drawn after superposition at the core region. These entries are both symmetric dimer. The hydrophobic contacts are depicted by yellow dotted lines (both of the contact lengths are 3.6 Å).

We now focus on the details of the dynamics that occur along with the change in distance 285-285. As shown in Fig. 6C, Thr(Ala)285 is in a 16-residue-long loop (residues 276-291). However, as illustrated in the Motion Tree (Fig. 2A), the fragment involving Thr(Ala)285 (residues 280-287) is in a section of the moving cluster that constitutes the rigid core of domain III. Thus, this fragment is rigid and does not change the conformation independently of domain III, probably due to its winding shape with intraloop hydrogen bonds. Therefore, the difference in distance is not caused by the internal motion of the loop but rather by the rigid body motion of domain III. The respective distributions of PC2(domain III dimer), a parameter describing the configuration of the domain III dimer (Fig. S1), of SARS-CoV 3CL^pro^ and SARS-CoV-2 3CL^pro^ are largely separated, similar to the distributions of distance 285-285 (Fig. 6B); domain III of SARS-CoV-2 3CL^pro^ has more closed arrangements relative to the open arrangements of SARS-CoV 3CL^pro^. Indeed, the mode structure of PC2(domain III dimer) agrees well with the direction of motion in Thr(Ala)285 departing from the counterpart of the other protomer (Fig. S1). In Fig. S8, a clear correlation between PC2(domain III dimer) and distance 285-285 is also shown.

#### SARS-CoV-2 3CL^pro^ has a higher activity than SARS-CoV 3CL^pro^: experimental evidence

We investigated whether the difference in the configuration of the domain III dimer, observed between SARS-CoV-2 3CL^pro^ and SARS-CoV 3CL^pro^ (Fig. 6A), affected the conformation of the C-loop or influenced catalytic activity. First, we analyzed the experimental data. Kinetic experimental data [24, 43, 44]) indicate that SARS-CoV-2 3CL^pro^ has a larger catalytic efficiency than SARS-CoV 3CL^pro^, although the kinetic parameters differ greatly among the three studies, 3-fold greater efficiency [43] and slightly greater efficiency [24, 44]. Furthermore, it was reported that the T285A mutant of SARS-CoV 3CL^pro^ had ~1.4-fold higher activity than the wild-type and that the triple mutant S284-T285-I286/A showed ~3.7-fold higher activity [44]. Given that distance 285-285 is reduced to 6.2 Å in the triple mutant S284-T285-I286/A (for the seven mutant dimers), down from the 7.7-Å of the average of all SARS-CoV 3CL^pro^ (Fig. 6A), it is reasonable to conclude that the motions of domain III influence catalytic activity.

#### Coupling between domain III and the C-loop: the uncharged N-terminus

Based on the analysis of the crystal structure ensemble, we assessed the connection between domain III and the C-loop (Fig. 7A and 7C). The probability of finding the C-loop in the active state, as well as the probability of HB 140-1 being formed, decreases with increasing distance 285-285, particularly over 8.5 Å (Fig. 7A), which clearly indicates the allosteric coupling between domain III and the C-loop. The chains with an uncharged N-terminus (which is highly unlikely to form HB 140-1) accumulate in the region of distance 285-285 over 8.5 Å (Fig. 7A). This observation can be understood as a causal relationship in which the absence of HB140-1 due to the uncharged N-terminus induces the opening motion of domain III dimer.

**Figure 7.**
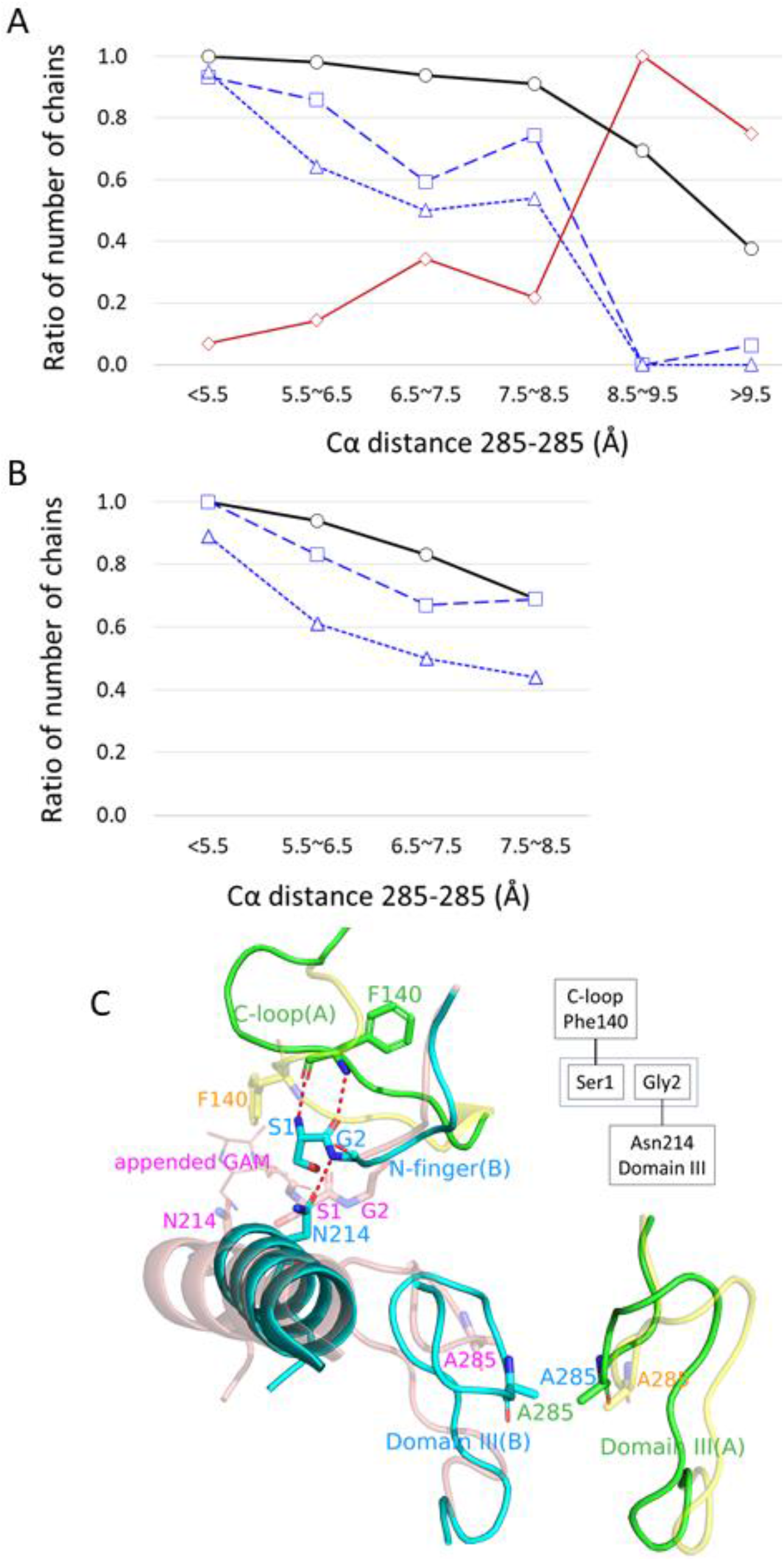
(A) Distributions along Cα distance 285-285: the probability of formation of the active C-loop conformation (thick black curve with circles); the probability of finding a chain with the uncharged N-terminus due to some appended amino acids (solid brown curve with diamonds); the probability of finding a chain with HB 140-1 (broken blue curve with squares); the probability of finding a chain with HB 214-2 (dotted blue curve with triangles). The total number of chains at each interval of distance 285-285 is, in ascending order, 59, 57, 32, 78, 13, and 15. (B) As in (A), but for ligand-free chains with natively charged N-termini without an appended amino acid. The intervals of 8.5 ~ 9.5 and > 9.5 are not presented because the number of chains in these intervals are not sufficient to calculate statistics. The total number of chains at each interval of distance 285-285 is, in ascending order, 18, 18, 6, and 16. (C) Two 3CL^pro^ structures that explain the scenario for the accumulation of chains with uncharged N-termini at large distance 285-285, drawn after superposition at the core region. These structures are PDB:6lu7 (SARS-CoV-2): A chain (green) and B chain (cyan); distance 285-285 = 5.314 Å, and PDB:7kfi (SARS-CoV-2) having A chain (salmon) and B chain (yellow) with transparency; distance = 9.858 Å. Structures are superimposed at the core region of 6lu7A and 7kfiB. Only the key parts are illustrated. The entry 6lu7 has a natively charged N-terminus, whereas 7kfi has an appended sequence at the N-terminus (Gly(−2), Ala(−1), and Met0) drawn as lines. 6lu7 has both HB 140-1 and HB 214-2 (red broken lines). However, 7kfi does not have either HB 140-1 or HB 214-2 and the C-loop is collapsed (Phe140 is oriented to the other direction) because of the uncharged N-terminus. Flexible Ser1 of 7kfi induces a shift of the position of Gly2 to break HB 214-2. The absence of HB214-2 causes the motion of domain III to separate Ala285. The upper right diagram shows a schematic diagram of the coupling between the C-loop and domain III via HB 140-1 and HB 214-2.

Here, we propose a possible scenario to explain these observations, using the structures illustrated in Fig. 7C together with a variety of experimental evidence. The uncharged N-terminus tends to break HB 140-1, as discussed above; thus, it destabilizes the active state in the C-loop. However, since neither Phe140 nor Ser1 directly interacts with domain III, it is necessary to identify a factor linking domain III and HB 140-1. As a possible factor, we identified an intraprotomer/interdomain hydrogen bond between Asn214 OD1 and Gly2 N (HB 214-2). This unique interdomain interaction forms and breaks in accordance with the position of domain III or distance 285-285; the opening motion along PC2(domain III dimer) separates Asn214 from Gly2 (Fig. S1), and distance 214-2 is well correlated with PC2(domain III dimer) (Fig. S9A). As shown in Fig. 7A, the probability of formation of HB 214-2 decreases with distance 285-285. In contrast, the other interdomain interactions occur at the hinge region of the domain motion and are stably maintained almost independently of the domain III position (Fig. S9B). Although these stable interactions do not operate as a switch, they have significant contributions to stabilizing the dimer structure; the mutations at the residues illustrated in Fig. S9B (Arg4, Ser123, Ser139, Glu290, Arg298, and Gln299) impair dimerization and catalysis [46, 47], and the mutations R298A (PDB:2qcy and 3m3t) and S139A (PDB:3f9e) produce the monomeric crystal structures.

The connection between HB 140-1 and HB 214-2 is explained as follows. The absence of HB 140-1 allows Ser1 to move freely and to be separated from Phe140. This conformational change accompanies the shift of Gly2 to destabilize HB 214-2 (the probability of formation of HB 214-2 decreases from the unconditional value of 0.59 (=150/254) for all chains to 0.19 (= 14/73) under the condition without HB 140-1). Finally, the influence of HB 214-2 to the domain III dimer can be observed in the mutant N214A disabling to form HB 214-2. The structures of N214A (PDB: 2qc2 and 3m3s; both are SARS-CoV 3CL^pro^) have large values of distance 285-285 (8.6 Å and 8.4 Å, respectively; the average distance of SARS-CoV 3CL^pro^ is 7.7 Å). These structures suggest that the loss of HB 214-2 tends to open the domain III dimer. In summary, we observed the following relationship: the absence of HB 140-1 decreases the probability of formation of HB 214-2 and then the absence of HB214-2 opens the interface of the domain III dimer.

#### Coupling between domain III and the C-loop: the charged N-terminus

The uncharged N-terminus is chemically unchangeable and makes the absence of HB 140-1 independent of the other components such as the configuration of the domain III dimer and HB 214-2. Therefore, the role of HB 140-1 in functional regulation is most evidently observed in chains with an uncharged N-terminus (Fig. 7A). Conversely, in the natively charged N-terminus, the occurrence of HB 140-1 is changed reversibly under the influence of other components. However, this complicated situation occurs in the native condition and should also be investigated. Furthermore, the ligand-induced activation is another factor obscuring the influence of domain III on the C-loop because the C-loop conformation is determined by interactions with the ligand molecule. Hence, we used the ligand-free chains with charged N-termini to recalculate the quantities shown in Fig. 7A, although the number of chains was significantly reduced from 254 to 62. Figure 7B shows the results of the recalculation (the values for > 8.5 Å are not shown because the number of chains in this distance range was not sufficient to calculate statistical quantities). The numbers of chains with HB 140-1 and HB 214-2 were shown to decrease with distance 285-285; these values did not differ largely from those shown in Fig. 7A. However, the number of chains in the active state, or chains with HB 143-28, clearly showed a monotonous decrease with distance 285-285 from 1.0 (distance < 5.5 Å) to 0.69 (~7.5–8.5 Å); these data contrast with those in Fig. 7A showing values kept close to unity due to ligand-induced activation. Overall, these data clearly demonstrate that the opening motion of the domain III dimer has a destabilizing effect on the active state of the C-loop through the dissociation of HB 214-2 and HB 140-1.

Based on the results presented above, we compared the numbers of formation of the hydrogen bonds for SARS-CoV 3CL^pro^ and SARS-CoV-2 3CLpro using 62 ligand-free chains with a natively charged N-terminus (Table 3). We found that the closed configuration of the domain III dimer in SARS-CoV-2 3CLpro results in a 0.54 greater probability of formation of HB 214-2 than the probability for SARS-CoV 3CL^pro^, as well as a 0.25 increase in the probability of formation of HB 140-1 and a 0.14 increase in the probability of occurrence of the active state. Although the influence is reduced by half for each interaction step connecting the four structural elements (the domain III dimer, HB 214-2, HB140-1, and C-loop), our analyses suggest that SARS-CoV-2 3CL^pro^ has slightly increased activity over that of SARS-CoV 3CL^pro^, which is largely consistent with the experimental data.

**Table 3.** Comparison of the two 3CL^pro^ using ligand-free chains with the natively charged N-terminus.

|  | active | HB 140-1 | HB 214-2 | All |
| --- | --- | --- | --- | --- |
| SARS-CoV | 0.79 (27) | 0.68 (23) | 0.35 (12) | 34 |
| SARS-CoV-2 | 0.93 (26) | 0.93 (26) | 0.89 (25) | 28 |

The probability was calculated as the number of chains having each HB (the number in the parenthesis) divided by the total number of chains listed in the column “All”. The column of “active” is those having HB 143-28 as in the above definition.

## Conclusion

The crystal structure ensemble, consisting of 258 independent chains, successfully describes the structural dynamics of SARS-CoV 3CL^pro^ and SARS-CoV-2 3CL^pro^ as well as elucidates the allosteric regulation of catalytic function. The structural dynamics is characterized by the motion of four loops (the C-loop, E-loop, H-loop, and Linker) and domain III on the rigid core. Among the four loops, the C-loop causes the order (active)– disorder (collapsed) transition, which is regulated cooperatively by the five hydrogen bonds with the surrounding residues. Three of the loops, the C-loop, E-loop, and Linker, constitute the major ligand binding sites with a limited variety of binding residues including the subsites. Ligand recognition at the main-chain NH groups of Gly143 and Cys145 induces the formation of an oxyanion hole-like structure to produce the active conformation of the C-loop (i.e., ligand-induced activation). Ligand binding also causes the ligand size dependent conformational changes to the E-loop and Linker, which further stabilize the C-loop through HB 138-172. Mutation T285A from SARS-CoV 3CL^pro^ to SARS-CoV-2 3CL^pro^ significantly closes the interface of the domain III dimer and affects the stability of the C-loop conformation allosterically via HB140-1 and HB 214-2.

Because of this allosteric regulation, the closed arrangement of the domain III dimer in SARS-CoV-2 3CL^pro^ increases the stability of the active state of the C-loop and yields a slightly higher activity than that of SARS-CoV 3CL^pro^. As a reference to the present results, the crystal structures of MERS-CoV 3CL^pro^, 3CL^pro^ of Middle East respiratory syndrome-related coronavirus (2012), were analyzed in a similar manner as in SARS-CoV and SARS-CoV-2 3CL^pro^. It was found that the structural and functional properties were basically the same as those of SARS-CoV and SARS-CoV-2 3CL^pro^. The details are summarized in Supporting text 4.

## Material and Methods

### Analysis of the crystal structure ensemble

As a scheme for the analysis of the crystal structure ensemble, the identification of the overall dynamic structure should precede the PCA, and then the PCA is applied separately to various moving parts of the protein. This scheme is to avoid the application of the PCA to the whole protein molecule. It is because the PCA tends to produce a mode structure in which the mode vector has non-zero elements at all atoms considered in the analysis, representing long-range correlation in motion, due to the orthonormal condition in the eigenvalue problem. Therefore, it is difficult to describe a localized motion by the PCA of the whole protein molecule. Motion Tree using the variance of the residue distance enables us to identify the moving clusters of any size from a domain level to a residue level without any prior knowledge (see below and Fig. 2A). Each of the moving clusters thus found is then separately subjected to the PCA (Fig. 2C and Fig. S1). Here, it is important that the PCA does not exclude the translation and rotation motions of the moving clusters; the motion should be defined as a relative motion against the core region of the protein via superimposition onto the core region.

### Data used in this study

The crystal structure ensemble was constructed for 3CL^pro^ of SARS-CoV-2 and SARS-CoV based on the PDB data of the version of 10/25/2020. The following entries were not used in the analysis: the entries from the PanDDA analysis (115 entries), the entry with domain swapping (PDB:3iwm), and the entries containing only domain III (PDB: 2k7x, 2liz, and 3ebn). The compiled data contains 83 entries/113 independent chains for SARS-CoV-2 3CL^pro^ and 101 entries/145 independent chains for SARS-CoV 3CL^pro^. These are listed in Supporting data S1 and S2 for the ligand-bound and ligand-free entries, respectively. The data after 10/25/2020 until 7/25/2021 were summarized in Supporting data S4, S5 and S6, which correspond to Supporting data S1, S2 and S3, respectively (SARS-CoV-2 3CL^pro^: 154 entries and 226 chains; SARS-CoV 3CL^pro^: 5 entries and 6 chains). The analyses of Supporting data S4, S5 and S6 were summarized in Fig. S10.

### Motion Tree

Protein dynamics, or the pattern of motions in the crystal structure ensemble, can be described by decomposing the overall motion into slow motions between the building blocks and fast vibrations within each building block. We developed a method to define the building blocks moving as rigid bodies, which we achieved through hierarchical clustering of interresidue distances (for pairwise comparisons) or their variances (for the comparison of many entries) and subsequent construction of a dendrogram, namely the Motion Tree [27, 28].

The Motion Tree illustrates, in a hierarchical manner, a pair of rigid-like clusters at each node that moves reciprocally with the amplitude of the tree height of the node named “Motion Tree (MT) score.” Because of the straightforward application to the structure ensemble without the need for a structural superposition procedure, a comprehensive understanding of the structural dynamics of various protein molecules can be achieved [17, 48–50].

We compared 258 chains in a single Motion Tree using the variance-based scheme. The variance of distance fluctuation, {*D_mn_*}, used as a metric for hierarchical clustering, is calculated as *D_mn_* =<Δ*d*^2^_*mn*_>^1/2^, where *d_mn_* is the distance between Cα atoms of residues *m* and *n*, Δ*d_mn_* is the associated deviation from the mean distance, and <…> is the average over the structural ensemble. We did not include highly mobile Ser1 and Gly2, as well as C-terminal residues 301–306, in the analysis because these residues are often in the list of missing residues. Since 3CL^pro^ is in the homodimeric form, *D_mn_* and the resulting clusters have to be symmetrical upon the exchange of the two protomers. However, asymmetric dimers exist in the crystal structure ensemble; they are considered to be under the influence of crystal packing or in different states of ligand binding. For the purpose of removing these influences, *D_mn_* was symmetrized using the duplicated structures of AB and BA, where AB is the original dimer and BA is the dimer with the protomers exchanged. Because of this symmetrization, two equivalent clusters corresponding to each protomer exist in the Motion Tree. This symmetrizing operation was also applied to the calculation of the PCs for the domain III dimers.

## Acknowledgments

This research was supported by Platform Project for Supporting Drug Discovery and Life Science Research (Basis for Supporting Innovative Drug Discovery and Life Science Research (BINDS)) from AMED under Grant Number JP21am0101109. We are grateful to Dr. R. Koike and Dr. M. Ota, Nagoya University, for their helpful comments.

## Author Contributions

A.K. and K.M. and designed research; A.K. and K.M. performed research and analyzed data; and A.K., K.M., T.E. and M.I. wrote the paper.

## References

[1] Blundell, T. L., Jhoti, H. & Abell, C. (2002). High-throughput crystallography for lead discovery in drug design. Nat. Rev. Drug Discov. 1, 45–54.

[2] Kuhn, P., Wilson, K., Patch, M. G. & Stevens, R. C. (2002). The genesis of high-throughput structure-based drug discovery using protein crystallography. Curr. Opin. Chem. Biol. 6, 704–710.

[3] Anderson, A. C. (2003). The process of structure-based drug design. Chem. Biol. 10, 787–797.

[4] Congreve, M., Murray, C. W. & Blundell, T. L. (2005). Keynote review: Structural biology and drug discovery. Drug Discov. Today. 10, 895–907.

[5] Mestres, J. (2005). Representativity of target families in the protein data bank: impact for family-directed structure-based drug discovery. Drug Discov. Today. 10, 1629–1637.

[6] Westbrook, J. D. & Burley, S. K. (2019). How Structural Biologists and the Protein Data Bank Contributed to Recent FDA New Drug Approvals. Structure. 27, 211–217.

[7] Rawlings, N. D., Alan, J., Thomas, P. D., Huang, X. D., Bateman, A. & Finn, R. D. (2018). The MEROPS database of proteolytic enzymes, their substrates and inhibitors in 2017 and a comparison with peptidases in the PANTHER database. Nucleic Acids Res. 46, D624–D632.

[8] Munk, C., Mutt, E., Isberg, V., Nikolajsen, L. F., Bibbe, J. M., Flock, T., Hanson, M. A., Stevens, R. C., Deupi, X. & Gloriam, D. E. (2019). An online resource for GPCR structure determination and analysis. Nat. Methods. 16, 151–162.

[9] Reau, M., Lagarde, N., Zagury, J. F. & Montes, M. (2019). Nuclear receptors database including negative data (NR-DBIND): A database dedicated to nuclear receptors binding data including negative data and pharmacological profile. J. Med. Chem. 62, 2894–904.

[10] Kanev, G. K., de Graaf, C., Westerman, B. A., de Esch, I. J. P. & Kooistra, A. J. (2021). KLIFS: an overhaul after the first 5 years of supporting kinase research. Nucleic Acids Res. 49, D562–D569.

[11] Boehr, D. D., Nussinov, R. & Wright, P. E. (2009). The role of dynamic conformational ensembles in biomolecular recognition. Nat. Chem. Biol. 5, 789–796.

[12] Feixas, F., Lindert, S., Sinko, W. & McCammon, J. A. (2014). Exploring the role of receptor flexibility in structure-based drug discovery. Biophys. Chem. 186, 31–45.

[13] Andrec, M., Snyder, D. A., Zhou, Z. Y., Young, J., Montellone, G. T. & Levy, R. M. (2007). A large data set comparison of protein structures determined by crystallography and NMR: Statistical test for structural differences and the effect of crystal packing. Proteins. 69, 449–465.

[14] Cavasotto, C. N. & Phatak, S. S. (2009). Homology modeling in drug discovery: current trends and applications. Drug Discov. Today. 14, 676–683.

[15] Ma, B. Y., Kumar, S., Tsai, C. J. & Nussinov, R. (1999). Folding funnels and binding mechanisms. Protein Eng. 12, 713–720.

[16] Ikeguchi, M., Ueno, J., Sato, M. & Kidera, A. (2005). Protein structural change upon ligand binding: Linear response theory. Phys. Rev. Lett. 94, Article 078102.

[17] Moritsugu, K., Nishino, Y. & Kidera, A. (2020). Inter-lobe motions allosterically regulate the structure and function of EGFR kinase. J. Mol. Biol. 432, 4561–4575.

[18] Goyal, B. & Goyal, D. (2020). Targeting the dimerization of the main protease of coronaviruses: A potential broad-spectrum therapeutic strategy. ACS Comb. Sci. 22, 297–305.

[19] Ullrich, S. & Nitsche, C. (2020). The SARS-CoV-2 main protease as drug target. Bioorg. Med. Chem. Lett. 30, Article 127377.

[20] Zhu, W., Xu, M., Chen, C. Z., Guo, H., Shen, M., Hu, X., Shinn, P., Klumpp-Thomas, C., Michael, S. G. & Zheng, W. (2020). Identification of SARS-CoV-2 3CL protease inhibitors by a quantitative high-throughput screening. ACS Pharmacol. Transl. 3, 1008–1016.

[21] Liu, C., Zhou, Q. Q., Li, Y. Z., Garner, L. V., Watkins, S. P., Carter, L. J., Smoot, J., Gregg, A.C., Daniels, A. D., Jervey, S. & Albaiu, D. (2020). Research and development on therapeutic agents and vaccines for COVID-19 and related human coronavirus diseases. ACS Central Sci. 6, 315–331.

[22] Li, G.D. & de Clercq, E. (2020). Therapeutic options for the 2019 novel coronavirus (2019-nCoV). Nat. Rev. Drug Discov. 19, 149–150.

[23] Sanders, J. M., Monogue, M. L., Jodlowski, T. Z. & Cutrell, J. B. (2020). Pharmacologic treatments for coronavirus disease 2019 (COVID-19) A review. JAMA 323, 1824–1836.

[24] Jin, Z. M., Du, X. Y., Xu, Y. C., Deng, Y.Q., Liu, M. Q., Zhao, Y., Zhang, B., Li, X., Zhang, L., Peng, C., Duan, Y., Yu, J., Wang, L., Yang, K., Liu, F., Jiang, R., Yang, X., You, T., Liu, X., Yang, X., Bai, F., Liu, H., Liu, X., Guddat. L. W., Xu, W., Xiao, G., Qin, C., Shi, Z., Jiang, H., Rao, Z. & Yang, H. (2020). Structure of M-pro from SARS-CoV-2 and discovery of its inhibitors. Nature. 582, 289–293.

[25] Yang, H. T., Yang, M. J., Ding, Y., Liu, Y. W., Lou, Z. Y., Zhou, Z., Sun, L., Mo, L., Ye, S., Pang, H., Gao, G. F., Anand, K., Bartlam, M., Hilgenfeld, R. & Rao, Z. (2003). The crystal structures of severe acute respiratory syndrome virus main protease and its complex with an inhibitor. Proc. Natl. Acad. Sci. U. S. A. 100, 13190–13195.

[26] Anand, K., Palm, G. J., Mesters, J. R., Siddell, S. G., Ziebuhr, J. & Hilgenfeld, R. (2002). Structure of coronavirus main proteinase reveals combination of a chymotrypsin fold with an extra alpha-helical domain. EMBO J. 21, 3213–3224.

[27] Koike, R., Ota, M. & Kidera, (2014). A hierarchical description and extensive classification of protein structural changes by motion tree. J. Mol. Biol. 426, 752–762.

[28] Moritsugu, K., Koike, R., Yamada, K., Kato, H. & Kidera, A. (2015). Motion tree delineates hierarchical structure of protein dynamics observed in molecular dynamics simulation. PLOS One. 10, Article e0131583.

[29] Tan, J.Z., Verschueren, K. H. G., Anand, K., Shen, J. H., Yang, M. J., Xu, Y. C., Rao, Z., Bigalke, J., Heisen, B., Mesters, J. R., Chen, K., Shen, X., Jiang, H. & Hilgenfeld, R. (2005). pH-dependent conformational flexibility of the SARS-CoV main proteinase (M-pro) dimer: Molecular dynamics simulations and multiple X-ray structure analyses. J. Mol. Biol. 354, 25–40.

[30] Chen, S., Chen, L. L., Tan, J. Z., Chen, J., Du, L., Sun, T., Shen, J., Chen, K., Jiang, H. & Shen, X. (2005). Severe acute respiratory syndrome coronavirus 3C-like proteinase N terminus is indispensable for proteolytic activity but not for enzyme dimerization - Biochemical and thermodynamic investigation in conjunction with molecular dynamics simulations. J. Biol. Chem. 280, 164–173.

[31] Hsu, W. C., Chang, H. C., Chou, C. Y., Tsai, P. J., Lin, P. I. & Chang, G. G. (2005). Critical assessment of important regions in the subunit association and catalytic action of the severe acute respiratory syndrome coronavirus main protease. J. Biol. Chem. 280, 22741–22748.

[32] Wei, P., Fan, K.Q., Chen, H., Ma, L., Huang, C. K., Tan, L., Xi, D., Li, C., Liu, Y., Cao, A. & Lai, L. (2006). The N-terminal octapeptide acts as a dimerization inhibitor of SARS coronavirus 3C-like proteinase. Biochem. Biophys. Research Comm. 339, 865–872.

[33] Lee, R.A., Razaz, M. & Hayward, S. (2003). The DynDom database of protein domain motions. Bioinformatics, 19, 1290–1291.

[34] Otto, H. H. & Schirmeister, T. (1997). Cysteine proteases and their inhibitors. Chemical Reviews. 97, 133–171.

[35] Stroud, R. M., Kossiakoff, A. A. & Chambers, J. L. (1977). Mechanisms of zymogen activation. Annu. Rev. Biophys. Bioeng. 6, 177–193.

[36] Wallace, A. C., Laskowski R. A. & Thornton, J. M. (1995). LIGPLOT - a program to generate schematic diagrams of protein ligand interactions. Protein Eng. 8, 127–134.

[37] Barrila, J., Gabelli, S. B., Bacha, U., Amzel, L.M. & Freire, E. (2010). Mutation of Asn28 disrupts the dimerization and enzymatic activity of SARS 3CL(pro). Biochemistry. 49, 4308–4317.

[38] Shi, J. H., Sivaraman, J. & Song, J. X. (2008). Mechanism for controlling the dimer-monomer switch and coupling dimerization to catalysis of the severe acute respiratory syndrome coronavirus 3C-like protease. J. Virol. 82, 4620–4629.

[39] Lee, T. W., Cherney, M. M., Huitema, C., Liu, J., James, K. E., Powers, J. C., Eltis L. D. & James, M. N. (2005). Crystal structures of the main peptidase from the SARS coronavirus inhibited by a substrate-like aza-peptide epoxide. J. Mol. Biol. 353, 1137–1151.

[40] Lee, T.W., Cherney, M. M., Liu, J., James, K. E., Powers, J. C., Eltis, L. D., James M.N. (2007). Crystal structures reveal an induced-fit binding of a substrate-like aza-peptide epoxide to SARS coronavirus main peptidase. J Mol Biol. 366, 916–932.

[41] Cheng, S. C., Chang, G. G. & Chou, C. Y. (2010). Mutation of Glu-166 blocks the substrate-induced dimerization of SARS coronavirus main protease. Biophys. J. 98, 1327–1336.

[42] Lim, L. Z., Shi, J. H., Mu, Y. G. & Song, J. X. (2014). Dynamically-driven enhancement of the catalytic machinery of the SARS 3C-like protease by the S284-T285-I286/A mutations on the extra domain. PLOS One. 9, Article e101941.

[43] Vuong, W., Khan, M. B., Fischer, C., Arutyunova, E., Lamer, T., Shields, J., Saffran H. A., McKay, R. T., van Belkum, M. J., Joyce, M. A., Young, H. S., Tyrrell, D. L., Vederas, J. C. & Lemieux, M. J. (2020). Feline coronavirus drug inhibits the main protease of SARS-CoV-2 and blocks virus replication. Nat. Commun. 11, Article 4282.

[44] Zhang, L. L., Lin, D. Z., Sun, X. Y. Y., Curth, U., Drosten, C., Sauerhering, L., Becker, S., Rox, K. & Hilgenfeld, R. (2020). Crystal structure of SARS-CoV-2 main protease provides a basis for design of improved alpha-ketoamide inhibitors. Science. 368, 409–412.

[45] Shi, J. & Song, J. (2006). The catalysis of the SARS 3C-like protease is under extensive regulation by its extra domain. FEBS J. 273, 1035–1045.

[46] Chou, C. Y., Chang, H. C., Hsu, W. C., Lin, T. Z., Lin, C. H. & Chang, G. G. (2004). Quaternary structure of the severe acute respiratory syndrome (SARS) coronavirus main protease. Biochemistry. 43, 14958–14970.

[47] Lin, P. Y., Chou, C. Y., Chang, H. C., Hsu, W. C. & Chang, G. G. (2008). Correlation between dissociation and catalysis of SARS-CoV main protease. Arch. Biochem. Biophys. 472, 34–42.

[48] Kobayashi, C., Matsunaga, Y., Koike, R., Ota, M. & Sugita, Y. (2015). Domain motion enhanced (DoME) model for efficient conformational sampling of multidomain proteins. J. Phys. Chem. B. 119, 14584–14593.

[49] Koike, R., Takeda, S., Maeda, Y. & Ota M. (2016). Comprehensive analysis of motions in molecular dynamics trajectories of the actin capping protein and its inhibitor complexes. Proteins. 84, 948–956.

[50] Moritsugu, K., Ito, T. & Kidera, A. (2019). Allosteric response to ligand binding: Molecular dynamics study of the N-terminal domains in IP3 receptor. Biophys Physicobiol. 16, 232–239.

