## Supporting Information for "Allosteric regulation of 3CL protease of SARS-CoV-2 and SARS-CoV observed in the crystal structure ensemble"

* Akinori Kidera

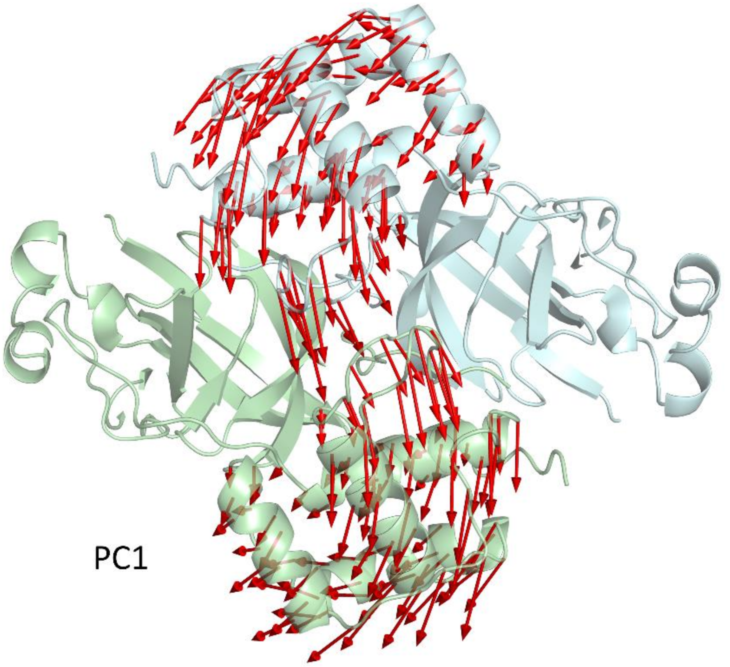


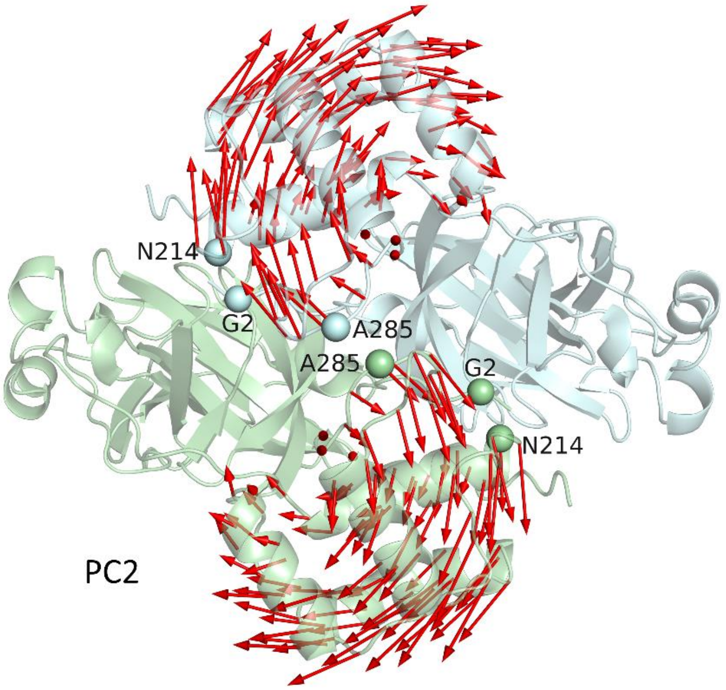


**Figure S1.** The first (top) and second (bottom) principal components (PCs) for domain III in the dimeric form (red arrows): chain A (light green) and chain B (light cyan). The variances explained by the PCs are 0.30 (PC1) and 0.25 (PC2). PC1(domain III dimer) represents the symmetry breaking heterogeneity in the crystal environment. Along PC2(domain III dimer), domain III separates from its counterpart of the other protomer, as does Ala285A from Ala285B (A and B denote two protomers) and Asn214 from Gly2.

A

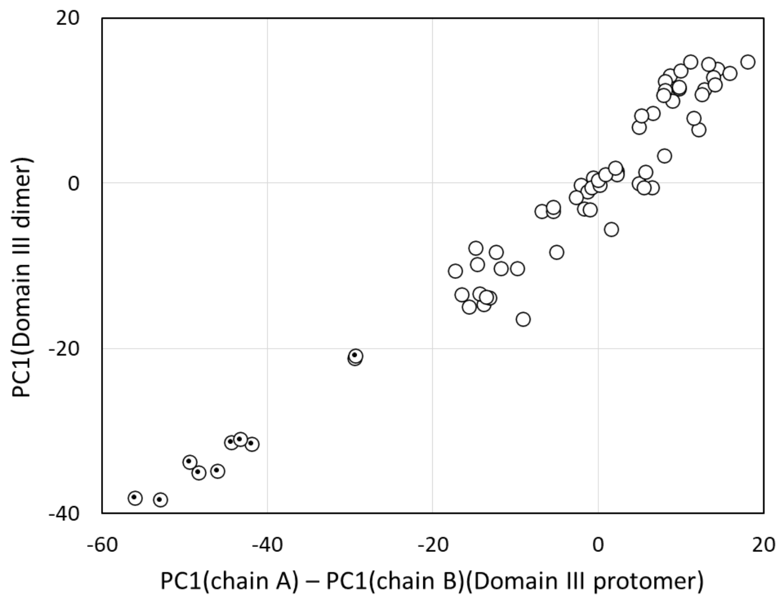


B

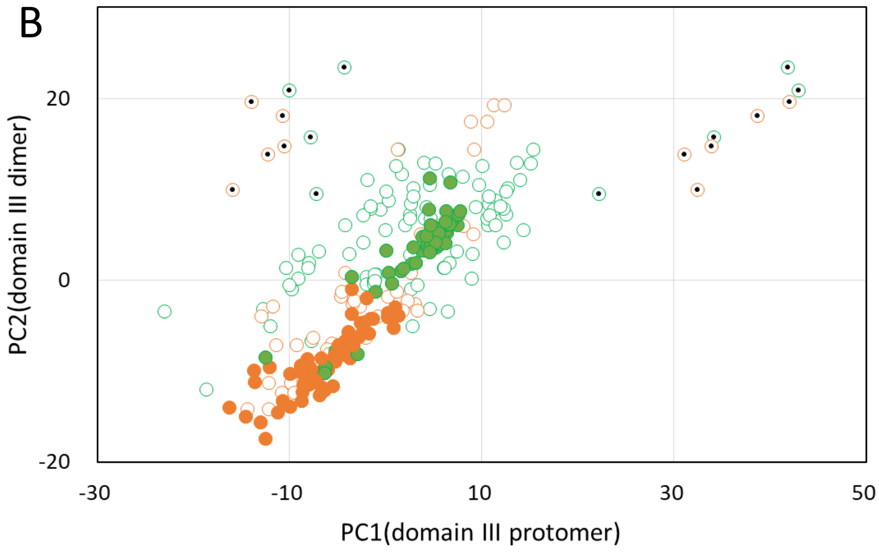


**Figure S2.** (A) PC1(domain III dimer) is well described by the difference of PC1(domain III protomer of chain A) – PC1(domain III protomer of chain B). The chain names used here are those of the original assignment in PDB. The inner dots indicate the chains with the space group P 1 $2_1$ 1 that have large heterogeneity in the environment of domain III (see SI Text 2). (B) PC2(domain III dimer) is plotted against PC1(domain III protomer): SARS-CoV $3CL^{pro}$ (green) and SARS-CoV-2 $3CL^{pro}$ (orange). The filled green and orange circles are those of the symmetric dimers that have a single value of PC1(domain III protomer) and show better linear relationships between the two variables. In contrast, the asymmetric dimers have different values of PC1(domain III protomer) and they have the following approximate relationship: PC2(domain III dimer) ~ [PC1(domain III protomer of chain A) + PC1(domain III protomer of chain B)]/2. Among the asymmetric dimers, the largest differences are found in the chains with the space group P 1 $2_1$ 1 with large heterogeneous crystal packing marked by the inner dots as in (A). The difference between the two species of $3CL^{pro}$ is discussed in the last section.

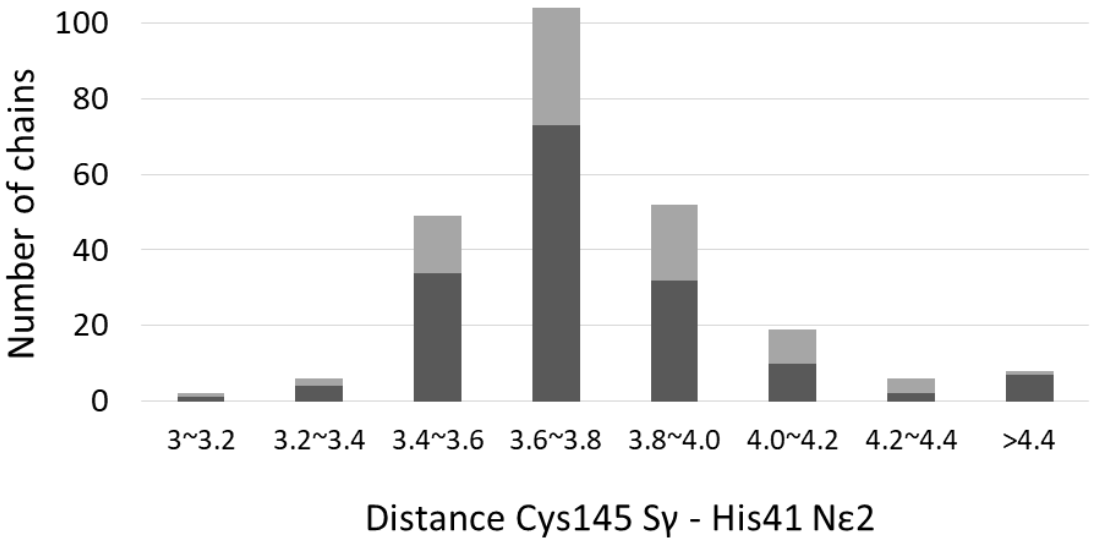


**Figure S3.** Histogram of the distance between Cys145 Sγ and His41 Nε2 based on 246 chains (12 chains with mutations at either Cys145 or His41); ligand-bound chains (dark gray) and ligand-free chains (light gray). The average distance is 3.7 ± 0.3Å; 83% of the chains are within the standard deviation, with no distinction between ligand-free and ligand-bound chains. Those with distances > 4.4 Å are mostly chains with ligand molecules inserted between Cys145 and His41.

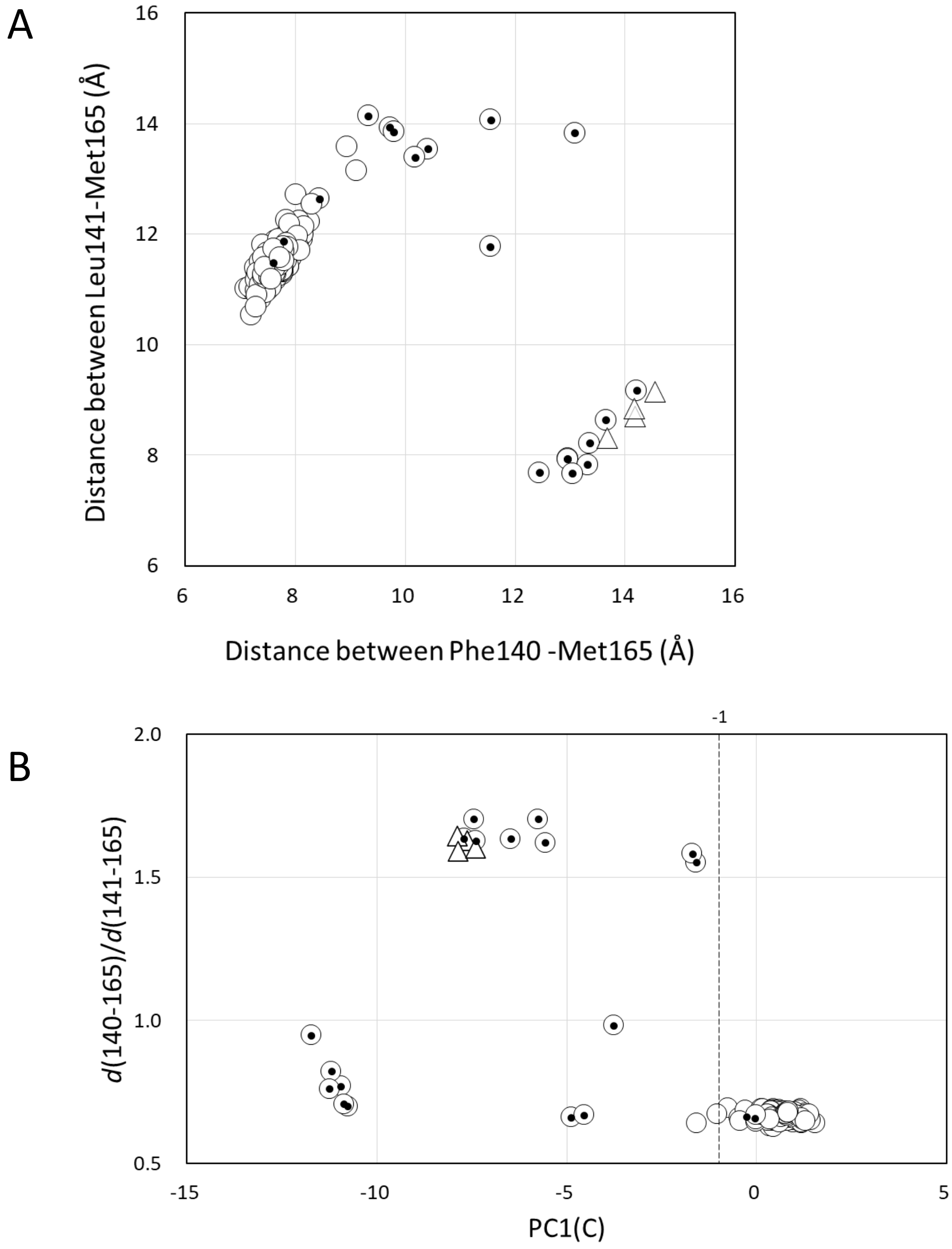


**Figure S4.** (A) Distance Phe140 Cβ-Met165 Cα vs. distance Leu141 Cβ-Met165 Cα; dimers (circles) and monomers (triangles). The inner dots are those of the collapsed chains. (B) The ratio of the two distances, $d(140–165)/d(141–165)$, is plotted against PC1(C). The vertical line at −1 indicates the threshold separating the active and collapsed states. The symbols are the same as those used in (A).

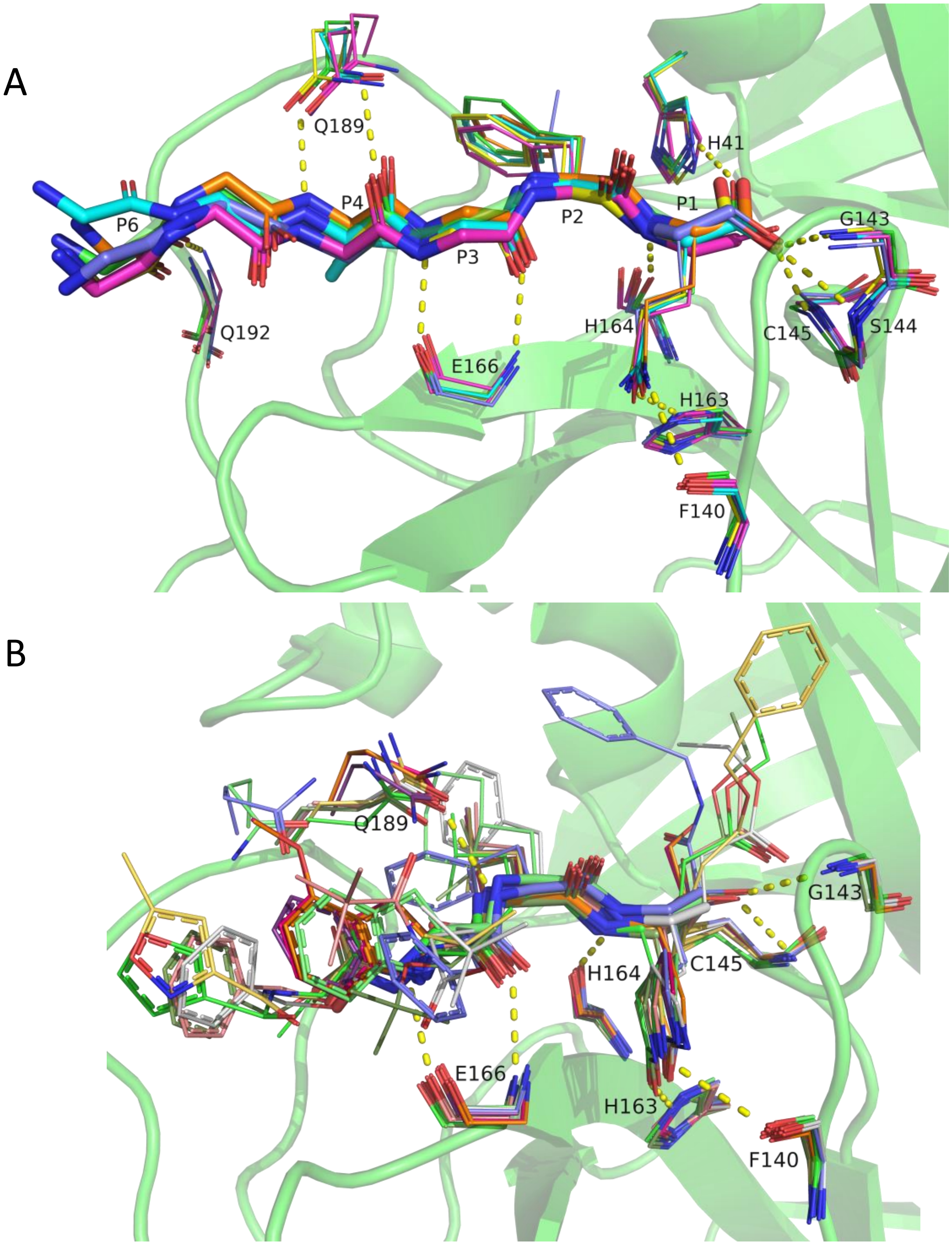
A
Q189
H41
P6
P4
P2
P1
G143
P3
H164
C145
S144
Q192
E166
H163
F140
B
Q189
G143
H164
C145
E166
H163
F140

C

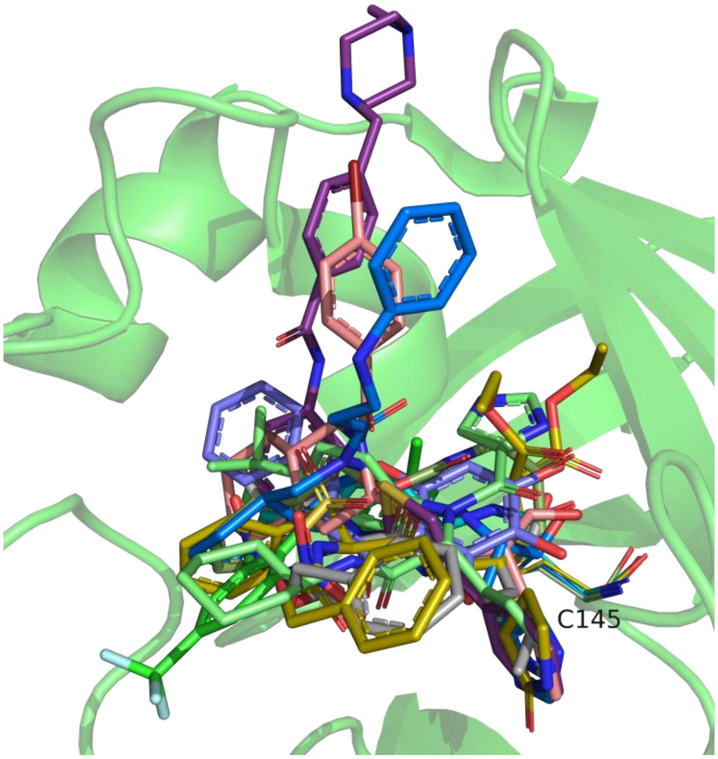


D

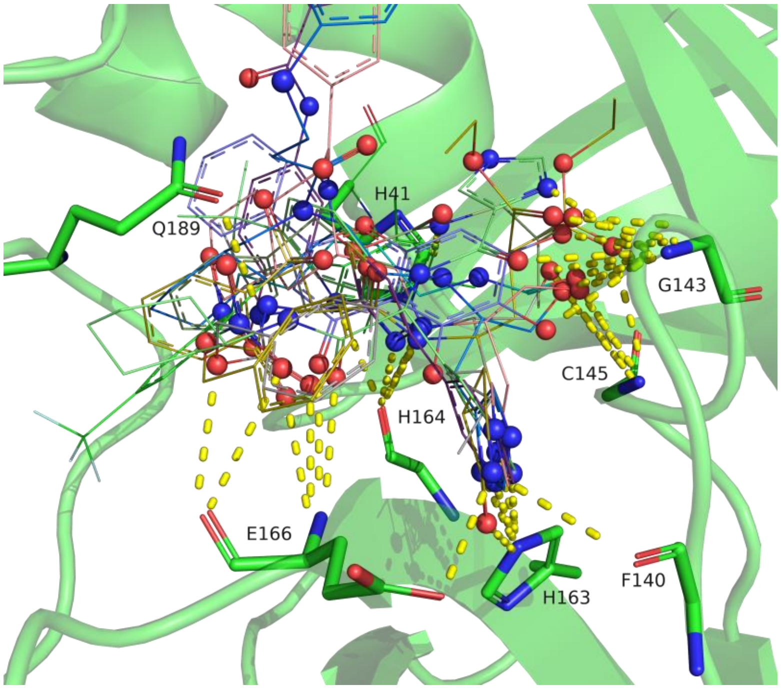

**Figure S5.** (A) The binding poses of the peptide substrates drawn after superposition at the core region. The main-chains of the substrates (six residues P6-P1) are drawn as sticks, whereas the side-chains of the substrates and the protein residues (either the main-chain or the side-chain) involved in ligand recognition are drawn as lines. Representative polar contacts are illustrated by yellow broken lines. Gly143 and Cys145 form the oxyanion hole to bind the carbonyl group at P1. Phe140 and His163 bind the sequence-specific side-chain of Gln at P1. The side-chain of P2 (Phe or Leu in these entries) is surrounded by a hydrophobic environment. The color scheme is as follows: PDB:1z1jA, green (the bound substrate: the C-terminal of $3CL^{pro}$, SGVTFQ); 2q6gA, slate (the C-terminal of Nsp4C or the upstream of the N-terminus of $3CL^{pro}$, TSAVLQ, for which the C-terminal five residues are not drawn for clarity); 5b6oA, cyan (the C-terminal of $3CL^{pro}$ for which the C-terminal four residues are not drawn), 6xoaA, orange (the C-terminal of $3CL^{pro}$); 7joyB, yellow (the C-terminal of $3CL^{pro}$), and 7khpB, magenta (the C-terminal of $3CL^{pro}$ covalently bound to Cys145). (B) The representative binding poses of peptide-mimic compounds. In these complexes, the recognition at the subsites is almost the same as that of the substrates: Gly143 and Cys145 also bind the carbonyl group at P1; Phe140 and His163 recognize the Gln-mimic moieties at P1; and the hydrophobic side-chains at P2 occupy almost the same positions as those of the substrate. However, the other hydrophobic side-chains are more randomly oriented, revealing less definite hydrophobic pockets other than the S2 site. The color scheme is as follows: PDB:2amdA, green; 2zu4, salmon; 3tiu, white; 5n19, slate; 6lze, pale green; 6xhlA, orange; 6xhmA, warm pink; 7bqy, yellow-orange; 7c8r, smudge; and 7d1mA, violet-purple. (C) The representative binding poses of nonpeptide compounds drawn as sticks. Cys145 is drawn as lines as the reference position. The color scheme is as follows: PDB:2gz7, green; 3d62, smudge; 3szn, olive; 4twwA, salmon; 5c5n, marine; 6m2nA, slate; 6w63, pale green; 7aku, gray; 7buy, teal; and 7ju7, violet. (D) Same as (C), but the ligands are drawn as lines, whereas N and O atoms are depicted as spheres, and the polar contacts with the subsite residues (stick) are represented by yellow dotted lines. The polar atoms of the ligands that have no interactions with the subsites are mostly exposed to the solvent.

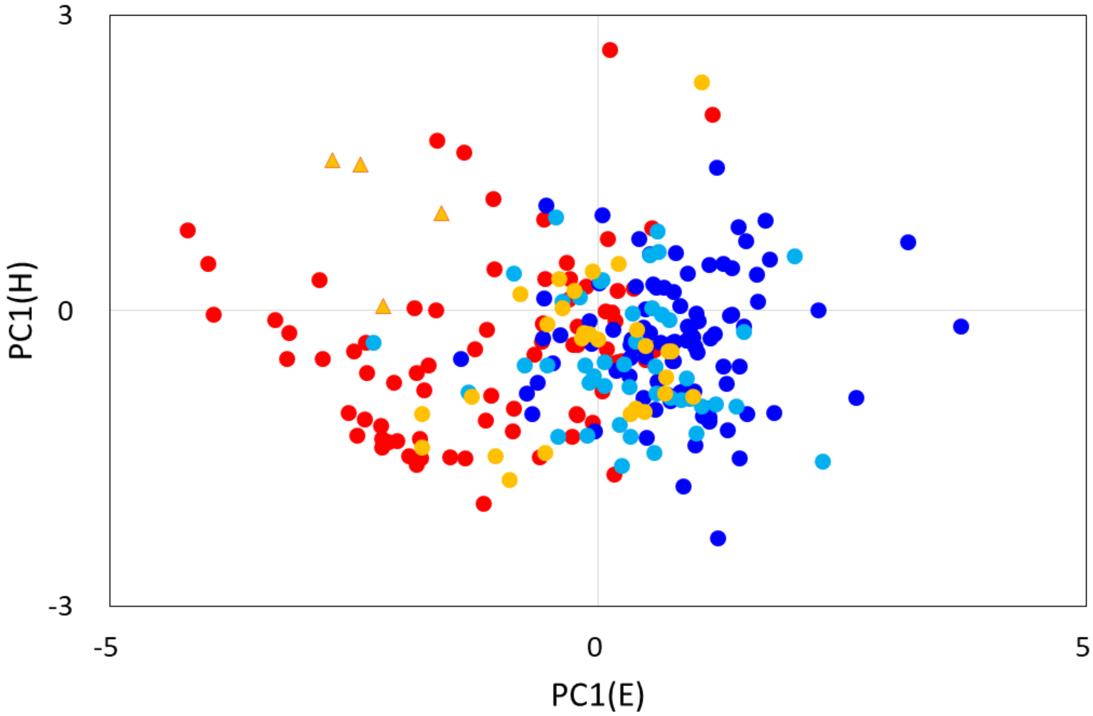


**Figure S6.** A scatter plot of PC1(E) vs. PC1(H). Same as Fig. 4A, but PC1(L) is replaced by PC1(H). The color scheme for the binding poses is as follows: the ligand-free, red; “w,” light orange; “E,” cyan; “EL,” blue; and the monomeric structure, orange triangle. Binding pose dependence is found along PC1(E), but such dependence is not found along PC1(H).

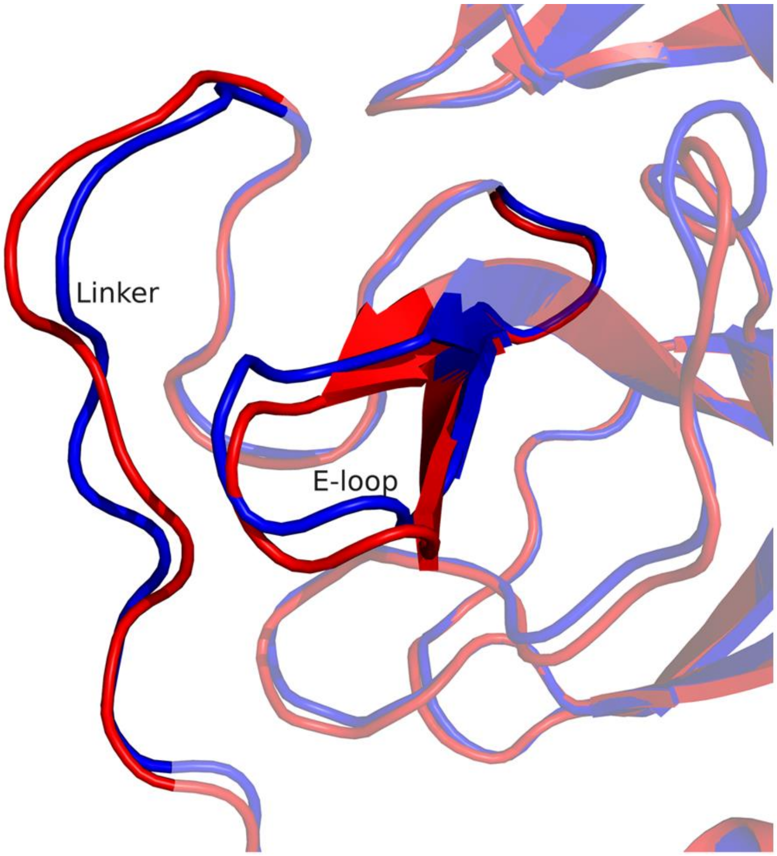


**Figure S7.** The representative structures with different values of PC1(E)+PC1(L); PDB:2qc2 (red, PC1(E)+PC1(L) = 2.179) and 7cwb (blue, −5.889) drawn after superposition at the core region. Only the E-loop and Linker are drawn without transparency. 2qc2 is in the open conformations of the E-loop and The N-terminal part of the Linker, whereas 7cwb is in the closed conformations.

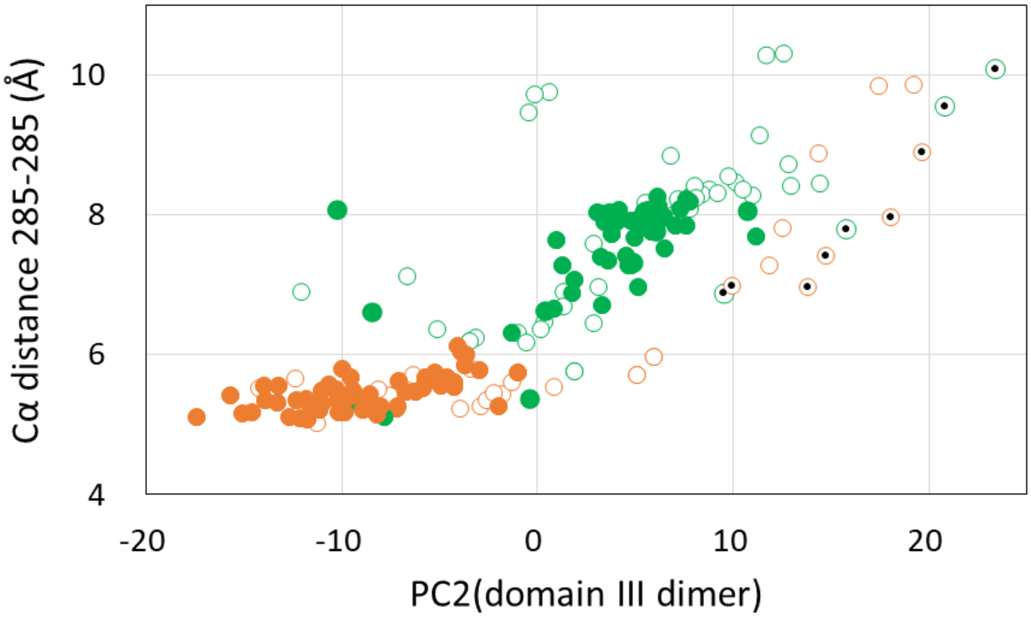


**Figure S8.** Correlation between PC2(domain III dimer) and distance 285-285: SARS-CoV 3CL$^{pro}$ (green) and SARS-CoV-2 3CL$^{pro}$ (orange). Filled circles indicate the symmetric dimers. Circles with inner dots are those of highly heterogeneous chains of the space group P 1 $2_1$ 1 (see SI Text 2).

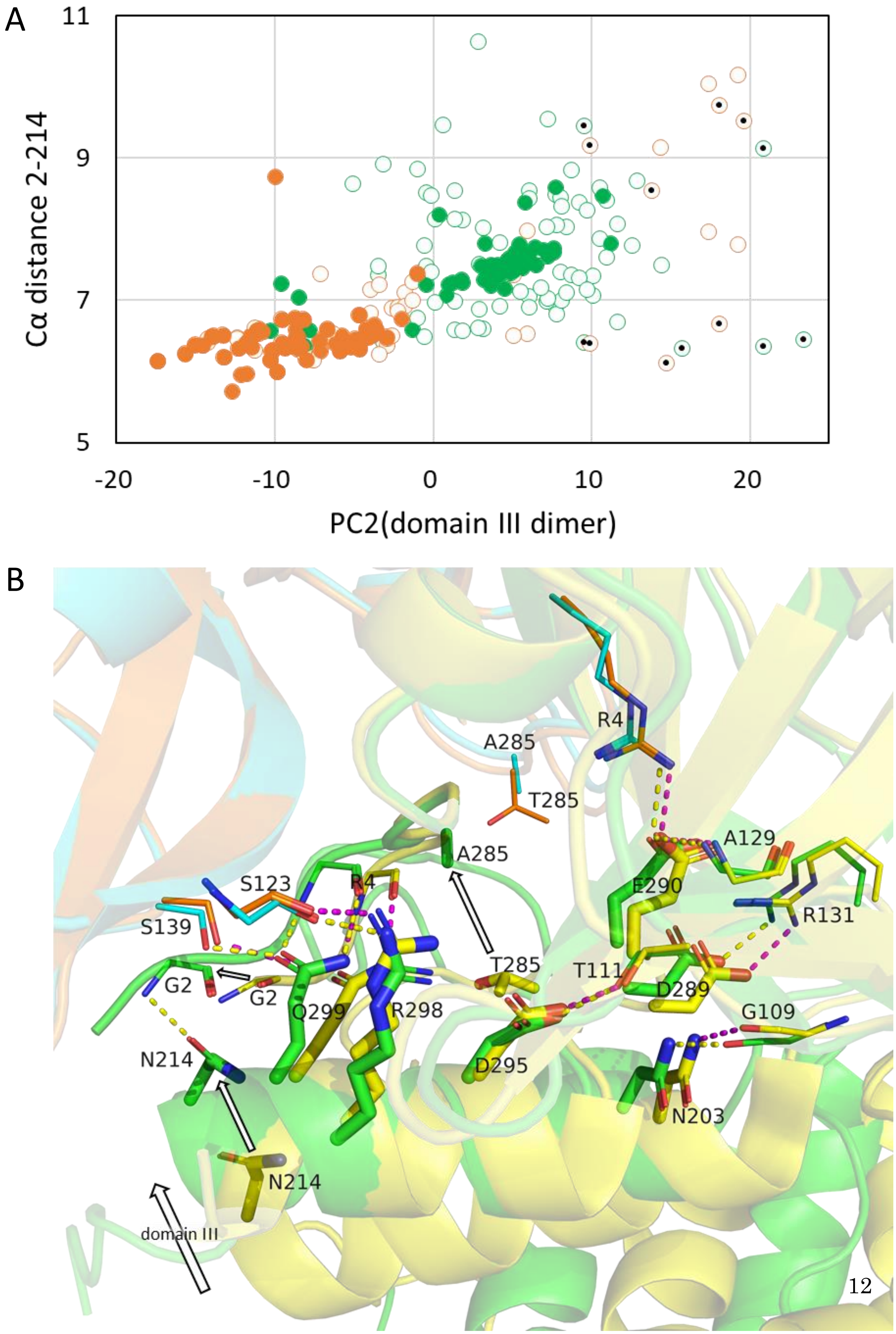
A
11
9
7
5
-20
-10
0
10
20
Cα distance 2-214
PC2(domain III dimer)
B
R4
A285
T285
A285
A129
S123
R4
E290
S139
R131
T285
T111
G2
G2
D289
Q299
R298
G109
N214
D295
N203
N214
domain III

**Figure S9.** (A) Correlation between PC2(domain III dimer) and Cα distance 2-214: SARS-CoV $3CL^{pro}$ (green) and SARS-CoV-2 $3CL^{pro}$ (orange). The filled circles are those of the symmetric dimers, showing linear relationship between the two variables (see the caption of Fig. S2B). The inner dots are those of the space group P 1 $2_1$ 1, which have large heterogeneity in the crystal environment. (B) The interdomain interactions with domain III in the representative structures, drawn after superposition at the core region of chain A. The major interdomain interactions are depicted here. The figure compares PDB:6lu7 (SARS-CoV-2; chain A (cyan) and chain B (green); distance 285-285 = 5.314 Å; PC2(domain III dimer) = −11.704; and distance 214-2 = 6.421 Å) with 6xhl (SARS-CoV; chain A (orange) and chain B (yellow); distance 285-285 = 9.541 Å; PC2(domain III dimer) = 20.824; and distance 214-2 = 9.131 Å (chain B)). 6xhl has an uncharged N-terminus with Gly0, but Gly0 and Ser1 are missing in the model. Here, chain A, for which the core region is superimposed, is not shown except for the interacting residues; two chain Bs are compared. Polar contacts are shown by dotted lines in yellow (6lu7) and magenta (6xhl). The arrows indicate the activating motion of Asn214 and Thr(Ala)285 along with the motion of domain III. At the same time, Gly2 in the N-finger shifts position along with the formation of H140-1.

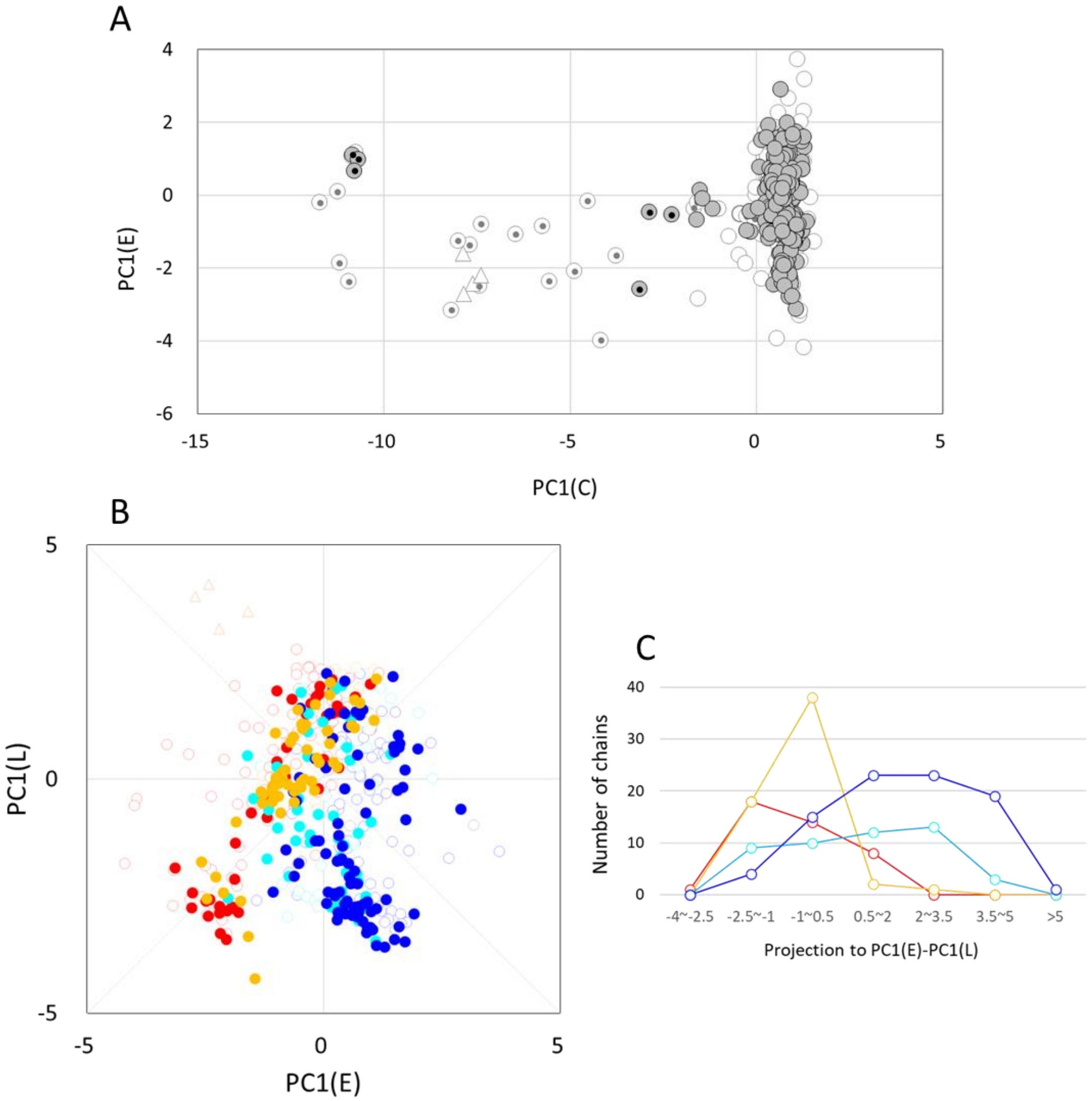


**Figure S10.** (A) The same figure as Fig. 3B, but for the entries listed in Dataset S4 and S5. Gray-shaded circles are for the new entries and thin open circles are for the entries in Dataset S1 and S2. The C-loop conformations show basically the same distribution as in Fig. 3B. (B) The same figure as Fig. 4A, but for the new entries. Colored circles are for those listed in Dataset S4 and S5 and thin open circles are for those in Dataset S1 and S2. (C) The projection of the data in Fig. S10B on PC1(E) – PC1(L). The ligand-size dependent distribution along PC1(E)-PC1(L) is also found in these entries.

Table S1 Correlation among the PC1's

| | C-loop | E-loop | H-loop | Linker |
|---|---|---|---|---|
| E-loop | 0.225 | | | |
| H-loop | 0.107 | 0.107 | | |
| Linker | -0.094 | 0.181 | 0.024 | |
| Domain III | -0.081 | 0.231 | 0.112 | 0.162 |

The values are correlation coefficients between each pair of PC1's.

### Supporting Text

#### *Supporting Text 1. Principal Components of the domain III dimer*

The principal components (PCs) of the domain III dimer relative to the core region are explained here. The first principal component (PC1) exhibits two domains that move translationally to the same direction to break symmetry (Fig. S1; variance explained = 0.30). This symmetry breaking motion is explained by the difference between the two values of PC1(domain III protomer), calculated respectively for chains A and B, which indicate the heterogeneity of the crystal environment in each of domain III (Fig. S2A). Therefore, the symmetric dimers that have two-fold symmetry within the dimer have the PC1 value of zero. Conversely, PC2 exhibits each domain III rotating around the rotation axis normal to the plane of the figure with no translational motion (Fig. S1; variance explained = 0.25). PC2 correlates well with the motion within a protomer, PC1(domain III protomer), in the symmetric dimers; in the asymmetric dimers, PC2(domain III dimer) correlates approximately with the average of PC1(domain III protomer) for chains A and B (Fig. S2B). As shown in Fig. S1, the motions along PC2(domain III dimer) separate Thr(Ala)285 from the counterpart of the other protomer and Asn214 from Gly2. Therefore, PC2 changes distance 285-285 as well as distance 214-2. This is the origin of the coupling between domain III and the C-loop via the N-finger (see the section entitled “Influences of T285A mutation ...” for more details).

#### *Supporting Text 2. The heterogeneous environment in the crystal structures of the space group P 1 $2_1$ 1*

Crystal packing is one of the factors affecting crystal structures. In $3CL^{pro}$, flexible domain III susceptibly changes configuration depending on crystal packing. Packing effects are evidently observed in the asymmetric dimers, in which two protomers are not related by the symmetry operation but have independent structures. In figures showing the behavior of the domain III dimers (Figs. S2A, S2B, S8, and S9A), it is notable that nine entries exhibit exceptionally large heterogeneity (PDB:6w2a, 6xhl, 6xhn, and 6xho of SARS-CoV $3CL^{pro}$; 6xbg, 6xhm, 6xmk, 7d1m, and 7jkv of SARS-CoV-2 $3CL^{pro}$). All of these are ligand-bound and belong to one of three different crystal forms of monoclinic P 1 $2_1$ 1.

#### *Supporting Text 3. Structural variation in the collapsed state of the C-loop*

It has been argued that the most typical collapsed structure is found in the monomeric state, which occurs in the crystals of dimerization-deficient mutants (PDB:2pwx, 2qcy, 3f9e, and 3m3t (1-3)). These structures are characterized by the large negative values of PC1(C) and the absence of the four intraprotomer hydrogen bonds listed in Table 1. Furthermore, the side-chain of Leu141 occupies the position of the side-chain of Phe140 situated in the active state in which His163 and His172 form tight hydrophobic packing (the importance of the hydrophobic interactions among Phe140, His163, and His172 is confirmed in the mutant F140A (PDB:3f9f, 3f9g, and 3f9h) in which the C-loop is collapsed). In addition, the intra main-chain HBs of the C-loop, namely HB 138-141 and HB 139-142, are formed to make a $3_{10}$-helix (residues 139–141).

To analyze the collapsed structure in the dimer, we first examined the position of the side-chains, Phe140 and Lue141, by measuring the distance between Cβ of Phe140/Leu141 and Cα of Met165 located in the stable domain II, d(140–165) and d(141–165). In the active state, it is expected that d(140–165) < d(141–165), whereas in the collapsed state, d(140–165) > d(141–165). Figure S4A shows various cases: eight collapsed chains have similar side-chain positions as those of the monomeric structures; another eight are found to have d(141–165) that is same or larger than the active state distance; and three maintain the active state distance. These chains are not necessarily weakly collapsed structures. When the ratio d(140–165)/d(141–165) (a small ratio in the active state and a large ratio in the collapsed state) is plotted against PC1(C) (Fig. S4B), some chains overshoot the PC1(C) values of the monomeric structures without increasing the ratio. These plots illustrate that the collapsed state takes diverse conformations of the C-loop. Thus, the structural change of the C-loop between the active and collapsed states can be understood as a type of the order–disorder transition.

When examining the $3_{10}$-helix (residues 139–141), we realize that only one chain (PDB:1uj1B) has a $3_{10}$-helix stabilized by the main-chain HBs 138–141 and 139–142, similar to the monomeric structures (Dataset

S1 and S2). This implies that the $3_{10}$-helix is not a requisite for the side-chain orientations that appear in monomeric structures. If the collapsed structures in the monomeric form are the most stable conformations, we can postulate a significant barrier for exchanging the side-chain positions that keeps the protein in an unstable collapsed structure.

***Supporting Text 4. The structural analysis of MERS-CoV 3CL$^{pro}$***

As a reference to the results of the structural analysis of SARS-CoV and SARS-CoV-2 3CL$^{pro}$, we also analyzed the crystal structures of homologous MERS-CoV 3CL$^{pro}$, 3C-like protease of Middle East respiratory syndrome–related coronavirus (2012). Protein Data Bank contains 16 PDB entries (33 independent chains) of MERS-CoV 3CL$^{pro}$, whose sequence has 51% identity to SARS-CoV-2 3CL$^{pro}$. However, as shown in the sequence alignment in Fig. S11, many of the important residues in the interactions are not mutated. Three amino acid differences, R298M, Y126F and A285T, are considered to affect the structure of the molecule (see Fig. S11). The experiments revealed that MERS-CoV 3CL$^{pro}$ is a weakly associated dimer [Kd = 7.7 μM (4) and 7.8 μM (5); these Kd values are much larger than those for SARS-CoV 3CL$^{pro}$, Kd = 0.7 μM (4) and 0.06 μM (5)], and is dimerized upon substrate binding (5).

The crystal structure of MERS-CoV 3CL$^{pro}$ (PDB: 4rsp) was compared with the structure of SARS-CoV-2 3CL$^{pro}$ (PDB:6lu7) using Motion Tree for pairwise comparison (6). Figure S12 shows the result of the comparison. The coordinate shifts mostly occur in the moving clusters found in SARS-CoV and SARS-CoV-2 3CL$^{pro}$. Largest shift is found in domain III. This is caused by the amino acid difference A285T (see below), as well as many mutations accumulated in domain III (Fig. S11).

Supporting data S7 summarizes the structural characteristics of the 16 PDB entries of MERS-CoV 3CL$^{pro}$. Because of the low sequence identity, we did not calculate the principal components of the moving clusters using the mode vectors derived from SARS-CoV and SARS-CoV-2 3CL$^{pro}$. However, the other quantities were calculated according to the alignment of Fig. S11. From this table, the followings were found:
(1) Y129F disables the formation of HB 142-129, and may destabilize the active conformation of the C-loop.
(2) The appended amino acids at the N-terminus disables the formation of HB 143-1. However, most of the chains without HB 143-1 have HB 142-2 instead. Similarly, the chains without HB 141-175 have HB 140-175. When HB 143-1 is absent, however, HB 142-2 cannot recover the inter-domain HB 2-217 and the distance between the two T285's increases. These hydrogen bonds, HB 142-2 and HB 140-175, are rarely observed in SARS-CoV and SARS-CoV-2 3CL$^{pro}$.
(3) The mutation A285T increases the distance between the two T285's as observed in SARS-CoV and SARS-CoV-2 3CL$^{pro}$; the average distance is 7.53 Å which is close to the average value of SARS-CoV 3CL$^{pro}$ (T285), 7.66 Å, which is larger than the average value of SARS-CoV-2 3CL$^{pro}$ (A285), 5.78 Å.
(4) The C-loop conformations observed in the side-chain positions, i.e., the distances 168Cα-143Cβ and 168Cα-144Cβ, appear to be similar to those of SARS-CoV and SARS-CoV-2 3CL$^{pro}$; the average distances are 7.41 Å and 11.00 Å for 168Cα-143Cβ and 168Cα-144Cβ, respectively (the corresponding average values of SARS-CoV and SARS-CoV-2 3CL$^{pro}$ are 7.65 and 11.45, respectively).
(5) C chain of 4wmf is in the monomeric state in the crystal. Nevertheless, the C-loop conformation stays in the active state (HB 146-28 exists), though HB 141-175 and HB 142-129 do not exist and the side-chain conformations of F143 and L144 are slightly distorted. The interaction that stabilizes the active C-loop conformation in 4wmfC can be ascribed to the hydrogen bonds between C-loop and Linker/domain III (7). 4wmfC has the largely different configuration of domain III due to the recognition of the C-terminal by chain A. Due to this position of domain III, Linker and the long loop (276-293) comes close to C-loop to form the hydrogen bonds, HB 142-202, HB 142-282, and HB 145-199. These HB may prevent collapsing of C-loop.

Supporting data S8 summarizes the ligand interactions. The major binding sites found in SARS-CoV and SARS-CoV-2 3CL$^{pro}$ are almost exactly conserved in MERS-CoV 3CL$^{pro}$ except for the minor binding sites of the interface region, where MERS-CoV 3CL$^{pro}$ scarcely shows ligand binding.

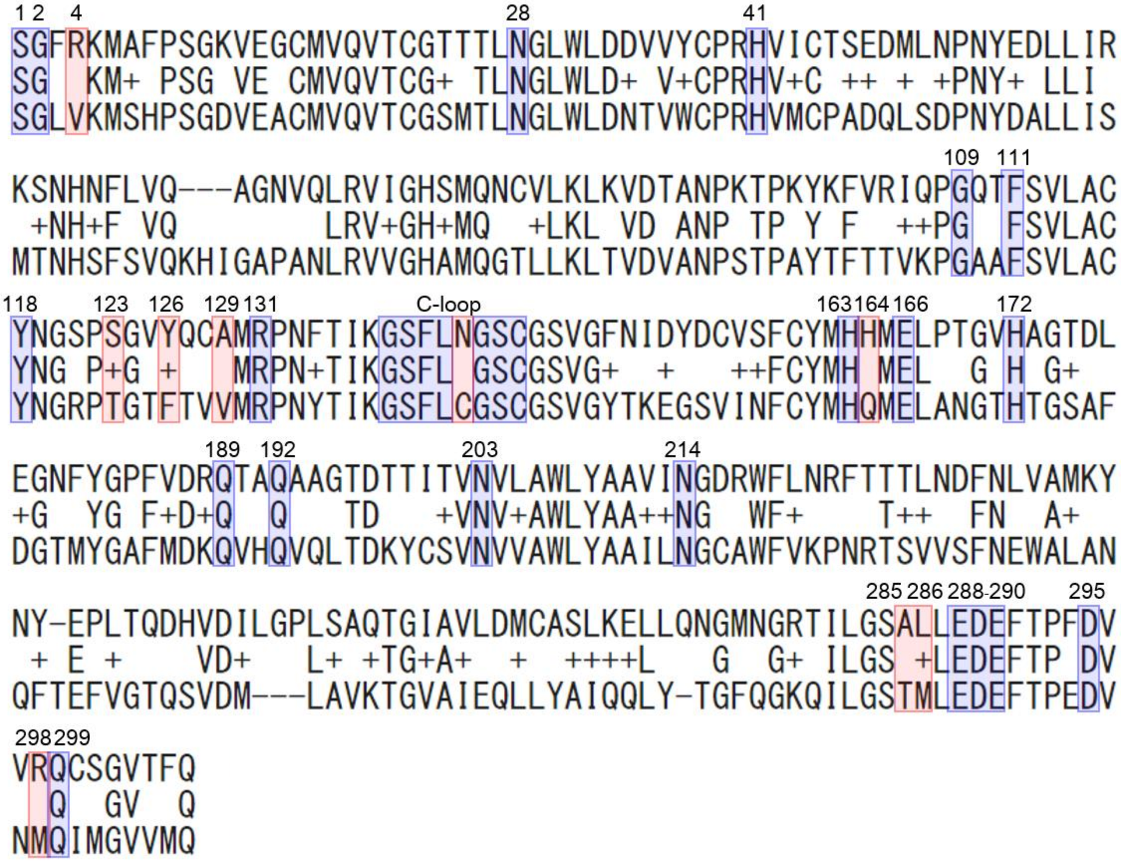


**Figure S11.** Sequence alignment between SARS-CoV-2 3CL$^{pro}$ and MERS CoV 3CL$^{pro}$ (51% identity; 157/310). Shaded amino acids are the residues having important interactions in SARS-CoV-2 3CL$^{pro}$ discussed in the text; blue shaded residues are not changed, and red shared ones are changed. The residue numbers are those of SARS-CoV-2 3CL$^{pro}$. R298M increases the dissociation constant of the two protomers by breaking the interprotomer HB 298A-123B (4). Y126F may destabilize the active conformation of the C-loop by breaking HB 126-139. A285T increase the distance between the interface of domain III as in the mutation A285T between SARS CoV-2 3CL$^{pro}$ and SARS CoV 3CL$^{pro}$ discussed in the text.

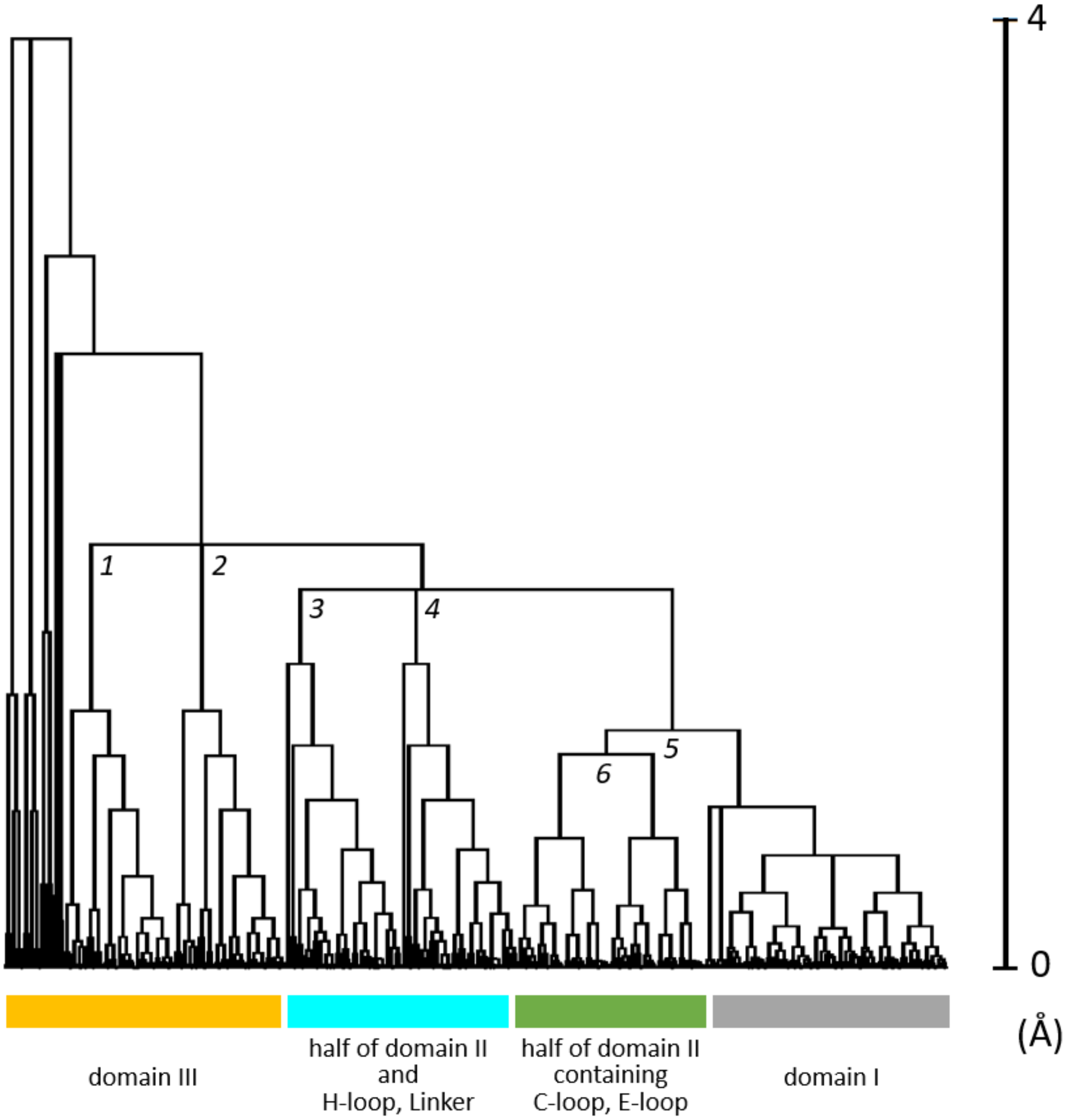


**Figure S12** Motion Tree comparing SARS-CoV-2 3CL$^{pro}$ (PDB: 6lu7) and MERS-CoV 3CL$^{pro}$ (PDB: 4rsp), the hierarchical clustering based on the absolute value of the difference in residue distance (4). The abscissa is the MT score representing the magnitude of the structural difference for each cluster. The tree is roughly separated into four regions, mostly corresponding to the moving clusters defined in SARS-CoV and SARS-CoV-2 3CL$^{pro}$. Both structures are symmetric dimers, and thus the tree have two equivalent nodes, one in chain A and the other in chain B (nodes *1* and *2*, and nodes *3* and *4*). However, nodes *5* and *6* appears only once because these nodes separate the interface region of the two chains into two clusters containing equivalent sets of residues. Nodes *1* and *2* separate domain III from domain I & II. Nodes *3* and *4* separate H-loop, Linker and a half of domain II (β-sheet located at the backside of domain II in Fig. 1) from domain I & II. Node *5* separates domain I from the rest of domain II containing C-loop and E-loop. Node *6* separates the interface region into chain A and chain B. The tallest branches at the left end of the tree are those for long flexible loops in domain III.

### SI References

1. S. Chen, T. Hu, J. Zhang, J. Chen, K. Chen, J. Ding, H. Jiang, X. Shen, Mutation of Gly-11 on the dimer interface results in the complete crystallographic dimer dissociation of severe acute respiratory syndrome coronavirus 3C-like protease: crystal structure with molecular dynamics simulations. *J. Biol. Chem.* **283**, 554-564 (2008).
2. J. Shi, J. Sivaraman, J. Song, Mechanism for controlling the dimer-monomer switch and coupling dimerization to catalysis of the severe acute respiratory syndrome coronavirus 3C-like protease. *J. Virol.* **82**, 4620-9 (2008).
3. T. Hu, Y. Zhang, L. Li, K. Wang, S. Chen, J. Chen, J. Ding, H. Jiang, X. Shen, Two adjacent mutations on the dimer interface of SARS coronavirus 3C-like protease cause different conformational changes in crystal structure. *Virology* **388**, 324-34 (2009).
4. B.-L. Ho, S.-C.Cheng, L. Shi, T.-Y. Wang , K-I Ho , C.-Y. Chou. Critical assessment of the important residues involved in the dimerization and catalysis of MERS coronavirus main protease. PLoS ONE 10, e0144865 (2015).
5. S. Tomar, M.L. Johnston, S.E. St John, H.L. Osswald, P.R. Nyalapatla, L.N. Paul, A.K. Ghosh, M.R. Denison, A.D. Mesecar. Ligand-induced dimerization of Middle East respiratory syndrome (MERS) coronavirus nsp5 protease (3CLpro): Implication for nsp5 regulation and the development of antivirals. J. Biol. Chem. 290, 19403-22 (2015).
6. R. Koike, M. Ota, A. Kidera, Hierarchical description and extensive classification of protein structural changes by Motion Tree. *Journal of Molecular Biology* **426**, 752-762 (2014).
7. D. Needle, G.T. Lountos, D.S. Waugh. Structures of the Middle East respiratory syndrome coronavirus 3C-like protease reveal insights into substrate specificity. Acta Cryst. D, 71 1102-11 (2015).
