## Supplementary material for "Allosteric regulation of 3CL protease of SARS-CoV-2 and SARS-CoV observed in the crystal structure ensemble": SI Datasets

Dataset S1 Summary of the ligand-bound entries.

| species | protein | | | | ligand | | | | comments | | | | | | | | | | | | | | | | | | | | | | | | | | | | | dimer | | |
|---|---|---|---|---|---|---|---|---|---|---|---|---|---|---|---|---|---|---|---|---|---|---|---|---|---|---|---|---|---|---|---|---|---|---|---|---|---|---|---|---|
| | | | | | | | | | | PC1 | | | | | C-loop conformation | | | | | | | | | PC1 | | | | | C-loop conformation | | | | | | | | | | | |
| | PDB ID | mutation | residues added to N-term | missing N-terminal residues# | name in PDB | name by authors | peptide or peptide mimic$ | Ki, IC$_{50}$*, EC$_{50}$$^{+}$ (μM) | | C-loop | E-loop | H-loop | Linker | Domain III | main-chain HB within 138-144 | HB 138-172& | HB 139-126& | HB 140-1& | HB 141-118& | HB 143-28& | distance 165Cα-140Cβ | distance 165Cα-141Cβ | distance 2-214 | C-loop | E-loop | H-loop | Linker | Domain III | main-chain HB within 138-144 | HB 138-172& | HB 139-126& | HB 140-1& | HB 141-118& | HB 143-28& | distance 165Cα-140Cβ | distance 165Cα-141Cβ | distance 2-214 | PC1 Domain III | PC2 Domain III | distance between two 285 Cα's |
| SARS-CoV | 1uk4 | | | SG of A | 5-mer peptide of inhibitor | hexa-peptidyl CMK inhibitor | P | 2000* | Cbz-VNSTLQ-CMK becomes NSTLQ-C-C145 in the complex. Cbz-V is not modeled. | 0.554 | -0.087 | -0.725 | 1.827 | 13.677 | | 1 | 1 | 1 | 1 | 1 | 8.311 | 12.537 | | -5.743 | -0.861 | -0.055 | -0.542 | 3.948 | 141O-143N | 0 | 0 | 0 | 0 | 0 | 13.049 | 7.664 | | 11.47 | 12.946 | 8.407 |
| | 1wof | | GPLGS | G(-5) of A, GPL of B | I12 | N1 | P | 10.7 | | 0.642 | 1.136 | -1.080 | 0.003 | 10.680 | 141O-144N | 1 | 1 | 1 | 1 | 1 | 7.590 | 11.529 | 8.547 | 0.123 | -0.472 | -0.523 | -1.754 | 2.011 | 141O-144N | 1 | 1 | 0 | 1 | 1 | 7.684 | 11.473 | 6.905 | 13.014 | 6.032 | 7.907 |
| | 1z1j_A | C145A | | | C-term of another chain A | | P | | Chain A binds a symmetrically related A chain (301-306) while chain B is in free state. C145A makes the protein inactive. | 1.129 | 0.756 | -0.317 | 2.256 | 6.628 | 141O-144N | 0 | 1 | 1 | 1 | 1 | 7.215 | 10.533 | 8.134 | | | | | | | | | | | | | | | 8.53 | 1.861 | 5.767 |
| | 2a5i | | | | AZP | Aza-peptide epoxide (APE) | P | 18 | | 0.310 | 1.442 | -1.495 | 0.703 | 2.678 | 141O-144N | 1 | 1 | 1 | 1 | 1 | 8.069 | 11.768 | 7.296 | | | | | | | | | | | | | | | -0.001 | 1.775 | 6.885 |
| | 2a5k | | A | A of A,B | AZP | Aza-peptide epoxide (APE) | P | | | 0.313 | 0.878 | -0.808 | -2.591 | 4.083 | 141O-144N | 1 | 1 | 0 | 1 | 1 | 7.886 | 11.504 | 7.595 | 0.284 | 0.461 | -0.885 | 0.783 | 4.706 | 141O-144N | 1 | 1 | 0 | 1 | 1 | 7.885 | 11.684 | 7.707 | 0.64 | 6.765 | 8.839 |
| | 2alv | | | S | CY6 | 4 | P | 70* | Mutations of T304V,F305V,Q306V. | 0.371 | 0.496 | -1.282 | 0.718 | 0.009 | 141O-144N | 1 | 1 | 0 | 1 | 1 | 7.708 | 11.899 | 7.805 | | | | | | | | | | | | | | | 0 | 3.262 | 6.705 |
| | 2amd | | GPLGS | G(-5) of A, GPL of B | 9IN | N9 | P | 6.7 | | 0.617 | 1.326 | -1.212 | -0.300 | 12.519 | 141O-144N | 1 | 1 | 1 | 1 | 1 | 7.556 | 11.471 | 9.548 | 0.257 | -0.738 | -0.832 | -1.998 | 2.543 | 141O-144N | 1 | 1 | 0 | 1 | 1 | 7.612 | 11.370 | 7.026 | 13.631 | 7.25 | 8.085 |
| | 2amq | | GPLGS | GPLGSSG of A, GPLGS of B | PRD_002214 | N3 | P | 9 | | 0.654 | 1.128 | -0.986 | -0.029 | 10.525 | 141O-144N | 1 | 1 | 1 | 1 | 1 | 7.871 | 11.795 | | 0.306 | 0.182 | -0.609 | -0.682 | 2.501 | 141O-144N | 1 | 1 | 0 | 1 | 1 | 7.590 | 11.427 | 7.184 | 12.321 | 6.807 | 7.979 |
| | 2d2d | | GPLGS | GPLGSSG of A, GPLGS of B | ENB | I2 | P | | | 0.598 | 1.191 | -0.232 | 0.765 | 14.349 | 141O-144N | 1 | 1 | 1 | 1 | 1 | 8.165 | 12.139 | | 0.174 | 0.766 | 0.200 | 0.084 | -0.058 | 141O-144N | 1 | 1 | 0 | 1 | 1 | 8.096 | 11.699 | 7.125 | 13.812 | 5.508 | 8.180 |
| | 2gtb | | A | AS | AZP | Aza-peptide epoxide (APE) | P | | | 0.467 | 0.317 | -0.665 | -0.653 | 4.546 | 141O-144N | 1 | 1 | 0 | 1 | 1 | 8.031 | 11.725 | 7.793 | | | | | | | | | | | | | | | 0.001 | 11.181 | 7.694 |
| | 2gx4 | | | | NOL | TG-0205221 | P | 0.053 | | 0.756 | 1.011 | -0.422 | 1.167 | 3.838 | 141O-144N | 1 | 1 | 1 | 1 | 1 | 7.470 | 11.118 | 7.296 | | | | | | | | | | | | | | | 0.002 | 3.231 | 8.018 |
| | 2gz7 | | | | D3F | 3 | | 0.3* | D3F makes H41 flip out, making distance to C145 Sg more than 9A. | 0.439 | 0.715 | -0.404 | 1.613 | 4.980 | 141O-144N | 1 | 1 | 1 | 1 | 1 | 7.935 | 11.560 | 8.381 | | | | | | | | | | | | | | | 0 | 5.83 | 7.754 |
| | 2gz8 | | | | F3F | 15 | | 3* | | 0.591 | 0.484 | -0.357 | 1.504 | 4.664 | 141O-144N | 1 | 1 | 1 | 1 | 1 | 7.734 | 11.447 | 7.57 | | | | | | | | | | | | | | | 0 | 5.363 | 7.841 |
| | 2hob | | | | PRD_002214 | N3 | P | 1.9 | | 0.743 | 1.143 | -0.273 | 0.544 | 5.270 | 141O-144N | 1 | 1 | 1 | 1 | 1 | 7.579 | 11.346 | 7.501 | | | | | | | | | | | | | | | 0.002 | 3.636 | 8.032 |
| | 2op9 | | A | A of B | WR1 | WRR 183 | P | 2.2 | Non-covalent (distance between WR1 and C145 is 2.8Å) in chain A; covalent (1.8Å) in chain B. Both epoxide is eliminated. C-terminal 5 residues do not exist. | 0.244 | -0.056 | 0.401 | 1.264 | -0.522 | 141O-144N | 1 | 1 | 0 | 1 | 1 | 7.619 | 11.651 | 6.911 | 0.554 | 0.232 | | 2.344 | 11.283 | 141O-144N | 1 | 1 | 1 | 1 | 1 | 7.845 | 11.832 | 8.493 | -10.339 | 7.748 | 8.094 |
| | 2q6g | H41A | | | polypeptide chain | N-terminal substrate | P | | Chain A has the intact 11 residues of the substrate; chain B has only N-terminal 8 residues. H41A makes the protein inactive. | 0.390 | 0.777 | -0.304 | 0.046 | 11.445 | 141O-144N | 1 | 1 | 0 | 1 | 1 | 7.287 | 11.001 | 8.83 | -1.018 | -0.389 | -0.248 | -1.348 | 0.339 | | 1 | 1 | 0 | 1 | 1 | 7.319 | 10.873 | 7.364 | 14.75 | 8.742 | 8.360 |
| | 2qiq | S1A | | | CYV | 3 | P | 80* | Non-covalent (distance between CYV and C145 is 2.5Å), while the bond length of C5-C6 of CYV is 1.5 A of a reacted single bond. C-terminal 5 residues are note modeled. | 0.312 | 0.580 | -1.004 | 1.112 | 4.454 | 141O-144N | 1 | 1 | 0 | 1 | 1 | 7.571 | 11.692 | 8.587 | | | | | | | | | | | | | | | 0 | 7.748 | 8.195 |
| | 2v6n | | | | XP1 | XP-59 | | 1.38 | XP-59 becomes XP1 upon binding. | 0.606 | 0.973 | -0.866 | 1.049 | 7.478 | 141O-144N | 1 | 1 | 1 | 1 | 1 | 7.553 | 11.364 | 7.658 | | | | | | | | | | | | | | | 0 | 6.027 | 7.998 |
| | 2vj1 | S1 del | S1 del | S1 del | BEZ | XP-27 | | | XP-27 becomes BEZ upon binding. BEZ of chain A binds to M49 and M165A, BEZ of chain B binds to C145 and E166B. After 303, 6 His are added. Deletion of S1 imapirs dimerization. | -0.426 | -0.526 | -0.131 | 2.364 | 13.997 | 141O-143N | 1 | 1 | 0 | 1 | 1 | 7.900 | 12.173 | 7.607 | -7.382 | -0.803 | 0.179 | 0.763 | -1.926 | 141O-144N | 0 | 0 | 0 | 0 | 0 | 13.360 | 8.212 | 8.408 | 13.364 | 10.982 | 8.279 |
| | 2z3c | | | | PRD_000250 | 1 | P | 0.25 | Soaking. Phthalhydrazide group of 1 is eliminated to make PRD_000250 upon binding. | 0.691 | 0.905 | -0.677 | 0.682 | 5.671 | 141O-144N | 1 | 1 | 1 | 1 | 1 | 7.388 | 11.211 | 7.165 | | | | | | | | | | | | | | | -0.001 | 4.515 | 7.413 |
| | 2z3d | | | | PRD_000235 | 2 | P | | Phthalhydrazide group of 2 is eliminated to make PRD_000250 upon binding. | 0.559 | 1.002 | -1.247 | 1.107 | 6.870 | 141O-144N | 1 | 1 | 1 | 1 | 1 | 7.654 | 11.477 | 7.497 | | | | | | | | | | | | | | | 0 | 6.507 | 7.514 |
| | 2z3e | | | | PRD_000250 | 1 | P | 0.25 | Co-crystallized. Phthalhydrazide group of 1 is eliminated to make PRD_000250 upon binding. | 0.540 | 0.996 | -0.362 | 1.072 | 5.326 | | 1 | 1 | 1 | 1 | 1 | 7.598 | 11.558 | 7.565 | | | | | | | | | | | | | | | 0.001 | 4.673 | 7.284 |
| | 2z94 | | | | TLD+Zn | TDT | | 1.4 | C145 binds to Zn of TDT+Zn. | 0.554 | 0.752 | -0.410 | 1.940 | 6.344 | 141O-144N | 1 | 1 | 1 | 1 | 1 | 7.711 | 11.469 | 7.674 | | | | | | | | | | | | | | | 0 | 5.208 | 7.949 |
| | 2z9g | | | | BNZ+Hg | PMA | | 0.7 | C145 is far from BNZ, but Hg binds to C44 and Y54.PMA becomes benzen upon binding. | 0.603 | 0.329 | -1.043 | 1.482 | 5.224 | 141O-144N | 1 | 1 | 1 | 1 | 1 | 7.627 | 11.394 | 7.589 | | | | | | | | | | | | | | | 0 | 4.793 | 7.912 |
| | 2z9j | | | | DTZ | EPDTC | | 1 | C145 binds to Zn of DTZ. EPDTC becomes DTZ upon binding. | -0.008 | -0.004 | -0.282 | 1.391 | -1.589 | 141O-144N | 1 | 0 | 1 | 1 | 1 | 7.429 | 11.523 | 6.811 | 0.491 | 0.382 | -0.992 | 2.232 | 12.257 | 141O-144N | 1 | 1 | 1 | 1 | 1 | 7.607 | 11.508 | 8.064 | -14.666 | 7.814 | 8.272 |
| | 2z9k | | | | DOZ | JMF1600 | | 0.32 | C145 binds to Zn of DAZ. JMF1600 becomes DOZ upon binding. | 0.558 | 0.683 | -0.835 | 2.374 | 10.733 | 141O-144N | 1 | 1 | 1 | 1 | 1 | 7.687 | 11.475 | 8.038 | 0.306 | 0.399 | -0.183 | 1.284 | 2.638 | 141O-144N | 1 | 1 | 1 | 1 | 1 | 7.662 | 11.518 | 7.108 | 11.317 | 8.405 | 8.291 |
| | 2z9l | | | | DAZ | JMF1586 | | 0.05 | C145 binds to Zn of DAZ. JMF1586 becomes DAZ upon binding. | 0.526 | 0.463 | | 2.162 | 8.985 | 141O-144N | 1 | 1 | 1 | 1 | 1 | 7.595 | 11.472 | 7.51 | 0.508 | -0.096 | -0.228 | 0.857 | -3.806 | 141O-144N | 1 | 1 | 1 | 1 | 1 | 7.772 | 11.344 | 6.882 | 11.382 | 2.886 | 7.591 |
| | 2zu4 | | | | PRD_000568 | TG-0204998 | P | 0.038 | | 0.847 | 1.277 | -0.556 | 0.556 | 1.560 | 141O-144N | 1 | 1 | 1 | 1 | 1 | 7.348 | 11.011 | 7.221 | | | | | | | | | | | | | | | -0.002 | 0.966 | 7.633 |
| | 2zu5 | | | | PRD_000569 | TG-0205486 | P | 0.099 | | 0.723 | 1.492 | -0.152 | 0.630 | 5.711 | 141O-144N | 1 | 1 | 1 | 1 | 1 | 7.501 | 11.207 | 7.483 | | | | | | | | | | | | | | | 0.001 | 3.954 | 8.014 |
| | 3atw | R188I | | | PRD_000815 | 35 | P | 0.098* | Non-covalent (distance between the ligand and C145 is 2.5Å in chain A and 2.7Å in chain B). R188I is to prevent proteolysis and increases the activity. | 0.491 | -0.560 | 0.128 | -2.476 | -12.038 | 141O-144N | 1 | 1 | 1 | 1 | 1 | 7.348 | 10.864 | 6.405 | 0.605 | 0.007 | 0.287 | -0.251 | 2.820 | 141O-144N | 1 | 1 | 1 | 1 | 1 | 7.549 | 11.096 | 8.642 | -7.775 | -5.11 | 6.368 |
| | 3avz | R188I | | | PRD_000990 | 31 | P | 0.065* | Side-chains of Thr285 clashes each other (2.2Å between two Cβ's). R188I is to prevent proteolysis and increases the activity. | 0.418 | 0.641 | -0.323 | 0.382 | -5.214 | 141O-144N | 1 | 1 | 1 | 1 | 1 | 7.386 | 10.822 | 6.569 | | | | | | | | | | | | | | | 0.001 | -7.823 | 5.107 |
| | 3aw0 | R188I | | | PRD_001000 | 16 | P | 5.7* | Non-covalent (distance between the ligand and C145 is 2.4Å).R188I is to prevent proteolysis and increases the activity. | 0.763 | 0.739 | -0.875 | 0.604 | 6.195 | 141O-144N | 1 | 1 | 1 | 1 | 1 | 7.530 | 11.190 | 7.79 | | | | | | | | | | | | | | | 0.001 | 5.429 | 8.050 |
| | 3d62 | S1, G2 del | S1, G2 del | S1, G2 del | 959 | 4 | | 0.4 | Br of compound 4 is replaced by Sg of C145. The Boc moiety is not modeled. C-terminal 7 residues do not exist. | 1.561 | -1.293 | -0.870 | 0.073 | -7.747 | | 1 | 1 | 0 | 0 | 1 | 7.369 | 11.498 | | | | | | | | | | | | | | | | -0.001 | -6.66 | 7.120 |
| | 3sn8 | | | | PRD_001114 | Cm-FF-H | P | 2.24 | | 0.489 | 0.468 | -1.020 | 1.850 | 2.843 | 141O-144N | 1 | 1 | 1 | 1 | 1 | 7.674 | 11.252 | 7.201 | | | | | | | | | | | | | | | 0.001 | 3.62 | 7.353 |
| | 3sna | | | | PRD_001117 | Ac-NSFSQ-H | P | 72.73 | C-terminal 5 residues do not exist. | 0.657 | -0.408 | -1.277 | 2.345 | 0.332 | 141O-144N | 1 | 1 | 1 | 1 | 1 | 7.600 | 11.463 | 7.067 | | | | | | | | | | | | | | | 0 | 0.849 | 6.665 |
| | 3snb | | | | PRD_001118 | Ac-DSFDQ-H | P | 41.24 | | 0.565 | 0.608 | -0.718 | 1.312 | 6.155 | 141O-144N | 1 | 1 | 1 | 1 | 1 | 7.718 | 11.421 | 7.554 | | | | | | | | | | | | | | | 0.001 | 6.034 | 7.984 |
| | 3snc | | | | PRD_000771 | Ac-NSTSQ-H | P | 40.98 | | 0.531 | 0.217 | -1.155 | 1.588 | 3.781 | 141O-144N | 1 | 1 | 1 | 1 | 1 | 7.621 | 11.177 | 7.446 | | | | | | | | | | | | | | | -0.001 | 4.794 | 7.283 |
| | 3snd | | | | PRD_000772 | Ac-ESTLQ-H | P | 8.27 | Cocrystalization. | 0.381 | 0.067 | -0.514 | 0.684 | 4.218 | 141O-144N | 1 | 1 | 1 | 1 | 1 | 7.752 | 11.537 | 7.402 | 0.651 | 0.530 | 0.571 | 1.612 | 9.325 | 141O-144N | 1 | 1 | 1 | 1 | 1 | 7.567 | 11.313 | 8.452 | -8.31 | 8.024 | 8.418 |
| | 3sne | | | | PRD_000772 | Ac-ESTLQ-H | P | 8.27 | Soaking. | 0.641 | 0.262 | -0.535 | 0.772 | 5.418 | 141O-144N | 1 | 1 | 1 | 1 | 1 | 7.747 | 11.441 | 7.451 | | | | | | | | | | | | | | | 0.001 | 5.628 | 8.070 |
| | 3szn | | | | G75 | SG75 | P | | | 0.617 | 1.412 | -0.968 | 1.728 | 6.202 | 141O-144N | 1 | 1 | 1 | 1 | 1 | 7.724 | 11.434 | 7.491 | | | | | | | | | | | | | | | 0 | 4.057 | 7.892 |
| | 3tit | | | | G81 | SG81 | P | | | 0.582 | 1.200 | -0.944 | 1.934 | 5.073 | 141O-144N | 1 | 1 | 1 | 1 | 1 | 7.651 | 11.337 | 7.449 | | | | | | | | | | | | | | | 0.001 | 4.933 | 7.672 |
| | 3tiu | | | | G82 | SG82 | P | | | 0.679 | 0.983 | -0.814 | 1.047 | 4.364 | 141O-144N | 1 | 1 | 1 | 1 | 1 | 7.590 | 11.358 | 7.479 | | | | | | | | | | | | | | | 0 | 3.789 | 7.726 |
| | 3tns | | | | G83 | SG83 | P | | | 0.524 | 1.316 | -0.737 | 0.448 | 3.098 | 141O-144N | 1 | 1 | 1 | 1 | 1 | 7.648 | 11.367 | 7.253 | | | | | | | | | | | | | | | -0.001 | 1.863 | 7.078 |
| | 3tnt | | | | G85 | SG85 | P | | | 0.417 | 1.340 | -0.042 | 0.668 | 4.651 | 141O-144N | 1 | 1 | 1 | 1 | 1 | 7.815 | 11.596 | 7.475 | | | | | | | | | | | | | | | 0 | 3.233 | 7.404 |
| | 3v3m | | | | 0EN | ML188(16-(R)) | | 1.5* | | 0.338 | 0.236 | -1.572 | 1.814 | 5.557 | 141O-144N | 1 | 1 | 1 | 1 | 1 | 7.706 | 11.234 | 7.418 | | | | | | | | | | | | | | | 0.001 | 5.151 | 6.974 |
| | 3vb4 | | | | PRD_000910 | B4Z | P | 49 | | 0.845 | -0.626 | -0.723 | -0.583 | -6.993 | 141O-144N | 1 | 1 | 1 | 1 | 1 | 7.890 | 11.846 | 6.604 | 0.817 | 0.046 | 0.980 | 0.379 | 7.397 | | 1 | 1 | 1 | 1 | 1 | 7.729 | 11.677 | 8.018 | -13.389 | 3.127 | 6.970 |
| | 3vb5 | | | | PRD_000911 | C4Z | P | 4.6 | | 0.754 | 0.480 | 0.026 | 0.697 | -2.385 | 141O-144N | 1 | 1 | 1 | 1 | 1 | 7.605 | 11.118 | 6.988 | 0.853 | 0.530 | 0.582 | 1.159 | 10.814 | 141O-144N | 1 | 1 | 1 | 1 | 1 | 7.656 | 11.278 | 8.031 | -13.905 | 7.166 | 8.228 |
| | 3vb6 | | | | PRD_000912 | C6Z | P | 39 | Cbz, T, and S of C6Z are not modeled. | 0.003 | -0.571 | -0.281 | 0.649 | -4.244 | 141O-144N | 1 | 1 | 1 | 1 | 1 | 7.483 | 11.460 | 7.049 | 0.619 | 0.154 | -0.186 | 1.183 | 11.381 | 141O-144N | 1 | 1 | 1 | 1 | 1 | 7.427 | 11.565 | 8.428 | -14.917 | 6.049 | 7.844 |
| | 3vb7 | | | | PRD_000913 | M4Z | P | 49 | Mic is not modeled. | 0.553 | -0.032 | -1.220 | 0.338 | -1.550 | 141O-144N | 1 | 1 | 1 | 1 | 1 | 7.624 | 11.529 | 6.997 | 0.769 | 0.387 | 0.249 | 0.899 | 11.965 | 141O-144N | 1 | 1 | 1 | 1 | 1 | 7.653 | 11.385 | 8.329 | -13.757 | 8.092 | 8.244 |
| | 4mds | | | | 23H | 7 | | 6.2* | V303A. C-terminal 3 residues do not exist. | 0.508 | -0.435 | 0.952 | 1.937 | -1.091 | 141O-144N | 1 | 1 | 1 | 1 | 1 | 7.594 | 11.103 | 6.585 | | | | | | | | | | | | | | | 0 | -1.317 | 6.313 |
| | 4tww | R188I | | | 3A7 | 41 | | 63* | Non-covalent (distance between 3A7 and C145 is 2.3Å in chain A and 2.4Å in chain B). R188I prevents from proteolysis and increases the activity. | 0.199 | -0.368 | 0.096 | -2.507 | -9.745 | 141O-144N | 1 | 1 | 1 | 1 | 1 | 7.535 | 11.050 | 6.746 | 0.401 | 0.021 | 0.303 | -0.061 | 2.633 | 141O-144N | 0 | 1 | 1 | 1 | 1 | 7.574 | 11.031 | 8.845 | -8.363 | -1.021 | 6.310 |
| | 4twy | R188I | | | 3BL | 40 | | 108* | R188I prevents from proteolysis and increases the activity. | 0.490 | 0.582 | -0.566 | 0.598 | 7.188 | 141O-144N | 1 | 1 | 1 | 1 | 1 | 7.664 | 11.165 | 7.704 | | | | | | | | | | | | | | | -0.001 | 7.018 | 7.849 |
| | 4wy3 | R188I | | | 3X5 | 44 | | 240* | Non-covalent (distance between 3X5 and C145 is 2.5Å). R188I prevents from proteolysis and increases the activity. | 0.525 | 0.593 | -0.832 | 1.118 | 7.545 | 141O-144N | 1 | 1 | 1 | 1 | 1 | 7.829 | 11.423 | 7.62 | | | | | | | | | | | | | | | 0 | 7.262 | 8.092 |
| | 5b6o_A | C145A | | | C-term of another chain A | | P | | Chain A binds a symmetrically related chain A (301-310), while chain B is in free state. After C-term, GKFKKIVKGT are added. C145A makes the protein inactive. | 0.587 | 0.835 | -0.821 | 0.884 | -23.013 | 141O-144N | 1 | 1 | 1 | 1 | 1 | 7.385 | 11.113 | 7.364 | | | | | | | | | | | | | | | -21.195 | -3.44 | 6.189 |
| | 5c5n | R188I | | | SLH | phenyl-beta-alanyl (R,S)-N-decalin type inhibitor | | 275* | R188I prevents from proteolysis and increases the activity. | 0.463 | 0.693 | -0.672 | 1.817 | 7.759 | 141O-144N | 1 | 1 | 1 | 1 | 1 | 7.773 | 11.261 | 7.674 | | | | | | | | | | | | | | | 0.002 | 7.618 | 7.851 |
| | 5c5o | R188I | | | SDJ | phenyl-beta-alanyl (S,R)-N-decalin type inhibitor | | 215* | R188I prevents from proteolysis and increases the activity. | 0.299 | -0.197 | 0.147 | -2.599 | -12.742 | 141O-144N | 1 | 1 | 1 | 1 | 1 | 7.519 | 10.979 | 6.417 | 0.401 | 0.040 | 0.322 | -0.367 | 4.554 | 141O-144N | 1 | 1 | 1 | 1 | 1 | 7.467 | 10.937 | 8.909 | -10.587 | -3.132 | 6.253 |
| | 5n19 | | | | D03 | | P | | | 0.522 | 0.939 | -0.886 | 2.081 | 6.459 | 141O-144N | 1 | 1 | 1 | 1 | 1 | 7.686 | 11.221 | 7.543 | | | | | | | | | | | | | | | 0 | 6.165 | 8.122 |
| | 5n5o | | | | 8O5 | | P | | | 0.654 | 0.975 | -0.262 | -0.250 | 4.475 | 141O-144N | 1 | 1 | 1 | 1 | 1 | 7.550 | 11.299 | 7.401 | | | | | | | | | | | | | | | 0 | 3.436 | 7.899 |
| | 6lnq | | | | EJF | M7 | P | | | 0.850 | 1.020 | -0.097 | 0.470 | 4.736 | 141O-144N | 1 | 1 | 1 | 1 | 1 | 7.575 | 11.363 | 7.489 | | | | | | | | | | | | | | | 0.001 | 3.887 | 8.022 |
| | 6lny | | | | EOC | 8a (M15) | P | 0.508* | | 0.787 | 0.731 | -0.091 | 0.256 | 5.136 | 141O-144N | 1 | 1 | 1 | 1 | 1 | 7.490 | 11.296 | 7.344 | | | | | | | | | | | | | | | 0.001 | 4.135 | 8.078 |
| | 6lo0 | | | | EOF | 8b (M14) | P | 0.830* | | 0.816 | 1.368 | 0.441 | -2.230 | 4.580 | 141O-144N | 1 | 1 | 1 | 1 | 1 | 7.471 | 11.247 | 7.488 | | | | | | | | | | | | | | | 0 | 3.031 | 8.035 |
| | 6w2a | | MHHHHHH | MHHHHHH(-1) of A,B | VDJ/QYS | 7j | | 1.1* | Sulfoxide of 7j is replaced by Sg of C145. The two ligands are stereoisomers, and cannnot be distinguished after binding. C-terminal 3 residues do not exist. Highly heterogeneous crystal environment of P 1 21 1. | 0.485 | 1.004 | -0.022 | -3.233 | -7.187 | 141O-144N | 1 | 1 | 0 | 1 | 1 | 7.246 | 10.969 | 6.41 | 0.346 | 0.902 | -0.302 | -2.652 | 22.177 | 141O-144N | 1 | 1 | 1 | 1 | 1 | 7.871 | 11.761 | 9.456 | -20.855 | 9.505 | 6.884 |
| | 6w79 | | | | X77 | X77 | | | | 0.496 | 0.329 | -1.274 | 1.296 | 6.618 | 141O-144N | 1 | 1 | 1 | 1 | 1 | 7.801 | 11.379 | 7.671 | | | | | | | | | | | | | | | 0.001 | 6.13 | 8.256 |
| | 6wco | | | | X47 | X47 | | | | 0.571 | 0.571 | -1.437 | 1.448 | 6.294 | 141O-144N | 1 | 1 | 1 | 1 | 1 | 7.836 | 11.534 | 7.736 | | | | | | | | | | | | | | | -0.002 | 7.613 | 8.217 |
| | 6xhl | | G | GS of A,B | V2M | 4 | P | 0.004*, 5$^{+}$ | Highly heterogeneous crystal environment of P 1 21 1. | 0.333 | 1.432 | 0.860 | -2.811 | -10.014 | 141O-144N | 1 | 1 | 0 | 1 | 1 | 7.560 | 11.421 | 6.348 | 0.351 | 1.139 | 0.467 | -2.708 | 42.960 | 141O-144N | 1 | 1 | 0 | 1 | 1 | 7.757 | 11.690 | 9.131 | -38.32 | 20.824 | 9.541 |
| | 6xhn | | G | GS of A, GSG of B | V3D | 28 | P | 0.017* | Highly heterogeneous crystal environment of P 1 21 1. | 0.384 | 1.627 | 0.376 | -2.882 | -4.269 | 141O-144N | 1 | 1 | 0 | 1 | 1 | 7.505 | 11.329 | 6.444 | 0.304 | 1.636 | 0.099 | -3.092 | 41.826 | 141O-144N | 1 | 1 | 0 | 1 | 1 | 7.645 | 11.503 | | -34.819 | 23.401 | 10.082 |
| | 6xho | | G | GS of A, GSG of B | V34 | 2 | P | | Highly heterogeneous crystal environment of P 1 21 1. | 0.333 | 1.707 | 0.919 | -3.474 | -7.808 | 141O-144N | 1 | 1 | 0 | 1 | 1 | 7.591 | 11.513 | 6.326 | 0.340 | 1.366 | -0.032 | -3.001 | 34.138 | 141O-144N | 1 | 1 | 0 | 1 | 1 | 7.809 | 11.682 | | -31.549 | 15.747 | 7.795 |
| | 6y7m | | | | OEW | 13a | P | | | 0.592 | 0.845 | -0.904 | 1.219 | 1.878 | 141O-144N | 1 | 1 | 1 | 1 | 1 | 7.575 | 11.323 | 7.25 | | | | | | | | | | | | | | | 0.001 | 1.258 | 7.271 |
| SARS-CoV-2% | 6lu7 | | | | PRD_002214 | N3 | P | 16.77$^{+}$ | | 0.810 | 0.732 | -0.909 | -1.766 | -5.522 | 141O-144N | 1 | 1 | 1 | 1 | 1 | 7.303 | 11.101 | 6.421 | | | | | | | | | | | | | | | 0 | -11.704 | 5.314 |
| | 6lze | | | | PRD_002347 | 11a | P | 0.053* | C-terminal 3 reidues do not exist. | 0.845 | 0.314 | -0.773 | 1.870 | -8.648 | 141O-144N | 1 | 1 | 1 | 1 | 1 | 7.466 | 11.174 | 6.328 | | | | | | | | | | | | | | | 0.001 | -12.306 | 5.355 |
| | 6m0k | | | | PRD_002349 | 11b | P | 0.040* | C-terminal 2 residues do not exist. | 0.832 | 0.348 | -0.023 | 1.907 | -8.517 | 141O-144N | 1 | 1 | 1 | 1 | 1 | 7.492 | 11.197 | 6.403 | | | | | | | | | | | | | | | 0 | -11.853 | 5.366 |
| | 6m2n_AC | | | | 3WL | Baicalein | | 0.94* | Tetramer (two asymmetric dimers). | 0.623 | -0.172 | -0.280 | -0.496 | 0.217 | 141O-144N | 1 | 1 | 1 | 1 | 1 | 7.471 | 11.322 | 7.116 | 0.764 | -0.407 | 0.323 | 1.063 | -4.649 | 141O-144N | 1 | 1 | 1 | 1 | 1 | 7.642 | 11.302 | 6.911 | -0.03 | -1.811 | 5.429 |
| | 6m2n_BD | | | | 3WL | Baicalein | | | | 0.632 | -0.152 | -0.226 | -0.650 | 1.125 | 141O-144N | 1 | 1 | 1 | 1 | 1 | 7.454 | 11.198 | 7.257 | 0.755 | -0.243 | 0.211 | 1.436 | -4.548 | 141O-144N | 1 | 1 | 1 | 1 | 1 | 7.682 | 11.431 | 7.004 | 1.358 | -1.292 | 5.608 |
| | 6w63 | | | | X77 | X77 | | | | 0.576 | -1.331 | -0.820 | 0.274 | -7.867 | 141O-144N | 1 | 1 | 1 | 1 | 1 | 7.526 | 11.098 | 6.404 | | | | | | | | | | | | | | | -0.001 | -9.567 | 5.684 |
| | 6wnp | | | | PRD_002382 | Boceprevir | P | | | 0.706 | 0.326 | -0.471 | -0.537 | -2.562 | 141O-144N | 1 | 1 | 1 | 1 | 1 | 7.716 | 11.332 | 6.799 | | | | | | | | | | | | | | | 0.001 | -4.702 | 5.614 |
| | 6wtj | | | | K36 | GC376 | P | 0.19* | Sulfoxide is replaced by Sg of C145. Upon binding to C145, the prodrug, GC376, is converted to GC373. | 0.997 | -0.759 | -0.548 | -2.950 | -8.744 | 141O-144N | 1 | 1 | 1 | 1 | 1 | 7.409 | 11.218 | 6.283 | | | | | | | | | | | | | | | 0 | -10.198 | 5.350 |
| | 6wtk | | | | UED | GC373 | P | 0.40*, 1.5$^{+}$ | | 0.936 | -0.131 | -0.554 | -2.714 | -8.547 | 141O-144N | 1 | 1 | 1 | 1 | 1 | 7.340 | 11.261 | 6.35 | | | | | | | | | | | | | | | 0.001 | -11.494 | 5.220 |
| | 6wtt_AA | | | | K36 | GC376 | P | 0.03* 3.37$^{+}$ | Tetramer. Chain A is symmetric, while chain B&C are assymmetric. Sulfoxide in K36 is replaced by Sg of C145. Upon binding, GC376 becomes K36 in chain A&B and B1S (stereoisomer of K36) in chain C. After 304, 6 His are added. | 1.270 | 0.919 | -0.157 | 1.061 | -4.006 | 141O-144N | 1 | 1 | 1 | 1 | 1 | 7.364 | 10.897 | 6.17 | | | | | | | | | | | | | | | -0.001 | -8.032 | 5.262 |
| | 6wtt_BC | | | | K36, B1S | GC376 | | | | 1.444 | -0.069 | -0.329 | 1.382 | -9.852 | 141O-144N | 1 | 1 | 1 | 1 | 1 | 7.186 | 11.036 | 6.293 | 1.397 | -0.673 | -1.046 | -0.744 | -12.140 | 141O-144N | 1 | 1 | 1 | 1 | 1 | 7.314 | 10.892 | 6.618 | 1.459 | -11.265 | 5.024 |
| | 6xa4 | | HM | HMS | PRD_002391 | calpain inhibitor II | P | 0.97* | | 0.919 | -0.870 | 0.378 | -2.261 | -12.065 | 141O-144N | 1 | 1 | 0 | 1 | 1 | 7.785 | 11.646 | 6.734 | | | | | | | | | | | | | | | -0.001 | -9.569 | 5.410 |
| | 6xb1 | | | | NEN | | | | C145 is di-oxydized, 2CO ( SOO), and C156 is capped by NEN. | 0.608 | -1.803 | -1.380 | -2.465 | -2.332 | 141O-144N | 1 | 1 | 1 | 1 | 1 | 7.904 | 11.778 | 6.46 | | | | | | | | | | | | | | | 0 | -4.951 | 5.556 |
| | 6xb2 | | | | NEN | | | | C145 and C156 are capped by NEN. | 0.639 | -1.802 | -1.042 | -2.207 | -1.586 | 141O-144N | 1 | 1 | 1 | 1 | 1 | 7.644 | 11.596 | 6.487 | | | | | | | | | | | | | | | 0 | -4.277 | 5.611 |
| | 6xbg | | HM | HMS of A, HMSG of B | PRD_002392 | UAW246 | P | 0.045* 15.13$^{+}$ | Highly heterogeneous crystal environment of P 1 21 1. | 0.604 | 0.913 | 0.382 | -2.689 | -10.519 | 141O-144N | 1 | 1 | 0 | 1 | 1 | 7.614 | 11.309 | 6.113 | 0.293 | 1.053 | -0.967 | -0.113 | 33.895 | 141O-144N | 1 | 1 | 0 | 1 | 1 | 7.696 | 11.431 | | -31.358 | 14.717 | 7.416 |
| | 6xbh | | HM | HMS | PRD_002393 | UAW247 | P | 0.045* 6.81$^{+}$ | | 1.104 | -1.412 | -0.486 | -2.591 | -13.650 | 141O-144N | 1 | 1 | 0 | 1 | 1 | 7.539 | 11.261 | 6.584 | | | | | | | | | | | | | | | -0.001 | -11.172 | 5.215 |
| | 6xbi | | HM | H of A | PRD_002394 | UAW248 | P | 20.49$^{+}$ | | 0.644 | 0.874 | -1.776 | 0.654 | -11.376 | 141O-144N | 1 | 1 | 0 | 1 | 1 | 7.434 | 11.366 | 6.312 | 1.030 | 1.224 | -2.310 | 1.442 | -9.291 | 141O-144N | 1 | 1 | 0 | 1 | 1 | 7.531 | 11.248 | 7.375 | -0.21 | -7.152 | 5.476 |
| | 6xch | | | | PRD_000216 | Leupeptin | P | 92* | | 0.664 | -0.049 | -0.661 | -1.849 | -9.910 | 141O-144N | 1 | 1 | 1 | 1 | 1 | 7.882 | 11.630 | 6.5 | | | | | | | | | | | | | | | 0 | -13.895 | 5.346 |
| | 6xfn | | HM | HMS | | calpain inhibitor XII | P | 0.453* | | 0.137 | -2.309 | -0.315 | -2.766 | -3.527 | 141O-144N | 1 | 1 | 0 | 1 | 1 | 7.887 | 11.413 | 7.384 | | | | | | | | | | | | | | | 0.001 | -0.995 | 5.746 |
| | 6xhm | | | | V2M | 4 | P | 0.00027 | Highly heterogeneous crystal environment of P 1 21 1. | 0.563 | 1.214 | 1.455 | -2.823 | -15.900 | 141O-144N | 1 | 1 | 1 | 1 | 1 | 7.557 | 11.418 | 6.387 | 0.423 | 1.510 | 0.709 | -3.217 | 32.433 | 141O-144N | 1 | 1 | 1 | 1 | 1 | 7.787 | 11.775 | 9.178 | -34.993 | 9.904 | 6.982 |
| | 6xmk | | SNIG | SNIGSGF of A, SNIG of B | QYS | 7j | P | 0.45* | Q306 do not exist. Highly heterogeneous crystal environment of P 1 21 1. | 0.668 | 0.422 | 0.735 | -2.507 | -13.981 | 141O-144N | 1 | 1 | 0 | 1 | 1 | 7.307 | 10.998 | | 0.658 | 0.457 | -0.168 | -2.528 | 42.060 | 141O-144N | 1 | 1 | 0 | 1 | 1 | 7.431 | 11.221 | 9.52 | -38.069 | 19.635 | 8.892 |
| | 6xoa_AC | C145S | | | C-term of another protein | | P | | Tetramer (two asym dimers AB and CD). Chains A&C bind symmetrically related C-term of C&A, respectively. Dimer data are of AB. Chains B&D are free. C145S makes the protein inactive. | 0.799 | -0.531 | 1.075 | 1.935 | -11.780 | 141O-144N | 1 | 1 | 1 | 1 | 1 | 7.769 | 11.468 | 6.37 | 0.795 | 0.599 | 0.242 | 0.255 | -2.678 | 141O-144N | 1 | 1 | 1 | 1 | 1 | 7.478 | 11.202 | 6.46 | -16.415 | -2.912 | 5.273 |
| | 6xqs | | | | SV6 | Telaprevir | P | 18* | | 0.586 | 0.343 | -0.285 | -1.472 | 0.222 | 141O-144N | 1 | 1 | 1 | 1 | 1 | 7.785 | 11.312 | 6.593 | | | | | | | | | | | | | | | 0 | -3.656 | 5.998 |
| | 6xqt | | | | NNA | Narlaprevir | P | 5.1* | | 1.014 | -0.093 | -0.104 | -2.576 | -10.810 | 141O-144N | 1 | 1 | 1 | 1 | 1 | 7.733 | 11.354 | 6.472 | 0.833 | 1.074 | -1.062 | -2.523 | -9.538 | 141O-144N | 1 | 1 | 1 | 1 | 1 | 7.720 | 11.430 | 6.464 | -1.011 | -12.379 | 5.658 |
| | 6xqu | | | | PRD_002382 | Boceprevir | P | 3.1* | | 0.772 | 1.803 | -1.035 | -1.535 | -8.750 | 141O-144N | 1 | 1 | 1 | 1 | 1 | 7.670 | 11.423 | 6.512 | | | | | | | | | | | | | | | 0.001 | -13.286 | 5.313 |
| | 6xr3 | | | | V7G | GRL-024-20 | P | | | 0.504 | 0.764 | -0.505 | -2.556 | -1.300 | 141O-144N | 1 | 1 | 1 | 1 | 1 | 7.717 | 11.390 | 6.595 | | | | | | | | | | | | | | | 0 | -4.239 | 5.540 |
| | 6y2f | | | | O6K | 13b | P | 0.67* 4-5$^{+}$ | | 0.991 | 0.791 | | 1.606 | -5.243 | 141O-144N | 1 | 1 | 1 | 1 | 1 | 7.295 | 10.972 | 6.357 | | | | | | | | | | | | | | | 0.001 | -8.942 | 5.209 |
| | 6y2g | | | | O6K | 13b | P | 0.67* 4-5$^{+}$ | | 0.954 | 1.489 | -0.209 | 0.899 | -0.775 | 141O-144N | 1 | 1 | 1 | 1 | 1 | 7.399 | 11.270 | 7.158 | 1.199 | 2.009 | 0.557 | 1.410 | -12.923 | 141O-144N | 1 | 1 | 1 | 1 | 1 | 7.423 | 11.273 | 6.823 | 6.551 | -3.949 | 5.227 |
| | 6ynq | | | | P6N | 2-Methyl-1-tetralone | | 24.05$^{+}$ | HEAT is cleaved into 2-Methyl-1-tetralone and Tyramine. EC50 is for HEAT. | 0.884 | -1.053 | -1.476 | -0.541 | -5.978 | 141O-144N | 1 | 1 | 1 | 1 | 1 | 7.729 | 11.527 | 6 | | | | | | | | | | | | | | | 0.001 | -9.846 | 5.183 |

| | | | | | | | | | | | | | | | | | | | | | | | | | | | | | | | | | | | | | | | | |
|---|---|---|---|---|---|---|---|---|---|---|---|---|---|---|---|---|---|---|---|---|---|---|---|---|---|---|---|---|---|---|---|---|---|---|---|---|---|---|---|---|
| | 6yvf | | | | A82 | AZD6482 | | | A82 is on the molecualr surface, which is not contained in the major binding sites. | 0.880 | -0.551 | -1.436 | 0.332 | -5.541 | 141O-144N | 1 | 1 | 1 | 1 | 1 | 7.731 | 11.584 | 6.473 | | | | | | | | | | | | | | | 0.001 | -8.7 | 5.308 |
| | 6z2e | | | | Q5T | biotin-PEG(4)-Abu-Tle-Leu-Gln-vinylsulfone | P | | Q5T contain a long polyether chain. | 0.859 | 0.838 | 0.051 | -1.018 | -2.176 | 141O-144N | 1 | 1 | 1 | 1 | 1 | 7.427 | 10.983 | 6.321 | | | | | | | | | | | | | | | 0.001 | -4.654 | 5.671 |
| | 6zrt | | | | PRD_002448 | Telaprevir | P | 55.76* | | 0.776 | 0.330 | -0.340 | -1.444 | 1.028 | 141O-144N | 1 | 1 | 1 | 1 | 1 | 7.567 | 11.482 | 6.483 | | | | | | | | | | | | | | | 0 | -2.954 | 5.783 |
| | 6zru | | | | PRD_002382 | Boceprevir | P | 1.59* | | 0.852 | 1.447 | -0.563 | -1.332 | -11.195 | 141O-144N | 1 | 1 | 1 | 1 | 1 | 7.611 | 11.594 | 6.37 | | | | | | | | | | | | | | | -0.001 | -14.568 | 5.172 |
| | 7aku | | | | RN2 | Calpeptin | | 0.072$^{+}$ | | 0.643 | -0.113 | -1.265 | -2.456 | -9.968 | 141O-144N | 1 | 1 | 1 | 1 | 1 | 7.552 | 11.144 | 6.165 | | | | | | | | | | | | | | | -0.001 | -10.285 | 5.499 |
| | 7b83 | | | | PK8 | pyrithione zinc | | | PK8 covalently binds both C145 and H41 at Zn. | 1.020 | -0.909 | -1.713 | -0.256 | -6.838 | 141O-144N | 1 | 1 | 1 | 1 | 1 | 7.756 | 11.595 | 6.297 | | | | | | | | | | | | | | | 0 | -10.158 | 5.175 |
| | 7bqy | | | | PRD_002214 | N3 | P | 16.77$^{+}$ | Ligand binding is improved over 6lu7. | 0.798 | 1.130 | -1.121 | -0.430 | -3.718 | 141O-144N | 1 | 1 | 1 | 1 | 1 | 7.378 | 11.151 | 6.5 | | | | | | | | | | | | | | | 0 | -8.561 | 5.435 |
| | 7brp | | G | G of A,B | PRD_002382 | Boceprevir | P | 8.0* | | 1.102 | 3.712 | -0.151 | -1.539 | -12.159 | 141O-144N | 1 | 1 | 1 | 1 | 1 | 7.650 | 11.712 | 6.408 | 1.301 | 3.165 | 0.697 | -0.988 | -14.419 | 141O-144N | 1 | 1 | 1 | 1 | 1 | 7.611 | 11.724 | 6.458 | 1.075 | -14.237 | 5.518 |
| | 7buy | | | | JRY | Carmofur | | 1.82* 24.30$^{+}$ | The fluorouracil moiety of carmofur is eliminated to make JRY upond binding. | 0.441 | 0.202 | 0.481 | 1.601 | -7.960 | 141O-144N | 1 | 1 | 1 | 1 | 1 | 7.560 | 11.219 | 6.501 | | | | | | | | | | | | | | | 0.002 | -11.529 | 5.270 |
| | 7c6s | | | | PRD_002382 | Boceprevir | P | 8.0* | | 0.689 | 0.558 | 0.273 | -1.000 | -4.263 | 141O-144N | 1 | 1 | 1 | 1 | 1 | 7.666 | 11.444 | 6.598 | | | | | | | | | | | | | | | 0.001 | -6.71 | 5.476 |
| | 7c6u | | | | K36 | GC376 | | 0.15* | Sulfoxide is replaced by Sg of C145. | 1.050 | 0.649 | -0.039 | -1.400 | -10.703 | 141O-144N | 1 | 1 | 1 | 1 | 1 | 7.297 | 11.162 | 6.203 | | | | | | | | | | | | | | | 0 | -13.249 | 5.552 |
| | 7c7p | | | | PRD_002448 | Telaprevir | P | | Almost a half of PRD_002448 is missing in chain B, and named FK3, while it is fully modeled in chain A. | 0.510 | 0.706 | -0.844 | -0.428 | 3.052 | 141O-144N | 1 | 1 | 1 | 1 | 1 | 7.770 | 11.324 | 6.887 | 0.746 | 0.404 | | 1.973 | -3.453 | 141O-144N | 1 | 1 | 1 | 1 | 1 | 7.646 | 11.384 | 6.51 | -0.455 | -2.592 | 5.356 |
| | 7c8b | | | | PRD_002352 | Z-VAD (OMe)-FMK | P | | | 0.524 | 0.064 | -0.754 | 2.114 | -5.474 | 141O-144N | 1 | 1 | 1 | 1 | 1 | 7.850 | 11.617 | 6.745 | | | | | | | | | | | | | | | -0.001 | -8.732 | 5.313 |
| | 7c8r | | | | TG3 | TG-0203770 | P | | | 0.587 | 2.254 | 0.015 | 0.779 | 0.790 | 141O-144N | 1 | 1 | 1 | 1 | 1 | 7.410 | 11.093 | 6.349 | | | | | | | | | | | | | | | -0.001 | -5.224 | 5.752 |
| | 7c8t | | | | NOL | TG-0205221 | P | | | 0.895 | 1.518 | -1.047 | 1.433 | -1.767 | 141O-144N | 1 | 1 | 1 | 1 | 1 | 7.556 | 11.113 | 6.295 | | | | | | | | | | | | | | | 0 | -5.886 | 5.530 |
| | 7c8u | | | SG | K36 | GC376 | P | | Sulfoxide is replaced by Sg of C145. | 0.811 | -0.524 | -0.553 | -2.401 | -5.604 | 141O-144N | 1 | 1 | 0 | 1 | 1 | 7.323 | 11.298 | | | | | | | | | | | | | | | | 0 | -6.932 | 5.607 |
| | 7cbt | | 27 res | 27 res+ SGF of A,B | K36 | GC376 | P | 0.643$^{+}$ 0.881$^{+}$ | Sulfoxide is replaced by Sg of C145. | 0.896 | 0.602 | 0.816 | -2.104 | -4.228 | 141O-144N | 1 | 1 | 0 | 1 | 1 | 7.460 | 10.963 | | 0.774 | 0.615 | 0.606 | -1.354 | 2.620 | | 1 | 1 | 0 | 1 | 1 | 7.542 | 11.217 | | -3.417 | 0.837 | 5.545 |
| | 7com | | | | PRD_002382 | Boceprevir | P | | | 0.674 | 0.488 | -0.462 | 1.660 | 2.253 | 141O-144N | 1 | 1 | 1 | 1 | 1 | 7.854 | 11.507 | 6.907 | 0.819 | 0.191 | | 1.942 | -3.242 | 141O-144N | 1 | 1 | 1 | 1 | 1 | 7.797 | 11.516 | 6.851 | -0.504 | -2.223 | 5.446 |
| | 7cx9 | | | | GKF | INZ-1 | | | | 0.214 | 1.000 | | 1.389 | -4.238 | 141O-144N | 1 | 1 | 1 | 1 | 1 | 8.053 | 11.962 | 6.732 | | | | | | | | | | | | | | | -0.001 | -8.227 | 5.137 |
| | 7d1m | | G | GS of A, GSG of B | K36 | GC376 | P | 0.15* | Sulfoxide is replaced by Sg of C145. Highly heterogeneous crystal environment of P 1 21 1. | 0.833 | 0.791 | 0.592 | -2.880 | -12.259 | 141O-144N | 1 | 1 | 0 | 1 | 1 | 7.362 | 11.082 | 8.549 | 0.349 | 0.554 | 0.029 | -2.161 | 31.074 | 141O-144N | 1 | 1 | 0 | 1 | 1 | 7.454 | 11.396 | | -31.017 | 13.809 | 6.972 |
| | 7d1o | | | | NNA | Narlaprevir | P | 16.11* | | 1.196 | 0.667 | 0.236 | -1.369 | -16.306 | 141O-144N | 1 | 1 | 1 | 1 | 1 | 7.506 | 11.658 | 6.491 | | | | | | | | | | | | | | | -0.002 | -13.997 | 5.564 |
| | 7d3i | | | | GQU | MI-23 | P | 0.0076* | GQU is hydrogen bonded with S1 of chain B. | 1.280 | 2.294 | -1.533 | 0.015 | -8.860 | 141O-144N | 1 | 1 | 1 | 1 | 1 | 7.479 | 11.172 | 6.335 | | | | | | | | | | | | | | | -0.001 | -9.409 | 5.493 |
| | 7jkv | | G | G of A | V7G | 5h | P | 0.0176 4.2$^{+}$ | Highly heterogeneous crystal environment of P 1 21 1. | 0.248 | 1.751 | 0.531 | -3.052 | -10.714 | 141O-144N | 1 | 1 | 0 | 1 | 1 | 7.506 | 11.311 | 6.666 | -0.012 | 1.283 | 0.479 | -3.065 | 38.703 | 141O-144N | 1 | 1 | 1 | 1 | 1 | 7.729 | 11.566 | 9.737 | -33.707 | 18.045 | 7.974 |
| | 7joy_B | C145A | | | C-term of another chain B | | P | | Chain B binds a symmetrically related C-term, while chain A are in free state. C145A makes the protein inactive. | | | | | | | | | | | | | | | 0.844 | 0.369 | -0.369 | 0.808 | 8.088 | 141O-144N | 1 | 1 | 1 | 1 | 1 | 7.298 | 10.675 | 7.975 | -1.691 | 5.914 | 5.965 |
| | 7ju7 | | | | G65 | Masitinib | | 2.6,2.5* 2.1$^{+}$,3.2$^{+}$ | | 0.455 | 0.378 | -0.314 | 1.819 | -7.308 | 141O-144N | 1 | 1 | 1 | 1 | 1 | 7.784 | 11.370 | 6.596 | | | | | | | | | | | | | | | 0 | -11.037 | 5.484 |
| | 7jyc | | | | NNA | Narlaprevir | P | | | 1.214 | 0.992 | -1.365 | -2.237 | -13.017 | 141O-144N | 1 | 1 | 1 | 1 | 1 | 7.528 | 11.504 | 6.252 | | | | | | | | | | | | | | | 0 | -15.669 | 5.428 |
| | 7k40 | | | | PRD_002382 | Boceprevir | P | | | 0.872 | 2.636 | -0.879 | -1.686 | -12.475 | 141O-144N | 1 | 1 | 1 | 1 | 1 | 7.539 | 11.449 | 6.151 | | | | | | | | | | | | | | | 0 | -17.424 | 5.113 |
| | 7k6d | | | | PRD_002448 | Telaprevir | P | | Acoustic droplet ejection. Cryo-protected. | 0.713 | 0.494 | -0.368 | -1.482 | 0.183 | 141O-144N | 1 | 1 | 1 | 1 | 1 | 7.673 | 11.389 | 6.485 | | | | | | | | | | | | | | | 0 | -4.091 | 6.131 |
| | 7k6e | | | | PRD_002448 | Telaprevir | P | | Acoustic droplet ejection. Direct vitrification. | 0.599 | 0.531 | -0.223 | -1.323 | 1.348 | 141O-144N | 1 | 1 | 1 | 1 | 1 | 7.618 | 11.448 | 6.517 | | | | | | | | | | | | | | | 0 | -3.897 | 6.040 |
| | 7khp_B | | | | C-term of another chain B | | P | | Chain B binds a symmetrically related C-term, while chain A are in free state. Q306 forms a covalent bond with C145 to form the acyl-enzyme intermediate. | | | | | | | | | | | | | | | 0.836 | 0.939 | -0.173 | 1.437 | 9.127 | 141O-144N | 1 | 1 | 1 | 1 | 1 | 7.579 | 11.183 | 7.356 | -3.358 | 5.057 | 5.710 |
| | 7lfp | | GAM | GAM of B | XY4 | F2X Entry Library G05 fragment | | | A single XY4 is at the protomer interface between the two N-fingers, and located out side the major binding sites. | -1.680 | -0.364 | 0.029 | 1.703 | 8.856 | | 1 | 0 | 0 | 1 | 0 | 13.658 | 8.634 | 10.043 | -10.861 | 1.054 | 2.326 | 2.327 | 10.531 | 139O-142N | 0 | 0 | 0 | 0 | 0 | 9.802 | 13.847 | 7.971 | -3.084 | 17.418 | 9.847 |
| | 7nev | | | | PRD_000216 | Leupeptin | P | | | 1.119 | 0.767 | -0.509 | 0.180 | -14.576 | 141O-144N | 1 | 1 | 1 | 1 | 1 | 7.569 | 11.364 | 6.375 | | | | | | | | | | | | | | | 0 | -15.051 | 5.156 |

PDB data are of the version of 10/25/2020. Lightgray colored cells indicate either chain B of symmetric dimer or ligand-unbound chain. Distances are in Å.

[$]Peptide or peptide-mimic ligand is defined when a ligand contains more than or equal to two peptide moieties. However, it is admitted that a carbonyl group is replaced by alcohol, and that Cα is in a part of an aromatic group.

[#]The residues with underline indicate that they are of the native sequence.

[%]Data from the PanDDA analysis (115entries) and the entry having swapping, 3iwm, were not used in this study.

[&]HBs: 1 for formed and 0 for not formed. HBs are intra-protomer hydrogen bonds except HB 140-1.

#### Dataset S2. Summary of the ligand-free entries.

| specirs | protein | | | | comments | chain A | | | | | | | | | | | | | | chain B | | | | | | | | | | | | | | dimer | | |
|---|---|---|---|---|---|---|---|---|---|---|---|---|---|---|---|---|---|---|---|---|---|---|---|---|---|---|---|---|---|---|---|---|---|---|---|---|
| | | | | | | PC1 | | | | | C-loop conformation | | | | | | | | | PC1 | | | | | C-loop conformation | | | | | | | | | | | |
| | PDB ID | mutation | residues added to N-term | missing N-term residues | | C-loop | E-loop | helical loop | linker | Domain III | main-chain HB within 138-144 | HB 138-172[&] | HB 139-126[&] | HB 140-1[&] | HB 141-118[&] | HB 143-28[&] | distance 165Cα-140Cβ | distance 165Cα-141Cβ | distance 2-214 | C-loop | E-loop | H-loop | Linker | Domain III | main-chain HB within 138-144 | HB 138-172[&] | HB 139-126[&] | HB 140-1[&] | HB 141-118[&] | HB 143-28[&] | distance 165Cα-140Cβ | distance 165Cα-141Cβ | distance 2-214 | PC1 Domain III | PC2 Domain III | distance between two 285 Cα's |
| SARS-CoV[%] | 1q2w | F305E, Q306G | SL | SLSG of A | | 0.278 | 0.087 | | 1.137 | 5.219 | 141O-144N | 1 | 1 | 0 | 1 | 1 | 7.650 | 11.874 | | 0.463 | 0.272 | | 1.729 | 15.065 | 141O-144N | 1 | 1 | 0 | 0 | 1 | 7.408 | 11.805 | 8.684 | -10.325 | 12.841 | 8.717 |
| | 1uj1 | | | SG of A | | 0.347 | -0.304 | 0.121 | 2.396 | 15.308 | | 1 | 1 | 1 | 1 | 1 | 8.024 | 12.716 | | -7.437 | -2.501 | -0.408 | -1.858 | 1.369 | 138O-141N 139O-142N [illegible] | 0 | 0 | 0 | 0 | 0 | 13.326 | 7.826 | 7.495 | 12.888 | 14.396 | 8.446 |
| | 1uk2 | | | | | -0.739 | -1.139 | -0.188 | 0.027 | 6.582 | | 0 | 0 | 1 | 0 | 1 | 9.119 | 13.142 | 8.081 | -4.879 | -2.099 | -0.724 | 0.412 | 1.699 | 138O-140N | 0 | 0 | 0 | 0 | 0 | 9.335 | 14.134 | 6.703 | 6.776 | 11.641 | 10.291 |
| | 1uk3 | | | SG of A | | -0.456 | -1.656 | 1.736 | 0.147 | 10.022 | | 1 | 0 | 0 | 0 | 1 | 8.949 | 13.583 | | -3.759 | -1.666 | 0.009 | 1.131 | 1.041 | 138O-140N 139O-141N | 0 | 0 | 0 | 0 | 0 | 11.552 | 11.767 | 7.765 | 9.945 | 12.545 | 10.300 |
| | 1z1i | | | | | 1.164 | -3.305 | -0.087 | 0.735 | -12.480 | 141O-143N 141O-144N | 1 | 1 | 1 | 1 | 1 | 7.888 | 11.419 | 7.047 | | | | | | | | | | | | | | | 0 | -8.503 | 6.615 |
| | 1z1j_B | C145A | | | Chain A binds a symmetrically related chain A (301-306) while chain B is in free state. C145A makes the protein inactive. | | | | | | | | | | | | | | | 0.260 | -0.254 | -0.343 | 1.299 | -7.947 | | 0 | 1 | 0 | 1 | 1 | 8.076 | 12.230 | 6.572 | 8.53 | 1.861 | 5.767 |
| | 2a5a | | | | | 0.557 | 0.173 | -0.104 | 1.668 | 4.216 | 141O-144N | 1 | 1 | 1 | 1 | 1 | 8.060 | 11.824 | 7.389 | | | | | | | | | | | | | | | 0.001 | 4.883 | 7.323 |
| | 2bx3 | | | S | | -6.466 | -1.075 | 1.136 | 1.481 | 6.677 | 138O-140N 141O-144N | 0 | 0 | 0 | 0 | 0 | 12.959 | 7.929 | 8.47 | | | | | | | | | | | | | | | -0.001 | 10.701 | 8.055 |
| | 2bx4 | | | SG | | -5.559 | -2.370 | -0.629 | -0.376 | -1.983 | 138O-140N 141O-144N | 1 | 0 | 0 | 0 | 0 | 12.447 | 7.681 | | | | | | | | | | | | | | | | 0 | 0.356 | 6.472 |
| | 2c3s | | | | | 0.556 | -3.945 | -0.031 | -0.420 | -3.520 | 141O-144N | 0 | 0 | 0 | 0 | 1 | 7.637 | 11.672 | 8.194 | | | | | | | | | | | | | | | -0.001 | 0.362 | 6.628 |
| | 2duc | | | | | 0.530 | 0.545 | 0.847 | 2.152 | 12.658 | 141O-144N | 1 | 1 | 1 | 1 | 1 | 7.956 | 11.716 | 7.065 | -4.531 | -0.173 | -0.302 | -1.623 | 2.899 | 141O-144N | 1 | 0 | 0 | 0 | 0 | 8.448 | 12.633 | 7.338 | 11.684 | 10.101 | 8.463 |
| | 2gt7 | | | | | 0.718 | 0.147 | | 2.076 | 10.744 | 141O-144N | 1 | 1 | 1 | 1 | 1 | 7.929 | 11.755 | 8.381 | 0.443 | 0.136 | -0.016 | 1.249 | 2.885 | 141O-144N | 1 | 1 | 1 | 1 | 1 | 8.079 | 11.894 | 7.314 | 10.657 | 9.188 | 8.313 |
| | 2gt8 | | A | ASG | | -7.697 | -1.377 | 1.613 | 1.098 | 7.986 | 138O-140N 141O-144N | 0 | 0 | 0 | 0 | 0 | 12.958 | 7.926 | | | | | | | | | | | | | | | | -0.001 | 11.365 | 9.131 |
| | 2gz9 | | | | | 0.480 | 0.486 | -0.499 | 1.540 | 4.689 | 141O-144N | 1 | 1 | 1 | 1 | 1 | 7.810 | 11.646 | 7.581 | | | | | | | | | | | | | | | 0.001 | 6.026 | 7.767 |
| | 2h2z | | | | | 0.609 | 0.576 | -0.399 | 1.206 | 6.193 | 141O-144N | 1 | 1 | 1 | 1 | 1 | 8.137 | 11.908 | 7.749 | | | | | | | | | | | | | | | 0.001 | 6.409 | 7.968 |
| | 2qc2 | N214A | GS | G of A,GS of B | N214A breaks HB 214-2 to destabilizes the active state. | -0.259 | -0.581 | -0.306 | 2.760 | 12.571 | | 1 | 0 | 0 | 0 | 0 | 7.604 | 11.472 | 8.266 | -1.561 | -2.854 | 0.317 | 0.526 | 0.061 | | 0 | 0 | 0 | 1 | 1 | 7.848 | 12.251 | 7.151 | 10.82 | 9.722 | 8.557 |
| | 3aw1 | R188I | | | R188I prevents from proteolysis and increases the activity. | 0.684 | -1.739 | -0.546 | -1.844 | -2.379 | 141O-144N | 1 | 1 | 1 | 1 | 1 | 7.808 | 11.947 | 6.922 | 0.529 | 0.038 | -0.815 | 1.092 | 12.289 | 141O-144N | 1 | 1 | 1 | 1 | 1 | 8.094 | 11.934 | 7.807 | -9.809 | 4.175 | 7.926 |
| | 3e91 | S284A,T285A,I286A | | | T285A is the same mutation as SARS-CoV-2 3CLpro. | 0.382 | -0.287 | 0.326 | 1.146 | 8.903 | 141O-144N | 1 | 1 | 1 | 1 | 1 | 7.372 | 11.395 | 8.14 | 0.190 | -1.063 | 0.426 | 0.629 | -9.126 | 141O-144N | 1 | 1 | 0 | 1 | 1 | 7.458 | 11.224 | 6.973 | 14.701 | 0.145 | 6.367 |
| | 3ea7 | S284A,T285A,I286A | | | T285A is the same mutation as SARS-CoV-2 3CLpro. | 0.694 | -0.058 | -1.132 | 2.346 | 3.286 | 141O-144N | 1 | 1 | 1 | 1 | 1 | 7.669 | 11.758 | 7.767 | 0.404 | -0.861 | -0.985 | 0.975 | -10.042 | 141O-144N | 1 | 1 | 1 | 1 | 1 | 7.502 | 11.572 | 6.493 | 14.459 | -0.552 | 6.180 |
| | 3ea8 | S284A,T285A,I286A | | | T285A is the same mutation as SARS-CoV-2 3CLpro. | 0.345 | 0.241 | -0.507 | 1.547 | 0.616 | 141O-144N | 1 | 1 | 1 | 1 | 1 | 7.632 | 11.385 | 7.216 | | | | | | | | | | | | | | | -0.001 | -0.411 | 5.375 |
| | 3ea9 | S284A,T285A,I286A,F291A | | | T285A is the same mutation as SARS-CoV-2 3CLpro. | 0.566 | -0.225 | -1.043 | 1.166 | -6.287 | 141O-144N | 1 | 1 | 1 | 1 | 1 | 7.326 | 11.163 | 7.239 | | | | | | | | | | | | | | | -0.001 | -9.603 | 5.356 |
| | 3eaj_AC | S284A,T285A,I286A,F291A | | | Chains A&C and B&D are two symmetric dimers. T285A is the same mutation as SARS-CoV-2 3CLpro. | 0.537 | -0.596 | -1.486 | 0.314 | -6.363 | 141O-144N | 1 | 1 | 1 | 1 | 1 | 7.113 | 11.011 | 6.576 | | | | | | | | | | | | | | | 0.001 | -10.259 | 8.074 |
| | 3eaj_BD | S284A,T285A,I286A,F291A | | | T285A is the same mutation as SARS-CoV-2 3CLpro. | 0.723 | -0.272 | -1.28 | 0.93 | -2.966 | 141O-144N | 1 | 1 | 1 | 1 | 1 | 7.822 | 11.887 | 6.366 | | | | | | | | | | | | | | | 0 | -8.14 | 5.234 |
| | 3f9f | F140A | GS | GSS of A,B | F140A breaks the hydrophobic interactions with H163 and H172 to destabilize the active state. | -10.938 | -2.379 | -0.315 | -1.933 | -2.022 | 139O-143N 141O-144N | 0 | 0 | 0 | 0 | 0 | 10.416 | 13.537 | 7.482 | -11.237 | 0.089 | -0.386 | 0.196 | -1.197 | 139O-143N 141O-144N | 0 | 0 | 0 | 0 | 0 | 10.181 | 13.384 | 8.51 | -0.483 | -0.429 | 9.456 |
| | 3f9g | F140A | GS | GSS of A,GSSSG of B | C-terminal 5 residues do not exist. F140A breaks the hydrophobic interactions with H163 and H172 to destabilize the active state. | -8.002 | -1.266 | -0.389 | -0.222 | -0.990 | | 0 | 0 | 0 | 0 | 0 | 12.698 | | 9.47 | -8.184 | -3.168 | -0.224 | -1.945 | -1.179 | | 1 | 0 | 0 | 0 | 0 | 9.547 | | | -0.187 | 0.609 | 9.756 |
| | 3f9h | F140A | GS | GSS of A, GS of B | F140A breaks the hydrophobic interactions with H163 and H172 to destabilize the active state. | -11.189 | -1.867 | -0.628 | -2.651 | -1.228 | | 0 | 0 | 0 | 0 | 0 | 11.559 | 14.061 | 7.399 | -11.715 | -0.211 | -0.154 | -0.117 | -1.160 | | 0 | 0 | 0 | 0 | 0 | 13.090 | 13.823 | 8.472 | 0.339 | -0.118 | 9.720 |
| | 3fzd | N28A | | SG | N28A breaks both HB 145-28 and HB 143-28 to destabilize the active state. Residues 139-141 in C-loop are missing. C145 and C117 has S-S bond. C-terminal 5 residues do not exist. | | -3.991 | 0.485 | -0.570 | -18.669 | | 0 | 0 | 0 | 0 | 0 | | | | | | | | | | | | | | | | | | -0.001 | -12.071 | 6.906 |
| | 3m3s | N214A | | | N214A breaks HB 214-2. | 0.785 | -0.214 | -1.060 | 2.191 | 9.714 | 141O-144N | 1 | 1 | 1 | 1 | 1 | 7.802 | 11.773 | 8.58 | -0.011 | 0.197 | -0.513 | 1.231 | 4.544 | 141O-144N | 1 | 1 | 1 | 1 | 1 | 8.016 | 11.816 | 7.871 | 8.248 | 10.49 | 8.370 |
| | 3m3v | S284A,T285A,I286A | GS | GS of A, G of B | T285A is the same mutation as SARS-CoV-2 3CLpro. | -0.041 | -0.655 | -0.438 | 1.384 | 6.051 | | 1 | 1 | 0 | 1 | 0 | 7.790 | 11.864 | 8.546 | -0.281 | -1.883 | 0.028 | 1.992 | -8.071 | | 0 | 0 | 0 | 1 | 1 | 7.841 | 11.445 | 6.991 | 11.989 | 1.341 | 6.898 |
| | 3vb3 | | | | | 0.425 | -1.155 | -1.111 | 0.253 | -10.374 | 141O-144N | 1 | 1 | 1 | 1 | 1 | 7.958 | 12.034 | 6.587 | 0.454 | 0.117 | 2.655 | 2.291 | 6.127 | 141O-144N | 1 | 1 | 1 | 1 | 1 | 7.761 | 11.672 | 8.143 | -13.491 | 1.334 | 6.698 |
| | 4hi3 | R298A | M | M of A, B | R298A impairs dimerization. Crystalized with 1mM substrate (ligand induced dimerization), but still density was not observed. After C-term, LE+6His are added. | 0.477 | -0.544 | 0.333 | 1.012 | 2.519 | 141O-144N | 1 | 1 | 1 | 1 | 1 | 7.456 | 11.379 | 10.639 | 0.205 | -0.557 | 0.937 | -1.804 | -9.058 | 141O-144N | 1 | 1 | 1 | 1 | 1 | 7.484 | 11.111 | 6.61 | 7.898 | 2.832 | 6.455 |
| | 5b6o_B | C145A | | | Chain A binds a symmetrically related chain A (301-310), while chain B is in free state. C145A makes the protein inactive. | | | | | | | | | | | | | | | 0.410 | 0.098 | 0.735 | 0.995 | 6.439 | 141O-144N | 1 | 1 | 1 | 1 | 1 | 8.158 | 11.982 | 7.478 | -21.195 | -3.44 | 6.189 |
| mono | 2pwx | G11A | GS | GSGSF | Monomeric in crystal. G11A destroys the tight binding in the dimerization interface. | -7.626 | -2.426 | 1.488 | 4.155 | | 138O-141N 139O-142N | 0 | 0 | | 0 | 0 | 14.180 | 8.704 | | | | | | | | | | | | | | | | | | |
| | 2qcy | R298A | | | Monomeric in crystal. R298A impairs dimerization. | -7.877 | -1.606 | 0.993 | 3.573 | -154.30 | 138O-141N 139O-142N 141O-144N | 0 | 0 | | 0 | 0 | 13.672 | 8.315 | 9.831 | | | | | | | | | | | | | | | | | |
| | 3f9e | S139A | GS | GSSG | Monomeric in crystal. S139A impairs dimerization. | -7.866 | -2.720 | 1.536 | 3.898 | -156.16 | 138O-141N 139O-142N 141O-144N | 0 | 0 | | 0 | 0 | 14.541 | 9.141 | | | | | | | | | | | | | | | | | | |
| | 3m3t | R298A | GS | | Monomeric in crystal.R298A impairs dimerization. | -7.400 | -2.202 | 0.048 | 3.211 | -153.83 | 138O-141N 139O-142N 141O-144N | 0 | 0 | | 0 | 0 | 14.174 | 8.847 | 10.111 | | | | | | | | | | | | | | | | | |
| SARS-CoV-2 | 6m03 | | | | | 0.611 | -1.789 | -0.808 | -1.452 | -6.662 | 141O-144N | 1 | 1 | 1 | 1 | 1 | 7.614 | 11.577 | 6.314 | | | | | | | | | | | | | | | -0.001 | -8.573 | 5.227 |
| | 6m2q | | | | | 0.798 | -2.391 | -1.098 | -2.643 | -4.920 | 141O-144N | 1 | 1 | 1 | 1 | 1 | 7.689 | 11.762 | 6.422 | | | | | | | | | | | | | | | 0 | -7.188 | 5.231 |
| | 6wqf | | | | | 0.633 | -1.865 | -1.559 | -2.187 | -2.464 | 141O-144N | 1 | 1 | 1 | 1 | 1 | 7.845 | 11.555 | 6.431 | | | | | | | | | | | | | | | 0.001 | -5.765 | 5.684 |
| | 6wtm | | | | | 0.978 | -2.470 | -1.268 | -2.894 | -5.314 | 141O-144N | 1 | 1 | 1 | 1 | 1 | 7.297 | 11.335 | 6.481 | 0.928 | -2.561 | -1.036 | -2.518 | -6.130 | 141O-144N | 1 | 1 | 1 | 1 | 1 | 7.380 | 11.334 | 6.161 | 1.098 | -7.571 | 5.418 |
| | 6xb0 | | | | C145 is di-oxydized, 2CO (SOO). | 0.672 | -1.936 | -1.468 | -2.108 | -2.746 | 141O-144N | 1 | 1 | 1 | 1 | 1 | 7.723 | 11.519 | 6.387 | | | | | | | | | | | | | | | 0 | -6.211 | 5.470 |
| | 6xhu | | | | | 1.011 | -2.824 | -0.483 | -2.575 | -7.561 | 141O-144N 142O-144N | 1 | 1 | 1 | 1 | 1 | 7.655 | 11.429 | 6.473 | 0.688 | -2.151 | -1.329 | -3.017 | -2.107 | 141O-144N | 1 | 1 | 1 | 1 | 1 | 7.643 | 11.412 | 6.527 | -2.874 | -6.344 | 5.706 |
| | 6xkf | | | | C145 is mono-oxydized, CSX. | 0.616 | -2.060 | -1.318 | -3.305 | -3.840 | 141O-144N | 1 | 1 | 1 | 1 | 1 | 7.333 | 11.385 | 6.658 | 0.557 | -2.220 | -1.292 | -2.859 | -5.935 | 141O-144N | 1 | 1 | 1 | 1 | 1 | 7.537 | 11.452 | 6.252 | 1.856 | -8.154 | 5.499 |
| | 6xkh | | | | C145 is di-oxydized, CSD (OSO). | 0.519 | -0.321 | 0.488 | 2.373 | -1.963 | 141O-144N | 1 | 1 | 1 | 1 | 1 | 7.674 | 11.323 | 6.736 | | | | | | | | | | | | | | | 0 | -1.966 | 5.265 |
| | 6xoa_BD | C145S | | | Tetramer (two asym dimers, AB&CD). Chains A&C bind symmetrically related C-term of C&A, respectively. Chains BD are free. Dimer data are of CD. C145S makes the protein inactive. | 0.266 | 0.167 | -1.656 | 0.317 | 3.339 | 141O-144N | 1 | 1 | 1 | 1 | 1 | 8.027 | 11.767 | 7.227 | 0.860 | 0.160 | -0.524 | -0.133 | 1.765 | 141O-144N | 0 | 1 | 1 | 1 | 1 | 7.510 | 11.278 | 6.245 | -5.535 | -3.383 | 5.805 |
| | 6y2e | | | | | 0.765 | -2.218 | -1.382 | -2.748 | -5.350 | 141O-144N | 1 | 1 | 1 | 1 | 1 | 7.745 | 11.778 | 6.350 | | | | | | | | | | | | | | | -0.001 | -8.21 | 5.216 |
| | 6y84 | | | | | 1.040 | -1.365 | -1.495 | -0.534 | -6.701 | 141O-144N | 1 | 1 | 1 | 1 | 1 | 7.412 | 11.240 | 5.959 | | | | | | | | | | | | | | | 0 | -11.786 | 5.077 |
| | 6yb7 | | | | | 1.077 | -1.514 | -1.486 | -0.499 | -6.374 | 141O-144N | 1 | 1 | 1 | 1 | 1 | 7.447 | 11.361 | 5.947 | | | | | | | | | | | | | | | 0 | -12.153 | 5.085 |
| | 7bro | | G | GS | | 1.286 | -4.201 | 0.826 | -1.805 | -13.749 | 141O-144N | 0 | 0 | 0 | 0 | 1 | 7.656 | 11.822 | 8.738 | | | | | | | | | | | | | | | 0.001 | -9.963 | 5.796 |
| | 7c2q | S1,G2 del | S1,G2 del | S1,G2 del | C-terminal 8 residues do not exist. Deletion of S1 and G2 imapirs dimerization. | | -0.571 | -0.118 | 2.372 | | | 1 | 1 | 0 | 0 | 1 | | | | | -0.256 | | 1.908 | | | 0 | 0 | 0 | 0 | 1 | | | | 0.559 | 11.858 | 7.284 |
| | 7c2y | | | SGF of A, SG of B | C-terminal 8 residues do not exist. | | -0.571 | -0.128 | 2.240 | | | 1 | 1 | 0 | 0 | 0 | 7.824 | | | | -0.211 | | 1.967 | | | 0 | 0 | 0 | 0 | 0 | | | | 0.235 | 12.501 | 7.812 |
| | 7cwb | | | | XFEL. | 1.169 | -3.185 | -0.480 | -2.704 | -4.164 | 141O-144N | 1 | 1 | 1 | 1 | 1 | 7.615 | 11.716 | 6.276 | | | | | | | | | | | | | | | 0 | -7.107 | 5.623 |
| | 7cwc | | GSHM | GSHM of A,GSHMSG of B | XFEL. | 0.138 | -1.094 | -0.860 | -0.261 | 9.187 | 141O-144N | 0 | 0 | 0 | 1 | 1 | 8.272 | 12.225 | 9.149 | 0.706 | -0.126 | 0.253 | -0.267 | 1.260 | 141O-144N | 0 | 1 | 0 | 1 | 1 | 7.911 | 11.829 | | 3.31 | 14.361 | 8.884 |
| | 7jfq | | | | | 0.617 | -0.875 | -1.219 | 0.549 | -8.191 | 141O-144N | 1 | 1 | 1 | 1 | 1 | 7.971 | 11.825 | 6.581 | | | | | | | | | | | | | | | -0.002 | -8.645 | 5.296 |
| | 7joy_A | C145A | | | Chain B binds a symmetrically related C-term of chain B, while chain A are in free state. C145A makes the protein inactive. | 0.528 | 0.078 | -0.005 | 0.484 | 5.424 | 141O-144N | 1 | 1 | 1 | 1 | 1 | 7.365 | 10.859 | 6.533 | | | | | | | | | | | | | | | -1.691 | 5.914 | 5.965 |
| | 7jp1 | | | | | 0.874 | -1.179 | -1.955 | -0.886 | -3.401 | 141O-144N | 1 | 1 | 1 | 1 | 1 | 7.860 | 11.797 | 6.437 | | | | | | | | | | | | | | | -0.002 | -7.179 | 5.272 |
| | 7jr3 | | | | | 0.512 | 0.199 | 0.205 | 1.933 | -7.733 | 141O-144N | 1 | 1 | 1 | 1 | 1 | 7.681 | 11.348 | 6.555 | | | | | | | | | | | | | | | -0.001 | -10.682 | 5.579 |
| | 7jr4 | | | | K61 is methylated, MLZ. | 0.427 | 0.249 | | 1.728 | -7.280 | 141O-144N | 1 | 1 | 1 | 1 | 1 | 7.587 | 11.343 | 6.525 | | | | | | | | | | | | | | | 0 | -11.003 | 5.294 |
| | 7jun | | | | Neutron diffraction determining the H positions. | 0.816 | -2.226 | -1.164 | -2.774 | -3.892 | 141O-144N | 1 | 1 | 1 | 1 | 1 | 7.595 | 11.466 | 6.499 | | | | | | | | | | | | | | | -0.001 | -5.745 | 5.586 |
| | 7jvz | | | | XFEL. | 0.542 | -1.830 | -1.301 | -2.281 | -3.543 | 141O-144N | 1 | 1 | 1 | 1 | 1 | 7.700 | 11.720 | 6.652 | | | | | | | | | | | | | | | -0.001 | -3.708 | 5.849 |
| | 7k3t | | | | | 1.200 | -1.820 | -1.498 | -0.537 | -6.848 | 141O-144N | 1 | 1 | 1 | 1 | 1 | 7.277 | 11.381 | 5.717 | | | | | | | | | | | | | | | 0 | -12.684 | 5.105 |
| | 7kfi | | GAM | GAM of B | | -1.587 | -0.209 | -0.341 | 1.858 | 11.263 | | 0 | 0 | 0 | 1 | 0 | 14.216 | 9.164 | 10.172 | -10.762 | 1.168 | 2.003 | 2.270 | 12.321 | 139O-142N | 0 | 0 | 0 | 0 | 0 | 9.731 | 13.923 | 7.78 | -3.175 | 19.214 | 9.858 |
| | 7khp_A | | | | Chain B binds a symmetrically related C-term of chain B, while chain A are in free state. | 0.498 | 0.364 | 0.231 | 0.525 | 3.636 | 141O-144N | 1 | 1 | 1 | 1 | 1 | 7.583 | 11.273 | 6.507 | | | | | | | | | | | | | | | -3.358 | 5.057 | 5.710 |

PDB data are of the version of 10/25/2020. Lightgray colored cells indicate either non-exsisting chain B of symmetric dimer or ligand-bound chains. Cells colored in darkgray are those with undetermined data due to missing residues. Distances are in Å.

[%]The entries containing only domain III, 2k7x, 2liz, and 3ebn, were not used in this study.

[&]HBs: 1 for formed and 0 for not formed. HBs are intra-protomer hydrogen bonds except HB 140-1.

**Dataset S3. Ligand binding.**

| species | PDBid | MC | interface | | | H-loop | | | C-loop | | | | interface | | I&II | | E-loop | | | | | I&II | Linker | | | | | Ligand interactions |
|---|---|---|---|---|---|---|---|---|---|---|---|---|---|---|---|---|---|---|---|---|---|---|---|---|---|---|---|---|
| | | BC | T25 | T26 | L27 | H41 | M49 | Y54 | F140 | L141 | N142 | G143 | S144 | C145 | H163 | H164 | M165 | E166 | L167 | P168 | H172 | D187 | R188 | Q189 | T190 | A191 | Q192 | |
| SARS-CoV | 1uk4 | A | 1 | 0 | 0 | 1 | 1 | 0 | 11 | 1 | 1 | 1 | 11 | 21 | 11 | 0 | 1 | 11 | 0 | 0 | 1 | 0 | 0 | 11 | 0 | 0 | 1 | CE |
| | 1wof | A | 1 | 1 | 0 | 1 | 1 | 0 | 1 | 1 | 1 | 1 | 0 | 21 | 11 | 11 | 1 | 11 | 0 | 1 | 1 | 1 | 1 | 11 | 11 | 1 | 1 | CEL |
| | 1z1j | A | 0 | 0 | 0 | 11 | 1 | 0 | 11 | 1 | 0 | 11 | 1 | 11 | 11 | 11 | 1 | 11 | 0 | 1 | 1 | 1 | 1 | 11 | 0 | 1 | 11 | CEHL |
| | 2a5i | AB | 1 | 0 | 0 | 1 | 1 | 1 | 11 | 1 | 1 | 11 | 1 | 21 | 11 | 11 | 1 | 11 | 0 | 1 | 1 | 1 | 1 | 11 | 1 | 1 | 1 | CEHL |
| | 2a5k | A | 1 | 0 | 0 | 1 | 1 | 1 | 11 | 1 | 1 | 11 | 1 | 21 | 11 | 11 | 1 | 11 | 1 | 0 | 1 | 1 | 1 | 11 | 1 | 0 | 1 | CEHL |
| | 2alv | AB | 0 | 1 | 1 | 1 | 0 | 1 | 11 | 1 | 1 | 1 | 0 | 21 | 11 | 11 | 1 | 11 | 0 | 1 | 1 | 1 | 1 | 1 | 1 | 1 | 1 | CEL |
| | 2amd | A | 1 | 1 | 0 | 1 | 1 | 0 | 1 | 0 | 1 | 1 | 0 | 21 | 11 | 11 | 1 | 11 | 0 | 1 | 1 | 1 | 1 | 11 | 1 | 1 | 1 | EL |
| | 2amq | A | 1 | 1 | 0 | 1 | 1 | 0 | 11 | 1 | 1 | 1 | 0 | 21 | 11 | 11 | 1 | 11 | 0 | 1 | 1 | 1 | 1 | 11 | 11 | 1 | 1 | CEL |
| | 2d2d | A | 0 | 1 | 1 | 1 | 1 | 0 | 11 | 1 | 1 | 1 | 1 | 21 | 11 | 11 | 1 | 11 | 0 | 1 | 1 | 0 | 0 | 1 | 1 | 1 | 1 | CEL |
| | 2gtb | AB | 1 | 0 | 0 | 1 | 1 | 1 | 11 | 1 | 1 | 11 | 1 | 21 | 11 | 11 | 1 | 11 | 1 | 1 | 1 | 1 | 1 | 11 | 1 | 0 | 1 | CEHL |
| | 2gx4 | AB | 0 | 0 | 0 | 1 | 1 | 1 | 11 | 0 | 0 | 11 | 0 | 21 | 11 | 11 | 1 | 11 | 1 | 1 | 1 | 1 | 1 | 11 | 1 | 1 | 1 | CEHL |
| | 2gz7 | AB | 0 | 0 | 0 | 1 | 0 | 0 | 0 | 0 | 0 | 0 | 0 | 1 | 0 | 1 | 1 | 1 | 0 | 0 | 0 | 1 | 1 | 0 | 0 | 0 | 1 | |
| | 2gz8 | AB | 0 | 0 | 0 | 1 | 0 | 0 | 0 | 1 | 11 | 1 | 1 | 11 | 0 | 0 | 1 | 1 | 0 | 1 | 0 | 0 | 0 | 1 | 1 | 0 | 1 | C |
| | 2hob | AB | 1 | 1 | 1 | 1 | 1 | 0 | 11 | 1 | 1 | 1 | 1 | 21 | 11 | 11 | 1 | 11 | 0 | 1 | 1 | 1 | 1 | 11 | 11 | 1 | 1 | RCEL |
| | 2op9 | A | 0 | 0 | 0 | 11 | 1 | 0 | 0 | 1 | 1 | 11 | 1 | 11 | 0 | 0 | 1 | 11 | 0 | 0 | 0 | 1 | 1 | 1 | 0 | 0 | 0 | CH |
| | 2q6g | A | 1 | 11 | 0 | 0 | 1 | 1 | 11 | 1 | 1 | 11 | 0 | 11 | 11 | 11 | 1 | 11 | 0 | 1 | 1 | 1 | 0 | 11 | 11 | 1 | 0 | RCEL |
| | 2qiq | AB | 0 | 1 | 1 | 1 | 1 | 1 | 11 | 1 | 1 | 1 | 0 | 11 | 11 | 11 | 1 | 11 | 0 | 1 | 1 | 1 | 1 | 1 | 11 | 0 | 1 | CEHL |
| | 2v6n | AB | 0 | 0 | 0 | 1 | 1 | 0 | 0 | 0 | 0 | 0 | 0 | 21 | 0 | 1 | 1 | 0 | 0 | 0 | 0 | 1 | 1 | 1 | 0 | 0 | 0 | |
| | 2vj1 | A | 0 | 0 | 0 | 1 | 21 | 0 | 0 | 0 | 0 | 0 | 0 | 0 | 0 | 0 | 21 | 0 | 0 | 0 | 0 | 1 | 1 | 1 | 0 | 0 | 0 | H |
| | 2z3c | AB | 0 | 0 | 0 | 1 | 0 | 0 | 11 | 1 | 1 | 11 | 1 | 21 | 11 | 11 | 1 | 11 | 0 | 0 | 1 | 0 | 0 | 0 | 0 | 0 | 0 | CE |
| | 2z3d | AB | 0 | 0 | 0 | 11 | 1 | 0 | 11 | 1 | 1 | 11 | 1 | 21 | 11 | 11 | 1 | 11 | 1 | 1 | 1 | 0 | 0 | 0 | 1 | 0 | 1 | CEH |
| | 2z3e | AB | 0 | 0 | 0 | 1 | 1 | 0 | 11 | 1 | 1 | 11 | 1 | 21 | 11 | 11 | 1 | 11 | 1 | 1 | 1 | 0 | 0 | 11 | 1 | 0 | 1 | CEL |
| | 2z94 | AB | 0 | 0 | 0 | 1 | 0 | 0 | 0 | 0 | 1 | 0 | 0 | 20 | 0 | 0 | 0 | 0 | 0 | 0 | 0 | 0 | 0 | 0 | 0 | 0 | 0 | |
| | 2z9g | AB | 0 | 0 | 0 | 1 | 1 | 20 | 0 | 0 | 0 | 0 | 0 | 0 | 0 | 0 | 1 | 0 | 0 | 0 | 0 | 1 | 1 | 1 | 0 | 0 | 0 | H |
| | 2z9j | A | 0 | 0 | 1 | 1 | 0 | 0 | 0 | 0 | 0 | 0 | 0 | 20 | 0 | 0 | 0 | 0 | 0 | 0 | 0 | 0 | 0 | 0 | 0 | 0 | 0 | |
| | 2z9k | A | 1 | 0 | 1 | 1 | 0 | 0 | 0 | 0 | 0 | 0 | 0 | 20 | 0 | 0 | 0 | 0 | 0 | 0 | 0 | 0 | 0 | 0 | 0 | 0 | 0 | |
| | 2z9l | A | 0 | 0 | 0 | 1 | 0 | 0 | 0 | 0 | 0 | 0 | 0 | 20 | 0 | 0 | 0 | 0 | 0 | 0 | 0 | 0 | 0 | 0 | 0 | 0 | 0 | |
| | 2zu4 | AB | 1 | 1 | 1 | 1 | 1 | 0 | 11 | 1 | 1 | 11 | 0 | 21 | 11 | 11 | 1 | 11 | 0 | 1 | 1 | 1 | 0 | 11 | 1 | 1 | 1 | RCEL |
| | 2zu5 | AB | 1 | 1 | 1 | 1 | 1 | 0 | 11 | 1 | 1 | 1 | 0 | 21 | 11 | 11 | 1 | 11 | 0 | 1 | 1 | 1 | 0 | 11 | 1 | 1 | 1 | RCEL |
| | 3atw | A | 0 | 0 | 0 | 1 | 1 | 1 | 1 | 1 | 1 | 11 | 1 | 11 | 11 | 11 | 1 | 11 | 1 | 1 | 0 | 1 | 1 | 1 | 11 | 1 | 0 | CEHL |
| | 3avz | AB | 0 | 0 | 0 | 1 | 1 | 0 | 1 | 0 | 1 | 11 | 1 | 21 | 11 | 11 | 1 | 11 | 0 | 1 | 0 | 1 | 1 | 1 | 11 | 1 | 0 | CEL |
| | 3aw0 | AB | 0 | 0 | 0 | 1 | 1 | 0 | 1 | 1 | 1 | 1 | 1 | 11 | 11 | 1 | 1 | 11 | 0 | 1 | 0 | 1 | 0 | 0 | 0 | 1 | 1 | CE |
| | 3d62 | AB | 0 | 0 | 0 | 11 | 1 | 0 | 0 | 0 | 1 | 11 | 1 | 21 | 0 | 1 | 1 | 0 | 0 | 0 | 0 | 0 | 1 | 1 | 0 | 0 | 0 | CH |
| | 3sn8 | AB | 0 | 0 | 0 | 0 | 1 | 0 | 1 | 1 | 1 | 11 | 1 | 21 | 0 | 1 | 1 | 11 | 0 | 0 | 0 | 1 | 1 | 1 | 0 | 0 | 0 | C |
| | 3sna | AB | 0 | 0 | 0 | 11 | 1 | 0 | 11 | 1 | 1 | 11 | 11 | 21 | 11 | 0 | 1 | 11 | 0 | 1 | 0 | 0 | 0 | 0 | 11 | 1 | 0 | CEH |
| | 3snb | AB | 0 | 0 | 0 | 1 | 1 | 0 | 11 | 1 | 0 | 1 | 0 | 21 | 11 | 1 | 1 | 11 | 0 | 1 | 0 | 0 | 1 | 1 | 11 | 0 | 0 | CEL |
| | 3snc | AB | 0 | 0 | 0 | 1 | 1 | 0 | 11 | 1 | 1 | 0 | 0 | 21 | 11 | 11 | 1 | 11 | 0 | 1 | 0 | 0 | 0 | 1 | 1 | 1 | 0 | CE |
| | 3snd | A | 0 | 0 | 0 | 0 | 0 | 0 | 11 | 1 | 0 | 1 | 1 | 21 | 11 | 11 | 1 | 11 | 0 | 0 | 1 | 0 | 0 | 0 | 0 | 0 | 0 | CE |
| | 3sne | AB | 0 | 0 | 0 | 1 | 1 | 0 | 11 | 1 | 1 | 11 | 0 | 21 | 11 | 1 | 1 | 11 | 1 | 1 | 0 | 1 | 0 | 1 | 1 | 0 | 0 | CE |
| | 3szn | AB | 0 | 1 | 1 | 1 | 1 | 0 | 11 | 1 | 1 | 11 | 1 | 21 | 11 | 11 | 1 | 11 | 0 | 0 | 1 | 1 | 1 | 1 | 0 | 0 | 0 | CE |
| | 3tit | AB | 0 | 0 | 0 | 0 | 1 | 0 | 11 | 1 | 1 | 11 | 1 | 21 | 11 | 11 | 1 | 11 | 0 | 0 | 1 | 1 | 1 | 1 | 0 | 0 | 0 | CE |
| | 3tiu | AB | 1 | 0 | 0 | 1 | 0 | 1 | 11 | 1 | 1 | 11 | 1 | 21 | 11 | 11 | 1 | 11 | 0 | 1 | 1 | 1 | 1 | 11 | 1 | 0 | 1 | CEL |
| | 3tns | AB | 0 | 0 | 0 | 1 | 1 | 1 | 11 | 0 | 1 | 1 | 1 | 21 | 11 | 11 | 1 | 11 | 0 | 1 | 1 | 1 | 1 | 11 | 1 | 1 | 1 | CEHL |
| | 3tnt | AB | 0 | 0 | 0 | 1 | 1 | 1 | 11 | 1 | 1 | 11 | 1 | 21 | 11 | 11 | 1 | 11 | 0 | 1 | 1 | 1 | 1 | 11 | 1 | 1 | 1 | CEHL |
| | 3v3m | AB | 1 | 1 | 0 | 1 | 1 | 0 | 1 | 1 | 1 | 11 | 11 | 1 | 11 | 1 | 1 | 11 | 0 | 0 | 0 | 1 | 0 | 0 | 0 | 0 | 0 | CE |
| | 3vb4 | A | 0 | 0 | 1 | 1 | 1 | 0 | 11 | 1 | 0 | 0 | 0 | 21 | 11 | 11 | 1 | 11 | 1 | 1 | 1 | 1 | 1 | 11 | 11 | 1 | 1 | CEL |
| | 3vb5 | A | 0 | 0 | 0 | 0 | 0 | 0 | 1 | 1 | 0 | 1 | 1 | 21 | 11 | 11 | 1 | 11 | 1 | 1 | 1 | 0 | 0 | 11 | 11 | 1 | 1 | CEL |
| | 3vb6 | A | 0 | 0 | 0 | 1 | 1 | 0 | 11 | 1 | 11 | 1 | 1 | 21 | 11 | 11 | 1 | 11 | 0 | 0 | 1 | 1 | 1 | 1 | 1 | 0 | 1 | CEL |
| | 3vb7 | A | 0 | 0 | 0 | 1 | 1 | 0 | 11 | 1 | 0 | 1 | 1 | 21 | 11 | 11 | 1 | 11 | 0 | 1 | 1 | 1 | 0 | 11 | 1 | 0 | 1 | CEL |
| | 4mds | AB | 0 | 0 | 0 | 1 | 1 | 0 | 1 | 1 | 1 | 0 | 0 | 11 | 11 | 0 | 1 | 11 | 1 | 1 | 0 | 0 | 1 | 1 | 0 | 0 | 0 | E |
| | 4tww | A | 0 | 0 | 0 | 1 | 1 | 0 | 1 | 1 | 1 | 11 | 1 | 11 | 11 | 11 | 0 | 1 | 0 | 0 | 0 | 1 | 1 | 1 | 0 | 0 | 0 | CE |
| | 4twy | AB | 1 | 0 | 0 | 1 | 1 | 0 | 1 | 1 | 1 | 11 | 1 | 21 | 11 | 10 | 0 | 1 | 0 | 0 | 0 | 1 | 0 | 0 | 0 | 0 | 0 | CE |
| | 4wy3 | AB | 0 | 0 | 0 | 1 | 1 | 0 | 1 | 1 | 1 | 1 | 1 | 11 | 11 | 1 | 1 | 1 | 0 | 0 | 1 | 1 | 1 | 1 | 0 | 0 | 0 | CE |
| | 5b6o | A | 1 | 11 | 0 | 11 | 1 | 0 | 1 | 1 | 1 | 11 | 10 | 11 | 11 | 10 | 1 | 11 | 1 | 1 | 1 | 1 | 1 | 11 | 1 | 1 | 1 | RCEHL |

| PDBid | MC | interface | | | H-loop | | | C-loop | | | | interface | | I&II | | E-loop | | | | | I&II | Linker | | | | | Ligand interactions |
|---|---|---|---|---|---|---|---|---|---|---|---|---|---|---|---|---|---|---|---|---|---|---|---|---|---|---|---|
| | BC | T25 | T26 | L27 | H41 | M49 | Y54 | F140 | L141 | N142 | G143 | S144 | C145 | H163 | H164 | M165 | E166 | L167 | P168 | H172 | D187 | R188 | Q189 | T190 | A191 | Q192 | |
| 1uk4 | B | 1 | 0 | 0 | 11 | 1 | 0 | 0 | 0 | 0 | 1 | 1 | 21 | 0 | 1 | 1 | 11 | 0 | 1 | 0 | 0 | 11 | 1 | 11 | 0 | 11 | HL |
| 1wof | B | 0 | 1 | 0 | 1 | 1 | 0 | 1 | 0 | 1 | 11 | 0 | 21 | 11 | 11 | 1 | 11 | 0 | 1 | 1 | 1 | 1 | 11 | 11 | 1 | 1 | CEL |
| 2a5k | B | 1 | 0 | 0 | 1 | 1 | 1 | 11 | 1 | 1 | 11 | 1 | 21 | 11 | 11 | 1 | 11 | 0 | 1 | 1 | 1 | 1 | 11 | 1 | 1 | 1 | CEHL |
| 2amd | B | 1 | 1 | 1 | 1 | 1 | 0 | 1 | 0 | 1 | 1 | 0 | 21 | 11 | 11 | 1 | 11 | 0 | 1 | 1 | 1 | 1 | 11 | 1 | 1 | 1 | REL |
| 2amq | B | 1 | 1 | 1 | 1 | 0 | 0 | 1 | 0 | 0 | 1 | 1 | 21 | 11 | 11 | 1 | 11 | 0 | 1 | 1 | 1 | 1 | 11 | 11 | 0 | 1 | REL |
| 2d2d | B | 0 | 1 | 1 | 1 | 1 | 0 | 11 | 1 | 1 | 1 | 1 | 21 | 11 | 11 | 1 | 11 | 0 | 1 | 1 | 0 | 0 | 11 | 1 | 1 | 1 | CEL |
| 2op9 | B | 0 | 0 | 0 | 11 | 1 | 0 | 0 | 1 | 0 | 11 | 1 | 21 | 0 | 1 | 1 | 11 | 0 | 0 | 0 | 1 | 1 | 0 | 0 | 0 | 0 | CH |
| 2q6g | B | 1 | 11 | 1 | 1 | 1 | 1 | 11 | 1 | 1 | 11 | 0 | 11 | 11 | 1 | 1 | 11 | 0 | 1 | 1 | 1 | 1 | 11 | 11 | 0 | 1 | RCEHL |
| 2vj1 | B | 0 | 0 | 0 | 0 | 0 | 0 | 0 | 1 | 1 | 1 | 0 | 21 | 1 | 0 | 0 | 21 | 0 | 0 | 0 | 0 | 0 | 0 | 0 | 0 | 0 | |
| 2z9j | B | 0 | 0 | 0 | 1 | 0 | 0 | 0 | 0 | 0 | 0 | 0 | 20 | 0 | 1 | 0 | 0 | 0 | 0 | 0 | 0 | 0 | 0 | 0 | 0 | 0 | |
| 2z9k | B | 0 | 0 | 1 | 1 | 0 | 0 | 0 | 0 | 0 | 0 | 0 | 21 | 0 | 1 | 0 | 0 | 0 | 0 | 0 | 0 | 0 | 0 | 0 | 0 | 0 | |
| 2z9l | B | 0 | 0 | 1 | 1 | 0 | 0 | 0 | 0 | 0 | 0 | 0 | 20 | 0 | 0 | 0 | 0 | 0 | 0 | 0 | 0 | 0 | 0 | 0 | 0 | 0 | |
| 3atw | B | 0 | 0 | 0 | 1 | 1 | 0 | 1 | 1 | 1 | 11 | 1 | 11 | 11 | 11 | 1 | 11 | 1 | 1 | 0 | 1 | 1 | 1 | 11 | 0 | 0 | CEL |
| 3snd | B | 0 | 0 | 0 | 0 | 1 | 0 | 11 | 1 | 1 | 11 | 0 | 21 | 11 | 11 | 1 | 11 | 0 | 0 | 0 | 1 | 0 | 0 | 0 | 0 | 0 | CE |
| 3vb4 | B | 0 | 0 | 0 | 1 | 1 | 1 | 11 | 11 | 0 | 0 | 1 | 21 | 11 | 11 | 1 | 11 | 1 | 0 | 1 | 0 | 0 | 11 | 11 | 1 | 0 | CEHL |
| 3vb5 | B | 0 | 0 | 0 | 1 | 1 | 0 | 11 | 1 | 0 | 1 | 0 | 21 | 11 | 11 | 1 | 11 | 1 | 1 | 1 | 0 | 0 | 11 | 11 | 1 | 1 | CEL |
| 3vb6 | B | 0 | 0 | 0 | 1 | 1 | 0 | 11 | 11 | 0 | 0 | 1 | 21 | 11 | 11 | 1 | 11 | 0 | 0 | 1 | 0 | 1 | 1 | 11 | 0 | 1 | CEL |
| 3vb7 | B | 0 | 0 | 0 | 1 | 1 | 0 | 11 | 1 | 0 | 0 | 1 | 21 | 11 | 11 | 1 | 11 | 0 | 0 | 1 | 1 | 1 | 11 | 1 | 0 | 1 | CEL |
| 4tww | B | 0 | 0 | 0 | 0 | 1 | 0 | 1 | 1 | 1 | 1 | 11 | 11 | 11 | 10 | 0 | 1 | 0 | 0 | 1 | 1 | 1 | 1 | 0 | 0 | 0 | CE |

| | | | | | | | | | | | | | | | | | | | | | | | | | | | | | | | | | | | | | | | | | | | | | | | | | | | | | | |
|---|---|---|---|---|---|---|---|---|---|---|---|---|---|---|---|---|---|---|---|---|---|---|---|---|---|---|---|---|---|---|---|---|---|---|---|---|---|---|---|---|---|---|---|---|---|---|---|---|---|---|---|---|---|---|---|
| | 5c5n | AB | 0 | 0 | 0 | 1 | 1 | 0 | 1 | 1 | 1 | 11 | 0 | 21 | 11 | 1 | 1 | 1 | 0 | 0 | 0 | 1 | 1 | 1 | 0 | 0 | 0 | C | | | | | | | | | | | | | | | | | | | | | | | | | | | |
| | 5c5o | A | 1 | 0 | 0 | 1 | 1 | 0 | 1 | 1 | 1 | 11 | 11 | 21 | 11 | 10 | 0 | 1 | 0 | 0 | 0 | 1 | 1 | 1 | 0 | 0 | 0 | CE | B | 1 | 0 | 0 | 1 | 1 | 0 | 1 | 1 | 1 | 11 | 0 | 21 | 11 | 10 | 0 | 1 | 0 | 0 | 0 | 1 | 0 | 0 | 0 | 0 | 0 | CE |
| | 5n19 | AB | 1 | 0 | 0 | 11 | 1 | 0 | 11 | 0 | 1 | 11 | 1 | 21 | 11 | 11 | 1 | 11 | 0 | 0 | 1 | 1 | 1 | 1 | 0 | 0 | 0 | CEH | | | | | | | | | | | | | | | | | | | | | | | | | | | |
| | 5n5o | AB | 0 | 1 | 0 | 11 | 0 | 0 | 11 | 1 | 1 | 11 | 0 | 21 | 11 | 11 | 1 | 11 | 0 | 0 | 1 | 1 | 0 | 11 | 1 | 0 | 0 | CEL | | | | | | | | | | | | | | | | | | | | | | | | | | | |
| | 6lnq | AB | 0 | 0 | 0 | 1 | 0 | 0 | 11 | 1 | 1 | 0 | 1 | 21 | 11 | 11 | 1 | 11 | 0 | 1 | 1 | 0 | 0 | 0 | 11 | 1 | 1 | CEL | | | | | | | | | | | | | | | | | | | | | | | | | | | |
| | 6lny | AB | 0 | 0 | 0 | 1 | 0 | 0 | 11 | 1 | 1 | 1 | 0 | 21 | 11 | 11 | 1 | 11 | 0 | 0 | 1 | 0 | 0 | 11 | 0 | 0 | 0 | CE | | | | | | | | | | | | | | | | | | | | | | | | | | | |
| | 6lo0 | AB | 0 | 0 | 0 | 1 | 0 | 0 | 11 | 1 | 1 | 0 | 1 | 21 | 11 | 11 | 1 | 11 | 0 | 0 | 1 | 1 | 1 | 11 | 1 | 1 | 0 | CEL | | | | | | | | | | | | | | | | | | | | | | | | | | | |
| | 6w2a | A | 0 | 0 | 0 | 11 | 1 | 0 | 11 | 0 | 1 | 0 | 1 | 21 | 11 | 11 | 1 | 11 | 0 | 0 | 1 | 1 | 1 | 11 | 0 | 0 | 0 | CEHL | B | 0 | 0 | 0 | 1 | 1 | 0 | 1 | 0 | 1 | 1 | 1 | 21 | 11 | 11 | 1 | 11 | 0 | 0 | 1 | 1 | 0 | 11 | 1 | 1 | 0 | CEL |
| | 6w79 | AB | 1 | 1 | 0 | 1 | 0 | 0 | 1 | 1 | 1 | 11 | 11 | 1 | 11 | 1 | 1 | 11 | 0 | 0 | 0 | 1 | 1 | 1 | 0 | 0 | 0 | CE | | | | | | | | | | | | | | | | | | | | | | | | | | | |
| | 6wco | AB | 1 | 1 | 0 | 1 | 1 | 0 | 1 | 1 | 1 | 11 | 1 | 1 | 11 | 1 | 1 | 11 | 0 | 0 | 0 | 1 | 1 | 1 | 0 | 0 | 0 | CE | | | | | | | | | | | | | | | | | | | | | | | | | | | |
| | 6xhl | A | 0 | 0 | 1 | 11 | 0 | 0 | 11 | 1 | 1 | 1 | 1 | 21 | 11 | 11 | 1 | 11 | 0 | 0 | 1 | 1 | 0 | 11 | 1 | 1 | 0 | CEL | B | 0 | 0 | 1 | 11 | 0 | 0 | 1 | 1 | 1 | 1 | 1 | 21 | 11 | 11 | 1 | 11 | 0 | 0 | 1 | 0 | 0 | 11 | 1 | 1 | 0 | CEL |
| | 6xhn | A | 0 | 0 | 0 | 1 | 0 | 0 | 11 | 1 | 1 | 11 | 1 | 21 | 11 | 11 | 1 | 11 | 0 | 0 | 1 | 0 | 0 | 11 | 1 | 1 | 0 | CEL | B | 0 | 0 | 0 | 1 | 1 | 0 | 11 | 1 | 1 | 11 | 1 | 21 | 11 | 11 | 1 | 11 | 0 | 1 | 1 | 0 | 1 | 11 | 1 | 1 | 0 | CEL |
| | 6xho | A | 1 | 1 | 0 | 1 | 0 | 0 | 11 | 1 | 1 | 1 | 0 | 21 | 11 | 11 | 1 | 11 | 0 | 0 | 1 | 1 | 0 | 11 | 1 | 1 | 0 | CEL | B | 1 | 1 | 0 | 1 | 0 | 0 | 11 | 1 | 1 | 1 | 1 | 21 | 11 | 11 | 1 | 11 | 0 | 0 | 1 | 1 | 0 | 11 | 1 | 1 | 0 | CEL |
| | 6y7m | AB | 0 | 1 | 0 | 11 | 1 | 0 | 11 | 1 | 1 | 11 | 0 | 21 | 11 | 10 | 1 | 11 | 1 | 1 | 1 | 1 | 1 | 1 | 0 | 0 | 0 | CEH | | | | | | | | | | | | | | | | | | | | | | | | | | | |
| SARS-CoV-2 | 6lu7 | AB | 1 | 1 | 0 | 1 | 0 | 0 | 11 | 1 | 1 | 11 | 0 | 21 | 11 | 11 | 1 | 11 | 0 | 1 | 1 | 0 | 0 | 11 | 11 | 1 | 1 | CEL | | | | | | | | | | | | | | | | | | | | | | | | | | | |
| | 6lze | AB | 0 | 0 | 0 | 1 | 1 | 1 | 11 | 1 | 1 | 1 | 1 | 21 | 11 | 10 | 1 | 11 | 0 | 0 | 1 | 1 | 1 | 1 | 0 | 0 | 0 | CEH | | | | | | | | | | | | | | | | | | | | | | | | | | | |
| | 6m0k | AB | 0 | 0 | 0 | 1 | 1 | 0 | 11 | 1 | 1 | 1 | 1 | 21 | 11 | 11 | 1 | 11 | 0 | 0 | 1 | 1 | 1 | 0 | 0 | 0 | 0 | CE | | | | | | | | | | | | | | | | | | | | | | | | | | | |
| | 6m2n | A | 0 | 0 | 0 | 1 | 1 | 0 | 0 | 0 | 1 | 11 | 0 | 1 | 0 | 1 | 1 | 10 | 0 | 0 | 0 | 1 | 0 | 1 | 0 | 0 | 0 | | C | 0 | 0 | 0 | 1 | 1 | 0 | 0 | 1 | 1 | 11 | 0 | 1 | 0 | 0 | 1 | 10 | 0 | 0 | 0 | 0 | 0 | 0 | 0 | 0 | 0 | |
| | 6m2n | B | 0 | 0 | 0 | 1 | 1 | 0 | 0 | 1 | 1 | 11 | 0 | 1 | 0 | 1 | 1 | 11 | 0 | 0 | 0 | 1 | 1 | 1 | 0 | 0 | 0 | | D | 0 | 0 | 0 | 1 | 1 | 0 | 0 | 0 | 1 | 11 | 0 | 1 | 0 | 0 | 1 | 10 | 0 | 0 | 0 | 0 | 0 | 0 | 0 | 0 | 0 | |
| | 6w63 | AB | 1 | 1 | 0 | 1 | 0 | 0 | 1 | 1 | 1 | 11 | 11 | 1 | 11 | 1 | 1 | 11 | 0 | 0 | 0 | 1 | 0 | 1 | 0 | 0 | 0 | CE | | | | | | | | | | | | | | | | | | | | | | | | | | | |
| | 6wnp | AB | 0 | 0 | 0 | 1 | 0 | 1 | 0 | 0 | 1 | 11 | 1 | 21 | 0 | 11 | 1 | 11 | 1 | 0 | 0 | 1 | 1 | 1 | 1 | 0 | 1 | CEL | | | | | | | | | | | | | | | | | | | | | | | | | | | |
| | 6wtj | AB | 0 | 0 | 0 | 1 | 1 | 0 | 11 | 0 | 1 | 1 | 0 | 21 | 11 | 11 | 1 | 11 | 0 | 0 | 1 | 0 | 0 | 0 | 0 | 0 | 0 | CE | | | | | | | | | | | | | | | | | | | | | | | | | | | |
| | 6wtk | AB | 0 | 0 | 0 | 1 | 1 | 0 | 11 | 1 | 1 | 11 | 0 | 21 | 11 | 11 | 1 | 11 | 0 | 0 | 1 | 1 | 0 | 11 | 0 | 0 | 0 | CE | | | | | | | | | | | | | | | | | | | | | | | | | | | |
| | 6wtt | AB | 0 | 0 | 0 | 1 | 1 | 0 | 11 | 1 | 1 | 1 | 0 | 21 | 11 | 11 | 1 | 11 | 0 | 1 | 1 | 1 | 0 | 11 | 1 | 1 | 1 | CEL | | | | | | | | | | | | | | | | | | | | | | | | | | | |
| | 6wtt | B | 0 | 0 | 0 | 1 | 0 | 0 | 0 | 1 | 1 | 1 | 1 | 21 | 11 | 11 | 1 | 11 | 0 | 0 | 1 | 1 | 0 | 11 | 1 | 0 | 1 | CEL | C | 0 | 0 | 1 | 11 | 1 | 0 | 11 | 1 | 1 | 0 | 1 | 21 | 11 | 11 | 1 | 11 | 0 | 1 | 1 | 1 | 1 | 11 | 1 | 0 | 1 | CEHL |
| | 6xa4 | AB | 0 | 0 | 0 | 1 | 1 | 0 | 1 | 1 | 1 | 1 | 0 | 21 | 1 | 11 | 1 | 11 | 0 | 0 | 0 | 0 | 0 | 1 | 1 | 0 | 0 | CE | | | | | | | | | | | | | | | | | | | | | | | | | | | |
| | 6xb1 | AB | 0 | 0 | 1 | 1 | 0 | 0 | 0 | 0 | 0 | 1 | 1 | 11 | 1 | 1 | 0 | 0 | 0 | 0 | 0 | 0 | 0 | 0 | 0 | 0 | 0 | | | | | | | | | | | | | | | | | | | | | | | | | | | | |
| | 6xb2 | AB | 0 | 0 | 0 | 1 | 0 | 0 | 0 | 0 | 0 | 1 | 0 | 21 | 0 | 0 | 0 | 0 | 0 | 0 | 0 | 0 | 0 | 0 | 0 | 0 | 0 | | | | | | | | | | | | | | | | | | | | | | | | | | | | |
| | 6xbg | A | 0 | 1 | 0 | 11 | 1 | 0 | 11 | 1 | 1 | 11 | 0 | 21 | 11 | 11 | 1 | 11 | 0 | 1 | 1 | 1 | 0 | 11 | 1 | 1 | 0 | CEHL | B | 0 | 1 | 1 | 11 | 1 | 0 | 1 | 1 | 1 | 11 | 0 | 21 | 11 | 11 | 1 | 11 | 0 | 0 | 1 | 0 | 0 | 1 | 0 | 0 | 0 | CEH |
| | 6xbh | AB | 0 | 0 | 0 | 1 | 1 | 1 | 11 | 0 | 1 | 1 | 0 | 21 | 11 | 11 | 1 | 11 | 0 | 0 | 1 | 1 | 1 | 11 | 0 | 0 | 0 | CEHL | | | | | | | | | | | | | | | | | | | | | | | | | | | |
| | 6xbi | A | 1 | 1 | 0 | 11 | 0 | 0 | 1 | 0 | 1 | 11 | 0 | 21 | 11 | 11 | 1 | 11 | 0 | 1 | 1 | 1 | 0 | 11 | 1 | 1 | 1 | CEL | B | 1 | 1 | 0 | 11 | 0 | 0 | 1 | 0 | 1 | 11 | 1 | 21 | 11 | 11 | 1 | 11 | 0 | 1 | 1 | 1 | 0 | 11 | 1 | 1 | 1 | CEL |
| | 6xch | AB | 0 | 0 | 0 | 1 | 1 | 1 | 0 | 0 | 11 | 1 | 0 | 21 | 0 | 11 | 1 | 11 | 0 | 0 | 0 | 1 | 0 | 1 | 1 | 0 | 0 | CEH | | | | | | | | | | | | | | | | | | | | | | | | | | | |
| | 6xfn | AB | 0 | 1 | 0 | 11 | 0 | 0 | 1 | 1 | 1 | 11 | 11 | 21 | 11 | 11 | 0 | 1 | 0 | 0 | 0 | 0 | 0 | 1 | 0 | 0 | 0 | CE | | | | | | | | | | | | | | | | | | | | | | | | | | | |
| | 6xhm | A | 0 | 0 | 1 | 1 | 0 | 0 | 11 | 1 | 1 | 0 | 1 | 21 | 11 | 11 | 1 | 11 | 0 | 1 | 1 | 1 | 0 | 11 | 1 | 1 | 0 | CEL | B | 0 | 0 | 1 | 11 | 0 | 0 | 11 | 1 | 1 | 0 | 1 | 21 | 11 | 11 | 1 | 11 | 0 | 0 | 1 | 0 | 1 | 11 | 1 | 1 | 0 | CEL |
| | 6xmk | A | 0 | 0 | 0 | 1 | 0 | 0 | 11 | 1 | 1 | 1 | 0 | 21 | 11 | 11 | 1 | 11 | 0 | 0 | 1 | 1 | 0 | 11 | 1 | 1 | 0 | CEL | B | 0 | 0 | 0 | 1 | 0 | 0 | 11 | 1 | 1 | 1 | 0 | 21 | 11 | 11 | 1 | 11 | 0 | 1 | 1 | 1 | 1 | 11 | 1 | 0 | 0 | CEL |
| | 6xoa | A | 0 | 0 | 0 | 1 | 1 | 0 | 11 | 1 | 1 | 11 | 11 | 11 | 1 | 0 | 1 | 11 | 0 | 1 | 1 | 1 | 1 | 11 | 0 | 1 | 11 | CEL | C | 0 | 0 | 0 | 1 | 1 | 0 | 1 | 1 | 0 | 11 | 11 | 11 | 11 | 0 | 1 | 11 | 0 | 1 | 1 | 1 | 1 | 11 | 1 | 0 | 1 | CEL |
| | 6xqs | AB | 0 | 1 | 0 | 11 | 1 | 0 | 0 | 0 | 1 | 11 | 0 | 21 | 0 | 11 | 1 | 11 | 1 | 1 | 0 | 1 | 1 | 11 | 1 | 1 | 1 | CEHL | | | | | | | | | | | | | | | | | | | | | | | | | | | |
| | 6xqt | A | 1 | 1 | 0 | 11 | 1 | 1 | 0 | 1 | 1 | 11 | 0 | 21 | 0 | 11 | 1 | 11 | 1 | 1 | 0 | 1 | 1 | 1 | 1 | 1 | 1 | CEHL | B | 0 | 1 | 0 | 11 | 1 | 1 | 0 | 1 | 1 | 11 | 0 | 21 | 1 | 11 | 1 | 11 | 1 | 1 | 0 | 1 | 1 | 1 | 1 | 1 | 1 | CEHL |
| | 6xqu | AB | 0 | 0 | 0 | 11 | 1 | 0 | 0 | 1 | 1 | 11 | 0 | 21 | 1 | 11 | 1 | 11 | 1 | 1 | 0 | 1 | 1 | 1 | 1 | 0 | 1 | CEHL | | | | | | | | | | | | | | | | | | | | | | | | | | | |
| | 6xr3 | AB | 1 | 0 | 0 | 1 | 0 | 0 | 11 | 0 | 1 | 1 | 0 | 21 | 11 | 11 | 1 | 11 | 0 | 1 | 1 | 0 | 0 | 11 | 1 | 1 | 0 | CEL | | | | | | | | | | | | | | | | | | | | | | | | | | | |
| | 6y2f | AB | 0 | 1 | 0 | 11 | 1 | 0 | 11 | 1 | 1 | 11 | 0 | 21 | 11 | 11 | 1 | 11 | 1 | 1 | 1 | 1 | 0 | 1 | 0 | 0 | 0 | CEH | | | | | | | | | | | | | | | | | | | | | | | | | | | |
| | 6y2g | A | 0 | 1 | 0 | 11 | 1 | 0 | 11 | 1 | 1 | 1 | 1 | 21 | 11 | 11 | 1 | 11 | 0 | 0 | 1 | 1 | 0 | 1 | 0 | 0 | 0 | CEH | B | 0 | 1 | 0 | 11 | 1 | 0 | 11 | 1 | 1 | 11 | 1 | 21 | 11 | 11 | 1 | 11 | 1 | 1 | 0 | 1 | 1 | 0 | 0 | 0 | 0 | CEH |
| | 6ynq | AB | 0 | 0 | 0 | 0 | 0 | 0 | 1 | 1 | 1 | 0 | 0 | 21 | 11 | 1 | 0 | 1 | 0 | 0 | 0 | 0 | 0 | 0 | 0 | 0 | 0 | | | | | | | | | | | | | | | | | | | | | | | | | | | | |
| | 6yvf | AB | | | | | | | | | | | | | | | | | | | | | | | | | | | | | | | | | | | | | | | | | | | | | | | | | | | | | |
| | 6z2e | AB | 0 | 1 | 1 | 1 | 0 | 0 | 11 | 0 | 1 | 11 | 1 | 21 | 11 | 11 | 1 | 11 | 0 | 0 | 1 | 0 | 1 | 11 | 11 | 1 | 1 | CEL | | | | | | | | | | | | | | | | | | | | | | | | | | | |
| | 6zrt | AB | 0 | 1 | 0 | 11 | 0 | 0 | 0 | 0 | 1 | 11 | 0 | 21 | 1 | 11 | 1 | 11 | 1 | 1 | 0 | 0 | 1 | 11 | 1 | 1 | 1 | CEL | | | | | | | | | | | | | | | | | | | | | | | | | | | |
| | 6zru | AB | 0 | 0 | 0 | 11 | 1 | 1 | 0 | 0 | 0 | 11 | 0 | 21 | 0 | 11 | 1 | 11 | 1 | 1 | 0 | 1 | 1 | 1 | 1 | 0 | 1 | EHL | | | | | | | | | | | | | | | | | | | | | | | | | | | |
| | 7aku | AB | 0 | 0 | 0 | 1 | 0 | 0 | 1 | 1 | 1 | 1 | 0 | 21 | 1 | 11 | 1 | 11 | 0 | 0 | 0 | 1 | 0 | 11 | 0 | 0 | 0 | CE | | | | | | | | | | | | | | | | | | | | | | | | | | | |
| | 7b83 | AB | 1 | 1 | 1 | 21 | 0 | 0 | 0 | 0 | 1 | 0 | 0 | 21 | 0 | 0 | 0 | 0 | 0 | 0 | 0 | 0 | 0 | 0 | 0 | 0 | 0 | R | | | | | | | | | | | | | | | | | | | | | | | | | | | |
| | 7bqy | AB | 0 | 1 | 1 | 1 | 0 | 0 | 11 | 1 | 1 | 11 | 0 | 21 | 11 | 11 | 1 | 11 | 0 | 1 | 1 | 0 | 1 | 11 | 11 | 0 | 1 | CEL | | | | | | | | | | | | | | | | | | | | | | | | | | | |
| | 7brp | A | 0 | 0 | 0 | 11 | 1 | 0 | 0 | 0 | 11 | 11 | 0 | 21 | 0 | 11 | 1 | 11 | 0 | 0 | 0 | 1 | 0 | 1 | 1 | 0 | 1 | CEHL | B | 0 | 0 | 0 | 11 | 1 | 0 | 0 | 0 | 11 | 11 | 0 | 21 | 0 | 11 | 1 | 11 | 0 | 0 | 0 | 1 | 1 | 1 | 1 | 0 | 1 | CEHL |
| | 7buy | AB | 0 | 0 | 0 | 1 | 1 | 0 | 0 | 0 | 0 | 11 | 0 | 21 | 0 | 1 | 1 | 0 | 0 | 0 | 0 | 1 | 0 | 0 | 0 | 0 | 0 | | | | | | | | | | | | | | | | | | | | | | | | | | | | |
| | 7c6s | AB | 0 | 0 | 0 | 11 | 1 | 1 | 0 | 0 | 1 | 11 | 0 | 21 | 0 | 11 | 1 | 11 | 0 | 0 | 0 | 1 | 1 | 1 | 1 | 0 | 1 | CEHL | | | | | | | | | | | | | | | | | | | | | | | | | | | |
| | 7c6u | AB | 0 | 0 | 0 | 1 | 1 | 1 | 11 | 1 | 1 | 1 | 1 | 21 | 11 | 11 | 1 | 11 | 0 | 0 | 1 | 1 | 0 | 1 | 0 | 0 | 0 | CEH | | | | | | | | | | | | | | | | | | | | | | | | | | | |
| | 7c7p | A | 0 | 1 | 0 | 11 | 1 | 0 | 0 | 0 | 1 | 11 | 0 | 21 | 0 | 11 | 1 | 11 | 1 | 1 | 0 | 1 | 1 | 11 | 1 | 0 | 1 | CEHL | B | 1 | 1 | 0 | 11 | 0 | 0 | 0 | 0 | 0 | 11 | 1 | 21 | 0 | 1 | 0 | 0 | 0 | 0 | 0 | 1 | 1 | 1 | 0 | 0 | 0 | C |
| | 7c8b | AB | 0 | 0 | 0 | 1 | 1 | 1 | 1 | 0 | 1 | 11 | 1 | 21 | 11 | 0 | 1 | 11 | 0 | 0 | 1 | 1 | 1 | 1 | 0 | 0 | 0 | CEH | | | | | | | | | | | | | | | | | | | | | | | | | | | |
| | 7c8r | AB | 1 | 1 | 1 | 11 | 0 | 0 | 11 | 1 | 1 | 1 | 1 | 21 | 11 | 11 | 1 | 11 | 1 | 1 | 1 | 0 | 1 | 11 | 1 | 1 | 1 | RCEL | | | | | | | | | | | | | | | | | | | | | | | | | | | |
| | 7c8t | AB | 0 | 0 | 0 | 1 | 1 | 1 | 11 | 1 | 1 | 11 | 1 | 21 | 11 | 10 | 1 | 11 | 1 | 1 | 1 | 1 | 1 | 1 | 1 | 1 | 1 | CEHL | | | | | | | | | | | | | | | | | | | | | | | | | | | |

| PDBid | BC | T25 | T26 | L27 | H41 | M49 | Y54 | F140 | L141 | N142 | G143 | S144 | C145 | H163 | H164 | M165 | E166 | L167 | P168 | H172 | D187 | R188 | Q189 | T190 | A191 | Q192 | Ligand interactions |
|---|---|---|---|---|---|---|---|---|---|---|---|---|---|---|---|---|---|---|---|---|---|---|---|---|---|---|---|
| | MC | interface | | | H-loop | | | C-loop | | | | interface | | I&II | | E-loop | | | | | I&II | Linker | | | | | |
| 7c8u | AB | 0 | 0 | 0 | 1 | 1 | 0 | 11 | 0 | 1 | 11 | 0 | 21 | 11 | 11 | 1 | 11 | 0 | 0 | 1 | 1 | 0 | 0 | 0 | 0 | 0 | CE |
| 7cbt | A | 0 | 0 | 0 | 1 | 0 | 0 | 11 | 1 | 1 | 0 | 1 | 21 | 11 | 11 | 1 | 11 | 0 | 0 | 1 | 1 | 1 | 1 | 0 | 0 | 0 | CE |
| 7com | A | 0 | 0 | 1 | 11 | 1 | 0 | 0 | 0 | 1 | 11 | 0 | 21 | 0 | 11 | 1 | 11 | 1 | 1 | 0 | 1 | 1 | 11 | 1 | 0 | 1 | CEHL |
| 7cx9 | AB | 0 | 0 | 0 | 1 | 0 | 0 | 0 | 0 | 0 | 0 | 0 | 21 | 0 | 1 | 1 | 0 | 0 | 0 | 0 | 0 | 0 | 0 | 0 | 0 | 0 | |
| 7d1m | A | 0 | 0 | 0 | 11 | 1 | 0 | 11 | 0 | 1 | 1 | 0 | 21 | 11 | 11 | 1 | 11 | 0 | 1 | 1 | 1 | 0 | 11 | 1 | 1 | 0 | CEHL |
| 7d1o | AB | 1 | 1 | 0 | 11 | 1 | 1 | 1 | 1 | 1 | 11 | 0 | 21 | 0 | 11 | 1 | 11 | 1 | 1 | 0 | 1 | 1 | 1 | 1 | 1 | 1 | CEHL |
| 7d3i | AB | 0 | 0 | 0 | 1 | 0 | 0 | 11 | 0 | 1 | 1 | 0 | 21 | 11 | 11 | 1 | 11 | 0 | 0 | 1 | 1 | 1 | 1 | 0 | 0 | 0 | CE |
| 7jkv | A | 1 | 0 | 0 | 1 | 1 | 0 | 11 | 1 | 1 | 1 | 1 | 21 | 11 | 11 | 1 | 11 | 0 | 1 | 1 | 1 | 0 | 11 | 1 | 1 | 0 | CEL |
| 7joy | | | | | | | | | | | | | | | | | | | | | | | | | | | |
| 7ju7 | AB | 1 | 0 | 0 | 1 | 1 | 0 | 1 | 1 | 1 | 0 | 0 | 10 | 11 | 11 | 1 | 1 | 0 | 0 | 0 | 0 | 0 | 0 | 0 | 0 | 0 | E |
| 7jyc | AB | 0 | 1 | 0 | 11 | 1 | 1 | 1 | 1 | 1 | 11 | 0 | 21 | 0 | 11 | 1 | 11 | 1 | 1 | 0 | 1 | 1 | 1 | 1 | 1 | 1 | CEHL |
| 7k40 | AB | 0 | 0 | 0 | 11 | 1 | 1 | 0 | 0 | 1 | 11 | 1 | 21 | 0 | 11 | 1 | 11 | 1 | 1 | 0 | 1 | 1 | 1 | 1 | 0 | 1 | CEHL |
| 7k6d | AB | 0 | 1 | 0 | 11 | 1 | 0 | 0 | 0 | 1 | 11 | 0 | 21 | 0 | 11 | 1 | 11 | 1 | 1 | 0 | 0 | 1 | 11 | 1 | 1 | 1 | CEHL |
| 7k6e | AB | 0 | 1 | 0 | 11 | 0 | 0 | 0 | 0 | 1 | 11 | 0 | 21 | 0 | 11 | 1 | 11 | 1 | 1 | 0 | 0 | 1 | 11 | 1 | 1 | 1 | CEL |
| 7khp | | | | | | | | | | | | | | | | | | | | | | | | | | | |
| 7lfp | A | | | | | | | | | | | | | | | | | | | | | | | | | | |
| 7nev | AB | 0 | 0 | 0 | 1 | 0 | 1 | 0 | 0 | 1 | 1 | 0 | 21 | 0 | 11 | 1 | 11 | 0 | 0 | 0 | 1 | 1 | 1 | 1 | 0 | 0 | EL |

| PDBid | BC | T25 | T26 | L27 | H41 | M49 | Y54 | F140 | L141 | N142 | G143 | S144 | C145 | H163 | H164 | M165 | E166 | L167 | P168 | H172 | D187 | R188 | Q189 | T190 | A191 | Q192 | Ligand interactions |
|---|---|---|---|---|---|---|---|---|---|---|---|---|---|---|---|---|---|---|---|---|---|---|---|---|---|---|---|
| | MC | interface | | | H-loop | | | C-loop | | | | interface | | I&II | | E-loop | | | | | I&II | Linker | | | | | |
| 7c8u | | | | | | | | | | | | | | | | | | | | | | | | | | | |
| 7cbt | B | 0 | 0 | 0 | 1 | 0 | 0 | 11 | 1 | 1 | 0 | 0 | 21 | 11 | 11 | 1 | 11 | 1 | 0 | 1 | 0 | 0 | 11 | 0 | 0 | 0 | CE |
| 7com | B | 0 | 0 | 0 | 11 | 0 | 0 | 0 | 0 | 1 | 11 | 0 | 21 | 0 | 11 | 1 | 11 | 0 | 1 | 0 | 1 | 1 | 1 | 1 | 0 | 1 | CEL |
| 7cx9 | | | | | | | | | | | | | | | | | | | | | | | | | | | |
| 7d1m | B | 0 | 0 | 0 | 11 | 0 | 0 | 11 | 0 | 1 | 1 | 0 | 21 | 11 | 11 | 1 | 11 | 0 | 0 | 1 | 1 | 0 | 11 | 0 | 0 | 0 | CE |
| 7d1o | | | | | | | | | | | | | | | | | | | | | | | | | | | |
| 7d3i | | | | | | | | | | | | | | | | | | | | | | | | | | | |
| 7jkv | B | 1 | 0 | 1 | 1 | 1 | 0 | 1 | 0 | 1 | 1 | 1 | 21 | 11 | 11 | 1 | 11 | 0 | 0 | 1 | 0 | 0 | 11 | 1 | 1 | 0 | CEL |
| 7joy | B | 0 | 0 | 0 | 11 | 1 | 0 | 11 | 1 | 0 | 11 | 1 | 11 | 11 | 11 | 1 | 11 | 1 | 1 | 1 | 1 | 1 | 11 | 0 | 1 | 11 | CEHL |
| 7ju7 | | | | | | | | | | | | | | | | | | | | | | | | | | | |
| 7jyc | | | | | | | | | | | | | | | | | | | | | | | | | | | |
| 7k40 | | | | | | | | | | | | | | | | | | | | | | | | | | | |
| 7k6d | | | | | | | | | | | | | | | | | | | | | | | | | | | |
| 7k6e | | | | | | | | | | | | | | | | | | | | | | | | | | | |
| 7khp | B | 0 | 0 | 0 | 0 | 1 | 0 | 11 | 1 | 0 | 11 | 1 | 20 | 11 | 1 | 1 | 1 | 1 | 1 | 1 | 1 | 1 | 11 | 0 | 1 | 1 | CEL |
| 7lfp | | | | | | | | | | | | | | | | | | | | | | | | | | | |
| 7nev | | | | | | | | | | | | | | | | | | | | | | | | | | | |

The assignments of contacts are as follows: 1 for nonpolar contact, 10 for polar contact, 11 for those containing both polar cond nonpolar contacts, 20 for covalent bond, and 21 for those containing both covalent and nonpolar contacts. Coordination between Metal ion and S or O is classified as covalent here. Polar and nonpolar contacts were assigned by LigPlot (35). Covalent bond is defined according to PyMol. MC and BC are moving cluster and binding cluster, respectively. The definition of binding mode is as follows: C: (H >= 2 and H+N >= 3) or (H+N >= 5) ; E: (H >= 2 and H+N >= 3) or (H+N >= 5) ; L: (H >= 1 and H+N >= 3) or (H+N>= 4) ; H: (H >= 1 and H+N >= 2) or (H+N>= 3); Core: (H >= 1 and H+N >= 2) or (H+N >= 3). Here H and N dinote the numbers of polar and nonpolar contacts, respectively. The enttries, 6yvf and 7lfp, bind ligands at portions of the protein other than the major binding sites, and no imformatiton is given in the table; instead the binding sites are listted here: 6yvf A:63,78,79,80,81,88,90; 7lfp A:4,5,7,125,126,127, B:5,7,127 (residue numbers without underline are for nonpolar contacts, and with underline for polar contact). The minor 16 binding sites are Thr21 (1), Thr24 (6), Asn28 (1), Pro39 (1), Cys44 (10), Thr45 (5), Ala46 (7), Asn51 (1), Tyr67 (1), Lys102 (1), Gly146 (1), Ser147 (1), Cys156 (2), Phe185 (2), Val186 (10), Ala193 (1), where the numbers of the parentheses are the number of occurrence of the contacts. The liagnd interaction in 6xoa was chosen for the interaction between chain A (donor) and chain C (acceptor), and the interaction between chain A (acceptor) and chain C (donor) was not calculated here.

**Dataset S4. Summary of the ligand-bound entries supplemented to Dataset S1.**

| species | protein | | | | ligand | | | | comments | | | | | | | | | | | | | | | | | | | | | | | | | | | | | dimer | | |
|---|---|---|---|---|---|---|---|---|---|---|---|---|---|---|---|---|---|---|---|---|---|---|---|---|---|---|---|---|---|---|---|---|---|---|---|---|---|---|---|---|
| | | | | | | | | | | PC1 | | | | | C-loop conformation | | | | | | | | | PC1 | | | | | C-loop conformation | | | | | | | | | | | |
| species | PDB ID | mutation | residues added to N-term | missing N-terminal residues# | name in PDB | name by authors | peptide or mimic@ | Ki, $IC_{50}$*, $EC_{50}$+ (μM) | comments | C-loop | E-loop | H-loop | Linker | Domain III | main-chain HB within 138-144 | HB 138-172& | HB 139-126& | HB 140-1& | HB 141-118& | HB 143-28& | distance 165Cα-140Cβ | distance 165Cα-141Cβ | distance 2-214 | C-loop | E-loop | H-loop | Linker | Domain III | main-chain HB within 138-144 | HB 138-172& | HB 139-126& | HB 140-1& | HB 141-118& | HB 143-28& | distance 165Cα-140Cβ | distance 165Cα-141Cβ | distance 2-214 | PC1 Domain III | PC2 Domain III | distance between two 285 Cα's |
| SARS-CoV | 6w79 | | | | X77 | X77 | | | | 0.571 | 0.571 | -1.274 | 1.296 | 6.618 | 141O-144N | 1 | 1 | 1 | 1 | 1 | 7.801 | 11.379 | 7.671 | | | | | | | | | | | | | | | 0.001 | 6.13 | 8.255 |
| | 7k0g | | MH6 | MHHHH(-2) of A,B | B1S/K36 | 2 | P | | B1S and K36 are stereoisomers, which are not distinguished. | 0.669 | 0.055 | -1.17 | -2.642 | -11.425 | 141O-144N | 1 | 1 | 0 | 1 | 1 | 7.309 | 11.083 | 8.611 | | | | | | | | | | | | | | | 0.001 | -2.789 | 5.257 |
| | 7k0h | | MH6 | MH6S̲ of A,B | VR4 | 5 | P | | | 0.482 | 0.914 | 0.238 | -2.613 | -5.659 | 141O-144N | 1 | 1 | 0 | 1 | 1 | 7.544 | 11.364 | 7.017 | 0.452 | 0.302 | -0.308 | -1.268 | 36.635 | 141O-144N | 1 | 1 | 0 | 1 | 1 | 7.437 | 11.284 | 10.048 | -29.283 | 19.527 | 8.908 |
| | 7lcp | | | | UED | GC373 | P | | | 0.738 | 0.546 | -0.203 | 0.41 | 3.343 | 141O-144N | 1 | 1 | 0 | 1 | 1 | 7.463 | 11.246 | 7.255 | | | | | | | | | | | | | | | 0.001 | 9.125 | 7.831 |
| | 7lcq | | | | UED | GC373 | P | | | 0.581 | 0.485 | -0.493 | 0.686 | 2.609 | 141O-144N | 1 | 1 | 0 | 1 | 1 | 7.452 | 11.350 | 7.084 | | | | | | | | | | | | | | | 0.002 | 8.19 | 7.844 |
| SARS-CoV-2 | 7a1u | | | | FUA | Fusidic acid | | | | 0.591 | -0.627 | -1.131 | -0.49 | -4.356 | 141O-144N | 1 | 1 | 1 | 1 | 1 | 7.577 | 11.358 | 6.111 | | | | | | | | | | | | | | | -0.001 | -10.207 | 5.014 |
| | 7abu | | | | R6Q | RS102895 | | 19.8+ 16.73+ | R6Q is between the surface of domain III and C-loop of another protomer | 1.237 | -1.33 | -1.399 | -0.274 | -6.221 | 141O-144N | 1 | 1 | 1 | 1 | 1 | 7.713 | 11.548 | 6.256 | | | | | | | | | | | | | | | 0.002 | -10.44 | 5.341 |
| | 7adw | | | | R7Q | Dimethyl propio phenone | | | | 1.109 | -0.625 | -1.718 | -0.25 | -6.739 | 141O-144N | 1 | 1 | 1 | 1 | 1 | 7.766 | 11.511 | 6.047 | | | | | | | | | | | | | | | -0.001 | -11.421 | 5.133 |
| | 7aeg | | | | F2G | 5 | P | | Q306 do not exist. | 0.955 | -0.188 | -1.467 | 0.434 | -9.039 | 141O-144N | 1 | 1 | 1 | 1 | 1 | 7.629 | 11.552 | 6.291 | | | | | | | | | | | | | | | -0.001 | -13.825 | 5.250 |
| | 7aeh | | | | R8H | 1 | P | 3* | Q306 do not exist. | 0.418 | -0.594 | -1.089 | -1.314 | -1.344 | 141O-144N | 1 | 1 | 1 | 1 | 1 | 8.140 | 11.656 | 6.561 | | | | | | | | | | | | | | | -0.001 | -6.262 | 5.139 |
| | 7af0 | | | | R9W | Ipidacrine | | | R9W is a small compound located on the surface of domain III. | 1.061 | -0.879 | -1.829 | 0.017 | -8.358 | 141O-144N | 1 | 1 | 1 | 1 | 1 | 7.823 | 11.672 | 6.235 | | | | | | | | | | | | | | | 0 | -11.839 | 5.140 |
| | 7aga | | | | LZE | AT7519 | | 25.16+ 43.95+ | LZE is at the interface of domain II and III. | 0.969 | -1.09 | -1.071 | -0.484 | -5.861 | 141O-144N | 1 | 1 | 1 | 1 | 1 | 7.544 | 11.335 | 6.146 | | | | | | | | | | | | | | | -0.001 | -10.886 | 5.358 |
| | 7aha | | | | SIN | Maleate | | | | 1.063 | -1.29 | -1.728 | -0.321 | -8.223 | 141O-144N | 1 | 1 | 1 | 1 | 1 | 7.673 | 11.530 | 5.956 | | | | | | | | | | | | | | | 0.001 | -11.482 | 5.239 |
| | 7ak4 | | | | CB1 | Tretazicar | | | | 1.04 | -0.522 | -1.629 | 0.031 | -5.522 | 141O-144N | 1 | 1 | 1 | 1 | 1 | 7.332 | 11.049 | 6.125 | | | | | | | | | | | | | | | -0.001 | -11.268 | 5.125 |
| | 7amj | | | | RMZ | PD 168568 | | | RMZ is between the surface of domain III and C-loop of another protomer. | 1.162 | -1.085 | -1.617 | -0.246 | -7.536 | 141O-144N | 1 | 1 | 1 | 1 | 1 | 7.631 | 11.463 | 6.034 | | | | | | | | | | | | | | | 0 | -11.772 | 5.151 |
| | 7ans | | | | RNW | Adrafinil | | | | 0.802 | -0.585 | -1.548 | -0.19 | -5.445 | 141O-144N | 1 | 1 | 1 | 1 | 1 | 7.779 | 11.528 | 6.312 | | | | | | | | | | | | | | | 0 | -8.487 | 5.130 |
| | 7aol | N72L | | | RQH | Climbazole | | | RQH is a small compound located on the molecular surface. | 1.044 | -1.006 | -1.725 | 0.004 | -7.533 | 141O-144N | 1 | 1 | 1 | 1 | 1 | 7.730 | 11.627 | 6.063 | | | | | | | | | | | | | | | 0 | -11.516 | 5.186 |
| | 7ap6 | | | | RQN | MUT056399 | | 38.24+ 36.73+ | M49(Alt B) and M65 clashes with RQH. | 0.848 | -0.488 | -2.251 | -0.06 | -5.364 | 141O-144N | 1 | 1 | 1 | 1 | 1 | 7.701 | 11.436 | 6.341 | | | | | | | | | | | | | | | -0.002 | -8.891 | 5.163 |
| | 7aph | | | | RT2 | Tofogliflozin | | | RT2 is on the surface of domain III. | 0.858 | -0.492 | -1.66 | -0.163 | -6.477 | 141O-144N | 1 | 1 | 1 | 1 | 1 | 7.430 | 11.643 | 5.859 | | | | | | | | | | | | | | | 0 | -11.15 | 5.275 |
| | 7aqe | | | | RV5 | UNC-2327 | | | | 0.76 | -1.062 | -1.56 | -0.4 | -6.909 | 141O-144N | 1 | 1 | 1 | 1 | 1 | 7.807 | 11.574 | 6.499 | | | | | | | | | | | | | | | 0.002 | -10.237 | 5.080 |
| | 7aqi | | | | QEL | Ifenprodil | | | QEL is on the surface of domain III. | 1.141 | -0.966 | -1.791 | -0.09 | -7.78 | 141O-144N | 1 | 1 | 1 | 1 | 1 | 7.826 | 11.614 | 6.055 | | | | | | | | | | | | | | | 0.001 | -10.521 | 5.117 |
| | 7aqj | | | | S7H,RV8 | Triglycidyl isocyanurate | | 30.02+ 31.51+ | S7H and RV8 are stereoisomers. The epoxide group of RV8 is elimiated to make alcohol of S7H. | 0.571 | -1.096 | -1.731 | -0.155 | -6.182 | 141O-144N | 1 | 1 | 1 | 1 | 1 | 7.325 | 11.269 | 6.302 | | | | | | | | | | | | | | | 0.001 | -7.911 | 5.411 |
| | 7arf | | | | RVW | thioglucose | | | RVW is a small compound located on the molecular surface. C145 is di-oxydized to be CSD (3-sulfinoalanine). | 1.082 | -1.854 | -1.093 | -0.906 | -5.948 | 141O-144N | 1 | 1 | 1 | 1 | 1 | 7.778 | 11.647 | 6.374 | | | | | | | | | | | | | | | 0 | -9.718 | 5.365 |
| | 7avd_B | | | | S1W | SEN1269 | | | Chain A is ligand-unbound. | | | | | | | | | | | | | | | 0.928 | -1.453 | -2.084 | -4.254 | -3.084 | 141O-144N | 1 | 1 | 1 | 1 | 1 | 7.210 | 11.149 | 6.641 | -6.19 | -9.655 | 5.526 |
| | 7awr | | | | S7W | Tegafur | | | S7W is a small compound located on the molecular surface. | 0.984 | -0.964 | -1.436 | -0.363 | -6.332 | 141O-144N | 1 | 1 | 1 | 1 | 1 | 7.778 | 11.616 | 6.413 | | | | | | | | | | | | | | | 0 | -11.495 | 5.142 |
| | 7aws | | | | S8E | TH-302 | | | | 1.249 | -0.99 | -1.743 | -0.032 | -7.853 | 141O-144N | 1 | 1 | 1 | 1 | 1 | 7.812 | 11.594 | 6.121 | | | | | | | | | | | | | | | 0.001 | -11.281 | 5.338 |
| | 7awu | | | | S8B | LSN2463359 | | | Cys145 is mono-oxydized to be CSO (hydroxycysteine). N142(Alt A) and M165(Alt A) clashes with S8B. | 0.959 | -0.781 | -1.683 | -0.112 | -5.146 | 141O-144N | 1 | 1 | 1 | 1 | 1 | 7.740 | 11.475 | 6.137 | | | | | | | | | | | | | | | -0.001 | -8.529 | 5.272 |
| | 7aww | | | | CLU | Clonidine | | | CLU is a small compound located on the molecular surface. | 1.091 | -1.031 | -1.812 | -0.129 | -6.623 | 141O-144N | 1 | 1 | 1 | 1 | 1 | 7.799 | 11.737 | 6.444 | | | | | | | | | | | | | | | 0 | -10.695 | 5.113 |
| | 7ax6 | | | | S8H | Glutathione isopropyl ester | P | | S8H is a small compound located on the molecular surface. | 1.124 | -0.824 | -2.058 | 0.206 | -5.323 | 141O-144N | 1 | 1 | 1 | 1 | 1 | 7.604 | 11.278 | 6.271 | | | | | | | | | | | | | | | 0 | -7.787 | 5.369 |
| | 7axm | | | | 93J | Pelitinib | | 1.25+ 1.24+ | 93J is between the surface of domain III and C-loop of another protomer. | 0.908 | -0.638 | -1.607 | -0.182 | -7.143 | 141O-144N | 1 | 1 | 1 | 1 | 1 | 7.781 | 11.551 | 6.310 | | | | | | | | | | | | | | | -0.001 | -10.848 | 5.186 |
| | 7axo | | | | QCP | AR-42 | | | QCP is on the surface of domain III. | 1.067 | -1.034 | -1.649 | -0.081 | -8.229 | 141O-144N | 1 | 1 | 1 | 1 | 1 | 7.730 | 11.583 | 6.279 | | | | | | | | | | | | | | | 0 | -11.13 | 5.305 |
| | 7ay7 | | | | S8T | Isofloxythepin | | 4.8+ 4.92+ | S8T is double configurations. | 1.035 | -1.305 | -1.69 | -0.514 | -7.277 | 141O-144N | 1 | 1 | 1 | 1 | 1 | 7.733 | 11.598 | 6.065 | | | | | | | | | | | | | | | -0.001 | -10.844 | 5.237 |
| | 7b3e | | | | MYC | myricetin | | | | 0.242 | 0.123 | -0.022 | 0.768 | 4.312 | 141O-144N | 1 | 1 | 1 | 1 | 1 | 7.830 | 11.660 | 7.348 | 0.49 | 0.07 | 0.336 | 1.034 | -1.446 | 141O-144N | 1 | 1 | 1 | 1 | 1 | 7.735 | 11.620 | 6.953 | 0.553 | 2.512 | 5.801 |
| | 7bak | | | | Se | ebselen | | 0.670* 4.67+ | Only Se is modeled. | 0.917 | -2.287 | -1.024 | -2.079 | -4.729 | 141O-144N | 1 | 1 | 1 | 1 | 1 | 7.456 | 11.651 | 6.271 | | | | | | | | | | | | | | | -0.001 | -8.554 | 5.263 |
| | 7bal | | | | Se | MR6-31-2 | | 0.824* 1.78+ | Only Se is modeled. | 0.614 | -2.121 | -1.137 | -2.421 | -4.218 | 141O-144N | 1 | 1 | 1 | 1 | 1 | 7.658 | 11.922 | 6.163 | | | | | | | | | | | | | | | 0.001 | -8.751 | 5.431 |
| | 7be7 | | | | ALD | MG-132 | P | | | 0.809 | 0.816 | 0.264 | 1.484 | 0.619 | 141O-144N | 1 | 1 | 1 | 1 | 1 | 7.765 | 11.495 | 6.668 | 1.017 | 0.677 | 0.418 | 1.447 | -3.208 | 141O-144N | 1 | 1 | 1 | 1 | 1 | 7.643 | 11.526 | 6.667 | -3.594 | -2.657 | 5.429 |
| | 7bfb | | | | 9JT | ebselen | | | There are five 9JT's (3 in chain A and 2 in chain B). Only a fragments of 9JT is modeled. Se-C145 is only in chain A. | 0.551 | -0.644 | -0.781 | 0.9 | 5.914 | 141O-144N | 1 | 1 | 1 | 1 | 1 | 7.642 | 11.455 | 7.639 | 0.622 | -0.188 | 0.424 | 1.601 | -6.68 | 141O-144N | 1 | 1 | 1 | 1 | 1 | 7.809 | 11.748 | 6.743 | 4.273 | 2.502 | 5.781 |
| | 7bgp | | | | ALD | MG-132 | P | | | 0.753 | 0.75 | 0.017 | 1.367 | 0.675 | 141O-144N | 1 | 1 | 1 | 1 | 1 | 7.808 | 11.525 | 6.778 | 0.975 | 0.719 | 0.215 | 1.426 | -2.979 | 141O-144N | 1 | 1 | 1 | 1 | 1 | 7.652 | 11.511 | 6.674 | -3.239 | -1.784 | 5.501 |
| | 7ca8_A | | | S̲G̲ of A,B | FNO | Shikonin | | | Chain B is ligand-free. C-terminal residues 299-306 do not exist. | -1.166 | -0.394 | -0.02 | 1.159 | 6.198 | | 1 | 0 | 0 | 0 | 1 | 8.472 | 13.067 | | | | | | | | | | | | | | | | 1.22 | 12.886 | 8.610 |
| | 7cb7 | | GSGGG | G̲S̲G̲G̲G̲ | B1S,K36 | GC376 | P | 0.89* | B1S,K36 are stereoisomers and are not distinguished. | 0.967 | 0.307 | 1.304 | -2.675 | -8.466 | 141O-144N | 1 | 1 | 0 | 1 | 1 | 7.214 | 10.835 | 9.220 | 0.509 | 1.402 | 0.868 | 0.532 | 27.909 | 141O-144N | 0 | 1 | 0 | 1 | 1 | 7.391 | 11.095 | 9.652 | -27.704 | 10.327 | 6.850 |
| | 7cut | | | | Z-VAD-FMK | Z-VAD-FMK | P | 0.59* | | 0.401 | 0.141 | -0.474 | 2.059 | -8.934 | 141O-144N | 1 | 1 | 1 | 1 | 1 | 7.758 | 11.545 | 6.730 | | | | | | | | | | | | | | | -0.001 | -10.874 | 5.124 |
| | 7cuu | | | | ALD | MG-132 | P | 3.91* | | 0.938 | 0.968 | -1.588 | -0.103 | -9.252 | 141O-144N | 1 | 1 | 1 | 1 | 1 | 7.451 | 11.158 | 6.033 | 1.248 | 0.706 | -0.33 | 0.517 | -9.227 | 141O-144N | 1 | 1 | 1 | 1 | 1 | 7.474 | 11.145 | 6.255 | 2.951 | -9.454 | 5.502 |
| | 7dk1 | | | | Zn | Zn | | 0.3251* | Three Zn exist (two at C145 and C156 of chain A and one at C145 of chain B). | 0.739 | 0.2 | -0.157 | 0.362 | -6.124 | 141O-144N | 1 | 1 | 1 | 1 | 1 | 7.670 | 11.425 | 6.857 | 0.648 | 0.276 | -0.416 | 0.257 | -3.16 | 141O-144N | 1 | 1 | 1 | 1 | 1 | 7.676 | 11.466 | 6.566 | -2.964 | -6.957 | 5.296 |
| | 7dpp | | | | MYC | Myricetin | | 0.63* | 302-306 do not exist | 0.09 | 0.753 | 18.39 | 1.63 | -4.801 | 141O-144N | 1 | 1 | 1 | 1 | 1 | 7.721 | 11.723 | 6.592 | | | | | | | | | | | | | | | 0 | -7.419 | 5.228 |
| | 7dpu | | | | HER | 3 | | 0.30* | | -3.132 | -2.594 | -0.605 | -1.758 | 2.383 | 140O-143N | 0 | 0 | 0 | 0 | 0 | 14.007 | 12.403 | 6.450 | 0.461 | -2.473 | -1.09 | -2.557 | -2.87 | 141O-144N | 1 | 1 | 1 | 1 | 1 | 7.291 | 11.557 | 6.402 | 1.948 | -0.731 | 5.830 |
| | 7dpv_AC | | | | HF0 | 7 | | 0.26* | Tetramer (two asymmetric dimers). | 0.294 | -0.261 | 0.74 | -0.04 | 4.338 | 141O-144N | 1 | 1 | 1 | 1 | 1 | 7.845 | 11.669 | 7.249 | -0.172 | -0.454 | 0.072 | 1.088 | -5.247 | 141O-144N | 0 | 1 | 0 | 0 | 1 | 7.326 | 12.125 | 6.577 | 3.38 | 0.051 | 5.968 |
| | 7dpv_BD | | | | HF0 | 7 | | 0.26* | | 0.263 | -0.203 | 0.925 | -0.241 | 4.505 | 141O-144N | 1 | 1 | 1 | 1 | 1 | 7.768 | 11.734 | 7.551 | -0.152 | -1.023 | 0.506 | 0.99 | -19.187 | 141O-144N | 0 | 1 | 0 | 0 | 1 | 7.375 | 11.987 | 6.451 | 12.614 | -12.208 | 5.882 |
| | 7e18 | | | | HUR | YH-53 | P | 0.0347 | | 0.142 | 1.494 | 0.476 | -3.419 | -4.251 | 141O-144N | 1 | 1 | 1 | 1 | 1 | 7.540 | 11.413 | 6.358 | | | | | | | | | | | | | | | 0 | -11.317 | 5.021 |
| | 7e19 | | | | HUO | SH-5 | P | 0.0145 | | 0.665 | 2.883 | -1.023 | -0.635 | -4.288 | 141O-144N | 1 | 1 | 1 | 1 | 1 | 7.453 | 11.253 | 6.358 | | | | | | | | | | | | | | | 0.001 | -8.024 | 5.516 |
| | 7ein | | | | PRD_000216 | Leupeptin | P | | | 0.917 | 0.696 | -0.461 | -2.025 | -12.461 | 141O-144N | 1 | 1 | 1 | 1 | 1 | 7.438 | 11.454 | 6.609 | 1.021 | 0.622 | -0.447 | -1.977 | -12.935 | 141O-144N | 1 | 1 | 1 | 1 | 1 | 7.406 | 11.402 | 6.686 | 0.245 | -13.643 | 5.297 |
| | 7fay | | | | 2XI | (R)-1a | | | | 0.31 | -0.519 | -1.852 | -1.004 | -5.241 | 141O-144N | 1 | 1 | 1 | 1 | 1 | 7.549 | 11.215 | 6.625 | | | | | | | | | | | | | | | 0.001 | -9.36 | 5.421 |
| | 7faz | | | | 2RI | Y180 | | | | 0.362 | -0.333 | 0.046 | 1.413 | -1.841 | 141O-144N | 1 | 1 | 1 | 1 | 1 | 7.663 | 11.225 | 6.692 | 0.415 | -0.563 | 0.296 | -0.655 | -3.518 | 141O-144N | 1 | 1 | 1 | 1 | 1 | 7.657 | 11.277 | 6.814 | -3.362 | -5.279 | 5.175 |
| | 7jpz | | | | GHX | MPI1 | P | 0.0979 0.100* | Cys128 and Cys300 are mono-oxydized to be CSO (hydroxycysteine). | 0.845 | -2.031 | -1.243 | -2.54 | -6.463 | 141O-144N | 1 | 1 | 1 | 1 | 1 | 7.272 | 11.287 | 6.236 | | | | | | | | | | | | | | | -0.001 | -9.468 | 5.392 |
| | 7jq0 | | | | VHV | MPI3 | P | 0.0083 0.0085* | Cys128 and Cys300 are mono-oxydized to be CSO (hydroxycysteine). | 1.244 | 0.71 | -0.893 | -2.235 | -7.62 | 141O-144N | 1 | 1 | 1 | 1 | 1 | 7.378 | 11.167 | 6.225 | | | | | | | | | | | | | | | -0.001 | -11.455 | 5.545 |
| | 7jq1 | | | | VHJ | MPI4 | P | 0.0147 0.015* | Cys128 and Cys300 are mono-oxydized to be CSO (hydroxycysteine). | 0.87 | 0.283 | -0.626 | -1.755 | -7.181 | 141O-144N | 1 | 1 | 1 | 1 | 1 | 7.393 | 11.308 | 6.446 | | | | | | | | | | | | | | | -0.002 | -10.807 | 5.370 |
| | 7jq2 | | | | VHM | MPI5 | P | 0.0322 0.033* | Cys128 is mono-oxydized to be CSO (hydroxycysteine). | 0.937 | 0.088 | -1.051 | -2.408 | -7.104 | 141O-144N | 1 | 1 | 1 | 1 | 1 | 7.395 | 11.199 | 6.370 | | | | | | | | | | | | | | | 0 | -10.158 | 5.364 |
| | 7jq3 | | | | VHP | MPI6 | P | 0.0586 0.060* | Cys128 and Cys300 are mono-oxydized to be CSO (hydroxycysteine). | 0.988 | -0.111 | -0.925 | -1.313 | -7.336 | 141O-144N | 1 | 1 | 1 | 1 | 1 | 7.348 | 11.339 | 6.377 | | | | | | | | | | | | | | | 0 | -10.078 | 5.278 |
| | 7jq4 | | | | XM2 | MPI7 | P | 0.0459 0.047* | Cys128 and Cys300 are mono-oxydized to be CSO (hydroxycysteine). | 0.99 | -0.538 | -0.844 | -1.791 | -8.135 | 141O-144N | 1 | 1 | 1 | 1 | 1 | 7.427 | 11.390 | 6.365 | | | | | | | | | | | | | | | -0.001 | -10.182 | 5.286 |
| | 7jq5 | | | | NOL | MPI8 | P | 0.1026 0.105* | Cys128 and Cys300 are mono-oxydized to be CSO (hydroxycysteine). | 0.949 | 0.04 | -0.735 | -1.583 | -8.522 | 141O-144N | 1 | 1 | 1 | 1 | 1 | 7.456 | 11.379 | 6.142 | | | | | | | | | | | | | | | 0.001 | -10.749 | 5.249 |

| | | | | | | | | | | | | | | | | | | | | | | | | | | | | | | | | | | | | | | | |
|---|---|---|---|---|---|---|---|---|---|---|---|---|---|---|---|---|---|---|---|---|---|---|---|---|---|---|---|---|---|---|---|---|---|---|---|---|---|---|---|
| 7jsu | | | | UED | GC376 | P | | | 1.047 | 0.879 | -0.052 | -0.719 | -9.114 | 141O-144N | 1 | 1 | 1 | 1 | 1 | 7.342 | 11.097 | 6.291 | | | | | | | | | | | | | | | 0.001 | -11.418 | 5.785 |
| 7jt0 | | | | LW1 | MAC-5576 | | | Chloropxypyridine in MAC-5576 is reduced to be LW1. | 0.481 | 1.061 | 3.173 | 1.267 | -6.907 | 141O-144N | 1 | 1 | 1 | 1 | 1 | 7.430 | 11.494 | 6.350 | | | | | | | | | | | | | | | 0 | -9.118 | 5.605 |
| 7jt7 | | | | TG3 | 4 | P | | | 0.601 | 1.57 | -0.764 | 0.948 | 1.359 | 141O-144N | 1 | 1 | 1 | 1 | 1 | 7.428 | 11.100 | 6.915 | | | | | | | | | | | | | | | 0.001 | -2.831 | 5.596 |
| 7jw8_AB | | | | TG3 | 4 | P | | Tetramer (two hheterodimers). Space group of P1. | 0.802 | 1.708 | -0.829 | 0.198 | 2.182 | 141O-144N | 1 | 1 | 1 | 1 | 1 | 7.338 | 11.067 | 6.892 | 0.734 | 1.531 | -0.397 | 0.687 | -0.389 | 141O-144N | 1 | 1 | 1 | 1 | 1 | 7.450 | 11.117 | 6.398 | -2.082 | -3.243 | 5.641 |
| 7jw8_CD | | | | TG3 | 4 | P | | | 0.886 | 1.597 | -0.47 | 0.756 | -3.862 | 141O-144N | 1 | 1 | 1 | 1 | 1 | 7.355 | 11.087 | 6.184 | 0.825 | 1.973 | -0.717 | 0.645 | 2.03 | 141O-144N | 1 | 1 | 1 | 1 | 1 | 7.324 | 11.025 | 6.669 | -0.69 | -4.698 | 5.655 |
| 7k0e | | SNIG | SNIGS of A,SNIGSGFR of B | K36 | 2 | P | 0.18*<br>0.086[+]<br>0.069[+] | | 0.563 | 0.438 | 0.591 | -2.568 | -11.146 | 141O-144N | 1 | 1 | 0 | 1 | 1 | 7.247 | 10.863 | 6.348 | 0.635 | 0.715 | 0.188 | -1.155 | 33.482 | 141O-144N | 1 | 1 | 0 | 1 | 1 | 7.438 | 11.131 | | -31.224 | 13.284 | 7.820 |
| 7k0f | | SNIG | SNIGSGF of A and B | VR4 | 5 | P | 1.31* | | 0.607 | 0.834 | 0.318 | -2.659 | -14.107 | 141O-144N | 1 | 1 | 0 | 1 | 1 | 7.599 | 11.283 | | 0.339 | 0.334 | 0.546 | -1.668 | 39.172 | 141O-144N | 1 | 1 | 0 | 1 | 1 | 7.377 | 11.222 | | -36.973 | 15.988 | 8.009 |
| 7kvl_A | | GAM | | X4P | FMAX E01 | | | X4P is a small compound located on the surface of domain III. | -2.274 | -0.545 | 0.161 | 1.481 | 9.464 | | 1 | 0 | 0 | 1 | 0 | 13.961 | 8.949 | 10.162 | | | | | | | | | | | | | | | -4.725 | 19.553 | 10.009 |
| 7kvr_B | | GAM | GAM of A, GAMS of B | XV4 | FMAX E09 | | | XV4 is a small compound located on the surface of domain III. | | | | | | | | | | | | | | | -1.509 | 0.119 | 2.26 | 1.815 | 7.42 | | 1 | 0 | 0 | 0 | 1 | 10.777 | 11.892 | 7.958 | -2.043 | 13.813 | 9.542 |
| 7kx5 | | | | X7V | Jun8-76-3A | | | 1-oxybenzotriazole in X7V is replaced by C145. | 0.797 | -0.994 | -1.472 | 0.267 | -10.026 | 141O-144N | 1 | 1 | 1 | 1 | 1 | 7.336 | 11.178 | 6.010 | | | | | | | | | | | | | | | 0.001 | -16.056 | 4.840 |
| 7kyu | | | | XC4 | 1H-indole-5-carboxylate | | | | 0.405 | 0.65 | 0.517 | 1.693 | -6.392 | 141O-144N | 1 | 1 | 1 | 1 | 1 | 7.717 | 11.485 | 6.643 | | | | | | | | | | | | | | | 0.001 | -10.877 | 5.351 |
| 7l0d | | | | 0EN | ML188 | | 2.5* | | 0.591 | -0.913 | -1.61 | -0.719 | -8.738 | 141O-144N | 1 | 1 | 1 | 1 | 1 | 7.736 | 11.540 | 6.218 | | | | | | | | | | | | | | | 0.001 | -12.377 | 5.134 |
| 7l10 | | | | XEY | 4 | | | | 0.621 | -0.484 | 1.411 | 1.861 | -4.572 | 141O-144N | 1 | 1 | 1 | 1 | 1 | 7.868 | 11.729 | 6.521 | | | | | | | | | | | | | | | 0.002 | -7.194 | 5.317 |
| 7l11 | | | | XF1 | 5 | | 2.5[+]<br>1.5[+] | | 0.434 | -0.35 | -1.164 | 0.484 | 4.795 | 141O-144N | 1 | 1 | 1 | 1 | 1 | 7.941 | 11.709 | 7.440 | 0.607 | -0.127 | 0.658 | -0.748 | 3.526 | 141O-144N | 1 | 1 | 1 | 1 | 1 | 7.947 | 11.671 | 6.894 | -3.581 | 2.524 | 5.903 |
| 7l12 | | | | XF4 | 14 | | 3.2[+] | | 0.329 | -1.203 | -0.984 | -1.702 | -2.678 | 141O-144N | 1 | 1 | 1 | 1 | 1 | 7.839 | 11.504 | 6.588 | | | | | | | | | | | | | | | 0 | -6.177 | 5.152 |
| 7l13_A | | | | XF7 | 21 | | 11.3[+] | Chain B is ligand-unbound. | 0.749 | -1.485 | -0.377 | -0.409 | 1.567 | 141O-144N | 1 | 1 | 1 | 1 | 1 | 7.731 | 11.369 | 7.493 | | | | | | | | | | | | | | | 4.833 | -1.553 | 5.504 |
| 7l14 | | | | XFD | 26 | | 2.0[+]<br>0.98[+] | | 0.524 | 0.429 | -1.199 | 1.401 | 4.677 | 141O-144N | 1 | 1 | 1 | 1 | 1 | 8.001 | 11.788 | 7.218 | 0.665 | 0.118 | 0.603 | 1.408 | -1.426 | 141O-144N | 1 | 1 | 1 | 1 | 1 | 7.930 | 11.675 | 6.835 | -0.38 | 0.494 | 5.480 |
| 7l5d | | | | XNJ | demethylated analog of masitinib | | | | 0.384 | 0.236 | -0.304 | 1.924 | -8.086 | 141O-144N | 1 | 1 | 1 | 1 | 1 | 7.725 | 11.260 | 6.430 | | | | | | | | | | | | | | | 0 | -12.961 | 5.177 |
| 7lb7 | | | | PRD_002448 | Telaprevir | P | | Neutron diffraction determining the H positions. | 0.679 | 0.387 | -0.173 | -1.419 | -0.27 | 141O-144N | 1 | 1 | 1 | 1 | 1 | 7.814 | 11.342 | 6.698 | | | | | | | | | | | | | | | 0 | -3.815 | 5.913 |
| 7lbn | | | | PRD_002460 | Calpain I | P | | | 0.968 | 0.371 | -0.416 | 2.009 | -9.944 | 141O-144N | 1 | 1 | 1 | 1 | 1 | 7.507 | 11.246 | 6.202 | | | | | | | | | | | | | | | 0.001 | -12.738 | 5.370 |
| 7lco | | | | XTJ | 1i | P | 0.14*<br>1.1[+] | | 1.163 | 0.157 | -0.723 | -2.727 | -9.175 | 141O-144N | 1 | 1 | 1 | 1 | 1 | 7.227 | 10.940 | 6.217 | | | | | | | | | | | | | | | 0 | -11.315 | 5.341 |
| 7lcs | | | | XTP | 1a | P | 0.05*<br>1.1[+] | | 0.962 | 0.283 | -1.216 | -2.999 | -10.364 | 141O-144N | 1 | 1 | 1 | 1 | 1 | 7.382 | 11.064 | 6.234 | | | | | | | | | | | | | | | 0.001 | -12.38 | 5.372 |
| 7lct | | | | XU4 | 1g | P | 0.27<br>1.1[+] | | 0.997 | 1.708 | -0.469 | -3.472 | -2.16 | 141O-144N | 1 | 1 | 1 | 1 | 1 | 7.206 | 10.920 | 6.488 | | | | | | | | | | | | | | | -0.001 | -6.006 | 5.892 |
| 7ldl | | | | XV4 | 1f | P | 0.06*<br>0.9[+] | | 0.836 | 0.717 | 1.008 | -2.714 | -5.606 | 141O-144N | 1 | 1 | 1 | 1 | 1 | 7.783 | 11.605 | 6.612 | 0.805 | 1.098 | 0.497 | -2.757 | -4.067 | 141O-144N | 1 | 1 | 1 | 1 | 1 | 7.774 | 11.674 | 6.551 | -1.203 | -8.473 | 5.407 |
| 7ldx | | GAM | GAM of B | R9V | F2X E06 | | | Small ligand R9V is on the interfavcce between domain I and III. | 0.25 | -0.703 | 0.217 | 0.799 | 9.302 | | 0 | 0 | 0 | 0 | 1 | 8.955 | 12.568 | 10.136 | -10.835 | 1.098 | 2.185 | 2.142 | 14.339 | 139O-142N<br>140O-143N | 0 | 0 | 0 | 0 | 0 | 9.885 | 14.054 | 7.643 | -4.945 | 18.997 | 9.873 |
| 7lfe_A | | GAM | | XWS | F2X E03 | | | Chain B is ligand-unbound. | -2.869 | -0.476 | 0.038 | 1.181 | 9.482 | | 0 | 0 | 0 | 0 | 0 | 13.092 | 11.059 | 10.211 | | | | | | | | | | | | | | | -2.15 | 17.03 | 9.750 |
| 7lkr | | SNIG | SNIGSGFR of A, SNIGSGF of B | Y91,Y8Y | 2a | P | 0.18* | Stereoisomers, Y91,Y8Y, are not distinguished. Sulfoxide of the ligand is replaced by Sg of C145. Q306 does not exist. | 0.565 | 0.288 | 0.801 | -2.534 | -12.944 | 141O-144N | 1 | 1 | 0 | 1 | 1 | 7.307 | 10.868 | | 0.588 | 0.628 | 0.363 | -2.218 | 36.895 | 141O-144N | 1 | 1 | 0 | 1 | 1 | 7.469 | 11.274 | | -35.029 | 16.177 | 7.912 |
| 7lks | | SNIG | SNIGSG of A, SNIGSG of B | Y4P,Y7G | 2f | P | 0.14* | Stereoisomers, Y4P,Y7G, are not distinguished. Sulfoxide of the ligand is replaced by Sg of C145. Q306 does not exist. | 0.359 | 0.726 | 0.509 | -3.016 | -13.803 | 141O-144N | 1 | 1 | 0 | 1 | 1 | 7.303 | 10.872 | | 0.309 | 0.212 | -0.067 | -2.441 | 38.365 | 141O-144N | 1 | 1 | 0 | 1 | 1 | 7.497 | 11.469 | | -34.904 | 16.69 | 7.806 |
| 7lkt | | SNIG | SNIGSGF of A, SNIGS of B | Y4V,Y7M | 2k | P | 0.18* | Stereoisomers, Y4V and Y7M, are not distinguished. Sulfoxide of the ligand is replaced by Sg of C145. Q306 does not exist. | 0.763 | 0.908 | 1.042 | -3.165 | -11.059 | 141O-144N | 1 | 1 | 0 | 1 | 1 | 7.370 | 10.958 | | 0.736 | 0.447 | -0.206 | -2.878 | 34.199 | 141O-144N | 1 | 1 | 0 | 1 | 1 | 7.554 | 11.405 | 9.509 | -31.798 | 14.154 | 7.904 |
| 7lku | | SNIG | SNIG of A, SNIGSG of B | Y91,Y8Y,Y5S, Y4D | 3b | P | 0.29* | Stereoisomers, Y91,Y8Y,Y5S,Y4D, are not distinguished. Sulfoxide of the ligand is replaced by Sg of C145. Q306 does not exist. | 0.605 | 0.378 | 0.639 | -2.385 | -16.332 | 141O-144N | 1 | 1 | 0 | 1 | 1 | 7.303 | 10.866 | 6.574 | 0.669 | 0.53 | 0.362 | -2.081 | 37.893 | 141O-144N | 1 | 1 | 0 | 1 | 1 | 7.623 | 11.523 | | -37.685 | 13.315 | 8.003 |
| 7lkv | | SNIG | SNIGS of A, SNIGSGF of B | Y64,Y4J | 3c | P | 0.26* | Stereoisomers, Y64,Y4J, are not distinguished. Sulfoxide of the ligand is replaced by Sg of C145. Q306 does not exist. | 0.616 | 0.342 | 1.186 | -2.436 | -13.375 | 141O-144N | 1 | 1 | 0 | 1 | 1 | 7.305 | 10.949 | 6.359 | 0.71 | 0.997 | 0.354 | -0.917 | 36.859 | 141O-144N | 1 | 1 | 0 | 1 | 1 | 7.356 | 11.149 | | -36.105 | 15.418 | 7.996 |
| 7lkw | | SNIG | SNIGSGF of A, SNIGSG of B | Y8V,Y8S | 3d | P | 0.30* | Stereoisomers, Y8V,Y8S, are not distinguished. Sulfoxide of the ligand is replaced by Sg of C145. Q306 does not exist. | 0.74 | 0.99 | -0.03 | -2.818 | -14.148 | 141O-144N | 1 | 1 | 0 | 1 | 1 | 7.316 | 10.890 | | 0.698 | 0.47 | 0.305 | -1.994 | 38.813 | 141O-144N | 1 | 1 | 0 | 1 | 1 | 7.382 | 11.251 | | -37.541 | 14.396 | 8.083 |
| 7lkx | | SNIG | SNIGSGFR of A, SNIGSG of B | Y71,Y51 | 3e | P | 0.28* | Stereoisomers, Y71,Y51, are not distinguished. Sulfoxide of the ligand is replaced by Sg of C145. Q306 does not exist. | 0.548 | 0.785 | 0.466 | -2.783 | -12.755 | 141O-144N | 1 | 1 | 0 | 1 | 1 | 7.318 | 10.921 | | 0.614 | 0.567 | 0.42 | -2.08 | 35.954 | 141O-144N | 1 | 1 | 0 | 1 | 1 | 7.516 | 11.392 | | -34.262 | 14.856 | 7.683 |
| 7lmc_AB | C145A | | | | C-term of chain C & C-term of Nsp4C | P | | Two heterodimers. Chain A binds the C-terminus of chain C in the asymmetric unit. Chain B binds 6 residue peptide of C-term of Nsp4C. | 0.809 | 0.43 | -0.219 | 2.103 | -7.777 | 141O-144N | 1 | 1 | 1 | 1 | 1 | 7.430 | 10.876 | 7.038 | 0.466 | 0.037 | -0.436 | 0.243 | 1.542 | 141O-144N | 1 | 1 | 1 | 1 | 1 | 7.950 | 11.597 | 6.375 | -9.102 | -4.462 | 6.047 |
| 7lmc_CD | C145A | | | | C-term of chain C & C-term of Nsp4C | P | | Two heterodimers. Chain C binds the C-terminus of chain C in the asymmetric unit. Chain D binds 6 residue peptide of C-term of Nsp4C. | 0.737 | 1.462 | -0.696 | 2.203 | -5.224 | 141O-144N | 0 | 1 | 1 | 1 | 1 | 7.619 | 10.761 | 7.501 | 0.999 | -0.65 | -0.925 | -0.264 | 2.282 | 141O-144N | 1 | 1 | 1 | 1 | 1 | 7.309 | 10.478 | 6.550 | -6.236 | -0.369 | 6.162 |
| 7ltj | | | | YD1 | Mcule-5948770040 | | | | 0.58 | -0.3 | -0.245 | -1.19 | -4.082 | 141O-144N | 1 | 1 | 1 | 1 | 1 | 7.717 | 11.533 | 6.586 | | | | | | | | | | | | | | | 0 | -6.481 | 5.433 |
| 7lyh | | | | YHJ | UAWJ9-36-1 | P | | | 1.294 | 1.305 | -1.032 | -2.567 | -11.197 | 141O-144N | 1 | 1 | 1 | 1 | 1 | 7.147 | 10.906 | 6.320 | | | | | | | | | | | | | | | 0 | -11.225 | 5.191 |
| 7lyi | | HM | HMS | YHI | UAWJ9-36-3 | P | | | 1.367 | -0.737 | 0.397 | -2.077 | -10.371 | 141O-144N | 1 | 1 | 0 | 1 | 1 | 7.439 | 11.148 | 6.573 | | | | | | | | | | | | | | | 0 | -6.746 | 5.567 |
| 7lzt | | SNIG | SNIGS | YN1,YMY | 8b | P | | | 1.223 | 0.895 | -1.006 | -3.284 | -15.355 | 141O-144N | 1 | 1 | 0 | 1 | 1 | 7.250 | 10.915 | 6.523 | | | | | | | | | | | | | | | 0 | -13.084 | 5.221 |
| 7lzu | | SNIG | SNIGSGFR of A,SNIGSG of B | YKP,YKM | 12b | P | | Stereoisomers,YKP,YKM, are not distinguished. Sulfoxide of the ligand is replaced by Sg of C145. Q306 does not exist. | 0.714 | 0.6 | 0.52 | -2.903 | -12.141 | 141O-144N | 1 | 1 | 0 | 1 | 1 | 7.394 | 10.976 | | 0.454 | 0.287 | 0.329 | -2.847 | 35.032 | 141O-144N | 1 | 1 | 0 | 1 | 1 | 7.438 | 11.315 | | -33.345 | 15.233 | 7.233 |
| 7lzv | | SNIG | SNIGSGF of A,B | YLS,YLM | 19b | P | | Stereoisomers,YLS,YLM, are not distinguished. Sulfoxide of the ligand is replaced by Sg of C145. Q306 does not exist. | 0.667 | 0.48 | 0.515 | -2.602 | -12.916 | 141O-144N | 1 | 1 | 0 | 1 | 1 | 7.321 | 10.891 | | 0.531 | 0.372 | 0.408 | -1.704 | 37.87 | 141O-144N | 1 | 1 | 0 | 1 | 1 | 7.414 | 11.159 | | -35.73 | 17.323 | 8.148 |
| 7lzw | | SNIG | SNIGSGF of A,B | YLS,YLM | 20b | P | | 20b is deuterated 19b. Stereoisomers,YLS,YLM, are not distinguished. Sulfoxide of the ligand is replaced by Sg of C145. Q306 does not exist. | 0.634 | 0.326 | 0.661 | -2.4 | -15.848 | 141O-144N | 1 | 1 | 0 | 1 | 1 | 7.254 | 10.871 | | 0.743 | 0.419 | 0.352 | -1.417 | 36.482 | 141O-144N | 1 | 1 | 0 | 1 | 1 | 7.319 | 10.906 | | -36.529 | 13.83 | 8.303 |
| 7lzx | | SNIG | SNIGSG of A,SNIG of B | YMD,YMG | 1c | P | | Stereoisomers,YMD,YMG,are not distinguished. Sulfoxide of the ligand is replaced by Sg of C145. Q306 does not exist. | 0.684 | 0.886 | 0.609 | -2.502 | -14.361 | 141O-144N | 1 | 1 | 0 | 1 | 1 | 7.294 | 10.962 | | 0.683 | 0.726 | 0.329 | -2.183 | 41.125 | 141O-144N | 1 | 1 | 0 | 1 | 1 | 7.340 | 11.047 | 9.617 | -37.902 | 18.769 | 8.896 |
| 7lzy | | SNIG | SNIGSG | YMM,YMJ | 3c | P | | Stereoisomers,YMM,YMJ,are not distinguished. Sulfoxide of the ligand is replaced by Sg of C145. Q306 does not exist. | 0.908 | 1.108 | -1.013 | -3.548 | -12.805 | 141O-144N | 1 | 1 | 0 | 1 | 1 | 7.455 | 11.150 | | | | | | | | | | | | | | | | -0.001 | -12.817 | 5.317 |
| 7lzz | | SNIG | SNIGSG | YMV,YMS | 5c | P | | Stereoisomers,YKY,YL7,are not distinguished. Sulfoxide of the ligand is replaced by Sg of C145. Q306 does not exist. | 0.856 | 1.09 | -1.007 | -3.447 | -13.016 | 141O-144N | 1 | 1 | 0 | 1 | 1 | 7.411 | 11.178 | | | | | | | | | | | | | | | | 0.001 | -14.511 | 5.057 |
| 7m00 | | SNIG | SNIGSG of A,B | YKD,YKA | 13c | P | | Stereoisomers,YKD,YKA,are not distinguished. Sulfoxide of the ligand is replaced by Sg of C145. Q306 does not exist. | 0.67 | 0.978 | -0.594 | -3.168 | -11.175 | 141O-144N | 1 | 1 | 0 | 1 | 1 | 7.473 | 11.246 | | 0.56 | 1.021 | -0.625 | -3.21 | -11.013 | 141O-144N | 1 | 1 | 0 | 1 | 1 | 7.494 | 11.239 | | 0.26 | -13.417 | 5.276 |
| 7m01 | | SNIG | SNIGSGF of A,SNIGSG of B | YKS,YKV | 14c | P | | Stereoisomers,YKS,YKV,are not distinguished. Sulfoxide of the ligand is replaced by Sg of C145. Q306 does not exist. | 0.701 | 0.716 | 0.215 | -2.854 | -13.576 | 141O-144N | 1 | 1 | 0 | 1 | 1 | 7.299 | 10.911 | | 0.496 | 0.57 | 0.362 | -2.23 | 33.27 | 141O-144N | 1 | 1 | 0 | 1 | 1 | 7.504 | 11.301 | | -33.242 | 12.345 | 7.179 |
| 7m02 | | SNIG | SNIGSGF of A,SNIGSG of B | YKY,YL7 | 17c | P | | Stereoisomers,YM1,YLV,are not distinguished. Sulfoxide of the ligand is replaced by Sg of C145. Q306 does not exist. | 0.511 | 0.709 | 0.506 | -2.936 | -14.387 | 141O-144N | 1 | 1 | 0 | 1 | 1 | 7.217 | 10.814 | | 0.436 | 0.629 | -0.059 | -2.73 | 36.926 | 141O-144N | 1 | 1 | 0 | 1 | 1 | 7.491 | 11.382 | | -34.551 | 14.718 | 7.625 |

| | | | | | | | | | | | | | | | | | | | | | | | | | | | | | | | | | | | | | | | |
|---|---|---|---|---|---|---|---|---|---|---|---|---|---|---|---|---|---|---|---|---|---|---|---|---|---|---|---|---|---|---|---|---|---|---|---|---|---|---|---|
| 7m03 | | SNIG | SNIGSGF of A,SNIGSG of B | YLJ,YLD | 18c | P | | Stereoisomers,YLJ,YLD,are not distinguished. Sulfoxide of the ligand is replaced by Sg of C145. Q306 does not exist. | 0.625 | 0.442 | 0.904 | -2.658 | -14.17 | 141O-144N | 1 | 1 | 0 | 1 | 1 | 7.301 | 10.782 | | 0.56 | 0.511 | -0.044 | -1.788 | 36.999 | 141O-144N | 1 | 1 | 0 | 1 | 1 | 7.455 | 11.204 | | -34.84 | 15.003 | 8.017 |
| 7m04 | | SNIG | SNIGS of A,SNIGSG of B | YM1,YLV | 21c | P | | Stereoisomers,YM1,YLV,are not distinguished. Sulfoxide of the ligand is replaced by Sg of C145. Q306 does not exist. | 0.69 | 0.742 | -0.098 | -2.787 | -14.142 | 141O-144N | 1 | 1 | 0 | 1 | 1 | 7.377 | 11.055 | 6.642 | 0.537 | 0.644 | 0.367 | -1.964 | 36.463 | 141O-144N | 1 | 1 | 0 | 1 | 1 | 7.569 | 11.390 | | -35.454 | 15.682 | 7.784 |
| 7m8m_A | | | | YSG | 11 | | 0.120* | Chain B is ligand-unbound. | 0.6 | -0.028 | 0.053 | 1.224 | 1.802 | 141O-144N | 1 | 1 | 1 | 1 | 1 | 7.694 | 11.292 | 7.057 | | | | | | | | | | | | | | | 4.404 | -2.185 | 5.293 |
| 7m8n | | | | YSP | 16 | | 0.1* | | 0.809 | -1.17 | -0.978 | -0.656 | 1.646 | 141O-144N | 1 | 1 | 1 | 1 | 1 | 7.607 | 11.102 | 7.228 | 0.771 | -1.094 | 0.478 | -0.305 | -1.808 | 141O-144N | 1 | 1 | 1 | 1 | 1 | 7.741 | 11.332 | 6.939 | -0.966 | 0.343 | 5.626 |
| 7m8o_A | | | | YSM | 19 | | 0.037* | Chain B is ligand-unbound. | 0.296 | -0.987 | -0.311 | -1.033 | 3.126 | 141O-144N | 1 | 1 | 1 | 1 | 1 | 7.650 | 11.282 | 7.548 | | | | | | | | | | | | | | | 4.64 | -3.14 | 5.541 |
| 7m8p_A | | | | YSJ | 23 | | 0.020* | Chain B is ligand-unbound. | 0.62 | -1.622 | -0.059 | 0.506 | 1.925 | 141O-144N | 1 | 1 | 1 | 1 | 1 | 7.566 | 11.255 | 7.581 | | | | | | | | | | | | | | | 1.054 | -0.663 | 5.795 |
| 7m8x | | | | YTJ | 6 | | 0.47* | | 0.42 | -0.576 | 4.787 | -0.452 | -4.038 | 141O-144N | 1 | 1 | 1 | 1 | 1 | 7.962 | 11.892 | 6.679 | | | | | | | | | | | | | | | -0.001 | -7.524 | 5.172 |
| 7m8y | | | | YTM | 15 | | 0.110* | | 0.439 | -1.013 | -1.348 | 0.032 | -2.055 | 141O-144N | 1 | 1 | 1 | 1 | 1 | 7.715 | 11.292 | 6.755 | | | | | | | | | | | | | | | 0.001 | -5.486 | 5.270 |
| 7m8z | | | | YTV | 29 | | 0.25-0.50* | | 0.569 | 0.178 | 4.584 | 0.874 | -3.911 | 141O-144N | 1 | 1 | 1 | 1 | 1 | 7.633 | 11.205 | 6.710 | | | | | | | | | | | | | | | 0 | -6.899 | 5.351 |
| 7m90 | | | | YTS | 50 | | 0.25-0.50* | | 0.436 | 0.51 | 1.331 | 1.115 | 0.464 | 141O-144N | 1 | 1 | 1 | 1 | 1 | 7.849 | 11.537 | 6.923 | | | | | | | | | | | | | | | 0.001 | -3.391 | 5.535 |
| 7m91 | | | | YU4 | 25 | | 0.025* | | 0.791 | -0.342 | 1.347 | 1.015 | -5.03 | 141O-144N | 1 | 1 | 1 | 1 | 1 | 7.754 | 11.448 | 6.649 | | | | | | | | | | | | | | | 0.001 | -7.138 | 5.542 |
| 7mbi_AD | | | | YWJ | 15l | P | 0.019* 0.30[+] | Tetramer (two heterodimers). Trimethylpyridine-3-carboxylate of 15l is replaced by Sg of C145 to be YWJ. | 0.63 | 0.822 | 0.23 | -3.042 | -10.168 | 141O-144N | 1 | 1 | 1 | 1 | 1 | 7.495 | 11.153 | 6.425 | 0.28 | 1.578 | -1.047 | -3.152 | 0.995 | 141O-144N | 1 | 1 | 1 | 1 | 1 | 7.503 | 11.331 | 6.436 | -9.862 | -5.929 | 5.809 |
| 7mbi_BC | | | | YWJ | 15l | P | 0.019* 0.30[+] | | 0.336 | 1.906 | 2.801 | -2.882 | -3.082 | 141O-144N | 1 | 1 | 1 | 1 | 1 | 7.473 | 11.116 | 6.413 | 0.497 | 1.277 | 0.344 | -3.586 | -0.519 | 141O-144N | 1 | 1 | 1 | 1 | 1 | 7.380 | 11.099 | 6.565 | 2.081 | -0.319 | 5.818 |
| 7mgr | C145A | | | AVKLQNNE | nsp8/9 substrate peptide | P | | The substrate is C-term 5res of Nsp8 and N-term 4res of Nsp9, AVKLQNNEL; the last L is missing. | 0.605 | -0.181 | -0.037 | -1.321 | -10.396 | 141O-144N | 1 | 1 | 1 | 1 | 1 | 7.282 | 10.685 | 6.460 | | | | | | | | | | | | | | | 0.001 | -12.234 | 5.648 |
| 7mgs | C145A | | | SAVLQSGF | N-term auto-processing substrate | P | | The substrate is the C-terminal 8res peptide of Nsp4C, SAVLQSGF | 0.965 | 1.489 | -0.624 | 0.594 | -9.242 | 141O-144N | 1 | 1 | 1 | 1 | 1 | 7.325 | 10.988 | 6.529 | | | | | | | | | | | | | | | 0.001 | -8.894 | 5.994 |
| 7mng | | | | ZL7 | VBY-825 | P | | ZL7 contains two models. | 0.651 | -1.598 | -1.33 | -3.353 | -6.069 | 141O-144N | 1 | 1 | 1 | 1 | 1 | 7.666 | 11.681 | 6.118 | | | | | | | | | | | | | | | -0.001 | -10.642 | 5.217 |
| 7mpb | | | | ASC | Ascorbate | | | After Q306, GPHHHHHH is added. | 0.857 | -1.766 | -1.173 | -2.607 | -8.102 | 141O-144N | 1 | 1 | 1 | 1 | 1 | 7.780 | 11.750 | 6.620 | 0.756 | -0.892 | -0.4 | 0.158 | 0.552 | 141O-144N | 1 | 1 | 1 | 1 | 1 | 7.507 | 11.469 | 6.692 | -8.769 | -6.705 | 5.730 |
| 7mrr | | | | PRD_000216 | Leupeptin | P | | | 0.875 | 0.202 | -0.849 | -0.781 | -13.296 | 141O-144N | 1 | 1 | 1 | 1 | 1 | 7.492 | 11.737 | 6.362 | | | | | | | | | | | | | | | 0.001 | -15.511 | 5.223 |
| 7n44 | | | | 06I | 13 | | 0.042* | | 0.384 | -1.078 | 3.139 | -2.401 | -1.979 | 141O-144N | 1 | 1 | 1 | 1 | 1 | 7.947 | 11.577 | 6.601 | | | | | | | | | | | | | | | 0 | -5.446 | 5.482 |
| 7n6n | C145S | GAMSAVLQ | GAMSAVLQ | GAMSAVLQ | N-terminal & C-terminal substrates | P | | Chain A binds SAVLQ, N-terminal of Nsp4C. Chain B binds C-terminus of chain A in another unit. GAM of GAMSAVLQ (C-term of Nsp4C) is missing. | 0.406 | 0.359 | -1.893 | -1.729 | -5.065 | 141O-144N | 1 | 1 | 1 | 1 | 1 | 7.520 | 11.045 | 6.469 | 1.189 | 0.047 | 0.248 | 2.267 | -11.621 | 141O-144N | 1 | 1 | 1 | 1 | 1 | 7.527 | 10.922 | 8.912 | 10.819 | -4.564 | 6.120 |
| 7n89 | C145A | | | SAVLQSGF | substrate | P | | The termials of the substrate are caped by ACE and NH2, respectively. | 0.88 | 0.23 | -1.096 | -2.482 | -14.168 | 141O-144N | 1 | 1 | 1 | 1 | 1 | 7.349 | 10.873 | 6.438 | 0.953 | 0.291 | -0.878 | -0.942 | -9.839 | 141O-144N | 1 | 1 | 1 | 1 | 1 | 7.323 | 10.859 | 6.606 | -3.043 | -11.271 | 5.525 |
| 7n8c | | | | YD1 | Mcule5948770040 | | | | 0.544 | -0.291 | -0.369 | -1.34 | -3.793 | 141O-144N | 1 | 1 | 1 | 1 | 1 | 7.736 | 11.563 | 6.807 | | | | | | | | | | | | | | | -0.001 | -5.717 | 5.508 |
| 7nbr | | | | PRD_002382 | Boceprevir | P | | | 1.076 | 1.73 | -0.473 | -0.867 | -12.363 | 141O-144N | 1 | 1 | 1 | 1 | 1 | 7.527 | 11.487 | 6.397 | | | | | | | | | | | | | | | 0 | -15.196 | 5.234 |
| 7nbs | | | | SV6 | Telaprevir | P | | | 0.779 | 0.308 | -0.269 | -1.2 | -2.199 | 141O-144N | 1 | 1 | 1 | 1 | 1 | 7.572 | 11.365 | 6.511 | | | | | | | | | | | | | | | 0.001 | -5.504 | 5.746 |
| 7nby[x] | | | | U88 | SU3327 | | | Four U88 per protomer bind to four distinct sites, respectively. | 0.578 | 0.647 | -0.285 | 1.104 | 5.446 | 141O-144N | 1 | 1 | 1 | 1 | 1 | 7.512 | 11.466 | 7.470 | 0.879 | 0.582 | 13.875 | 1.155 | -4.962 | 141O-144N | 1 | 1 | 1 | 1 | 1 | 7.524 | 11.407 | 6.943 | 3.622 | 1.667 | 5.753 |
| 7nf5 | | | | ALD | MG-132 | P | | | 0.949 | 0.439 | 0.063 | -0.209 | -1.309 | 141O-144N | 1 | 1 | 1 | 1 | 1 | 7.440 | 11.139 | 6.254 | | | | | | | | | | | | | | | -0.001 | -4.845 | 5.895 |
| 7ng3 | | | | ALD | MG-132 | P | | | 1.23 | 1.465 | -1.032 | 0.698 | -9.441 | 141O-144N | 1 | 1 | 1 | 1 | 1 | 7.452 | 11.137 | 6.359 | 1.004 | 1.544 | -0.734 | -0.242 | -8.807 | 141O-144N | 1 | 1 | 1 | 1 | 1 | 7.447 | 11.201 | 6.065 | -3.513 | -9.065 | 5.697 |
| 7ng6 | | | | ALD | MG-132 | P | | | 1.276 | 1.587 | -0.999 | 0.686 | -11.172 | 141O-144N | 1 | 1 | 1 | 1 | 1 | 7.420 | 11.193 | 6.350 | 0.987 | 1.661 | -0.537 | -0.177 | -11.629 | 141O-144N | 1 | 1 | 1 | 1 | 1 | 7.423 | 11.263 | 6.035 | -2.057 | -10.498 | 5.358 |
| 7nt1_B | | | | UQW | II-F10 | | 139.10* | Chain A is ligand-free. | | | | | | | | | | | | | | | 0.828 | -0.424 | -0.469 | 0.009 | 3.753 | 141O-144N | 1 | 1 | 1 | 1 | 1 | 7.718 | 11.478 | 7.578 | -5.269 | 1.449 | 5.668 |
| 7nt2_B | | | | URK | II-G11 | | 9.39* | Chain A is ligand-free. | | | | | | | | | | | | | | | 0.702 | -0.361 | -0.751 | 0.641 | 5.594 | 141O-144N | 1 | 1 | 1 | 1 | 1 | 7.656 | 11.482 | 7.585 | -5.61 | 2.08 | 5.805 |
| 7nt3_B | | | | UQZ | F3 | | >> 1000* | Chain A is ligand-free. | | | | | | | | | | | | | | | 0.841 | -0.528 | -0.406 | -0.418 | 4.253 | 141O-144N | 1 | 1 | 1 | 1 | 1 | 7.677 | 11.425 | 7.842 | -4.666 | 1.96 | 5.644 |
| 7ntv_B | | | | US8 | F2 | | 2.36* | Chain A is ligand-free. | | | | | | | | | | | | | | | 0.623 | -0.145 | -0.556 | 0.465 | 6.113 | 141O-144N | 1 | 1 | 1 | 1 | 1 | 7.770 | 11.552 | 7.595 | -4.559 | 1.334 | 5.772 |
| 7nuk_B | | | | USH | F1 | P | 3.96* | Chain A is ligand-free. | | | | | | | | | | | | | | | 0.702 | -0.11 | -0.71 | 0.361 | 4.47 | 141O-144N | 1 | 1 | 1 | 1 | 1 | 7.654 | 11.292 | 7.605 | -3.833 | 1.033 | 5.835 |
| 7nw2_A | | | | USZ | LON-WEI-adc59df6-47 | | | | 0.555 | -0.497 | -0.84 | 1.513 | 3.25 | 141O-144N | 1 | 1 | 1 | 1 | 1 | 7.652 | 11.288 | 7.067 | | | | | | | | | | | | | | | | | |
| 7p35 | | | | AG7 | Rupintrivir | P | | | 1.133 | -0.803 | -0.328 | -1.512 | -7.983 | 141O-144N | 1 | 1 | 1 | 1 | 1 | 7.672 | 11.571 | 6.325 | 0.775 | -0.627 | -1.242 | -2.066 | -4.604 | 141O-144N | 1 | 1 | 1 | 1 | 1 | 7.675 | 11.708 | 6.403 | -2.473 | -8.763 | 5.296 |

PDB data are between 10/25/2020 of Tables 1&2 and 7/25/2021. The meaning of the table is the same as in Table S1 and S2.

**Dataset S5. Summary of the ligand-free entries suppplemented to Dataset S2.**

| specirs | protein | | | | comments | chain A | | | | | | | | | | | | | | chain B | | | | | | | | | | | | | | dimer | | |
|---|---|---|---|---|---|---|---|---|---|---|---|---|---|---|---|---|---|---|---|---|---|---|---|---|---|---|---|---|---|---|---|---|---|---|---|---|
| | PDB ID | mutation | residues added to N-term | missing N-term residues | | PC1 | | | | | C-loop conformation | | | | | | | | distance 2-214 | PC1 | | | | | C-loop conformation | | | | | | | | distance 2-214 | PC1 Domain III | PC2 Domain III | distance between two 285 Cα's |
| | | | | | | C-loop | E-loop | helical loop | linker | Domain III | main-chain HB within 138-144 | HB 138-172[&] | HB 139-126[&] | HB 140-1[&] | HB 141-118[&] | HB 143-28[&] | distance 165Cα-140Cβ | distance 165Cα-141Cβ | | C-loop | E-loop | H-loop | Linker | Domain III | main-chain HB within 138-144 | HB 138-172[&] | HB 139-126[&] | HB 140-1[&] | HB 141-118[&] | HB 143-28[&] | distance 165Cα-140Cβ | distance 165Cα-141Cβ | | | | |
| SARS-CoV-2 | 7alh | | | | | 0.636 | -2.049 | -1.375 | -2.807 | -5.759 | 141O-144N | 1 | 1 | 1 | 1 | 1 | 7.680 | 11.709 | 6.348 | | | | | | | | | | | | | | | -0.001 | -8.365 | 5.311 |
| | 7ali | | | | | 0.753 | -2.213 | -1.198 | -2.865 | -3.978 | 141O-144N | 1 | 1 | 1 | 1 | 1 | 7.575 | 11.647 | 6.535 | 0.757 | -2.5 | -1.058 | -2.602 | -5.837 | 141O-144N | 1 | 1 | 1 | 1 | 1 | 7.639 | 11.711 | 6.228 | 1.462 | -8.126 | 5.371 |
| | 7ar5 | | | | Cys300 is mono-oxydized to be CSO (S-hydroxycysteine). | 0.811 | -2.061 | -1.426 | -3.421 | -6.709 | 141O-144N | 1 | 1 | 1 | 1 | 1 | 7.410 | 11.330 | 6.163 | | | | | | | | | | | | | | | 0.001 | -11.236 | 5.128 |
| | 7ar6 | | | | | 1.087 | -0.822 | -1.824 | 0.063 | -8.112 | 141O-144N | 1 | 1 | 1 | 1 | 1 | 7.725 | 11.573 | 6.056 | | | | | | | | | | | | | | | 0.001 | -11.978 | 5.224 |
| | 7avd_A | | | | Chain B binds a ligand. | 0.883 | -2.801 | -0.069 | -2.74 | -14.464 | 141O-144N | 1 | 1 | 1 | 1 | 1 | 7.373 | 11.221 | 5.982 | | | | | | | | | | | | | | | -6.19 | -9.655 | 5.526 |
| | 7baj | | | | | 0.942 | -2.449 | -1.349 | -2.915 | -4.169 | 141O-144N | 1 | 1 | 1 | 1 | 1 | 7.803 | 11.780 | 6.357 | | | | | | | | | | | | | | | 0 | -7.002 | 5.390 |
| | 7bb2 | | | | | 0.425 | 0.387 | -0.678 | 1.463 | 4.169 | 141O-144N | 1 | 1 | 1 | 1 | 1 | 7.709 | 11.547 | 7.300 | 0.569 | 0.165 | 0.549 | 2.133 | -2.63 | 141O-144N | 1 | 1 | 1 | 1 | 1 | 7.842 | 11.536 | 6.815 | 2.431 | 0.35 | 5.470 |
| | 7ca8_B | | | SG of A,B | Chain A is ligand-bound. C-terminal residues 299-306 do not exist. | | | | | | | | | | | | | | | -1.435 | -0.101 | 14.532 | 1.823 | 3.105 | | 1 | 0 | 0 | 0 | 1 | 8.494 | 12.241 | | 1.22 | 12.886 | 8.610 |
| | 7cam | | G | GSGF of A, GSG of B | | 0.059 | -0.376 | 0.895 | 1.398 | -7.559 | 141O-144N | 0 | 0 | 0 | 1 | 1 | 8.856 | 10.881 | | -0.245 | -0.991 | 0.698 | 1.895 | 1.817 | 141O-144N | 0 | 0 | 0 | 1 | 1 | 8.278 | 12.439 | | -4.699 | 3.107 | 5.443 |
| | 7djr | | | | | 0.938 | -1.533 | -1.517 | -0.712 | -5.647 | 141O-144N | 1 | 1 | 1 | 1 | 1 | 7.386 | 11.325 | 6.250 | | | | | | | | | | | | | | | 0 | -10.532 | 5.119 |
| | 7jpy | | | | Cys300 is mono-oxydized to be CSO (S-hydroxycysteine). | 0.7 | -2.417 | -0.97 | -2.571 | -5.07 | 141O-144N | 1 | 1 | 1 | 1 | 1 | 7.618 | 11.688 | 6.263 | | | | | | | | | | | | | | | 0.001 | -8.018 | 5.309 |
| | 7jst | | | | | 0.478 | 0.279 | -0.097 | 1.567 | -7.276 | 141O-144N | 1 | 1 | 1 | 1 | 1 | 7.566 | 11.629 | 6.373 | | | | | | | | | | | | | | | 0.001 | -9.11 | 5.644 |
| | 7kph | | | | | 1.023 | -1.26 | -1.82 | -0.413 | -5.845 | 141O-144N | 1 | 1 | 1 | 1 | 1 | 7.773 | 11.685 | 6.482 | | | | | | | | | | | | | | | 0 | -9.564 | 5.221 |
| | 7kvl_B | | GAM | GAM of B | | | | | | | | | | | | | | | | -10.682 | 0.978 | 2.764 | 2.033 | 13.676 | 139O-142N 140O-143N | 0 | 0 | 0 | 0 | 0 | 9.745 | 13.864 | 8.234 | -4.725 | 19.553 | 10.009 |
| | 7kvr_A | | GAM | GAM of A, GAMS of B | | -1.601 | -0.683 | 0.247 | 1.716 | 6.974 | 140O-143N | 1 | 0 | 0 | 0 | 1 | 11.843 | 11.777 | 8.794 | | | | | | | | | | | | | | | -2.043 | 13.813 | 9.542 |
| | 7l13_B | | | | Chain A is ligand-bound. | | | | | | | | | | | | | | | 0.663 | -0.788 | 1.062 | 0.677 | -9.457 | 141O-144N | 1 | 1 | 1 | 1 | 1 | 7.712 | 11.527 | 6.523 | 4.833 | -1.553 | 5.504 |
| | 7lfe_B | | GAM | GAMS of B | Chain A is ligand-bound. | | | | | | | | | | | | | | | -10.781 | 0.656 | 3.138 | 1.765 | 10.497 | 139O-142N 140O-143N | 0 | 0 | 0 | 0 | 0 | 9.799 | 14.050 | 8.583 | -2.15 | 17.03 | 9.750 |
| | 7lkd | | | | | 0.892 | -2.188 | -0.967 | -3.292 | -4.491 | 141O-144N | 1 | 1 | 1 | 1 | 1 | 7.177 | 11.307 | 6.567 | 0.969 | -2.785 | -0.625 | -2.418 | -4.213 | 141O-144N | 1 | 1 | 1 | 1 | 1 | 7.299 | 11.253 | 6.231 | 0.155 | -7.167 | 5.396 |
| | 7lke | | | | | 1.074 | -3.14 | -0.738 | -1.898 | -5.129 | 141O-144N | 1 | 1 | 1 | 0 | 1 | 7.453 | 11.530 | 6.356 | | | | | | | | | | | | | | | -0.001 | -7.963 | 5.243 |
| | 7m8m_B | | | | Chain A is ligand-bound. | | | | | | | | | | | | | | | 0.765 | -0.114 | 1.017 | -0.204 | -7.913 | 141O-144N | 1 | 1 | 1 | 1 | 1 | 7.779 | 11.348 | 6.764 | 4.404 | -2.185 | 5.293 |
| | 7m8o_B | | | | Chain A is ligand-bound. | | | | | | | | | | | | | | | 0.704 | -0.011 | 1.217 | 0.428 | -7.398 | 141O-144N | 0 | 1 | 1 | 1 | 1 | 7.997 | 11.491 | 6.997 | 4.64 | -3.14 | 5.541 |
| | 7m8p_B | | | | Chain A is ligand-bound. | | | | | | | | | | | | | | | 0.703 | -0.997 | 0.832 | 0.369 | -5.201 | 141O-144N | 1 | 1 | 1 | 1 | 1 | 7.753 | 11.618 | 6.774 | 1.054 | -0.663 | 5.795 |
| | 7mbg | | | | | 0.718 | -0.176 | 0.466 | 1.756 | 0.964 | 141O-144N | 1 | 1 | 1 | 1 | 1 | 7.697 | 11.435 | 6.699 | 0.682 | -0.094 | 10.458 | 1.979 | -3.146 | 141O-144N | 0 | 1 | 1 | 1 | 1 | 7.660 | 11.530 | 6.646 | -1.085 | -2.909 | 5.468 |
| | 7mhf | | | | 100K | 0.879 | -1.205 | -1.356 | -0.812 | -6.173 | 141O-144N | 1 | 1 | 1 | 1 | 1 | 7.672 | 11.642 | 6.248 | | | | | | | | | | | | | | | 0.002 | -10.255 | 5.086 |
| | 7mhg | | | | 240K | 0.818 | -1.878 | -1.289 | -1.365 | -4.237 | 141O-144N | 1 | 1 | 1 | 1 | 1 | 7.841 | 11.899 | 6.231 | | | | | | | | | | | | | | | 0.001 | -8.918 | 5.276 |
| | 7mhh | | | | 277K | 0.698 | -1.816 | -1.281 | -2.841 | -2.029 | 141O-144N | 1 | 1 | 1 | 1 | 1 | 7.353 | 11.274 | 6.333 | | | | | | | | | | | | | | | 0 | -5.477 | 5.760 |
| | 7mhi | | | | 298K | 0.736 | -1.974 | -1.442 | -2.767 | -2.689 | 141O-144N | 1 | 1 | 1 | 1 | 1 | 7.741 | 11.752 | 6.437 | | | | | | | | | | | | | | | 0 | -5.184 | 5.849 |
| | 7mhj | | | | 298K | 0.647 | -2.213 | -1.203 | -2.713 | -3.047 | 141O-144N | 1 | 1 | 1 | 1 | 1 | 7.412 | 11.336 | 6.478 | | | | | | | | | | | | | | | 0.001 | -5.647 | 5.772 |
| | 7mhk | | | | 310K | 0.671 | -1.889 | -1.331 | -2.135 | -2.195 | 141O-144N | 1 | 1 | 1 | 1 | 1 | 7.473 | 11.400 | 6.398 | | | | | | | | | | | | | | | 0 | -5.064 | 5.817 |
| | 7n5z | C145S | | | | 0.747 | -1.933 | -1.436 | -2.73 | -4.753 | 141O-144N | 1 | 1 | 1 | 1 | 1 | 7.753 | 11.452 | 6.439 | | | | | | | | | | | | | | | -0.002 | -6.214 | 5.228 |
| | 7nt1_A | | | | Chain B binds a ligand. | 0.854 | 0.182 | 0.757 | 1.399 | -6.322 | 141O-144N | 1 | 1 | 1 | 1 | 1 | 7.792 | 11.583 | 6.625 | | | | | | | | | | | | | | | -5.269 | 1.449 | 5.668 |
| | 7nt2_A | | | | Chain B binds a ligand. | 0.546 | -0.277 | 1.173 | 1.623 | -6.073 | 141O-144N | 1 | 1 | 1 | 1 | 1 | 7.942 | 12.021 | 6.580 | | | | | | | | | | | | | | | -5.61 | 2.08 | 5.805 |
| | 7nt3_A | | | | Chain B binds a ligand. | 0.858 | 0.31 | 0.924 | 0.243 | -6.297 | | 1 | 1 | 1 | 1 | 1 | 7.807 | 11.648 | 6.897 | | | | | | | | | | | | | | | -4.666 | 1.96 | 5.644 |
| | 7ntv_A | | | | Chain B binds a ligand. | 0.748 | 0.056 | 0.867 | 1.353 | -5.26 | 141O-144N | 1 | 1 | 1 | 1 | 1 | 7.862 | 11.879 | 6.444 | | | | | | | | | | | | | | | -4.559 | 1.334 | 5.772 |
| | 7nuk_A | | | | Chain B binds a ligand. | 0.757 | 0.346 | 0.836 | 1.444 | -4.113 | 141O-144N | 1 | 1 | 1 | 1 | 1 | 7.841 | 11.517 | 6.695 | | | | | | | | | | | | | | | -3.833 | 1.033 | 5.835 |
| | 7nw2_B | | | | Chain B binds a ligand. | | | | | | | | | | | | | | | 0.712 | 0.174 | 0.854 | 1.901 | -3.742 | 141O-144N | 1 | 1 | 1 | 1 | 1 | 7.686 | 11.324 | 6.962 | 2.171 | -0.607 | 5.577 |

**Dataset S6. Summary of ligand binding supplemented to Dataset S4.**

| species | PDBid | MC | interface | | | H-loop | | | C-loop | | | | interface | | I&II | | E-loop | | | | | I&II | Linker | | | | | Ligand interactions |
|---|---|---|---|---|---|---|---|---|---|---|---|---|---|---|---|---|---|---|---|---|---|---|---|---|---|---|---|---|
| | | BC | T25 | T26 | L27 | H41 | M49 | Y54 | F140 | L141 | N142 | G143 | S144 | C145 | H163 | H164 | M165 | E166 | L167 | P168 | H172 | D187 | R188 | Q189 | T190 | A191 | Q192 | |
| SARS-CoV | 6w79 | AB | 1 | 1 | 0 | 1 | 0 | 0 | 1 | 1 | 1 | 11 | 11 | 1 | 11 | 1 | 1 | 11 | 0 | 0 | 0 | 1 | 1 | 1 | 0 | 0 | 0 | CE |
| | 7k0g | AB | 0 | 0 | 0 | 1 | 1 | 1 | 11 | 1 | 1 | 1 | 1 | 21 | 11 | 11 | 1 | 11 | 0 | 0 | 1 | 1 | 1 | 11 | 0 | 0 | 0 | CEHL |
| | 7k0h | A | 0 | 1 | 0 | 11 | 0 | 0 | 11 | 1 | 1 | 11 | 0 | 21 | 11 | 11 | 1 | 11 | 0 | 0 | 1 | 1 | 0 | 11 | 1 | 1 | 0 | CEL |
| | 7lcp | AB | 0 | 0 | 0 | 1 | 1 | 0 | 11 | 1 | 1 | 0 | 1 | 21 | 11 | 11 | 1 | 11 | 0 | 0 | 1 | 0 | 0 | 11 | 0 | 0 | 0 | CE |
| | 7lcq | AB | 0 | 0 | 0 | 1 | 0 | 0 | 11 | 0 | 1 | 1 | 0 | 21 | 11 | 11 | 1 | 11 | 0 | 0 | 1 | 0 | 1 | 11 | 0 | 0 | 0 | CE |
| SARS-CoV-2 | 7a1u | AB | 0 | 0 | 0 | 1 | 1 | 0 | 0 | 0 | 1 | 0 | 0 | 1 | 0 | 1 | 1 | 0 | 0 | 0 | 0 | 1 | 1 | 11 | 0 | 0 | 0 | L |
| | 7abu[x] | AB | 0 | 0 | 0 | 0 | 0 | 0 | 0 | 1[#] | 11[#] | 0 | 0 | 0 | 0 | 0 | 0 | 0 | 0 | 0 | 0 | 0 | 0 | 0 | 0 | 0 | 0 | |
| | 7adw | AB | 0 | 0 | 0 | 0 | 0 | 0 | 1 | 1 | 1 | 0 | 0 | 20 | 11 | 1 | 0 | 1 | 0 | 0 | 0 | 0 | 0 | 0 | 0 | 0 | 0 | |
| | 7aeg | AB | 0 | 0 | 0 | 1 | 0 | 0 | 1 | 1 | 0 | 11 | 1 | 21 | 11 | 11 | 1 | 11 | 0 | 1 | 0 | 0 | 1 | 1 | 1 | 0 | 1 | CEL |
| | 7aeh | AB | 1 | 1 | 0 | 11 | 0 | 0 | 1 | 1 | 1 | 11 | 11 | 21 | 11 | 11 | 1 | 1 | 0 | 0 | 0 | 0 | 0 | 0 | 0 | 0 | 0 | CE |
| | 7af0[x] | AB | 0 | 0 | 0 | 0 | 0 | 0 | 0 | 0 | 0 | 0 | 0 | 0 | 0 | 0 | 0 | 0 | 0 | 0 | 0 | 0 | 0 | 0 | 0 | 0 | 0 | |
| | 7aga[x] | AB | 0 | 0 | 0 | 0 | 0 | 0 | 0 | 0 | 0 | 0 | 0 | 0 | 0 | 0 | 0 | 0 | 0 | 0 | 0 | 0 | 0 | 0 | 0 | 0 | 0 | |
| | 7aha | AB | 0 | 0 | 0 | 1 | 0 | 0 | 0 | 0 | 11 | 11 | 0 | 21 | 0 | 1 | 0 | 0 | 0 | 0 | 0 | 0 | 0 | 0 | 0 | 0 | 0 | C |
| | 7ak4 | AB | 0 | 0 | 0 | 0 | 1 | 0 | 0 | 0 | 0 | 0 | 0 | 0 | 0 | 0 | 21 | 11 | 0 | 1 | 0 | 1 | 1 | 1 | 10 | 0 | 11 | EL |
| | 7amj[x] | AB | 0 | 0 | 0 | 0 | 0 | 0 | 0 | 0 | 0 | 0 | 0 | 0 | 0 | 0 | 0 | 0 | 0 | 0 | 0 | 0 | 0 | 0 | 0 | 0 | 0 | |
| | 7ans | AB | 0 | 0 | 0 | 1 | 1 | 0 | 0 | 0 | 1 | 1 | 0 | 11 | 0 | 1 | 1 | 0 | 0 | 0 | 0 | 1 | 1 | 1 | 0 | 0 | 0 | |
| | 7aol[x] | AB | 0 | 0 | 0 | 0 | 0 | 0 | 0 | 0 | 0 | 0 | 0 | 0 | 0 | 0 | 0 | 0 | 0 | 0 | 0 | 0 | 0 | 0 | 0 | 0 | 0 | |
| | 7ap6 | AB | 0 | 0 | 0 | 1 | 1 | 1 | 11 | 1 | 0 | 0 | 0 | 1 | 11 | 1 | 1 | 1 | 0 | 0 | 0 | 1 | 0 | 0 | 0 | 0 | 0 | H |
| | 7aph[x] | AB | 0 | 0 | 0 | 0 | 0 | 0 | 0 | 0 | 0 | 0 | 0 | 0 | 0 | 0 | 0 | 0 | 0 | 0 | 0 | 0 | 0 | 0 | 0 | 0 | 0 | |
| | 7aqe | AB | 1 | 0 | 0 | 1 | 1 | 0 | 0 | 0 | 1 | 0 | 0 | 0 | 0 | 0 | 0 | 11 | 1 | 1 | 0 | 0 | 0 | 11 | 0 | 0 | 0 | |
| | 7aqi[x] | AB | 0 | 0 | 0 | 0 | 0 | 0 | 0 | 0 | 0 | 0 | 0 | 0 | 0 | 0 | 0 | 0 | 0 | 0 | 0 | 0 | 0 | 0 | 0 | 0 | 0 | |
| | 7aqj | AB | 0 | 0 | 0 | 0 | 1 | 0 | 0 | 0 | 1 | 1 | 0 | 21 | 0 | 1 | 1 | 0 | 0 | 0 | 0 | 0 | 0 | 0 | 0 | 0 | 0 | |
| | 7arf[x] | AB | 0 | 0 | 0 | 0 | 0 | 0 | 0 | 0 | 0 | 0 | 0 | 0 | 0 | 0 | 0 | 0 | 0 | 0 | 0 | 0 | 0 | 0 | 0 | 0 | 0 | |
| | 7avd | | | | | | | | | | | | | | | | | | | | | | | | | | | |
| | 7awr[x] | AB | 0 | 0 | 0 | 0 | 0 | 0 | 0 | 0 | 0 | 0 | 0 | 0 | 0 | 0 | 0 | 0 | 0 | 0 | 0 | 0 | 0 | 0 | 0 | 0 | 0 | |
| | 7aws | AB | 0 | 0 | 0 | 11 | 1 | 0 | 0 | 1 | 11 | 0 | 0 | 21 | 1 | 1 | 11 | 0 | 0 | 0 | 0 | 1 | 1 | 1 | 0 | 0 | 0 | CH |
| | 7awu | AB | 0 | 0 | 1 | 1 | 0 | 0 | 0 | 0 | 0 | 1 | 1 | 11 | 1 | 1 | 0 | 0 | 0 | 0 | 0 | 0 | 0 | 0 | 0 | 0 | 0 | |
| | 7aww[x] | AB | 0 | 0 | 0 | 0 | 0 | 0 | 0 | 0 | 0 | 0 | 0 | 0 | 0 | 0 | 0 | 0 | 0 | 0 | 0 | 0 | 0 | 0 | 0 | 0 | 0 | |
| | 7ax6[x] | AB | 0 | 0 | 0 | 0 | 0 | 0 | 0 | 0 | 0 | 0 | 0 | 0 | 0 | 0 | 0 | 0 | 0 | 0 | 0 | 0 | 0 | 0 | 0 | 0 | 0 | |
| | 7axm[x] | AB | 0 | 0 | 0 | 0 | 0 | 0 | 0 | 1[#] | 1[#] | 0 | 0 | 0 | 0 | 0 | 0 | 0 | 0 | 0 | 0 | 0 | 0 | 0 | 0 | 0 | 0 | |
| | 7axo[x] | AB | 0 | 0 | 0 | 0 | 0 | 0 | 0 | 0 | 0 | 0 | 0 | 0 | 0 | 0 | 0 | 0 | 0 | 0 | 0 | 0 | 0 | 0 | 0 | 0 | 0 | |
| | 7ay7 | AB | 1 | 0 | 0 | 1 | 1 | 0 | 1 | 1 | 1 | 0 | 1 | 21 | 1 | 0 | 0 | 1 | 0 | 0 | 0 | 0 | 0 | 0 | 0 | 0 | 0 | C |
| | 7b3e | A | 1 | 0 | 0 | 1 | 1 | 0 | 0 | 0 | 0 | 1 | 0 | 21 | 0 | 1 | 1 | 10 | 0 | 0 | 0 | 1 | 1 | 1 | 0 | 0 | 0 | |
| | 7bak | AB | 0 | 0 | 0 | 10 | 0 | 0 | 0 | 0 | 0 | 0 | 0 | 20 | 0 | 0 | 0 | 0 | 0 | 0 | 0 | 0 | 0 | 0 | 0 | 0 | 0 | |
| | 7bal | AB | 0 | 0 | 0 | 10 | 0 | 0 | 0 | 0 | 0 | 0 | 0 | 20 | 0 | 0 | 0 | 0 | 0 | 0 | 0 | 0 | 0 | 0 | 0 | 0 | 0 | |
| | 7be7 | A | 0 | 0 | 0 | 1 | 0 | 0 | 0 | 0 | 1 | 1 | 1 | 21 | 1 | 11 | 1 | 11 | 0 | 1 | 0 | 0 | 0 | 11 | 1 | 1 | 1 | EL |
| | 7bfb[x] | A | 0 | 0 | 0 | 0 | 1 | 0 | 0 | 0 | 0 | 1 | 0 | 21 | 0 | 0 | 0 | 0 | 0 | 0 | 0 | 1 | 0 | 1 | 0 | 0 | 0 | |
| | 7bgp | A | 0 | 0 | 0 | 1 | 0 | 0 | 0 | 0 | 1 | 1 | 0 | 21 | 1 | 11 | 1 | 11 | 0 | 1 | 0 | 0 | 0 | 11 | 1 | 1 | 1 | EL |
| | 7ca8 | A | 0 | 0 | 0 | 1 | 0 | 0 | 0 | 0 | 0 | 0 | 0 | 1 | 0 | 1 | 1 | 0 | 0 | 0 | 0 | 1 | 1 | 1 | 1 | 0 | 1 | L |
| | 7cb7 | A | 0 | 0 | 0 | 1 | 1 | 0 | 11 | 1 | 1 | 1 | 1 | 21 | 11 | 11 | 1 | 11 | 0 | 0 | 1 | 1 | 0 | 11 | 0 | 0 | 0 | CE |
| | 7cut | AB | 0 | 0 | 0 | 1 | 1 | 0 | 1 | 0 | 1 | 11 | 1 | 21 | 1 | 1 | 1 | 11 | 0 | 0 | 0 | 1 | 1 | 1 | 0 | 0 | 0 | C |
| | 7cuu | A | 0 | 0 | 0 | 1 | 1 | 0 | 1 | 0 | 1 | 1 | 0 | 21 | 1 | 11 | 1 | 11 | 0 | 1 | 0 | 1 | 0 | 11 | 1 | 1 | 1 | EL |
| | 7dk1 | A | 0 | 0 | 0 | 20 | 0 | 0 | 0 | 0 | 0 | 0 | 0 | 20 | 0 | 0 | 0 | 0 | 0 | 0 | 0 | 0 | 0 | 0 | 0 | 0 | 0 | |
| | 7dpp | AB | 1 | 1 | 1 | 1 | 0 | 0 | 0 | 0 | 0 | 1 | 0 | 21 | 0 | 1 | 1 | 0 | 0 | 0 | 0 | 1 | 0 | 0 | 0 | 0 | 0 | R |
| | 7dpu | A | 1 | 1 | 1 | 1 | 1 | 1 | 0 | 0 | 0 | 1 | 0 | 21 | 0 | 0 | 1 | 11 | 0 | 0 | 0 | 1 | 1 | 1 | 0 | 0 | 0 | RH |
| | 7dpv | A | 1 | 0 | 0 | 1 | 0 | 1 | 0 | 0 | 0 | 11 | 0 | 21 | 0 | 1 | 1 | 10 | 0 | 0 | 0 | 1 | 1 | 1 | 0 | 0 | 0 | |
| | 7dpv | B | 1 | 0 | 0 | 1 | 1 | 1 | 0 | 0 | 0 | 11 | 0 | 21 | 0 | 1 | 1 | 10 | 0 | 0 | 0 | 1 | 1 | 1 | 0 | 0 | 0 | H |
| | 7e18 | AB | 1 | 0 | 1 | 1 | 1 | 0 | 11 | 1 | 1 | 0 | 1 | 21 | 11 | 11 | 1 | 11 | 0 | 1 | 1 | 0 | 1 | 11 | 1 | 1 | 0 | CEL |
| | 7e19 | AB | 1 | 0 | 0 | 1 | 1 | 0 | 11 | 1 | 1 | 1 | 1 | 21 | 11 | 11 | 1 | 11 | 0 | 0 | 1 | 0 | 0 | 1 | 1 | 1 | 1 | CEL |
| | 7ein | A | 0 | 0 | 0 | 1 | 1 | 1 | 0 | 0 | 1 | 1 | 0 | 21 | 0 | 11 | 1 | 11 | 0 | 0 | 0 | 1 | 0 | 1 | 1 | 0 | 0 | EH |
| | 7fay | AB | 0 | 0 | 0 | 1 | 1 | 0 | 1 | 1 | 1 | 11 | 0 | 21 | 11 | 1 | 1 | 11 | 0 | 0 | 0 | 0 | 0 | 1 | 0 | 0 | 0 | CE |
| | 7faz | A | 0 | 0 | 0 | 1 | 1 | 1 | 1 | 1 | 1 | 11 | 11 | 21 | 11 | 1 | 1 | 11 | 0 | 0 | 0 | 1 | 1 | 1 | 0 | 0 | 0 | CEH |
| | 7jpz | AB | 0 | 0 | 0 | 1 | 1 | 1 | 11 | 1 | 1 | 11 | 1 | 21 | 11 | 1 | 1 | 11 | 1 | 1 | 1 | 1 | 1 | 1 | 0 | 0 | 0 | CEH |
| | 7jq0 | AB | 0 | 0 | 0 | 1 | 1 | 0 | 11 | 0 | 1 | 1 | 1 | 21 | 11 | 11 | 1 | 11 | 0 | 1 | 1 | 1 | 0 | 1 | 1 | 1 | 0 | CEL |
| | 7jq1 | AB | 0 | 0 | 0 | 1 | 1 | 1 | 11 | 1 | 1 | 11 | 1 | 21 | 11 | 11 | 1 | 11 | 0 | 1 | 1 | 1 | 1 | 1 | 1 | 1 | 1 | CEHL |

| PDBid | MC | interface | | | H-loop | | | C-loop | | | | interface | | I&II | | E-loop | | | | | I&II | Linker | | | | | Ligand interactions |
|---|---|---|---|---|---|---|---|---|---|---|---|---|---|---|---|---|---|---|---|---|---|---|---|---|---|---|---|
| | BC | T25 | T26 | L27 | H41 | M49 | Y54 | F140 | L141 | N142 | G143 | S144 | C145 | H163 | H164 | M165 | E166 | L167 | P168 | H172 | D187 | R188 | Q189 | T190 | A191 | Q192 | |
| 6w79 | | | | | | | | | | | | | | | | | | | | | | | | | | | |
| 7k0g | | | | | | | | | | | | | | | | | | | | | | | | | | | |
| 7k0h | B | 0 | 1 | 0 | 11 | 0 | 0 | 11 | 1 | 1 | 11 | 0 | 11 | 11 | 11 | 1 | 11 | 0 | 0 | 1 | 1 | 0 | 11 | 0 | 0 | 0 | CE |
| 7lcp | | | | | | | | | | | | | | | | | | | | | | | | | | | |
| 7lcq | | | | | | | | | | | | | | | | | | | | | | | | | | | |
| 7a1u | | | | | | | | | | | | | | | | | | | | | | | | | | | |
| 7abu[x] | | | | | | | | | | | | | | | | | | | | | | | | | | | |
| 7adw | | | | | | | | | | | | | | | | | | | | | | | | | | | |
| 7aeg | | | | | | | | | | | | | | | | | | | | | | | | | | | |
| 7aeh | | | | | | | | | | | | | | | | | | | | | | | | | | | |
| 7af0[x] | | | | | | | | | | | | | | | | | | | | | | | | | | | |
| 7aga[x] | | | | | | | | | | | | | | | | | | | | | | | | | | | |
| 7aha | | | | | | | | | | | | | | | | | | | | | | | | | | | |
| 7ak4 | | | | | | | | | | | | | | | | | | | | | | | | | | | |
| 7amj[x] | | | | | | | | | | | | | | | | | | | | | | | | | | | |
| 7ans | | | | | | | | | | | | | | | | | | | | | | | | | | | |
| 7aol[x] | | | | | | | | | | | | | | | | | | | | | | | | | | | |
| 7ap6 | | | | | | | | | | | | | | | | | | | | | | | | | | | |
| 7aph[x] | | | | | | | | | | | | | | | | | | | | | | | | | | | |
| 7aqe | | | | | | | | | | | | | | | | | | | | | | | | | | | |
| 7aqi[x] | | | | | | | | | | | | | | | | | | | | | | | | | | | |
| 7aqj | | | | | | | | | | | | | | | | | | | | | | | | | | | |
| 7arf[x] | | | | | | | | | | | | | | | | | | | | | | | | | | | |
| 7avd | B | 0 | 0 | 0 | 1 | 11 | 1 | 0 | 0 | 0 | 0 | 0 | 0 | 0 | 1 | 0 | 0 | 0 | 0 | 0 | 1 | 1 | 1 | 0 | 0 | 0 | H |
| 7awr[x] | | | | | | | | | | | | | | | | | | | | | | | | | | | |
| 7aws | | | | | | | | | | | | | | | | | | | | | | | | | | | |
| 7awu | | | | | | | | | | | | | | | | | | | | | | | | | | | |
| 7aww[x] | | | | | | | | | | | | | | | | | | | | | | | | | | | |
| 7ax6[x] | | | | | | | | | | | | | | | | | | | | | | | | | | | |
| 7axm[x] | | | | | | | | | | | | | | | | | | | | | | | | | | | |
| 7axo[x] | | | | | | | | | | | | | | | | | | | | | | | | | | | |
| 7ay7 | | | | | | | | | | | | | | | | | | | | | | | | | | | |
| 7b3e | B | 1 | 1 | 1 | 1 | 0 | 0 | 0 | 0 | 0 | 1 | 0 | 21 | 0 | 1 | 1 | 10 | 0 | 0 | 0 | 1 | 1 | 0 | 0 | 0 | 0 | R |
| 7bak | | | | | | | | | | | | | | | | | | | | | | | | | | | |
| 7bal | | | | | | | | | | | | | | | | | | | | | | | | | | | |
| 7be7 | B | 0 | 0 | 0 | 1 | 1 | 0 | 0 | 0 | 1 | 1 | 0 | 21 | 1 | 11 | 1 | 11 | 0 | 1 | 0 | 0 | 0 | 11 | 1 | 1 | 1 | EL |
| 7bfb[x] | B | 0 | 0 | 0 | 0 | 0 | 0 | 0 | 0 | 0 | 0 | 0 | 0 | 0 | 0 | 0 | 0 | 0 | 0 | 0 | 0 | 0 | 0 | 0 | 0 | 0 | |
| 7bgp | B | 0 | 0 | 0 | 1 | 0 | 0 | 0 | 0 | 1 | 1 | 0 | 21 | 1 | 11 | 1 | 11 | 0 | 0 | 0 | 0 | 0 | 11 | 1 | 1 | 1 | EL |
| 7ca8 | | | | | | | | | | | | | | | | | | | | | | | | | | | |
| 7cb7 | B | 0 | 0 | 0 | 11 | 1 | 0 | 11 | 0 | 1 | 0 | 1 | 21 | 11 | 11 | 1 | 11 | 0 | 0 | 1 | 0 | 0 | 11 | 0 | 0 | 0 | CEH |
| 7cut | | | | | | | | | | | | | | | | | | | | | | | | | | | |
| 7cuu | B | 0 | 0 | 0 | 1 | 1 | 0 | 1 | 1 | 1 | 11 | 0 | 21 | 1 | 11 | 1 | 11 | 1 | 1 | 0 | 1 | 0 | 11 | 1 | 1 | 1 | CEL |
| 7dk1 | B | 0 | 0 | 0 | 20 | 0 | 0 | 0 | 0 | 0 | 0 | 0 | 20 | 0 | 0 | 0 | 0 | 0 | 0 | 0 | 0 | 0 | 0 | 0 | 0 | 0 | |
| 7dpp | | | | | | | | | | | | | | | | | | | | | | | | | | | |
| 7dpu | B | 1 | 0 | 0 | 1 | 1 | 1 | 0 | 0 | 0 | 1 | 0 | 21 | 0 | 1 | 1 | 11 | 0 | 0 | 0 | 1 | 1 | 1 | 0 | 0 | 0 | H |
| 7dpv | C | 1 | 1 | 0 | 1 | 0 | 1 | 0 | 0 | 0 | 1 | 0 | 21 | 0 | 1 | 1 | 0 | 0 | 0 | 0 | 1 | 1 | 1 | 0 | 0 | 0 | |
| 7dpv | D | 1 | 1 | 0 | 1 | 0 | 1 | 0 | 0 | 0 | 1 | 0 | 21 | 0 | 1 | 1 | 10 | 0 | 0 | 0 | 1 | 1 | 1 | 0 | 0 | 0 | |
| 7e18 | | | | | | | | | | | | | | | | | | | | | | | | | | | |
| 7e19 | | | | | | | | | | | | | | | | | | | | | | | | | | | |
| 7ein | B | 0 | 0 | 0 | 1 | 1 | 1 | 0 | 0 | 0 | 1 | 0 | 21 | 0 | 11 | 1 | 11 | 0 | 0 | 0 | 1 | 0 | 1 | 1 | 0 | 0 | EH |
| 7fay | | | | | | | | | | | | | | | | | | | | | | | | | | | |
| 7faz | B | 0 | 0 | 0 | 1 | 1 | 1 | 1 | 1 | 1 | 11 | 11 | 21 | 11 | 1 | 1 | 11 | 0 | 0 | 0 | 1 | 1 | 0 | 0 | 0 | 0 | CEH |
| 7jpz | | | | | | | | | | | | | | | | | | | | | | | | | | | |
| 7jq0 | | | | | | | | | | | | | | | | | | | | | | | | | | | |
| 7jq1 | | | | | | | | | | | | | | | | | | | | | | | | | | | |

| | | | | | | | | | | | | | | | | | | | | | | | | | | | | | | | | | | | | | | | | | | | | | | | | | | | | | | |
|---|---|---|---|---|---|---|---|---|---|---|---|---|---|---|---|---|---|---|---|---|---|---|---|---|---|---|---|---|---|---|---|---|---|---|---|---|---|---|---|---|---|---|---|---|---|---|---|---|---|---|---|---|---|---|
| 7jq2 | AB | 0 | 0 | 0 | 1 | 1 | 1 | 11 | 1 | 1 | 11 | 0 | 21 | 11 | 10 | 1 | 11 | 0 | 1 | 1 | 1 | 1 | 1 | 1 | 1 | 1 | CEHL | | | | | | | | | | | | | | | | | | | | | | | | | | | |
| 7jq3 | AB | 0 | 0 | 0 | 1 | 0 | 1 | 11 | 1 | 1 | 11 | 0 | 21 | 11 | 11 | 1 | 11 | 0 | 1 | 0 | 1 | 1 | 1 | 1 | 1 | 1 | CEL | | | | | | | | | | | | | | | | | | | | | | | | | | | |
| 7jq4 | AB | 0 | 0 | 0 | 1 | 1 | 1 | 11 | 1 | 1 | 11 | 1 | 21 | 11 | 11 | 1 | 11 | 1 | 1 | 1 | 1 | 1 | 1 | 1 | 1 | 1 | CEHL | | | | | | | | | | | | | | | | | | | | | | | | | | | |
| 7jq5 | AB | 0 | 0 | 0 | 1 | 1 | 1 | 11 | 1 | 1 | 11 | 0 | 21 | 11 | 11 | 1 | 11 | 0 | 1 | 1 | 1 | 1 | 1 | 1 | 1 | 0 | CEHL | | | | | | | | | | | | | | | | | | | | | | | | | | | |
| 7jsu | AB | 0 | 0 | 0 | 1 | 0 | 0 | 11 | 1 | 1 | 1 | 0 | 21 | 11 | 11 | 1 | 11 | 0 | 0 | 1 | 1 | 1 | 11 | 0 | 0 | 0 | CEL | | | | | | | | | | | | | | | | | | | | | | | | | | | |
| 7jt0 | AB | 0 | 0 | 0 | 1 | 0 | 0 | 0 | 0 | 0 | 0 | 0 | 21 | 0 | 1 | 1 | 0 | 0 | 0 | 0 | 0 | 0 | 0 | 0 | 0 | 0 | | | | | | | | | | | | | | | | | | | | | | | | | | | | |
| 7jt7 | AB | 0 | 0 | 0 | 1 | 0 | 0 | 11 | 1 | 1 | 11 | 0 | 21 | 11 | 11 | 1 | 11 | 1 | 1 | 1 | 1 | 0 | 11 | 1 | 1 | 1 | CEL | | | | | | | | | | | | | | | | | | | | | | | | | | | |
| 7jw8 | A | 0 | 0 | 0 | 1 | 0 | 0 | 11 | 0 | 1 | 10 | 1 | 21 | 11 | 11 | 1 | 11 | 0 | 1 | 1 | 0 | 0 | 11 | 1 | 1 | 1 | CEL | B | 0 | 1 | 0 | 1 | 1 | 0 | 11 | 0 | 1 | 11 | 0 | 21 | 11 | 11 | 1 | 11 | 0 | 1 | 1 | 0 | 0 | 11 | 1 | 0 | 1 | CEL |
| 7jw8 | C | 1 | 1 | 1 | 1 | 0 | 0 | 11 | 1 | 1 | 1 | 0 | 21 | 11 | 11 | 1 | 11 | 0 | 1 | 1 | 0 | 0 | 11 | 1 | 0 | 1 | RCEL | D | 0 | 0 | 0 | 1 | 1 | 0 | 11 | 0 | 1 | 10 | 0 | 21 | 11 | 11 | 1 | 11 | 0 | 1 | 1 | 0 | 0 | 11 | 1 | 1 | 1 | CEL |
| 7k0e | A | 0 | 0 | 0 | 1 | 1 | 0 | 11 | 1 | 1 | 1 | 0 | 21 | 11 | 11 | 1 | 11 | 0 | 0 | 1 | 1 | 0 | 11 | 0 | 0 | 0 | CE | B | 0 | 0 | 0 | 1 | 0 | 0 | 11 | 1 | 1 | 1 | 0 | 21 | 11 | 11 | 1 | 11 | 0 | 0 | 1 | 1 | 0 | 11 | 0 | 0 | 0 | CE |
| 7k0f | A | 0 | 1 | 0 | 11 | 0 | 0 | 11 | 1 | 1 | 11 | 0 | 21 | 11 | 11 | 1 | 11 | 0 | 0 | 1 | 1 | 0 | 11 | 1 | 1 | 0 | CEL | B | 0 | 1 | 0 | 11 | 0 | 0 | 11 | 0 | 1 | 11 | 0 | 21 | 11 | 11 | 1 | 11 | 0 | 0 | 1 | 1 | 0 | 11 | 0 | 0 | 0 | CE |
| 7kvl[x] | A | 0 | 0 | 0 | 0 | 0 | 0 | 0 | 0 | 0 | 0 | 0 | 0 | 0 | 0 | 0 | 0 | 0 | 0 | 0 | 0 | 0 | 0 | 0 | 0 | 0 | | | | | | | | | | | | | | | | | | | | | | | | | | | | |
| 7kvr[x] | | | | | | | | | | | | | | | | | | | | | | | | | | | | B | 0 | 0 | 0 | 0 | 0 | 0 | 0 | 0 | 0 | 0 | 0 | 0 | 0 | 0 | 0 | 0 | 0 | 0 | 0 | 0 | 0 | 0 | 0 | 0 | 0 | |
| 7kx5 | AB | 1 | 1 | 1 | 1 | 1 | 1 | 1 | 1 | 1 | 11 | 11 | 1 | 11 | 1 | 1 | 11 | 0 | 0 | 0 | 1 | 1 | 1 | 0 | 0 | 0 | RCEH | | | | | | | | | | | | | | | | | | | | | | | | | | | |
| 7kyu | AB | 0 | 0 | 0 | 1 | 0 | 0 | 0 | 0 | 0 | 0 | 0 | 21 | 0 | 1 | 1 | 0 | 0 | 0 | 0 | 1 | 1 | 0 | 0 | 0 | 0 | | | | | | | | | | | | | | | | | | | | | | | | | | | | |
| 7l0d | AB | 1 | 1 | 1 | 1 | 1 | 0 | 1 | 1 | 1 | 11 | 1 | 1 | 11 | 1 | 0 | 1 | 0 | 0 | 0 | 1 | 1 | 1 | 0 | 0 | 0 | RC | | | | | | | | | | | | | | | | | | | | | | | | | | | |
| 7l10 | AB | 1 | 1 | 0 | 1 | 1 | 0 | 1 | 1 | 1 | 1 | 0 | 1 | 11 | 1 | 1 | 11 | 0 | 0 | 0 | 0 | 1 | 0 | 0 | 0 | 0 | CE | | | | | | | | | | | | | | | | | | | | | | | | | | | |
| 7l11 | A | 1 | 1 | 0 | 1 | 1 | 0 | 1 | 1 | 1 | 1 | 0 | 1 | 11 | 1 | 1 | 11 | 1 | 0 | 0 | 0 | 1 | 1 | 0 | 0 | 1 | CE | B | 1 | 1 | 0 | 1 | 1 | 0 | 1 | 1 | 1 | 1 | 11 | 1 | 11 | 1 | 1 | 11 | 1 | 0 | 0 | 0 | 1 | 1 | 1 | 0 | 0 | CE |
| 7l12 | AB | 1 | 11 | 0 | 1 | 1 | 0 | 1 | 1 | 1 | 11 | 0 | 11 | 11 | 1 | 1 | 11 | 0 | 1 | 0 | 0 | 1 | 1 | 1 | 0 | 0 | RCE | | | | | | | | | | | | | | | | | | | | | | | | | | | |
| 7l13 | A | 1 | 10 | 0 | 1 | 1 | 0 | 1 | 1 | 1 | 11 | 0 | 11 | 11 | 0 | 1 | 11 | 0 | 1 | 0 | 0 | 0 | 1 | 1 | 0 | 0 | RCE | | | | | | | | | | | | | | | | | | | | | | | | | | | |
| 7l14 | A | 1 | 1 | 0 | 1 | 1 | 0 | 1 | 1 | 1 | 1 | 0 | 1 | 11 | 1 | 1 | 11 | 1 | 0 | 0 | 0 | 1 | 1 | 1 | 0 | 1 | CEL | B | 1 | 1 | 0 | 0 | 1 | 0 | 1 | 1 | 1 | 1 | 1 | 1 | 11 | 1 | 1 | 11 | 0 | 1 | 0 | 0 | 1 | 1 | 1 | 0 | 1 | CEL |
| 7l5d | AB | 1 | 0 | 0 | 11 | 1 | 0 | 1 | 1 | 1 | 0 | 0 | 10 | 11 | 11 | 1 | 1 | 0 | 0 | 0 | 0 | 0 | 0 | 0 | 0 | 0 | EH | | | | | | | | | | | | | | | | | | | | | | | | | | | |
| 7lb7 | AB | 1 | 1 | 0 | 11 | 0 | 0 | 0 | 0 | 1 | 11 | 1 | 21 | 0 | 11 | 1 | 11 | 0 | 1 | 0 | 1 | 1 | 11 | 1 | 1 | 1 | CEL | | | | | | | | | | | | | | | | | | | | | | | | | | | |
| 7lbn | AB | 0 | 0 | 0 | 1 | 0 | 0 | 1 | 1 | 1 | 11 | 0 | 21 | 0 | 11 | 1 | 11 | 0 | 0 | 0 | 1 | 0 | 0 | 0 | 0 | 0 | CE | | | | | | | | | | | | | | | | | | | | | | | | | | | |
| 7lco | AB | 0 | 0 | 0 | 1 | 1 | 0 | 11 | 1 | 1 | 1 | 0 | 21 | 11 | 11 | 1 | 11 | 0 | 1 | 1 | 1 | 1 | 0 | 1 | 0 | 0 | CE | | | | | | | | | | | | | | | | | | | | | | | | | | | |
| 7lcs | AB | 0 | 0 | 0 | 1 | 1 | 1 | 11 | 1 | 1 | 1 | 0 | 21 | 11 | 11 | 1 | 11 | 0 | 0 | 1 | 1 | 1 | 11 | 0 | 0 | 0 | CEHL | | | | | | | | | | | | | | | | | | | | | | | | | | | |
| 7lct | AB | 0 | 0 | 0 | 1 | 1 | 0 | 11 | 1 | 1 | 1 | 1 | 21 | 11 | 11 | 1 | 11 | 0 | 0 | 1 | 1 | 1 | 1 | 1 | 1 | 0 | CEL | | | | | | | | | | | | | | | | | | | | | | | | | | | |
| 7ldl | A | 0 | 0 | 0 | 1 | 1 | 0 | 11 | 1 | 1 | 1 | 0 | 21 | 11 | 11 | 1 | 11 | 0 | 0 | 1 | 1 | 1 | 11 | 1 | 1 | 0 | CEL | B | 0 | 0 | 0 | 1 | 1 | 0 | 11 | 0 | 1 | 1 | 1 | 21 | 11 | 11 | 1 | 11 | 0 | 0 | 1 | 1 | 1 | 1 | 1 | 1 | 0 | CEL |
| 7ldx[x] | AB | 0 | 0 | 0 | 0 | 0 | 0 | 0 | 0 | 0 | 0 | 0 | 0 | 0 | 0 | 0 | 0 | 0 | 0 | 0 | 0 | 0 | 0 | 0 | 0 | 0 | | | | | | | | | | | | | | | | | | | | | | | | | | | | |
| 7lfe | A | 0 | 0 | 0 | 11 | 1 | 0 | 0 | 0 | 0 | 0 | 0 | 11 | 0 | 1 | 1 | 11 | 0 | 0 | 0 | 1 | 0 | 1 | 0 | 0 | 0 | H | | | | | | | | | | | | | | | | | | | | | | | | | | | |
| 7lkr | A | 0 | 0 | 0 | 11 | 0 | 0 | 11 | 1 | 1 | 0 | 0 | 21 | 11 | 11 | 1 | 11 | 0 | 0 | 1 | 1 | 0 | 11 | 0 | 0 | 0 | CE | B | 0 | 0 | 0 | 11 | 0 | 1 | 11 | 1 | 1 | 0 | 0 | 21 | 11 | 11 | 1 | 11 | 0 | 0 | 1 | 1 | 1 | 11 | 1 | 1 | 0 | CEHL |
| 7lks | A | 0 | 0 | 0 | 1 | 0 | 0 | 11 | 1 | 1 | 11 | 0 | 21 | 11 | 11 | 1 | 11 | 1 | 1 | 1 | 1 | 0 | 11 | 0 | 0 | 0 | CE | B | 0 | 0 | 0 | 1 | 0 | 0 | 0 | 0 | 1 | 1 | 1 | 21 | 11 | 11 | 1 | 11 | 0 | 0 | 1 | 1 | 0 | 11 | 1 | 0 | 0 | EL |
| 7lkt | A | 0 | 0 | 0 | 1 | 1 | 0 | 11 | 1 | 1 | 1 | 0 | 21 | 11 | 11 | 1 | 11 | 0 | 1 | 1 | 1 | 0 | 11 | 1 | 1 | 0 | CEL | B | 0 | 0 | 0 | 1 | 0 | 0 | 10 | 0 | 1 | 1 | 1 | 21 | 11 | 11 | 1 | 11 | 0 | 0 | 1 | 1 | 0 | 11 | 1 | 1 | 0 | CEL |
| 7lku | A | 0 | 0 | 0 | 11 | 0 | 0 | 11 | 1 | 1 | 0 | 0 | 21 | 11 | 11 | 1 | 11 | 0 | 0 | 1 | 1 | 0 | 11 | 0 | 0 | 0 | CE | B | 0 | 0 | 0 | 11 | 1 | 0 | 11 | 1 | 1 | 0 | 1 | 21 | 11 | 11 | 1 | 11 | 0 | 0 | 1 | 1 | 1 | 11 | 0 | 0 | 0 | CEHL |
| 7lkv | A | 0 | 0 | 0 | 11 | 1 | 0 | 11 | 0 | 1 | 0 | 1 | 21 | 11 | 11 | 1 | 11 | 0 | 0 | 1 | 1 | 0 | 11 | 0 | 0 | 0 | CEH | B | 0 | 0 | 0 | 11 | 0 | 0 | 11 | 0 | 1 | 0 | 1 | 21 | 11 | 11 | 1 | 11 | 1 | 1 | 1 | 1 | 0 | 11 | 0 | 0 | 0 | CE |
| 7lkw | A | 0 | 0 | 0 | 10 | 0 | 0 | 11 | 1 | 1 | 0 | 0 | 21 | 11 | 11 | 1 | 11 | 1 | 1 | 1 | 1 | 1 | 11 | 0 | 0 | 0 | CEL | B | 0 | 0 | 0 | 11 | 1 | 0 | 11 | 0 | 1 | 0 | 1 | 21 | 11 | 11 | 1 | 11 | 0 | 1 | 1 | 1 | 0 | 11 | 0 | 0 | 0 | CEH |
| 7lkx | A | 0 | 0 | 0 | 11 | 0 | 0 | 11 | 0 | 1 | 0 | 0 | 21 | 11 | 11 | 1 | 11 | 1 | 1 | 1 | 1 | 0 | 11 | 0 | 0 | 0 | CE | B | 0 | 0 | 0 | 11 | 1 | 0 | 1 | 0 | 1 | 0 | 0 | 21 | 11 | 11 | 1 | 11 | 0 | 0 | 1 | 1 | 0 | 11 | 0 | 0 | 0 | EH |
| 7lmc | A | 0 | 0 | 0 | 0 | 1 | 0 | 1 | 1 | 1 | 1 | 1 | 1 | 11 | 1 | 1 | 11 | 1 | 1 | 0 | 0 | 1 | 11 | 0 | 1 | 1 | CEL | B | 0 | 0 | 0 | 1 | 1 | 0 | 1 | 1 | 1 | 11 | 0 | 1 | 11 | 11 | 1 | 11 | 0 | 1 | 1 | 0 | 1 | 11 | 11 | 0 | 1 | CEL |
| 7lmc | C | 0 | 0 | 0 | 1 | 1 | 0 | 1 | 1 | 1 | 1 | 0 | 1 | 11 | 1 | 1 | 11 | 1 | 1 | 1 | 1 | 1 | 1 | 0 | 1 | 10 | CEL | D | 0 | 0 | 0 | 1 | 1 | 0 | 1 | 1 | 1 | 11 | 1 | 11 | 11 | 11 | 1 | 11 | 1 | 1 | 0 | 1 | 0 | 11 | 1 | 1 | 0 | CEL |
| 7ltj | AB | 0 | 0 | 0 | 1 | 1 | 0 | 1 | 1 | 1 | 11 | 1 | 1 | 11 | 0 | 0 | 11 | 0 | 0 | 1 | 0 | 0 | 1 | 0 | 0 | 0 | CE | | | | | | | | | | | | | | | | | | | | | | | | | | | |
| 7lyh | AB | 0 | 0 | 0 | 1 | 1 | 0 | 11 | 1 | 1 | 0 | 0 | 11 | 11 | 11 | 1 | 11 | 0 | 1 | 1 | 1 | 1 | 1 | 1 | 0 | 0 | CEL | | | | | | | | | | | | | | | | | | | | | | | | | | | |
| 7lyi | AB | 0 | 0 | 0 | 1 | 1 | 0 | 11 | 0 | 1 | 1 | 1 | 21 | 11 | 11 | 1 | 11 | 0 | 0 | 1 | 1 | 1 | 0 | 0 | 0 | 0 | CE | | | | | | | | | | | | | | | | | | | | | | | | | | | |
| 7lzt | AB | 0 | 0 | 0 | 11 | 0 | 0 | 11 | 1 | 1 | 0 | 0 | 21 | 11 | 11 | 1 | 11 | 0 | 1 | 1 | 1 | 1 | 11 | 0 | 0 | 0 | CEL | | | | | | | | | | | | | | | | | | | | | | | | | | | |
| 7lzu | A | 0 | 0 | 0 | 11 | 1 | 0 | 11 | 1 | 1 | 0 | 0 | 21 | 11 | 11 | 1 | 11 | 0 | 0 | 1 | 1 | 1 | 11 | 1 | 1 | 0 | CEHL | B | 0 | 0 | 0 | 1 | 1 | 0 | 11 | 1 | 1 | 1 | 0 | 21 | 11 | 11 | 1 | 11 | 0 | 0 | 1 | 1 | 1 | 11 | 1 | 0 | 0 | CEL |
| 7lzv | A | 0 | 0 | 0 | 11 | 0 | 0 | 11 | 1 | 1 | 0 | 0 | 21 | 11 | 11 | 1 | 11 | 0 | 0 | 1 | 1 | 0 | 11 | 1 | 1 | 0 | CEL | B | 0 | 0 | 0 | 11 | 0 | 1 | 11 | 1 | 1 | 0 | 0 | 21 | 11 | 11 | 1 | 11 | 0 | 0 | 1 | 1 | 1 | 11 | 0 | 0 | 0 | CEHL |
| 7lzw | A | 0 | 0 | 0 | 10 | 0 | 0 | 11 | 1 | 1 | 0 | 0 | 21 | 11 | 11 | 1 | 11 | 0 | 0 | 1 | 1 | 1 | 11 | 1 | 1 | 0 | CEL | B | 0 | 0 | 0 | 11 | 0 | 0 | 11 | 1 | 1 | 0 | 0 | 21 | 11 | 11 | 1 | 11 | 0 | 0 | 1 | 1 | 0 | 11 | 0 | 0 | 0 | CE |
| 7lzx | A | 0 | 0 | 0 | 1 | 0 | 0 | 11 | 1 | 1 | 1 | 0 | 21 | 11 | 11 | 1 | 11 | 0 | 1 | 1 | 1 | 0 | 11 | 0 | 0 | 0 | CE | B | 0 | 0 | 0 | 1 | 0 | 0 | 11 | 0 | 1 | 1 | 1 | 21 | 11 | 11 | 1 | 11 | 1 | 1 | 1 | 1 | 0 | 11 | 0 | 0 | 0 | CE |
| 7lzy | AB | 0 | 0 | 0 | 11 | 1 | 0 | 1 | 0 | 1 | 0 | 0 | 21 | 11 | 11 | 1 | 11 | 0 | 1 | 1 | 1 | 0 | 11 | 1 | 0 | 0 | EHL | | | | | | | | | | | | | | | | | | | | | | | | | | | |
| 7lzz | AB | 0 | 0 | 0 | 11 | 0 | 1 | 11 | 0 | 1 | 0 | 0 | 21 | 11 | 11 | 1 | 11 | 0 | 1 | 1 | 1 | 0 | 11 | 0 | 0 | 0 | CEH | | | | | | | | | | | | | | | | | | | | | | | | | | | |
| 7m00 | A | 0 | 0 | 0 | 11 | 0 | 1 | 1 | 0 | 1 | 0 | 0 | 21 | 11 | 11 | 1 | 11 | 1 | 1 | 1 | 1 | 1 | 11 | 1 | 0 | 0 | EHL | B | 0 | 0 | 0 | 11 | 0 | 1 | 1 | 0 | 1 | 0 | 0 | 21 | 11 | 11 | 1 | 11 | 1 | 1 | 1 | 1 | 1 | 11 | 1 | 0 | 0 | EHL |
| 7m01 | A | 0 | 0 | 0 | 1 | 0 | 0 | 11 | 1 | 1 | 1 | 0 | 21 | 11 | 11 | 1 | 11 | 0 | 0 | 1 | 1 | 1 | 11 | 1 | 1 | 0 | CEL | B | 0 | 0 | 0 | 11 | 1 | 0 | 11 | 0 | 1 | 0 | 1 | 21 | 11 | 11 | 1 | 11 | 0 | 0 | 1 | 1 | 1 | 11 | 1 | 0 | 0 | CEHL |
| 7m02 | A | 0 | 0 | 0 | 1 | 0 | 0 | 11 | 1 | 1 | 1 | 0 | 21 | 11 | 11 | 1 | 11 | 0 | 0 | 1 | 1 | 1 | 11 | 1 | 1 | 0 | CEL | B | 0 | 0 | 0 | 1 | 0 | 0 | 11 | 0 | 1 | 1 | 1 | 21 | 11 | 11 | 1 | 11 | 0 | 0 | 1 | 1 | 1 | 11 | 0 | 0 | 0 | CEL |
| 7m03 | A | 0 | 0 | 0 | 11 | 0 | 0 | 11 | 1 | 1 | 0 | 0 | 21 | 11 | 11 | 1 | 11 | 0 | 1 | 1 | 1 | 0 | 11 | 1 | 1 | 0 | CEL | B | 0 | 0 | 0 | 1 | 0 | 0 | 11 | 1 | 1 | 1 | 1 | 21 | 11 | 11 | 1 | 11 | 0 | 0 | 1 | 1 | 1 | 11 | 0 | 0 | 0 | CEL |
| 7m04 | A | 0 | 0 | 0 | 11 | 0 | 0 | 11 | 0 | 1 | 0 | 0 | 21 | 11 | 11 | 1 | 11 | 0 | 0 | 1 | 1 | 0 | 11 | 1 | 1 | 0 | CEL | B | 0 | 0 | 0 | 11 | 0 | 0 | 11 | 1 | 1 | 0 | 1 | 21 | 11 | 11 | 1 | 11 | 0 | 0 | 1 | 1 | 1 | 11 | 0 | 0 | 0 | CEL |
| 7m8m | A | 1 | 11 | 0 | 1 | 1 | 0 | 1 | 1 | 1 | 11 | 0 | 11 | 11 | 1 | 1 | 11 | 1 | 0 | 0 | 0 | 1 | 0 | 0 | 0 | 1 | RCE | | | | | | | | | | | | | | | | | | | | | | | | | | | |
| 7m8n | A | 1 | 11 | 0 | 1 | 1 | 0 | 1 | 1 | 1 | 11 | 0 | 11 | 11 | 0 | 1 | 11 | 1 | 1 | 0 | 0 | 0 | 1 | 1 | 0 | 1 | RCE | B | 1 | 11 | 0 | 1 | 0 | 0 | 1 | 1 | 1 | 11 | 0 | 11 | 11 | 0 | 1 | 11 | 1 | 1 | 0 | 0 | 1 | 1 | 1 | 0 | 1 | RCEL |
| 7m8o | A | 1 | 11 | 0 | 1 | 0 | 0 | 1 | 1 | 1 | 11 | 0 | 11 | 11 | 1 | 1 | 11 | 0 | 1 | 0 | 0 | 0 | 1 | 1 | 0 | 0 | RCE | | | | | | | | | | | | | | | | | | | | | | | | | | | |
| 7m8p | A | 1 | 11 | 0 | 1 | 0 | 0 | 1 | 1 | 1 | 11 | 10 | 11 | 11 | 1 | 1 | 11 | 0 | 1 | 0 | 0 | 0 | 1 | 1 | 0 | 0 | RCE | | | | | | | | | | | | | | | | | | | | | | | | | | | |
| 7m8x | AB | 1 | 1 | 0 | 1 | 1 | 0 | 1 | 1 | 1 | 1 | 1 | 1 | 11 | 1 | 1 | 11 | 1 | 0 | 0 | 0 | 1 | 1 | 1 | 0 | 1 | CEL | | | | | | | | | | | | | | | | | | | | | | | | | | | |
| 7m8y | AB | 1 | 0 | 0 | 0 | 1 | 0 | 1 | 1 | 1 | 11 | 0 | 11 | 0 | 0 | 1 | 11 | 0 | 1 | 0 | 0 | 1 | 1 | 1 | 0 | 0 | C | | | | | | | | | | | | | | | | | | | | | | | | | | | |

| PDBid | BC | T25 | T26 | L27 | H41 | M49 | Y54 | F140 | L141 | N142 | G143 | S144 | C145 | H163 | H164 | M165 | E166 | L167 | P168 | H172 | D187 | R188 | Q189 | T190 | A191 | Q192 | Ligand interactions |
|---|---|---|---|---|---|---|---|---|---|---|---|---|---|---|---|---|---|---|---|---|---|---|---|---|---|---|---|
| | MC | interface | | | H-loop | | | C-loop | | | | interface | | I&II | | E-loop | | | | | I&II | Linker | | | | | |
| 7m8z | AB | 1 | 0 | 0 | 0 | 0 | 0 | 1 | 1 | 11 | 11 | 0 | 11 | 11 | 1 | 1 | 11 | 1 | 1 | 0 | 0 | 1 | 1 | 1 | 0 | 1 | CEL |
| 7m90 | AB | 1 | 0 | 0 | 1 | 1 | 0 | 1 | 1 | 1 | 11 | 0 | 11 | 11 | 1 | 1 | 11 | 1 | 1 | 0 | 0 | 0 | 1 | 1 | 1 | 11 | CEL |
| 7m91 | AB | 1 | 0 | 0 | 1 | 0 | 0 | 1 | 1 | 11 | 11 | 0 | 11 | 11 | 0 | 1 | 11 | 0 | 0 | 0 | 0 | 0 | 1 | 0 | 0 | 1 | CE |
| 7mbi | A | 0 | 0 | 0 | 1 | 0 | 0 | 11 | 0 | 1 | 11 | 0 | 21 | 11 | 11 | 1 | 11 | 0 | 0 | 0 | 1 | 1 | 11 | 1 | 1 | 0 | CEL |
| 7mbi | B | 0 | 0 | 0 | 1 | 0 | 0 | 11 | 0 | 1 | 11 | 1 | 21 | 11 | 11 | 1 | 11 | 0 | 1 | 1 | 1 | 1 | 11 | 1 | 1 | 0 | CEL |
| 7mgr | AB | 1 | 11 | 0 | 1 | 1 | 0 | 11 | 1 | 11 | 11 | 1 | 11 | 11 | 11 | 1 | 11 | 1 | 1 | 1 | 0 | 1 | 1 | 11 | 0 | 1 | RCEL |
| 7mgs | AB | 1 | 11 | 0 | 1 | 1 | 0 | 11 | 1 | 11 | 11 | 0 | 11 | 11 | 11 | 1 | 11 | 0 | 1 | 1 | 0 | 1 | 1 | 11 | 0 | 1 | RCEL |
| 7mng | AB | 1 | 1 | 0 | 11 | 1 | 0 | 0 | 0 | 1 | 11 | 0 | 21 | 0 | 1 | 1 | 1 | 0 | 0 | 0 | 0 | 0 | 1 | 0 | 0 | 0 | CH |
| 7mpb | A | 0 | 0 | 0 | 0 | 0 | 0 | 1 | 1 | 11 | 1 | 1 | 11 | 10 | 0 | 0 | 1 | 0 | 0 | 0 | 0 | 0 | 0 | 0 | 0 | 0 | C |
| 7mrr | AB | 0 | 0 | 0 | 1 | 1 | 0 | 0 | 1 | 1 | 0 | 0 | 21 | 1 | 11 | 1 | 11 | 0 | 0 | 0 | 0 | 0 | 1 | 1 | 0 | 0 | E |
| 7n44 | AB | 1 | 11 | 0 | 1 | 1 | 0 | 1 | 1 | 1 | 11 | 0 | 11 | 11 | 1 | 1 | 11 | 1 | 1 | 0 | 0 | 1 | 1 | 1 | 1 | 1 | RCEL |
| 7n6n | A | 0 | 0 | 0 | 1 | 0 | 0 | 1 | 1 | 1 | 1 | 1 | 11 | 11 | 11 | 1 | 11 | 0 | 1 | 1 | 1 | 1 | 1 | 11 | 1 | 0 | CEL |
| 7n89 | A | 1 | 11 | 0 | 1 | 1 | 0 | 11 | 1 | 1 | 11 | 1 | 11 | 11 | 11 | 1 | 11 | 0 | 1 | 1 | 0 | 1 | 11 | 11 | 1 | 1 | RCEL |
| 7n8c | AB | 0 | 0 | 0 | 1 | 1 | 0 | 1 | 0 | 1 | 11 | 1 | 21 | 11 | 0 | 0 | 11 | 0 | 0 | 1 | 0 | 0 | 1 | 0 | 0 | 0 | CE |
| 7nbr | AB | 0 | 0 | 0 | 11 | 1 | 1 | 0 | 0 | 1 | 11 | 0 | 21 | 0 | 11 | 1 | 11 | 1 | 1 | 0 | 1 | 0 | 1 | 1 | 0 | 1 | CEHL |
| 7nbs | AB | 1 | 1 | 0 | 11 | 1 | 0 | 0 | 0 | 1 | 11 | 0 | 21 | 0 | 11 | 1 | 11 | 1 | 1 | 0 | 0 | 1 | 11 | 1 | 1 | 1 | CEHL |
| 7nby[x] | A | 0 | 0 | 0 | 1 | 1 | 1 | 0 | 0 | 0 | 0 | 0 | 21 | 0 | 1 | 1 | 0 | 0 | 0 | 0 | 1 | 11 | 11 | 0 | 0 | 0 | HL |
| 7nf5 | AB | 0 | 0 | 0 | 1 | 1 | 0 | 1 | 0 | 1 | 1 | 0 | 21 | 1 | 11 | 1 | 11 | 0 | 1 | 0 | 1 | 0 | 11 | 1 | 1 | 1 | EL |
| 7ng3 | A | 0 | 0 | 0 | 1 | 1 | 0 | 1 | 0 | 0 | 1 | 0 | 21 | 1 | 11 | 1 | 11 | 1 | 1 | 0 | 1 | 0 | 11 | 1 | 1 | 1 | EL |
| 7ng6 | A | 0 | 0 | 0 | 1 | 0 | 0 | 1 | 0 | 0 | 1 | 0 | 21 | 0 | 11 | 1 | 11 | 1 | 0 | 0 | 1 | 1 | 11 | 1 | 1 | 1 | EL |
| 7nt1 | | | | | | | | | | | | | | | | | | | | | | | | | | | |
| 7nt2 | | | | | | | | | | | | | | | | | | | | | | | | | | | |
| 7nt3 | | | | | | | | | | | | | | | | | | | | | | | | | | | |
| 7ntv | | | | | | | | | | | | | | | | | | | | | | | | | | | |
| 7nuk | | | | | | | | | | | | | | | | | | | | | | | | | | | |
| 7nw2 | A | 0 | 0 | 0 | 1 | 1 | 0 | 1 | 1 | 1 | 10 | 11 | 21 | 11 | 1 | 1 | 11 | 0 | 0 | 0 | 1 | 1 | 1 | 0 | 0 | 1 | CEL |
| 7p35 | A | 1 | 1 | 0 | 1 | 1 | 0 | 11 | 1 | 1 | 11 | 1 | 21 | 11 | 0 | 1 | 11 | 0 | 1 | 1 | 1 | 1 | 1 | 11 | 1 | 0 | CEL |

| PDBid | BC | T25 | T26 | L27 | H41 | M49 | Y54 | F140 | L141 | N142 | G143 | S144 | C145 | H163 | H164 | M165 | E166 | L167 | P168 | H172 | D187 | R188 | Q189 | T190 | A191 | Q192 | Ligand interactions |
|---|---|---|---|---|---|---|---|---|---|---|---|---|---|---|---|---|---|---|---|---|---|---|---|---|---|---|---|
| | MC | interface | | | H-loop | | | C-loop | | | | interface | | I&II | | E-loop | | | | | I&II | Linker | | | | | |
| 7m8z | | | | | | | | | | | | | | | | | | | | | | | | | | | |
| 7m90 | | | | | | | | | | | | | | | | | | | | | | | | | | | |
| 7m91 | | | | | | | | | | | | | | | | | | | | | | | | | | | |
| 7mbi | D | 0 | 0 | 0 | 1 | 0 | 0 | 11 | 0 | 1 | 11 | 0 | 21 | 11 | 11 | 1 | 11 | 0 | 0 | 1 | 1 | 1 | 1 | 1 | 1 | 0 | CEL |
| 7mbi | C | 0 | 0 | 0 | 1 | 0 | 0 | 11 | 1 | 0 | 1 | 0 | 21 | 11 | 11 | 1 | 11 | 0 | 0 | 1 | 1 | 1 | 11 | 1 | 1 | 0 | CEL |
| 7mgr | | | | | | | | | | | | | | | | | | | | | | | | | | | |
| 7mgs | | | | | | | | | | | | | | | | | | | | | | | | | | | |
| 7mng | | | | | | | | | | | | | | | | | | | | | | | | | | | |
| 7mpb | B | 0 | 0 | 0 | 0 | 0 | 0 | 1 | 1 | 1 | 1 | 0 | 1 | 11 | 0 | 0 | 1 | 0 | 0 | 1 | 0 | 0 | 0 | 0 | 0 | 0 | C |
| 7mrr | | | | | | | | | | | | | | | | | | | | | | | | | | | |
| 7n44 | | | | | | | | | | | | | | | | | | | | | | | | | | | |
| 7n6n | B | 0 | 0 | 0 | 1 | 1 | 0 | 1 | 1 | 1 | 1 | 1 | 1 | 1 | 0 | 1 | 11 | 0 | 1 | 1 | 1 | 1 | 1 | 0 | 1 | 0 | CEL |
| 7n89 | B | 1 | 11 | 0 | 1 | 1 | 0 | 1 | 1 | 1 | 11 | 0 | 11 | 11 | 11 | 1 | 11 | 0 | 1 | 0 | 1 | 1 | 11 | 11 | 0 | 1 | RCEL |
| 7n8c | | | | | | | | | | | | | | | | | | | | | | | | | | | |
| 7nbr | | | | | | | | | | | | | | | | | | | | | | | | | | | |
| 7nbs | | | | | | | | | | | | | | | | | | | | | | | | | | | |
| 7nby[x] | B | 0 | 0 | 0 | 1 | 0 | 1 | 0 | 0 | 0 | 0 | 0 | 21 | 0 | 11 | 1 | 0 | 0 | 0 | 0 | 1 | 1 | 0 | 0 | 0 | 0 | |
| 7nf5 | | | | | | | | | | | | | | | | | | | | | | | | | | | |
| 7ng3 | B | 0 | 0 | 0 | 1 | 0 | 0 | 1 | 0 | 1 | 1 | 0 | 21 | 1 | 11 | 1 | 11 | 1 | 1 | 0 | 1 | 0 | 11 | 1 | 1 | 1 | EL |
| 7ng6 | B | 0 | 0 | 0 | 1 | 0 | 0 | 1 | 0 | 1 | 1 | 0 | 21 | 1 | 11 | 1 | 10 | 0 | 1 | 0 | 1 | 0 | 11 | 1 | 1 | 1 | EL |
| 7nt1 | B | 0 | 0 | 0 | 1 | 1 | 0 | 1 | 0 | 1 | 1 | 1 | 21 | 11 | 0 | 1 | 1 | 0 | 0 | 0 | 0 | 0 | 1 | 0 | 0 | 0 | C |
| 7nt2 | B | 0 | 0 | 0 | 1 | 1 | 0 | 0 | 0 | 1 | 1 | 0 | 21 | 0 | 1 | 1 | 11 | 0 | 0 | 0 | 1 | 1 | 1 | 0 | 0 | 0 | |
| 7nt3 | B | 0 | 0 | 0 | 1 | 1 | 1 | 1 | 1 | 1 | 0 | 0 | 0 | 11 | 1 | 1 | 11 | 0 | 0 | 0 | 1 | 1 | 1 | 0 | 0 | 0 | EH |
| 7ntv | B | 1 | 1 | 0 | 1 | 1 | 0 | 0 | 0 | 1 | 1 | 0 | 21 | 0 | 0 | 0 | 0 | 0 | 0 | 0 | 0 | 0 | 0 | 0 | 0 | 0 | |
| 7nuk | B | 0 | 1 | 0 | 1 | 0 | 0 | 0 | 0 | 0 | 11 | 0 | 21 | 0 | 0 | 0 | 0 | 0 | 0 | 0 | 0 | 0 | 0 | 0 | 0 | 0 | |
| 7nw2 | | | | | | | | | | | | | | | | | | | | | | | | | | | |
| 7p35 | B | 0 | 1 | 1 | 1 | 1 | 0 | 11 | 1 | 0 | 11 | 1 | 21 | 11 | 10 | 1 | 11 | 0 | 1 | 1 | 1 | 1 | 1 | 1 | 1 | 0 | CEL |

PDB data are between 10/25/2020 of Tables 1&2 and 7/25/2021. The meaning of the table is the same as in Table S4. The minor 14 binding sites other than the extraordinary binding sites (Table shown below) are Lys5(1), Gly15(1), Thr24(6), Asn28(1), Pro39(3), Cys44(13), Thr45(3), Ser46(6), Ala70(1), Lys97(1), Gly146(1), Ser147(1), Gln256(1), Ser301(2), where the numbers of the parentheses are the number of contacts.

[#]the inter-protomer contact.

The ligand binding at sites outside the major binding sites (the entry name with x).

| | | | | | | | | | | | |
|---|---|---|---|---|---|---|---|---|---|---|---|
| 7abu | A:401 | | B:141,142 A: 252,253,256,297 | 7axm | A:502 | | B:118,141,142 A:213,252,253,256,296,297,300,301 | 7nby | A:401 | | A: 41,44,49,54,187,188,189 |
| 7af0 | A:504 | | A:273,274 | 7axo | A:506 | | A:237,271,272,273,274,275,276,285,287 | | A:402 | | A: 1,2,3,210,213,214,300 |
| 7aga | A:401 | | A:107,108,109,110,153,202,203,249,292,293,294,298 | 7bfb | A:401 | | A:44,45,49,187,189 | | A:403 | | A: 41,145,164,165,189 |
| 7amj | A:409 | | B:142 A:213,253,256,297,300,301 | | A:402 | | A:143,145 | | A:406 | | A: 100,101,102,156 |
| 7aol: | A:403 | | A:103,104,105,176,178 | | A403 | | A:100,101,102,156 | | B:406 | | B: 44,53,54,57,188 |
| 7aph | A:401 | | A:252,253,255,256,297,300,301 | | B:1002 | | B:100,102,156 | | B:401 | | B: 1,2,141,213,214,300 |
| 7aqi | A:401 | | A:252,253,256,297,300 | | B:1001 | | B:256,297,300,301 | | B:402 | | B: 39,41,145,164,165,187 |
| 7arf | A:401 | | A:100,101,155,156 | 7kvl | A:503 | | A:233,237,269,273 | | B:403 | | B: 101,156 |
| 7awr | A:401 | | A:33,99,100,101,102 | 7kvr | A:703 | | A:83,84 | | | | |
| 7aww | A:401 | | A:33,34,94,95,98 | 7ldx | A:509 | | A:132,133,196,198,235,239,240,241 | | | | |
| 7ax6 | A:401 | | A:100,101,102,156 | | B:402 | | B:132,133,196,198,235,239,240,241 | | | | |

Residue number without underline is for nonpolar contact, with single underline for polar contact, and with double underline for covalent bond.

**Supporting data S7** Summary of entries of MERS-CoV 3CL protease

| ligand | protein | | | | ligand | | | | comments | Chain A | | | | | | | | | Chain B | | | | | | | | | |
|---|---|---|---|---|---|---|---|---|---|---|---|---|---|---|---|---|---|---|---|---|---|---|---|---|---|---|---|---|
| | | | | | | | | | | C-loop conformation | | | | | | | | | C-loop conformation | | | | | | | | | |
| | PDB ID | mutation | residues added to N-term | missing N-terminal residues | name in PDB | name by authors | peptide or peptide mimic$ | Ki, $IC_{50}$*, $EC_{50}^{+}$ (μM) | | main-chain HB within 141-147 (138-145)# | HB 141-175 (138-172)# | HB 142-129 (139-126)# | HB 143-1 (140-1)# | HB 144-121 (141-118)# | HB 146-28 (143-28)# | distance 168Cα-143Cβ (165-140)# | distance 168Cα-144Cβ (165-141)# | HB 2-217 (2-214)# | main-chain HB within 141-147 (138-145)# | HB 141-175 (138-172)# | HB 142-129 (139-126)# | HB 143-1 (140-1)# | HB 144-121 (141-118)# | HB 146-28 (143-28)# | distance 168Cα-143Cβ (165-140)# | distance 168Cα-144Cβ (165-141)# | HB 2-217 (2-214)# | distance between two 285 Cα's |
| ligand bound | 4rsp | | | | PRD_002 174 | 6 | p | 3.6 | C145 is CSO and does not bind the ligand | 144O-147N | 1 | 0 | 1 | 1 | 1 | 7.414 | 10.957 | 1 | | | | | | | | | | 6.123 |
| | 4wme_AB | C148A | | | | C-term of another molecule | p | | chain BD are ligand-free. A(C) bind C-term of C(A) | 144O-147N | 1 | 0 | 1 | 1 | 1 | 7.485 | 10.867 | 1 | 144O-147N | 1 | 0 | 1 | 1 | 1 | 7.556 | 10.982 | 1 | 6.153 |
| | 4wmf_A | C148A | | | | C-term of another molecule | p | | chain BC are ligand free A binds C-term of C | 144O-147N | 1 | 0 | 1 | 1 | 1 | 7.294 | 10.603 | 1 | | | | | | | | | | 5.956 |
| | 4ylu_AB | | | | R30 | 11 | | >100* | | 144O-147N | 1 | 0 | 1 | 1 | 1 | 7.5 | 11.073 | 1 | 144O-147N | 1 | 0 | 1 | 1 | 1 | 7.549 | 11.173 | 1 | 7.174 |
| | 4ylu_CD | | | | R30 | 11 | | >100* | | 144O-147N | 1 | 0 | 1 | 1 | 1 | 7.589 | 11.175 | 1 | 144O-147N | 1 | 0 | 1 | 1 | 1 | 7.475 | 10.935 | 1 | 6.132 |
| | 5wkj | | MH6 | MH5 | K36,B1S | GC376 | p | | K36 and,B1S are stereoisomers, and not distinguished | 144O-147N | 1 | 0 | 0& | 1 | 1 | 7.225 | 10.955 | 0 | | | | | | | | | | 6.762 |
| | 5wkk | | MH6 | MH4 | B3G,AW4 | GC813 | p | | B3G and AW4 are stereoisomers, and not distinguished | 144O-147N | 1 | 0 | 0& | 1 | 1 | 7.278 | 10.744 | 0 | | | | | | | | | | 6.603 |
| | 5wkl | | MH6 | MH4 | B3J, AVY | 10c | p | 0.7* | B3J and AVY are stereoisomers, and not distinguished | 144O-147N | 1 | 0 | 0& | 1 | 1 | 7.332 | 10.946 | 0 | | | | | | | | | | 7.26 |
| | 5wkm | | MH6 | MH4 | B6Y, N02 | 10e | | 7.5* | B6Y and N02 are stereoisomers, and not distinguished | 144O-147N | 1 | 0 | 0& | 1 | 1 | 7.3 | 11.024 | 0 | | | | | | | | | | 7.636 |
| | 6vgy | | MH6 | MH4 of A,B | QZJ | 6b | | 0.33* | | 144O-147N | 0$ | 0 | 0& | 1 | 1 | 7.298 | 11.006 | 0 | 144O-147N | 0$ | 0 | 0& | 1 | 1 | 7.356 | 11.099 | 0 | 9.495 |
| | 6vgz | | MH6 | MH4 of A,B | QZG | 6d | | 0.41* | | 144O-147N | 0$ | 0 | 0& | 1 | 1 | 7.3 | 11.061 | 0 | 144O-147N | 0$ | 0 | 0& | 1 | 1 | 7.348 | 11.181 | 0 | 9.624 |
| | 6vh0 | | MH6 | MH4 of A,B | QZD | 6g | | 0.12* | | 144O-147N | 0$ | 0 | 0& | 1 | 1 | 7.193 | 10.862 | 0 | 144O-147N | 0$ | 0 | 0& | 1 | 1 | 7.352 | 11.144 | 0 | 9.289 |
| | 6vh1 | | MH6 | MH4 of A,B | QZ7 | 6h | | 0.07* | | 144O-147N | 0$ | 0 | 0& | 1 | 1 | 7.284 | 11.074 | 0 | 144O-147N | 0$ | 0 | 0& | 1 | 1 | 7.287 | 11.067 | 0 | 9.842 |
| | 6vh2 | | MH6 | MH4 of A,B | QZ4 | 7i | | 1.9* | | 144O-147N | 0$ | 0 | 0& | 1 | 1 | 7.317 | 11.066 | 0 | 144O-147N | 0$ | 0 | 0& | 1 | 1 | 7.434 | 11.123 | 0 | 9.758 |
| | 6vh3 | | MH6 | MH4 of A, MH6 of B | QYS | 7j | p | 0.1* | | 144O-147N | 0$ | 0 | 0& | 1 | 1 | 7.486 | 11.369 | 0 | 144O-147N | 0$ | 0 | 0 | 1 | 1 | 7.32 | 11.132 | 0 | 10.103 |
| ligand free | 4wmd_AB | C148A | | | | | | | | 144O-147N | 1 | 0 | 1 | 1 | 1 | 7.238 | 10.959 | 0 | 144O-147N | 1 | 0 | 1 | 1 | 1 | 7.277 | 10.989 | 1 | 6.293 |
| | 4wmd_C | C148A | | | | | | | | 144O-147N | 1 | 0 | 1 | 1 | 1 | 7.215 | 10.962 | 1 | | | | | | | | | | 6.116 |
| | 4wme_CD | C148A | | | | | | | chain AD are ligand bound | 144O-147N | 1 | 0 | 1 | 1 | 1 | 7.426 | 10.866 | 1 | 144O-147N | 1 | 0 | 1 | 1 | 1 | 7.4 | 10.817 | 1 | 6.191 |
| | 4wmf_B | C148A | | | | | | | chain A is ligand bound | | | | | | | | | | 144O-147N | 1 | 0 | 1 | 1 | 1 | 7.215 | 10.755 | 1 | 5.956 |
| | 4wmf_C | C148A | | 1-10 of C | | | | | chain C is monomeric. Its core is maintained, but domain III rotates about 180 deg. | 144O-147N | 0 | 0 | 0 | 1 | 1 | 8.071 | 11.316 | 0 | | | | | | | | | | |
| | 5c3n | | | | | | | | | 144O-147N | 0 | 0 | 1 | 1 | 1 | 7.87 | 11.145 | 0 | 144O-147N | 0 | 0 | 1 | 1 | 1 | 7.571 | 11.064 | 1 | 6.581 |

### The residue numbers in parenthesis is the corresponding number for SARS-CoV-2 $3CL^{pro}$.

$ HB 141-175 is replaced by HB 140-175.

& HB 143-1 is replaced by HB 142-2.

**Table S8** Ligand binding at protein residues of MERS-CoV 3CL protease.

| | | interfa ce | H-loop | | | C-loop | | | | | | E-loop | | | | | | | Linker | | | | | | Ligand interactions |
|---|---|---|---|---|---|---|---|---|---|---|---|---|---|---|---|---|---|---|---|---|---|---|---|---|---|
| entry | chain | M25 | H41 | L49 | Y54 | F143 | L144 | C145 | G146 | S147 | C148 | H166 | Q167 | M168 | E169 | L170 | A171 | H175 | D190 | K191 | Q192 | V193 | H194 | Q195 | |
| 4rsp | A | 0 | 1 | 0 | 0 | 11 | 1 | 1 | 10 | 1 | 21 | 11 | 11 | 1 | 11 | 0 | 0 | 1 | 1 | 10 | 11 | 11 | 1 | 11 | CEL |
| 4wme_AB | A | 0 | 11 | 0 | 0 | 11 | 1 | 1 | 11 | 1 | 11 | 11 | 11 | 1 | 11 | 1 | 0 | 1 | 1 | 1 | 11 | 11 | 1 | 1 | CEL |
| 4wme_CD | D | 0 | 11 | 0 | 0 | 11 | 1 | 0 | 11 | 0 | 11 | 11 | 11 | 1 | 11 | 1 | 0 | 1 | 1 | 1 | 1 | 1 | 1 | 0 | CEL |
| 4wmf_AB | A | 0 | 11 | 0 | 0 | 11 | 1 | 1 | 11 | 1 | 11 | 11 | 11 | 1 | 11 | 1 | 1 | 1 | 1 | 1 | 11 | 1 | 11 | 0 | CEL |
| 4ylu_AB | A | 1 | 1 | 1 | 1 | 1 | 1 | 1 | 0 | 0 | 11 | 11 | 0 | 1 | 11 | 0 | 0 | 0 | 1 | 1 | 1 | 0 | 0 | 0 | EH |
| 4ylu_AB | B | 1 | 1 | 1 | 0 | 1 | 1 | 1 | 0 | 0 | 1 | 11 | 0 | 1 | 11 | 0 | 0 | 0 | 1 | 1 | 1 | 0 | 0 | 0 | E |
| 4ylu_CD | C | 1 | 1 | 1 | 0 | 1 | 1 | 1 | 0 | 0 | 1 | 11 | 0 | 1 | 11 | 0 | 0 | 0 | 1 | 1 | 1 | 0 | 0 | 0 | EH |
| 4ylu_CD | D | 1 | 1 | 1 | 1 | 1 | 1 | 1 | 0 | 0 | 11 | 11 | 0 | 1 | 11 | 0 | 0 | 0 | 1 | 1 | 1 | 0 | 0 | 0 | EH |
| 5wkj | A | 0 | 11 | 1 | 0 | 11 | 1 | 1 | 0 | 0 | 21 | 11 | 11 | 1 | 11 | 0 | 0 | 1 | 1 | 0 | 11 | 0 | 0 | 0 | CE |
| 5wkk | A | 0 | 1 | 1 | 0 | 11 | 1 | 1 | 1 | 1 | 21 | 11 | 11 | 1 | 11 | 0 | 1 | 1 | 1 | 0 | 11 | 1 | 1 | 0 | CEL |
| 5wkl | A | 0 | 11 | 1 | 1 | 11 | 1 | 1 | 0 | 1 | 21 | 11 | 11 | 1 | 11 | 0 | 0 | 1 | 1 | 0 | 11 | 0 | 0 | 0 | CEH |
| 5wkm | A | 0 | 0 | 0 | 0 | 11 | 0 | 1 | 1 | 1 | 21 | 11 | 11 | 1 | 11 | 0 | 0 | 1 | 1 | 1 | 0 | 0 | 0 | 0 | CE |
| 6vgy | A | 0 | 1 | 1 | 0 | 1 | 1 | 1 | 1 | 1 | 21 | 11 | 11 | 1 | 11 | 0 | 1 | 1 | 1 | 0 | 11 | 0 | 0 | 0 | CE |
| | B | 0 | 0 | 1 | 0 | 11 | 0 | 0 | 1 | 1 | 21 | 11 | 11 | 1 | 11 | 1 | 1 | 1 | 1 | 1 | 11 | 0 | 0 | 0 | CEL |
| 6vgz | A | 0 | 1 | 0 | 0 | 11 | 0 | 1 | 1 | 0 | 21 | 11 | 11 | 1 | 11 | 0 | 1 | 1 | 1 | 1 | 11 | 0 | 0 | 0 | CEL |
| | B | 0 | 0 | 0 | 0 | 11 | 0 | 0 | 1 | 0 | 21 | 11 | 11 | 1 | 11 | 0 | 0 | 1 | 1 | 1 | 11 | 0 | 0 | 0 | CEL |
| 6vh0 | A | 0 | 1 | 0 | 0 | 11 | 1 | 1 | 1 | 1 | 21 | 11 | 11 | 1 | 11 | 0 | 0 | 1 | 1 | 1 | 11 | 0 | 0 | 0 | CEL |
| | B | 0 | 0 | 1 | 0 | 11 | 0 | 0 | 1 | 1 | 21 | 11 | 11 | 1 | 11 | 0 | 0 | 1 | 1 | 1 | 11 | 0 | 0 | 0 | CEL |
| 6vh1 | A | 0 | 1 | 1 | 0 | 11 | 0 | 0 | 1 | 1 | 21 | 11 | 11 | 1 | 11 | 0 | 0 | 1 | 0 | 0 | 11 | 0 | 0 | 0 | CE |
| | B | 0 | 0 | 1 | 0 | 11 | 0 | 0 | 1 | 1 | 21 | 11 | 11 | 1 | 11 | 0 | 0 | 1 | 1 | 0 | 11 | 0 | 0 | 0 | CE |
| 6vh2 | A | 0 | 1 | 1 | 1 | 11 | 1 | 1 | 1 | 1 | 21 | 11 | 10 | 1 | 11 | 0 | 0 | 1 | 1 | 1 | 11 | 1 | 0 | 11 | CELH |
| | B | 0 | 1 | 1 | 1 | 11 | 0 | 1 | 1 | 1 | 21 | 11 | 11 | 1 | 11 | 0 | 0 | 1 | 1 | 1 | 11 | 0 | 0 | 0 | CELH |
| 6vh3 | A | 0 | 1 | 0 | 0 | 11 | 0 | 0 | 1 | 1 | 21 | 11 | 11 | 1 | 11 | 1 | 1 | 1 | 1 | 1 | 11 | 1 | 0 | 11 | CEL |
| | B | 0 | 1 | 0 | 0 | 11 | 0 | 0 | 1 | 0 | 21 | 11 | 11 | 1 | 11 | 0 | 1 | 1 | 1 | 0 | 11 | 1 | 1 | 0 | CEL |
| MERS-CoV | | M25 | H41 | L49 | Y54 | F143 | L144 | C145 | G146 | S147 | C148 | H166 | Q167 | M168 | E169 | L170 | A171 | H175 | D190 | K191 | Q192 | V193 | H194 | Q195 | |
| ratio of occurrence | | 0.17 | 0.79 | 0.58 | 0.21 | 1.00 | 0.58 | 0.67 | 0.75 | 0.63 | 1.00 | 1.00 | 0.83 | 1.00 | 1.00 | 0.21 | 0.29 | 0.83 | 0.96 | 0.71 | 0.96 | 0.33 | 0.25 | 0.17 | |
| SARS-CoV-2 | | T25 | H41 | M49 | Y54 | F140 | L141 | N142 | G143 | S144 | C145 | H163 | H164 | M165 | E166 | L167 | P168 | H172 | D187 | R188 | Q189 | T190 | A191 | Q192 | |
| ratio of occurrence | | 0.25 | 0.94 | 0.66 | 0.18 | 0.72 | 0.65 | 0.80 | 0.83 | 0.49 | 0.99 | 0.74 | 0.88 | 0.89 | 0.90 | 0.22 | 0.47 | 0.56 | 0.68 | 0.54 | 0.80 | 0.56 | 0.38 | 0.43 | |

The contacts occurring in only one chain, L27, C44, S46 and A193, are not listed in the table.
